## Supplementary material for "Temporal Profile of Transporter mRNA Expression in the Brain after Traumatic Brain Injury in Developing Rats": Archive containing raw data, r functions, and manuscript (Rmd): analyses.R

```
# Prepare Environment
require(ggplot2)
source("compare_table.R")

#setwd() 

# Load Data
expression.data <- read.csv("all_tissues.csv", header = T, sep=",")
genes <- read.csv("genes.csv", sep=",", header=F, as.is = T)
times <- c(3,12,24,72,168,336)

# Generate Comparisons (Injury)

hippocampus.injury.data <- CompareGroupsInjury(expression.data, 
                                         times = times, 
                                         genes = genes, 
                                         tissue = "hippocampus", 
                                         group1 = "CCI", 
                                         group2 = "Sham")

cortex.injury.data <- CompareGroupsInjury(expression.data, 
                                         times = times, 
                                         genes = genes, 
                                         tissue = "cortex", 
                                         group1 = "CCI", 
                                         group2 = "Sham")

# Generate Comparisons (Naive)

cortex.naive.data <- CompareGroupsNaive(expression.data, 
                                        genes = genes, 
                                        times = c(336, 1000), 
                                        tissue =  "cortex", 
                                        base_time = 0)

hippocampus.naive.data <- CompareGroupsNaive(expression.data, 
                                        genes = genes, 
                                        times = c(336, 1000), 
                                        tissue =  "hippocampus", 
                                        base_time = 0)

CP.naive.data <- CompareGroupsNaive(expression.data, 
                                             genes = genes, 
                                             times = c(336), 
                                             tissue =  "CP", 
                                             base_time = 0)

liver.naive.data <- CompareGroupsNaive(expression.data, 
                                    genes = genes, 
                                    times = c(336), 
                                    tissue =  "liver", 
                                    base_time = 0)

kidney.naive.data <- CompareGroupsNaive(expression.data, 
                                       genes = genes, 
                                       times = c(336), 
                                       tissue =  "kidney", 
                                       base_time = 0)

# # Visualize Data
# 
# cortex.injury.plot <- ggplot(
#   data=cortex.injury.data, 
#   aes(
#     x=as.numeric(as.character(Time)), 
#     y=as.numeric(as.character(FC))))
# 
# cortex.injury.plot + 
#   facet_wrap(Gene~., nrow = 6) +
#   geom_line() + geom_point()
```
