## Supplementary material for "Temporal Profile of Transporter mRNA Expression in the Brain after Traumatic Brain Injury in Developing Rats": Archive containing raw data, r functions, and manuscript (Rmd): compare_table.R

```
CompareGroupsInjury <- function(dataset, times, genes, tissue, group1="CCI", group2="Sham") {
  # data: long data frame (defined previously)
  # times: vector of integer time points for comparison
  # genes: matrix - gene and group
  # tissue: string (what tissue for comparison?)
  # group1: numerator (main comparator)
  # group2: denominator (relative comparator)
  
  # This requires dplyr
  require(dplyr)
  
  # Define nested function
  MakeRow <- function(time, gene, gene.group) {
    # time: time point
    # gene: gene for comparison
    data1 <- dataset %>%
      filter(Group == "CCI" & Time == time & Gene == gene & Tissue == tissue)
    data2 <- dataset %>%
      filter(Group == "Sham" & Time == time & Gene == gene & Tissue == tissue)
    pval <- t.test(data1$Count, data2$Count)$p.value #T test p value
    fc <- mean(data1$Count) - mean(data2$Count) #FC (note that values for count need to be log2 transformed). 
                                    # Log2(x) - Log2(y) = Log2(x/y)
    fcsd <- sqrt(abs(sd(data1$Count)^2 - sd(data2$Count)^2))
    return(c("FC" = fc, "SD" = fcsd, "P" = pval, "Gene" = gene, "GeneGroup" = gene.group, "Time" = time, "Tissue" = tissue))
  }
  # Make matrix of gene + time comparators
  comparator <- cbind(rep(times, length(genes)), rep(genes$V1, each=length(times)), rep(genes$V2, each=length(times)))
  
  # Transpose, convert to df
  comparator.data <- data.frame(t(mapply(MakeRow, as.numeric(comparator[,1]), comparator[,2], comparator[,3])))
  
  # Add FDR and return
  comparator.data$FDR <- p.adjust(as.numeric(as.character(comparator.data$P)), method="fdr")
  return(comparator.data)
}

CompareGroupsNaive <- function(dataset, genes, times, tissue, base_time = 0) {
  # data: long data frame (defined previously)
  # genes: matrix - gene and group
  # times: vector of integer times to be compared to base_time
  # tissue: string (what tissue for comparison?)
  # base_time: time for comparison
  
  
  # This requires dplyr
  require(dplyr)
  
  # Define nested function
  MakeRow <- function(time, gene, gene.group) {
    # time: time point
    # gene: gene for comparison
    data1 <- dataset %>%
      filter(Group == "Naive" & Gene == gene & Tissue == tissue & Time == base_time)
    data2 <- dataset %>%
      filter(Group == "Naive" & Gene == gene & Tissue == tissue & Time == time)
    

    pval <- t.test(data1$Count, data2$Count)$p.value #T test p value
    fc <- mean(data2$Count) - mean(data1$Count) #FC (note that values for count need to be log2 transformed). 
                                                # Log2(x) - Log2(y) = Log2(x/y)
                                                # Reversed in this function! (All relative to T0)
    fcsd <- sqrt(abs(sd(data2$Count)^2 - sd(data1$Count)^2))
    return(c("FC" = fc, "SD" = fcsd, "P" = pval, "Gene" = gene, "GeneGroup" = gene.group, "Time" = time, "Tissue" = tissue))
    
  }
  
  # Make matrix of gene + time comparators
  comparator <- cbind(rep(times, length(genes)), rep(genes$V1, each=length(times)), rep(genes$V2, each=length(times)))
  
  # Transpose, convert to df
  comparator.data <- data.frame(t(mapply(MakeRow, as.numeric(comparator[,1]), comparator[,2], comparator[,3])))
  
  # Add FDR and return
  comparator.data$FDR <- p.adjust(as.numeric(as.character(comparator.data$P)), method="fdr")
  return(comparator.data)
  
}
```
