## Supplementary material for "Temporal Profile of Transporter mRNA Expression in the Brain after Traumatic Brain Injury in Developing Rats": Archive containing raw data, r functions, and manuscript (Rmd): functions.R

```
CompareGroupsInjury <- function(dataset, times, genes, tissue, group1="CCI", group2="Sham") {
  # data: long data frame (defined previously)
  # times: vector of integer time points for comparison
  # genes: matrix - gene and group
  # tissue: string (what tissue for comparison?)
  # group1: numerator (main comparator)
  # group2: denominator (relative comparator)
  
  # This requires dplyr
  require(dplyr)
  
  # Define nested function
  MakeRow <- function(time, gene, gene.group) {
    # time: time point
    # gene: gene for comparison
    data1 <- dataset %>%
      filter(Group == "CCI" & Time == time & Gene == gene & Tissue == tissue)
    data2 <- dataset %>%
      filter(Group == "Sham" & Time == time & Gene == gene & Tissue == tissue)
    pval <- t.test(data1$Count, data2$Count)$p.value #T test p value
    fc <- mean(data1$Count) - mean(data2$Count) #FC (note that values for count need to be log2 transformed). 
                                    # Log2(x) - Log2(y) = Log2(x/y)
    fcsd <- sqrt(abs(sd(data1$Count)^2 - sd(data2$Count)^2))
    return(c("FC" = fc, "FCSD" = fcsd, "P" = pval, "Gene" = gene, "GeneGroup" = gene.group, "Time" = time, "Tissue" = tissue))
  }
  # Make matrix of gene + time comparators
  comparator <- cbind(rep(times, length(genes)), rep(genes$V1, each=length(times)), rep(genes$V2, each=length(times)))
  
  # Transpose, convert to df
  comparator.data <- data.frame(t(mapply(MakeRow, as.numeric(comparator[,1]), comparator[,2], comparator[,3])))
  
  # Add FDR and return
  comparator.data$FDR <- p.adjust(as.numeric(as.character(comparator.data$P)), method="fdr")
  
  comparator.data[is.nan(comparator.data$FDR),]$FDR <- 1
  
  return(comparator.data)
}

CompareGroupsNaive <- function(dataset, genes, times, tissue, base_time = 0) {
  # data: long data frame (defined previously)
  # genes: matrix - gene and group
  # times: vector of integer times to be compared to base_time
  # tissue: string (what tissue for comparison?)
  # base_time: time for comparison
  
  
  # This requires dplyr
  require(dplyr)
  
  # Define nested function
  MakeRow <- function(time, gene, gene.group) {
    # time: time point
    # gene: gene for comparison
    data1 <- dataset %>%
      filter(Group == "Naive" & Gene == gene & Tissue == tissue & Time == base_time)
    data2 <- dataset %>%
      filter(Group == "Naive" & Gene == gene & Tissue == tissue & Time == time)
    

    pval <- t.test(data1$Count, data2$Count)$p.value #T test p value
    fc <- mean(data2$Count) - mean(data1$Count) #FC (note that values for count need to be log2 transformed). 
                                                # Log2(x) - Log2(y) = Log2(x/y)
                                                # Reversed in this function! (All relative to T0)
    fcsd <- sqrt(abs(sd(data2$Count)^2 - sd(data1$Count)^2))
    mean_ <- mean(data2$Count)
    sd_ <- sd(data2$Count)
    return(c("FC" = fc, "FCSD" = fcsd, "P" = pval, "Gene" = gene, "GeneGroup" = gene.group, "Time" = time, "Tissue" = tissue, "Mean" = mean_, "SD" = sd_))
    
  }
  
  # Make matrix of gene + time comparators
  comparator <- cbind(rep(times, length(genes)), rep(genes$V1, each=length(times)), rep(genes$V2, each=length(times)))
  
  # Transpose, convert to df
  comparator.data <- data.frame(t(mapply(MakeRow, as.numeric(comparator[,1]), comparator[,2], comparator[,3])))
  
  # Add FDR and return
  comparator.data$FDR <- p.adjust(as.numeric(as.character(comparator.data$P)), method="fdr")
  
  comparator.data[is.nan(comparator.data$FDR),]$FDR <- 1
  return(comparator.data)
  
}

UpDownArrow <- function(FC1, FC2, FC3, FC4, FC5, FC6, FDR1, FDR2, FDR3, FDR4, FDR5, FDR6) {
  
  # Takes the gene name plus FC and FDR, returns the gene name with an up, down, or sideways arrow
  # FC and FDR are both vectors, GENE is a string
  
  fdrc <- cbind(c(FC1, FC2, FC3, FC4, FC5, FC6), c(FDR1, FDR2, FDR3, FDR4, FDR5, FDR6))
  
  if (sum(fdrc[c(1,2,3),2] < 0.05) > 0) {
    if (mean(fdrc[c(1,2,3),1]) > 0) {
      pattern <- "↑"
    } else {
      pattern <- "↓"
    }
  } else {
    pattern <- "↔"
  }
  if (sum(fdrc[4,2] < 0.05) > 0) {
    if (mean(fdrc[4,1]) > 0) {
      pattern <- paste(pattern, "↑")
    } else {
      pattern <- paste(pattern,"↓")
    }
  } else {
    pattern <- paste(pattern,"↔")
  }
  if (sum(fdrc[c(5,6),2] < 0.05) > 0) {
    if (mean(fdrc[c(5,6),1]) > 0) {
      pattern <- paste(pattern, "↑")
    } else {
      pattern <- paste(pattern,"↓")
    }
  } else {
    pattern <- paste(pattern,"↔")
  }
}

InjuryTable <- function(df) {
    require(data.table)
    require(reshape2)
    require(tibble)
    time_order <- c("3_FC", "3_FCSD", "3_FDR", 
                    "12_FC", "12_FCSD", "12_FDR", 
                    "24_FC", "24_FCSD", "24_FDR", 
                    "72_FC", "72_FCSD", "72_FDR",
                    "168_FC", "168_FCSD", "168_FDR",
                    "336_FC", "336_FCSD", "336_FDR")
    wide.df <- melt(df, id.vars = c("Gene", "Time"), measure.vars = c("FDR", "FC", "FCSD"))
    wide.df.2 <- dcast(wide.df, Gene ~ Time + variable, value.var = c("value"))
    df.names <- wide.df.2[,-1] 
    row.names(df.names) <- wide.df.2[,1]
    df.ordered <- df.names[,time_order]
    final <- data.frame(round(data.matrix(df.ordered), 3))
    setDT(final, keep.rownames = TRUE)[]
    CHANGE <- mapply(FUN = function(FC1, FC2, FC3, FC4, FC5, FC6, FDR1, FDR2, FDR3, FDR4, FDR5, FDR6) {
      UpDownArrow(FC1, FC2, FC3, FC4, FC5, FC6, FDR1, FDR2, FDR3, FDR4, FDR5, FDR6)},
                            final$X3_FC, 
                            final$X12_FC, 
                            final$X24_FC, 
                            final$X72_FC, 
                            final$X168_FC, 
                            final$X336_FC,
                            final$X3_FDR, 
                            final$X12_FDR, 
                            final$X24_FDR, 
                            final$X72_FDR, 
                            final$X168_FDR, 
                            final$X336_FDR)
    final <- add_column(final, CHANGE, .after="rn")
    
    return(final)
}

HeatMapMatrix <- function(df) {
    require(reshape2)
    time_order <- c("3", "12", "24", "72", "168", "336")
    wide.df <- dcast(df, Gene ~ Time, value.var = "FC")
    df.names <- wide.df[,-1] 
    row.names(df.names) <- wide.df[,1]
    df.ordered <- df.names[,time_order]
    colnames(df.ordered) <- c("3 Hours", "12 Hours", "24 Hours", "72 Hours", "1 Week", "2 Weeks")
    return(data.matrix(df.ordered))
}
```
