## Supplementary material for "Temporal Profile of Transporter mRNA Expression in the Brain after Traumatic Brain Injury in Developing Rats": Archive containing raw data, r functions, and manuscript (Rmd): main.Rmd

````
---
always_allow_html: yes
bibliography: references/All.bib
fontsize: 12pt
geometry: margin=1in
mainfont: Arial
csl: references/american-society-for-pharmacology-and-experimental-therapeutics.csl
header-includes:
- \usepackage{setspace}
- \usepackage{lscape}
- \newcommand{\blandscape}{\begin{landscape}}
- \newcommand{\elandscape}{\end{landscape}}
- \usepackage{csvsimple}
output:
  pdf_document:
    latex_engine: xelatex
  word_document: default
---

```{r setup, include=FALSE, echo=FALSE}

#

if (!require(ggplot2)) {
  install.packages("ggplot2")
  library(ggplot2)
}

if (!require(gplots)) {
  install.packages("gplots")
  library(gplots)
}

if (!require(kableExtra)) {
  install.packages("kableExtra")
  library(kableExtra)
}

if (!require(gridExtra)) {
  install.packages("gridExtra")
  library(gridExtra)
}

if (!require(draw)) {
  install.packages("draw")
  library(draw)
}

if (!require(png)) {
  install.packages("png")
  library(png)
}

if (!require(dplyr)) {
  install.packages("dplyr")
  library(dplyr)
}

if (!require(formattable)) {
  install.packages("formattable")
  library(formattable)
}

if (!require(data.table)) {
  install.packages("data.table")
  library(data.table)
}

require(grid)
require(knitr)
require(reshape2)
source("functions/functions.R")

```


## Temporal Profile of Transporter mRNA Expression in the Brain after Traumatic Brain Injury in Developing Rats

&nbsp;
&nbsp;

Solomon M. Adams, Fanuel T. Hagos, Jeffrey P. Cheng, Robert S. B. Clark, Patrick M. Kochanek, Anthony E. Kline, Samuel M. Poloyac, Philip E. Empey

&nbsp;
&nbsp;

Department of Pharmacogenomics, School of Pharmacy, Shenandoah University, Fairfax, VA (S.M.A.);  
Department of Pharmaceutical Science, School of Pharmacy, University of Pittsburgh, Pittsburgh, PA (S.M.A., F.T.H., S.M.P.);  
Department of Pharmacy and Therapeutics, School of Pharmacy, University of Pittsburgh, Pittsburgh, PA (P.E.E.);  
Department of Critical Care Medicine, University of Pittsburgh School of Medicine (R.S.B.C., P.M.K.);  
Department of Physical Medicine and Rehabilitation, University of Pittsburgh School of Medicine (A.E.K.);  
Safar Center for Resuscitation Research, University of Pittsburgh, Pittsburgh, PA (J.P.C., R.S.B.C., P.M.K, A.E.K.)  

\newpage

# Running Title Page

**Running Title:** Pediatric Neurotrauma Associated Transporter Expression

&nbsp;
&nbsp;

**Corresponding Author:**  
Philip E. Empey, PharmD, PhD  
Department of Pharmacy and Therapeutics, School of Pharmacy  
University of Pittsburgh  
205 Salk Pavilion  
335 Sutherland Drive  
Pittsburgh, PA 15213  
Phone: (412) 648-7219  
Email: pempey\@pitt.edu  

&nbsp; 
&nbsp; 

**Text Pages: 12**  
**Tables: 3**  
**Figures: 2**  
**References: 34**  
**Abstract Word Count: 249**  
**Body Word Count: 3439**  

\newpage

\raggedright

\doublespacing

# ABSTRACT

Traumatic brain injury (TBI) is a leading cause of death in children and young adults; however, new pharmacologic approaches have failed to improve outcomes in clinical trials. 
Transporter proteins are central to the maintenance of homeostasis within the neurovascular unit, and regulate drug penetration into the brain. 
Our objective was to measure transporter temporal changes in expression in the hippocampus and cortex after experimental TBI in developing rats. 
We also evaluated the expression of transporters in brain, liver, and kidney across the age spectrum in both pediatric and adult rats. 
Eighty post-natal day (PND)-17 rats and four adult rats were randomized to receive controlled cortical impact (CCI), sham surgery, or no surgery. 
mRNA transcript counts for 27 ATP-binding cassette and solute carrier transporters were measured in the hippocampus, cortex, choroid plexus, liver, and kidney at 3h, 12h, 24h, 72h, 7d, and 14d post injury. 
After TBI, the expression of many transporters (*Abcc2, Slc15a2, Slco1a2*) decreased significantly in the first 24 hours, with a return to baseline over 7-14 days. 
Some transporters (*Abcc4, Abab1a/b, Slc22a4*) showed a delayed increase in expression. 
Baseline expression of transporters was of a similar order of magnitude in brain tissues relative to liver and kidney. 
Findings suggest that transporter-regulated processes may be impaired in the brain early after TBI and are potentially involved in the recovery of the neurovascular unit. 
Our data also suggest that transport-dependent processes in the brain are of similar importance as those seen in organs involved in drug metabolism and excretion.

## Significance Statement

Baseline transporter mRNA expression in the central nervous system is of similar magnitude as liver and kidney, and experimental traumatic brain injury is associated with acute decrease in expression of several transporters, while others show delayed increase or decrease in expression. 
Pharmacotherapy following traumatic brain injury should consider potential pharmacokinetic changes associated with transporter expression. 

\newpage

# INTRODUCTION

Traumatic brain injury (TBI) is a leading cause of morbidity and mortality in children and young adults, contributing to nearly a third of all injury-related deaths [@Coronado2011].
TBI comprises a primary (e.g. motor vehicle accident, improvised explosive device detonation) and secondary injury. 
Secondary injury is a progressive process that appears to be driven by inflammation, dysruption in neurovascular auto-regulation, hemorrhage, excitotoxicity, axonal failure, and oxidative stress, among other mechanisms [@Kenney2016]. 
Other than traditional guidelines-based therapies targeting intracranial hypertension [@Kochanek2019], there are no new FDA approved pharmacologic agents that improve clinical outcomes from TBI [@Kochanek2015; @Taylor2017].
Drug development for TBI is challenged by variable dysruption of blood-brain barrier (BBB) permeability and other barriers in the CNS. 
It is difficult to predict BBB integrity and function following TBI, which is regulated, in part, by active transport proteins in the brain parenchyma and within the neurovascular unit [@Muoio2014]. 
This lack of knowledge limits our ability to develop drugs with favorable pharmacokinetic profiles with regard to brain penetration to optimize target engagement for patients with TBI. 

ATP-Binding Cassette (ABC) and Solute Carrier (SLC) transporter proteins are critical to homeostasis within the CNS and its associated barriers. 
Transporters are expressed on all brain barriers and within the brain parenchyma on neurons, astrocytes, and microglia [@Stieger2015a].
Highly relevant to the need for our proposed studies, pharmacologic modulation (inhibition) of solute carrier organic anion transporters after TBI has been used to successfully increase the levels of brain-directed potential neuroprotective therapies (exogenous n-acetylcysteine)  [@Hagos2016; @Clark2017]. 
The role of transporters in the brain is generally within the context of pharmacotherapy and brain penetration (i.e. pharmacokinetics).
However, independent of a role with medications, genetic variation in the ATP-Binding cassette transporters *ABCB1* and *ABCG2* are associated with clinical outcomes following TBI [@Adams2017;@Cousar2013;@Adams2017a;@Hagos2019].

The inflammatory pathways activated by TBI might affect transporter expression in the brain and in distant tissues that could importantly impact drug pharmacokinetics. 
Within the brain, Willyerd and colleagues found that protein expression of ABCC1 was higher in the cortex of injured subjects than control, but that ABCB1 showed little change [@Willyerd2016].
Kalsotra, et al investigated the changes in hepatic and renal expression of cytochrome P-450 drug-metabolizing enzymes following experimental TBI and found that expression was significantly lower in early time points after injury, even in organs outside of the brain [@Kalsotra2003].
Cytochrome P450 enzyme (CYPs) expression in the liver is regulated in part by cytokines. 
For example, increases in circulating IL-6 levels are well known to be associated with decreased expression of multiple CYP families including CYP3A, CYP2C, and CYP1A [@Abdel-Razzak1993]. 
Transporters might follow similar patterns of cytokine-based regulation of expression, which suggests that transporter expression might vary along with inflammation following TBI. 

Little is known about the directionality, timing, and magnitude of transporter expression changes following TBI in the developing brain. 
There is also a paucity of data regarding the relative levels of the expression of different transporters at baseline in the pediatric brain and their ontogeny relative to other tissues with high transporter expression (e.g. liver, kidney). 
We sought to assess two objectives.
First we assessed absolute transporter mRNA expression in hippocampus and cortex after experimental TBI in developing rats.
Second we assessed absolute transporter mRNA expression in multiple tissues across the age spectrum in both developing and adult rats.
To address these objectives, we used a novel multiplex bar coding technology to provide absolute transcript counts.
Based on well-defined inflammatory pathways in TBI, we hypothesized that mRNA expression for the majority of ABC and SLC transporters would be decreased following experimental TBI in conjunction with enrichment of pathways associated with inflammation and hypoxia. 
We also hypothesized that transporter mRNA levels in brain would, however, be less robust than those seen in organs traditionally known to be essential to drug metabolism and excretion, namely, liver and kidney, respectively. 


# METHODS

## Animal Husbandry
All animal experiments were performed in accordance with the University of Pittsburgh Institutional Animal Care and Use Committee. 
Male Sprague-Dawley rats (N=84, Charles River, Wilmington, MA) were housed in a temperature (22&deg;C) controlled room with a 12h light/dark cycle, allowed access to water and chow *ad libitum*, and weighed daily. 
Seventy-two post-natal day (PND) 17 rats were randomized 2:1 to experimental TBI and sham injury. 
Eight PND-17 rats and four adult rats (300g) remained naïve to injury for evaluation of ontogeny. 
Animal groupings and time of sacrifice were randomized *a priori* to control for litter effects.

## Experimental TBI
Experimental TBI was carried out using the controlled cortical impact (CCI) model [@Dixon1991]
Rats were anesthetized with a 2:1 mixture of N\textsubscript{2}O and oxygen with 4% isoflurane for induction and 2% isoflurane for maintenance through a nosecone. 
Continuous rectal temperature was measured and maintained at 37&deg;C using a heated pad. 
The head was secured using a stereotaxic frame with ear pins. 
Using sterile aseptic technique, a mid-line sagittal incision was made followed by reflection of the scalp with retractors. 
A high speed dental drill was then used to make a 7mm craniotomy in the left parietal bone, which was removed to expose the dura. 
A calibrated pneumatic piston was then used to impact the intact dura with a 2.5mm deformation and impact velocity of 4m/s. 
Sham injured animals received the full procedure with the exception of the impact.

## Sacrifice and RNA Extraction
CCI and sham rats were sacrificed at 3, 12, 24, 72, 168, and 336h post-surgery. 
Naïve PND-17 rats were sacrificed at PND-17 and at PND-31. 
Adult rats were sacrificed after acclimation in our facility. 
Rats were placed in a plastic chamber and subjected to 4% isoflurane and 2:1 N\textsubscript{2}:O\textsubscript{2} for induction then transferred to a nosecone providing 2% isoflurane with 2:1 N\textsubscript{2}:O\textsubscript{2} for maintenance of anesthesia. 
Following cardiac puncture, rats were decapitated and the brain was dissected atop an inverted glass petri dish on a bed of wet ice. 
The ipsilateral and contralateral cortices were removed first, followed by the ipsilateral and contralateral hippocampi, the choroid plexus, a single kidney, and an approximately 50g section of liver tissue. 
Each tissue sample was immediately frozen with liquid nitrogen and stored at -80°C until further processing.

RNA was isolated from tissues with the RNeasy Mini Kit (QIAGEN, Hilden, Germany) using the manufacturer’s protocol with modification for hippocampus and cortex. 
Briefly, the entire ipsilateral cortex/hippocampus was homogenized in 300$\mu$L radioimmunoprecipitation assay buffer
(Tris HCl, EDTA, NaCl, Triton-X, Sodium Deoxycholate, Protease Inhibitor Cocktail) with 1% v/v beta-mercaptoethanol, from which a 150$\mu$L aliquot was introduced to the protocol. 
For liver and kidney, the standard procedure was used after manually pulverizing the liver and kidney tissue with a mortar and pestle cooled with liquid nitrogen. 
For the choroid plexus, we also used the standard procedure using the entirety of the tissue. 
RNA concentration and quality were assessed with optical density at 260nm and 260:280 ratio, respectively (NanoDrop, Thermo Fisher Scientific, Waltham, MA).

## Gene Expression Panel
We focused our evaluation on a targeted list of transporters of relevance to the pathophysiology of secondary injury and/or drug development.
First, we conducted a literature review to identify transporters, biomarkers, and transcription factors that were of probable importance to TBI pathophysiology. 
This was judged based on 1) reported expected expression in the brain, 2) hypothesized or documented roles in brain disease, and 3) role in transport for medications used in TBI patients. 
Next, we added transporters that are recommended for evaluation in the context of drug-development by the International Transporter Consortium and FDA guidelines [@Giacomini2013].
Biomarkers and transcription factors were added to the panel based on evidence for a role in TBI and data supporting their role in modulating expression of genes related to metabolism.
Finally, human genes were transposed to their corresponding rat homologues by using the NCBI HomoloGene (https://www.ncbi.nlm.nih.gov/homologene).
Housekeeping genes were selected based on evidence for stability post TBI [@Rhinn2008;@Julian2014;@Penna2011].
The final list of genes was developed into a custom panel for absolute quantitation of mRNA using the nCounter^&reg;^ (NanoString Technologies, Seattle, WA) platform.

## Measurement of mRNA Expression with Nanostring
mRNA quantification was conducted using the manufacturer’s instructions for Nanostring at the Genomics Research Core at the University of Pittsburgh. 
Briefly, 100ng mRNA (at 30ng/$\mu$L) was added to capture probes with a biotinylated tail and reporter probes with a fluorescent barcode complementary to target transcripts. 
Hybridization of each sample at 65°C occurred overnight, samples were loaded on the nCounter^&trade;^ Prep Station which is an automated liquid handler that removes excess sample and unbound codesets in a two-step bead clean-up protocol. 
After purification, probe/transcript complexes are eluted and transferred to the streptavidin coated nCounter® cartridge, which binds the biotinylated region of the probe/transcript complex. 
Next, a current was applied to align the reporter probes. 
The cartridge was then scanned with a Digital Analyzer, which scans 600 fields of view across each cartridge and counts the occurrence of each fluorescent barcode. 

## Statistical Methods
Statistical analyses were carried out using R v3.5.1 (R Development Core Team, Vienna, Austria) [@Rv3_5_1]. 
Data visualization was performed using ggplot2 [@Wickham2016]. 
Raw transcript counts were obtained from the nCounter^&reg;^ platform and normalized with the R package NanoStringNorm [@Waggott2012].
Technical variation was normalized by adjusting individual counts the geometric mean of all probe counts. 
The mean of negative controls was subtracted from each probe count to control for background variation. 
Variation in total RNA content was normalized to the geometric mean of absolute counts of housekeeping genes. 
Samples with a background value greater than three standard deviations from the mean excluded. 
Post-normalization counts were log2 transformed for downstream analysis.

```{r, eval=F, include=F}

# Here is the actual function call, note that the RCC_Files are not included

Data_Norm  <- NanoStringNorm(RCC_Files, 
                             CodeCount = "geo.mean", 
                             Background = "mean", 
                             SampleContent = "housekeeping.geo.mean", 
                             OtherNorm = "none", 
                             take.log = TRUE, 
                             return.matrix.of.endogenous.probes = FALSE
                             ) #Loads files into function
```

P values were corrected using Benjamini-Hochberg False Discovery Rate (FDR) to produce q values [@Benjamini1995].
For all analyses, a q-value of 0.05 was considered statistically significant. 
Gene ontogeny was determined from measurements in naïve rats. 
A baseline expression was established at PND-17, to which PND-31 and adult expression was compared with an unpaired t-test.
Data for naive rats are presented as fold change relative to PND-17. 
Expression changes associated with injury in the ipsilateral hippocampus and cortex were compared with sham animals at each time point with unpaired t-tests. 
Data for injured vs. sham rats are presented as fold change. 

# RESULTS

## Gene Expression Panel Design
The literature review identified 27 rat transporters corresponding to 24 human homologues. 
To provide a limited physiological context for changes in expression, we selected four transcription regulators known for their roles in injury and in modulation of transporter expression (*Nfe1l2, Hif1a, Nr1i2, Il6*). 
To serve as validation and to ensure adequate control for the model, we added five biomarkers (*Edn1, Gfap, Vim, Ngb, Icam1*) for injury that are well characterized post injury, and four housekeepers to normalize the expression (*Ppia, Gapdh, Tbp, Hprt1*). 
Genes included in the panel, their human homologues, and common names are included in **Table 1**.

## Transporter Ontogeny 
```{r, eval=F, include=F}

# Numbers obtained with the following

unique(as.character(ontogeny.transporters[ontogeny.transporters$FDR < 0.05 & ontogeny.transporters$Tissue=="cortex",]$Gene))
unique(as.character(ontogeny.transporters[ontogeny.transporters$FDR < 0.05 & ontogeny.transporters$Tissue=="hippocampus",]$Gene))
unique(as.character(ontogeny.transporters[ontogeny.transporters$FDR < 0.05 & ontogeny.transporters$Tissue=="CP",]$Gene))
unique(as.character(ontogeny.transporters[ontogeny.transporters$FDR < 0.05 & ontogeny.transporters$Tissue=="liver",]$Gene))
unique(as.character(ontogeny.transporters[ontogeny.transporters$FDR < 0.05 & ontogeny.transporters$Tissue=="kidney",]$Gene))

```

We found diversity in transporter expression across different tissues, organs, and time points in uninjured rats (**Figure 2**). 
*Slc7a5* and *Slco1a2* showed differential expression PND-31 and/or adulthood relative to PND-17 in the cortex. 
Among others, we found that *Abcb1a, Abcc2, Abcc4*, and *Slc22a4* were differentially expressed from PND-17 to adulthood in the hippocampus, liver, and kidney.

*Slc2a1*, a glucose transporter, was the most prominently expressed transporter in brain relative to non-CNS expression. 
In tissues where adult tissues were studies (cortex and hippocampus), little change was observed between PND-31 and adulthood.

## Expression Changes Due to Injury
Baseline expression of transporters (PND-17) are shown for all tissues in **Figure 1**. 
*Slc16a1* was the most highly expressed transporter at a consistent magnitude in all tissues. In brain, *Slc2a1* was the most highly expressed transporter.

Expression changes for all genes in cortex and hippocampus following experimental TBI are shown in **Tables 2 and 3**. 
These show the directionality of significant changes, grouped into early (3-24 hours), mid (72 hours), and late (1-2 weeks) following TBI. 
Fold change is calculated with $log_2(TBI)-log_2(Control)$ and FDR corresponds to the FDR corrected p value.
Expression of many transporters decreased in both hippocampus and cortex following TBI (e.g. *Abcc2, Abcc8, Slc22a6, Slc22a8*), while some showed increased expression (e.g. *Abcc4, Slc22a4*).
Decreased expression was primarily observed early (~3-24 hours) after TBI, followed by a gradual return to baseline with some transporters showing an inversion (i.e. decreased, then increased). 
Eight out of nine ABC transporters showed significantly differential expression from sham at a minimum of one time point in the hippocampus. 
Despite its negligible expression at baseline, *Abcb11* showed a significant spike in expression that was significant in the cortex. 
These patterns follow expression patterns similar to *Hif1*$\alpha$ and *Il6*. 
Surprisingly, we found significantly decreased expression for *Ngb* in both hippocampus and cortex following injury.

Many of the patterns in expression cluster with the transcription factors: *Il-6*, which showed an immediate spike in expression following TBI with a quick return to baseline. 
Others followed a trend set by the transcription factor *Nfe2l2*, which spiked in expression immediately following injury with a secondary spike at 72 hours.


# DISCUSSION

We used innovative mRNA expression profiling technology to determine absolute transcript counts of transporters in the brain following experimental TBI in developing rats to gain insight into this mechanism for pediatric TBI. 
Consistent with our first hypothesis, after TBI, expression was predominantly decreased immediately (3-24 hours) following TBI, followed by gradual return to baseline expression over 14 days and in some cases, increased expression from baseline following injury. 
Biomarkers and transcription factors showed expression patterns reflective of previous studies, with the exception of Ngb, which was acutely down-regulated following CCI in both cortex and hippocampus. 
Surprisingly, refuting our second hypothesis, we also found that baseline transporter expression in the brain often achieved levels similar or exceeding that found in tissues where they have a primary role in drug elimination.

TBI was associated with differential transporter mRNA expression in the hippocampus and cortex. 
In the hippocampus, transporters relevant to drug disposition including *Abcb1a/b, Abcc2, Abcg2, Slc22a6, and Slc22a8* were decreased early following TBI. 
However, *Abcb1a/b* was increased early after injury in the cortex, which may more accurately reflect BBB expression. 
Other transporters (e.g. *Abcc4, Slc22a4*) had increased expression in hippocampus and cortex following injury. 
Substrates for these transporters (e.g. phenytoin) might have altered distribution into the CNS compartment in patients after TBI. 
These findings are similar to work in tissues outside of the CNS by Kalsotra and colleagues, who found that hepatic and renal expression of CYP enzymes decrease following experimental TBI with a later increase in expression [@Kalsotra2003].
Changes in expression also mirror changes in expression of transcription factors that can impact the expression of metabolic proteins (e.g. enzymes, transporters). 
Particularly, early inhibitions in expression of *Abcb1a, Abcc2, Abcg2*, and *Slco* in the hippocampus mirror changes found in *Il6*, which is known to be associated with decreased expression of transporters and CYP enzymes [@Abdel-Razzak1993;@Poller2010]. 
This acute decrease in expression in injured brain might support protection of vulnerable etissue from the toxic byproducts associated with secondary injury and tissue repair [@Medzhitov2008]. 
*Abcb11*, which is normally not detectable in the brain, was increased with similar timing as a spike in *Nfe2l2* at 72 hours post injury [@Warren2009]. 

*Ngb* (neuroglobin) mRNA expression post TBI decreased in the hippocampus and cortex following TBI. 
Neuroglobin is a neuro-protective hemoprotein that binds O\textsubscript{2}, NO, and CO in the brain [@Lazzarino2014].
It can mitigate oxidative stress and hypoxia, and over-expression is neuroprotective [@Lazzarino2014;@Chuang2010].
Previous work by Di Pietro and colleagues studied neuroglobin expression following experimental diffuse (mild and severe) TBI in adult rats.
They found increased expression of neuroglobin immediately following injury in both the severe and mild TBI models [@Lazzarino2014].
This suggests possible divergence with the brain's response to our focal injury model (CCI) versus the response following a diffuse injury model. 
Alternatively, age- or injury severity-related differences could be important.
This contrasts with the other biomarkers evaluated on this panel, which followed expected patterns of expression changes.

To explore tissue differences in transporter expression and expression across age we compared expression in three brain regions (hippocampus, cortex, and choroid plexus) to liver and kidney.  
Baseline expression of transporters was similar in the cortex, hippocampus, and choroid plexus. 
Transporters related to metabolism (i.e. glucose, lactate, pyruvate) including *Slc16a1* and *Slc2a1* were among the highest expressed transporters in hippocampus, cortex, and choroid plexus. 
Expression of transporter mRNA in the brain was often found to be within the same order of magnitude as liver and kidney. 
We found non-zero expression of all probed transporters included in the draft FDA industry guidance for in vitro drug-drug interaction studies, including *Abcb1a, Abcg2, Slc22a2, Slc22a6, Slc22a8, Slc47a1*, and *Slco1a*. 
Of the transporters included in the FDA guidance Abcb1a and Abcg2 were found to be expressed at high and similar levels across the brain, liver, and kidneys [@FDA2017]. 
*ABCB1* and *ABCG2* are among the most highly expressed transporters on the BBB in humans, and this is mirrored in their rat homologues [@Wijaya2017]. 

Ontogeny findings are similar to those of Mooij and colleagues, which show changes associated with development alone in the gut [@Mooij2014].
We found changes with development to be more pronounced in the liver and kidney, wherein *Abcb1a/b, Abcc, Slc22a, Slc47a1*, and *Slco1a* transporters/subfamilies showed differential expression with development. 
Conversely, a novel observation in our work is that in contrast to liver and kidney, brain expression of transporters was relatively conserved, with few significant changes from PND-17 to adult seen in *Abcc, Abcb*, and *Slco* transporter subfamilies. 
Nevertheless, brain and systemic disposition of xenobiotics that are known substrates of these pathways should be considered when extrapolating adult doses to pediatrics.
However, this finding merits additional exploration given its potential impact on the brain PK and resultant target engagement and pharmacodynamics consequences comparing developing and adult brain.

There are several limitations of this work that should be considered. 
Our model of severe TBI (CCI) is a well studied focal model, but may not translate to other type of TBI such as diffuse injuries, mild TBI, or mild repetitive TBI. 
To minimize time from dissection to freezing, we rinsed tissue with saline rather than perfusing the rats. 
This raises the possibility of minor contamination of mRNA found in blood.
However, the variability in results between tissues suggests this was not an significant.
Finally, while mRNA data provides insight into the biological mechanisms and the method used allowed for absolute quantification without amplification, mRNA changes may not always translate to changes in protein levels, particularly in the injured brain, where protein synthesis may be impaired in TBI core or penumbral regions [@Kochanek2008].

To our knowledge, this is the first investigation in pediatric TBI that provides absolute transcript counts of xenobiotic transporters. 
We believe the multiple brain tissues evaluated in a robust time-course design are an important contribution toward greater understanding of a vastly understudied facet governing the secondary injury following TBI in children, along with impacting the response to current or future pharmacological therapies. 
Indeed, ontological changes associated with development from pediatric to adulthood suggest a dynamic biochemical environment that may have implications in drug development and dosing. 
These baseline expression levels also differ, in some cases, across brain regions suggesting that the disposition of xenobiotics and endogenous substrates is dynamic across brain regions and barriers. 
Importantly, experimental TBI in developing rats shows patterns of acutely decreased transporter expression in the cortex and hippocampus, with some divergent increases in expression at various time points. 
Changes in expression from the acute to chronic phase post TBI also support the need for greater evaluation of substrate/transporter interactions and ultimately suggest the potential need for the use of dynamic treatment protocols that call for variable drug dosing with time post injury.

# Acknowledgements

Support for this research from grants from the National Institutes of Health 1TL1 TR001858-01 (to S.M.A.), R01NS084967 (to A.E.K.), and R01HD069620 (to A.E.K.) for grant support. We also acknowledge support from the University of Pittsburgh Genomics Research Core for running the Nanostring panel. 

# Authorship Contribution

*Participated in research design*: Adams, Cheng, Clark, Kochanek, Kline, Poloyac, Empey

*Conducted experiments*: Adams, Cheng

*Performed data analysis*: Adams, Cheng, Hagos

*Wrote or contributed to the writing of the manuscript*: Adams, Hagos, Kochanek, Empey

\newpage

# REFERENCES

\setlength{\parindent}{-0.2in}
\setlength{\leftskip}{0.2in}
\setlength{\parskip}{8pt}
\noindent

<div id="refs"></div>

\newpage
\setlength{\parindent}{0in}


```{r, eval=T, include=F}

# Prepare Environment


#setwd() 

# Load Data
expression.data <- read.csv("data/all_tissues.csv", header = T, sep=",")
genes <- read.csv("data/genes.csv", sep=",", header=F, as.is = T)
times <- c(3,12,24,72,168,336)


# Generate Comparisons (Injury)

hippocampus.injury.data <- CompareGroupsInjury(expression.data, 
                                         times = times, 
                                         genes = genes, 
                                         tissue = "hippocampus", 
                                         group1 = "CCI", 
                                         group2 = "Sham")

cortex.injury.data <- CompareGroupsInjury(expression.data, 
                                         times = times, 
                                         genes = genes, 
                                         tissue = "cortex", 
                                         group1 = "CCI", 
                                         group2 = "Sham")

injury.data <- rbind(cortex.injury.data, hippocampus.injury.data)

# Generate Comparisons (Naive)

cortex.naive.data <- CompareGroupsNaive(expression.data, 
                                        genes = genes, 
                                        times = c(0, 336, 1000), 
                                        tissue =  "cortex", 
                                        base_time = 0)

hippocampus.naive.data <- CompareGroupsNaive(expression.data, 
                                        genes = genes, 
                                        times = c(0, 336, 1000), 
                                        tissue =  "hippocampus", 
                                        base_time = 0)

CP.naive.data <- CompareGroupsNaive(expression.data, 
                                             genes = genes, 
                                             times = c(0, 336), 
                                             tissue =  "CP", 
                                             base_time = 0)

liver.naive.data <- CompareGroupsNaive(expression.data, 
                                    genes = genes, 
                                    times = c(0, 336), 
                                    tissue =  "liver", 
                                    base_time = 0)

kidney.naive.data <- CompareGroupsNaive(expression.data, 
                                       genes = genes, 
                                       times = c(0, 336), 
                                       tissue =  "kidney", 
                                       base_time = 0)

naive.data <- rbind(kidney.naive.data, liver.naive.data, CP.naive.data, hippocampus.naive.data, cortex.naive.data)

```
\newpage

# Figure Legends


**Figure 1**  
Baseline transporter expression (PND-17) in kidney, liver, choroid plexus, hippocampus, and cortex. Error bars show standard deviation. 

**Figure 2**  
Ontogeny of transporters in cortex, hippocampus, choroid plexus, liver, and kidney. Point color indicates statistical significance (FDR < 0.05). 

\newpage

```{r eval=F, echo=F, error=F, out.width='.49\\linewidth', fig.width=5, fig.height=5,fig.show='hold',fig.align='center'}
# Heatmap

# Not included in manuscript. Can change eval to T to include. 

#**Figure x**

par(cex.main=0.75)

heatmap.2(
    HeatMapMatrix(cortex.injury.data), 
    dendrogram = "row",
    Colv = F, 
    cexRow = 0.5, 
    cexCol = 0.75,
    offsetRow = -0.4,
    offsetCol = -0.4,
    lhei = c(1,5), 
    lwid = c(1,5),
    key.title = NA,
    key.xlab = NA,
    key.ylab = NA,
    main = "Ipsilateral Cortex",
    density.info = "none"
)

heatmap.2(
    HeatMapMatrix(hippocampus.injury.data), 
    dendrogram = "row",
    Colv = F, 
    cexRow = 0.5, 
    cexCol = 0.75,
    offsetRow = -0.4,
    offsetCol = -0.4,
    lhei = c(1,5), 
    lwid = c(1,5),
    key.title = NA,
    key.xlab = NA,
    key.ylab = NA,
    main = "Ipsilateral Hippocampus",
    density.info = "none"
)


```

\newpage

**Figure 1**
```{r eval=T, echo=F, error=F, out.width="125%", out.height="125%", fig.height=10, fig.width=15, fig.show='hold',fig.align='center',out.extra='angle=90'}

baseline.transporters <- naive.data[naive.data$GeneGroup %in% c("ABC","SLC") & naive.data$Time == "0",]

ggplot(
  data=baseline.transporters, 
  aes(
    x = reorder(Gene, as.numeric(as.character(Mean))), 
    y=as.numeric(as.character(Mean)), 
    fill=Tissue)) +
  geom_bar(
    stat = "identity", 
    position=position_dodge(), 
    color="black", 
    width=0.75,
    size=0.25) +
  geom_errorbar(
    mapping=aes(x=Gene,
                ymin=as.numeric(as.character(Mean)),
                ymax=as.numeric(as.character(Mean)) + as.numeric(as.character(SD))
                            ),
                position=position_dodge(width=0.75), size=0.4, width=0.5) +
  theme_bw() +
  theme(axis.text.x = element_text(angle = 90, hjust = 1), axis.title.x=element_blank()) +
  ylab("Log2 Transcript Count") +
  scale_fill_discrete(name="Tissue",
                      breaks=c("kidney", "liver", "CP", "hippocampus", "cortex"),
                      labels=c("Kidney", "Liver", "Choroid Plexus", "Hippocampus", "Cortex"))


```

\newpage

**Figure 2**
```{r eval=T, echo=F, error=F, out.width="125%", out.height="125%", fig.height=10, fig.width=15, fig.show='hold',fig.align='center',out.extra='angle=90'}

# Transporter Ontogeny

ontogeny.transporters <- naive.data[naive.data$GeneGroup %in% c("ABC","SLC"),]

ggplot(ontogeny.transporters, aes(x=as.numeric(as.character(Time)), y=as.numeric(as.character(Mean)), color=Tissue)) +
    facet_wrap(Gene~., nrow=3) +
    geom_line() + 
    geom_point(aes(shape=as.numeric(as.character(FDR)) >= 0.05), na.rm=TRUE) + 
    theme_bw() + 
    ylab("Log2 Normalized Transcript Count") +
    scale_x_continuous(breaks=c(0,336,1000), labels = c("PND-17", "PND-31", "Adult"), expand = c(0.2,0.5)) +
    theme(axis.text.x = element_text(angle = 90, hjust = 1), axis.title.x=element_blank()) +
    scale_color_manual(name="Tissue",
                       labels=c("Kidney","Liver","Choroid Plexus", "Hippocampus", "Cortex"), 
                       values=c("red", "brown", "pink", "blue", "green")) + 
    scale_shape_manual(name="FDR",
                       labels=c("<0.05", ">=0.05"),
                       values=c(17,46))


```

\newpage

\fontsize{7}{7}\selectfont

**Table 1** Genes Included on Expression Panel


\csvreader[head to column names, tabular=ccc, 
    table head= \toprule \bfseries Rat~Gene & \bfseries Human~Gene & \bfseries Common~Name,
    table foot = \\\bottomrule]
    {data/table1.csv}{}
{
	\ifthenelse{\equal{\HGNCID}{}}
    {\\ \midrule & \HumanHomologue & \\ \midrule}
    {\textit{\HGNCID} & \textit{\HumanHomologue} & \CommonName}
}

\newpage

**Table 2** Expression Changes in the Ipsilateral Cortex Post TBI

```{r eval=T, echo=F, warning=F, message=F, out.extra='angle=90'}

# Injury

InjuryTable(cortex.injury.data) %>% 
  mutate(
      X3_FDR = ifelse(X3_FDR < 0.05,
                  cell_spec(sprintf("%.3f", round(X3_FDR, 3)), bold = T),
                  cell_spec(sprintf("%.3f", round(X3_FDR, 3)))),
      X12_FDR = ifelse(X12_FDR < 0.05,
                  cell_spec(sprintf("%.3f", round(X12_FDR, 3)), bold = T),
                  cell_spec(sprintf("%.3f", round(X12_FDR, 3)))),
      X24_FDR = ifelse(X24_FDR < 0.05,
                  cell_spec(sprintf("%.3f", round(X24_FDR, 3)), bold = T),
                  cell_spec(sprintf("%.3f", round(X24_FDR, 3)))),
      X72_FDR = ifelse(X72_FDR < 0.05,
                  cell_spec(sprintf("%.3f", round(X72_FDR, 3)), bold = T),
                  cell_spec(sprintf("%.3f", round(X72_FDR, 3)))),
      X168_FDR = ifelse(X168_FDR < 0.05,
                  cell_spec(sprintf("%.3f", round(X168_FDR, 3)), bold = T),
                  cell_spec(sprintf("%.3f", round(X168_FDR, 3)))),
      X336_FDR = ifelse(X336_FDR < 0.05,
                  cell_spec(sprintf("%.3f", round(X336_FDR, 3)), bold = T),
                  cell_spec(sprintf("%.3f", round(X336_FDR, 3))))
      
  ) %>%
  kable(format="latex", booktabs = TRUE, row.names=F, 
        col.names = c("", "EML*", rep(c("FC", "SD", "FDR"),6)),
         align = c("l", "l", rep(c("r"),18)), escape = FALSE) %>%
  kable_styling(latex_options = c("scale_down")) %>%
  add_header_above(header = c(" "= 2,
                              "3 Hours" = 3,
                              "12 Hours" = 3, 
                              "24 Hours" = 3, 
                              "72 Hours" = 3,
                              "1 Week" = 3,
                              "2 Weeks" = 3
  ), 
  bold = FALSE, italic = FALSE,
  monospace = FALSE, align = "c", escape = FALSE, line = TRUE) %>%
  column_spec(column = 1, italic = T) %>%
  footnote(general="EML*: Early (3-24 hours), Mid (72 hours), Late (7-14 Days); ↑: expression increased, ↓: expression decreased, ↔: expression did not change significantly", general_title = "", footnote_as_chunk = T)


```

\newpage

**Table 3** Expression Changes in the Ipsilateral Hippocampus Post TBI

```{r eval=T, echo=F, warning=F, message=F, out.extra='angle=90', results="asis"}

# Injury

InjuryTable(hippocampus.injury.data) %>% 
    mutate(
      X3_FDR = ifelse(X3_FDR < 0.05,
                  cell_spec(sprintf("%.3f", round(X3_FDR, 3)), bold = T),
                  cell_spec(sprintf("%.3f", round(X3_FDR, 3)))),
      X12_FDR = ifelse(X12_FDR < 0.05,
                  cell_spec(sprintf("%.3f", round(X12_FDR, 3)), bold = T),
                  cell_spec(sprintf("%.3f", round(X12_FDR, 3)))),
      X24_FDR = ifelse(X24_FDR < 0.05,
                  cell_spec(sprintf("%.3f", round(X24_FDR, 3)), bold = T),
                  cell_spec(sprintf("%.3f", round(X24_FDR, 3)))),
      X72_FDR = ifelse(X72_FDR < 0.05,
                  cell_spec(sprintf("%.3f", round(X72_FDR, 3)), bold = T),
                  cell_spec(sprintf("%.3f", round(X72_FDR, 3)))),
      X168_FDR = ifelse(X168_FDR < 0.05,
                  cell_spec(sprintf("%.3f", round(X168_FDR, 3)), bold = T),
                  cell_spec(sprintf("%.3f", round(X168_FDR, 3)))),
      X336_FDR = ifelse(X336_FDR < 0.05,
                  cell_spec(sprintf("%.3f", round(X336_FDR, 3)), bold = T),
                  cell_spec(sprintf("%.3f", round(X336_FDR, 3))))
      
  ) %>%
   kable(format="latex", booktabs = TRUE, row.names=F, 
         col.names = c("", "EML*", rep(c("FC", "SD", "FDR"),6)),
         align = c("l", "l", rep(c("r"),18)), escape = FALSE) %>%
   kable_styling(latex_options = c("scale_down")) %>%
    add_header_above(header = c(" "= 2,
                                "3 Hours" = 3,
                                "12 Hours" = 3, 
                                "24 Hours" = 3, 
                                "72 Hours" = 3,
                                "1 Week" = 3,
                                "2 Weeks" = 3
                                ), 
                     bold = FALSE, italic = FALSE,
  monospace = FALSE, align = "c", line = TRUE) %>%
    column_spec(column = 1, italic = T) %>%
  footnote(general="EML*: Early (3-24 hours), Mid (72 hours), Late (7-14 days); ↑: expression increased, ↓: expression decreased, ↔: expression did not change significantly", general_title = "", footnote_as_chunk = T)


```
````
