## Supplementary material for "Temporal Profile of Transporter mRNA Expression in the Brain after Traumatic Brain Injury in Developing Rats": Archive containing raw data, r functions, and manuscript (Rmd): All.bib

Automatically generated by Mendeley Desktop 1.19.4
Any changes to this file will be lost if it is regenerated by Mendeley.

BibTeX export options can be customized via Preferences -> BibTeX in Mendeley Desktop

@article{Kochanek2019,
author = {Kochanek, Patrick M. and Tasker, Robert C. and Carney, Nancy and Totten, Annette M. and Adelson, P. David and Selden, Nathan R. and Davis-O'Reilly, Cynthia and Hart, Erica L. and Bell, Michael J. and Bratton, Susan L. and Grant, Gerald A. and Kissoon, Niranjan and Reuter-Rice, Karin E. and Vavilala, Monica S. and Wainwright, Mark S.},
doi = {10.1097/PCC.0000000000001735},
issn = {1529-7535},
journal = {Pediatric Critical Care Medicine},
month = {mar},
pages = {S1--S82},
title = {{Guidelines for the Management of Pediatric Severe Traumatic Brain Injury, Third Edition}},
url = {http://insights.ovid.com/crossref?an=00130478-201903001-00001},
volume = {20},
year = {2019}
}
@article{Meng2017,
abstract = {The complexity of the traumatic brain injury (TBI) pathology, particularly concussive injury, is a serious obstacle for diagnosis, treatment, and long-term prognosis. Here we utilize modern systems biology in a rodent model of concussive injury to gain a thorough view of the impact of TBI on fundamental aspects of gene regulation, which have the potential to drive or alter the course of the TBI pathology. TBI perturbed epigenomic programming, transcriptional activities (expression level and alternative splicing), and the organization of genes in networks centered around genes such as Anax2, Ogn, and Fmod. Transcriptomic signatures in the hippocampus are involved in neuronal signaling, metabolism, inflammation, and blood function, and they overlap with those in leukocytes from peripheral blood. The homology between genomic signatures from blood and brain elicited by TBI provides proof of concept information for development of biomarkers of TBI based on composite genomic patterns. By intersecting with human genome-wide association studies, many TBI signature genes and network regulators identified in our rodent model were causally associated with brain disorders with relevant link to TBI. The overall results show that concussive brain injury reprograms genes which could lead to predisposition to neurological and psychiatric disorders, and that genomic information from peripheral leukocytes has the potential to predict TBI pathogenesis in the brain.},
author = {Meng, Qingying and Zhuang, Yumei and Ying, Zhe and Agrawal, Rahul and Yang, Xia and Gomez-Pinilla, Fernando},
doi = {10.1016/j.ebiom.2017.01.046},
file = {:Users/solomon/Documents/Mendeley Desktop/Meng et al. - 2017 - Traumatic Brain Injury Induces Genome-Wide Transcriptomic, Methylomic, and Network Perturbations in Brain and Blood.pdf:pdf},
issn = {23523964},
journal = {EBioMedicine},
keywords = {Cognition,DNA methylome,Gene networks,Hippocampus,Leukocytes,Transcriptome,Traumatic brain injury,traumatic brain injury},
pages = {184--194},
pmid = {28174132},
publisher = {The Authors},
title = {{Traumatic Brain Injury Induces Genome-Wide Transcriptomic, Methylomic, and Network Perturbations in Brain and Blood Predicting Neurological Disorders}},
url = {http://dx.doi.org/10.1016/j.ebiom.2017.01.046},
volume = {16},
year = {2017}
}
@article{Dinsmore2013,
author = {Dinsmore, Judith},
doi = {10.1093/bjaceaccp/mkt010},
file = {:Users/solomon/Documents/Mendeley Desktop/Dinsmore - 2013 - Traumatic brain injury an evidence-based review of management.pdf:pdf},
isbn = {1743-1816$\backslash$r1743-1824},
issn = {1743-1816},
journal = {Continuing Education in Anaesthesia, Critical Care {\&} Pain },
pages = {1--7},
title = {{Traumatic brain injury: an evidence-based review of management}},
url = {http://ceaccp.oxfordjournals.org/content/early/2013/02/24/bjaceaccp.mkt010.short},
year = {2013}
}
@article{Perry2016,
author = {Perry, David C and Sturm, Virginia E and Peterson, Matthew J and Pieper, Carl F and Bullock, Thomas and Boeve, Bradley F and Miller, Bruce L and Guskiewicz, Kevin M and Berger, Mitchel S and Kramer, Joel H and Welsh-Bohmer, Kathleen A},
doi = {10.3171/2015.2.JNS14503.Traumatic},
file = {:Users/solomon/Documents/Mendeley Desktop/Perry et al. - 2016 - Traumatic brain injury is associated with subsequent neurologic and psychiatric disease a meta-analysis.pdf:pdf},
journal = {Journal of neurosurgery},
number = {2},
pages = {511--526},
title = {{Traumatic brain injury is associated with subsequent neurologic and psychiatric disease: a meta-analysis}},
volume = {124},
year = {2016}
}
@article{Romagnoli2016,
archivePrefix = {arXiv},
arxivId = {15334406},
author = {Romagnoli, Katrina M. and Boyce, Richard D. and Empey, Philip E. and Adams, Solomon and Hochheiser, Harry},
doi = {10.1016/j.ijmedinf.2015.11.015},
eprint = {15334406},
file = {:Users/solomon/Documents/Mendeley Desktop/Romagnoli et al. - 2016 - Bringing clinical pharmacogenomics information to pharmacists A qualitative study of information needs and res.pdf:pdf},
isbn = {1106401207082},
issn = {13865056},
journal = {International Journal of Medical Informatics},
month = {feb},
number = {12},
pages = {54--61},
pmid = {28642624},
title = {{Bringing clinical pharmacogenomics information to pharmacists: A qualitative study of information needs and resource requirements}},
url = {http://linkinghub.elsevier.com/retrieve/pii/S1386505615300666},
volume = {86},
year = {2016}
}
@article{Appelboom2015,
abstract = {INTRODUCTION: Aquaporin-4 (AQP4) is the prominent water-channel protein in the brain playing a critical role in controlling cell water content. After intracerebral haemorrhage (ICH), perihematomal oedema (PHE) formation leads to a rapid increase in intracranial pressure (ICP) after the initial bleed. We sought to investigate the effect of a common genomic variant in the AQP4 gene on PHE formation after ICH. METHODS: We reviewed the literature and identified a candidate polymorphism in AQP4 genes previously reported in Genome Wide Association Studies (GWAS). Between February 2009 and March 2011, 128 patients consented to genetic testing and were genotyped for single nucleotide polymorphism (SNP) on the AQP4 gene. Genomic DNA was extracted from buccal swabs using MasterAmp extraction kits (Epicentre, Madison, WI, USA). DNA extracted from buffy coats of whole blood samples was amplified via PCR. Linear regression with log-transformed ICH + PHE volume as the response variable was used to determine the association of SNP controlled for admission variables age, GCS, infratentorial location, hypertension, systolic blood pressure (SBP), blood urea nitrogen (BUN), glucose and alkaline phosphatase. RESULTS: Nine of 128 patients had the minor allele for SNP rs1058427. Presence of the minor allele was significant in the model (P = 0.021), and associated with an increase of 88{\%} in ICH + PHE volume (beta = 0.632, exp(beta) = 1.88) after controlling for admission variables. The only other significant variables included in the model was GCS (P {\textless} 0.001). CONCLUSION: The establishment of an independent association between rs1054827 and ICH + PHE volume provides evidence implicating the AQP4 gene in haematoma and oedema formation after ICH. Further investigation is needed to characterise this link.},
author = {Appelboom, Geoff and Bruce, Sam and Duren, Andrew and Piazza, Matt and Monahan, Aimee and Christophe, Brandon and Zoller, Steve and LoPresti, Melissa and Connolly, E Sander},
doi = {10.1179/1743132815Y.0000000047},
file = {:Users/solomon/Documents/Mendeley Desktop/Appelboom et al. - 2015 - Aquaporin-4 gene variant independently associated with oedema after intracerebral haemorrhage.pdf:pdf},
isbn = {0161-6412},
issn = {1743-1328 (Electronic)},
journal = {Neurological research},
keywords = {Aquaporin 4,Brain Edema,Cerebral Hemorrhage,Female,Gene Frequency,Genotyping Techniques,Humans,Linear Models,Male,Middle Aged,Polymorphism, Single Nucleotide,Prospective Studies,complications,etiology,genetics,pathology,physiopathology},
number = {8},
pages = {657--661},
pmid = {26000774},
title = {{Aquaporin-4 gene variant independently associated with oedema after intracerebral haemorrhage.}},
volume = {37},
year = {2015}
}
@article{Baber2015,
abstract = {The objective of this study was to examine interindividual variability in codeine requirements and pain management by examining select genetic polymorphisms in the codeine pharmacological pathway. The study included a nested cohort of 98 women who were prescribed codeine following cesarean section. Participants were genotyped for select polymorphisms of the COMT, ABCB1, CYP2D6, UGT2B7 and OPRM1 genes and instructed to describe their level of pain using the visual analog scale (mm) 1 h following each dose of codeine. Analysis revealed that reported pain increases with maternal age (P = 0.041). Asians required more codeine than Caucasians (P = 0.048). Significant differences in mean dose consumption were seen among the genotypic groups of the OPRM1 A118G (P = 0.001) and UGT2B7 C802T (P = 0.015) variants. These variants were found to predict codeine consumption in the cohort overall (P = 0.000) and among Caucasians (P = 0.001). These findings will assist in customizing therapy to effectively manage postpartum pain.},
author = {Baber, M and Chaudhry, S and Kelly, L and Ross, C and Carleton, B and Berger, H and Koren, G},
doi = {10.1038/tpj.2015.3},
file = {:Users/solomon/Documents/Mendeley Desktop/Baber et al. - 2015 - The pharmacogenetics of codeine pain relief in the postpartum period.pdf:pdf},
issn = {1470-269X},
journal = {Pharmacogenomics J.},
month = {oct},
number = {5},
pages = {430--435},
title = {{The pharmacogenetics of codeine pain relief in the postpartum period}},
url = {https://www-nature-com.pitt.idm.oclc.org/articles/tpj20153.pdf http://www.nature.com/articles/tpj20153},
volume = {15},
year = {2015}
}
@article{Tayag1996,
abstract = {Following traumatic brain injury (TBI), cortical and thalamic areas were analyzed histologically and by high-performance liquid chromatography with electrochemical detection for uric acid at various survival times. Following TBI, cortical uric acid was elevated by ten-fold at 24 and 48 h, but not at 1 h post-TBI. Histological evidence of neurodegeneration was found not only in cortex but also in the anteroventral thalamus. These data suggest that as in stroke, uric acid measurements may be a convenient and sensitive method for measuring peroxidative status in TBI.},
author = {Tayag, Emilio C. and Nair, Somnath N. and Wahhab, Samina and Katsetos, Christos D. and Lighthall, James W. and Lehmann, John C.},
doi = {10.1016/0006-8993(96)00669-5},
file = {:Users/solomon/Documents/Mendeley Desktop/Tayag et al. - 1996 - Cerebral uric acid increases following experimental traumatic brain injury in rat.pdf:pdf},
issn = {00068993},
journal = {Brain Research},
keywords = {Cortex,Microdialysis,Rat,Thalamus,Traumatic brain injury,Uric acid},
number = {2},
pages = {287--291},
pmid = {8891312},
title = {{Cerebral uric acid increases following experimental traumatic brain injury in rat}},
volume = {733},
year = {1996}
}
@article{Olivecrona2013,
abstract = {OBJECT: The aim of the study was to evaluate the early changes in pituitary hormone levels after severe traumatic brain injury (sTBI) and compare hormone levels to basic neuro-intensive care data, a systematic scoring of the CT-findings and to evaluate whether hormone changes are related to outcome.$\backslash$n$\backslash$nMETHODS: Prospective study, including consecutive patients, 15-70 years, with sTBI, Glasgow Coma Scale (GCS) score ≤ 8, initial cerebral perfusion pressure {\textgreater} 10 mm Hg, and arrival to our level one trauma university hospital within 24 hours after head trauma (n = 48). Serum samples were collected in the morning (08-10 am) day 1 and day 4 after sTBI for analysis of cortisol, growth hormone (GH), prolactin, insulin-like growth factor 1 (IGF-1), thyroid-stimulating hormone (TSH), free triiodothyronine (fT3), free thyroxine (fT4), follicular stimulating hormone (FSH), luteinizing hormone (LH), testosterone and sex hormone-binding globulin (SHBG) (men). Serum for cortisol and GH was also obtained in the evening (17-19 pm) at day 1 and day 4. The first CT of the brain was classified according to Marshall. Independent staff evaluated outcome at 3 months using GOS-E.$\backslash$n$\backslash$nRESULTS: Profound changes were found for most pituitary-dependent hormones in the acute phase after sTBI, i.e. low levels of thyroid hormones, strong suppression of the pituitary-gonadal axis and increased levels of prolactin. The main findings of this study were: 1) A large proportion (54{\%} day 1 and 70{\%} day 4) of the patients showed morning s-cortisol levels below the proposed cut-off levels for critical illness related corticosteroid insufficiency (CIRCI), i.e. {\textless}276 nmol/L (=10 ug/dL), 2) Low s-cortisol was not associated with higher mortality or worse outcome at 3 months, 3) There was a significant association between early (day 1) and strong suppression of the pituitary-gonadal axis and improved survival and favorable functional outcome 3 months after sTBI, 4) Significantly lower levels of fT3 and TSH at day 4 in patients with a poor outcome at 3 months. 5) A higher Marshall CT score was associated with higher day 1 LH/FSH- and lower day 4 TSH levels 6) In general no significant correlation between GCS, ICP or CPP and hormone levels were detected. Only ICPmax and LH day 1 in men was significantly correlated.$\backslash$n$\backslash$nCONCLUSION: Profound dynamic changes in hormone levels are found in the acute phase of sTBI. This is consistent with previous findings in different groups of critically ill patients, most of which are likely to be attributed to physiological adaptation to acute illness. Low cortisol levels were a common finding, and not associated with unfavorable outcome. A retained ability to a dynamic hormonal response, i.e. fast and strong suppression of the pituitary-gonadal axis (day 1) and ability to restore activity in the pituitary-thyroid axis (day 4) was associated with less severe injury according to CT-findings and favorable outcome.},
author = {Olivecrona, Zandra and Dahlqvist, Per and Koskinen, Lars-Owe D},
doi = {10.1186/1757-7241-21-33},
file = {:Users/solomon/Documents/Mendeley Desktop/Olivecrona, Dahlqvist, Koskinen - 2013 - Acute neuro-endocrine profile and prediction of outcome after severe brain injury.pdf:pdf},
issn = {1757-7241},
journal = {Scandinavian journal of trauma, resuscitation and emergency medicine},
keywords = {Adolescent,Adult,Aged,Brain Injuries,Brain Injuries: blood,Brain Injuries: radiography,Female,Growth Hormone,Growth Hormone: blood,Humans,Hydrocortisone,Hydrocortisone: blood,Hypothalamo-Hypophyseal System,Hypothalamo-Hypophyseal System: physiopathology,Insulin-Like Growth Factor I,Insulin-Like Growth Factor I: analysis,Luteinizing Hormone,Luteinizing Hormone: blood,Male,Middle Aged,Pituitary Hormones,Pituitary Hormones: blood,Pituitary-Adrenal System,Pituitary-Adrenal System: physiopathology,Prognosis,Prolactin,Prolactin: blood,Prospective Studies,Sex Hormone-Binding Globulin,Sex Hormone-Binding Globulin: analysis,Testosterone,Testosterone: blood,Thyrotropin,Thyrotropin: blood,Treatment Outcome,Young Adult},
pages = {33},
pmid = {23601250},
title = {{Acute neuro-endocrine profile and prediction of outcome after severe brain injury.}},
url = {http://www.pubmedcentral.nih.gov/articlerender.fcgi?artid=3637196{\&}tool=pmcentrez{\&}rendertype=abstract},
volume = {21},
year = {2013}
}
@misc{Adams2014a,
author = {Adams, Solomon M and Empey, Philip E},
title = {{Experimental TBI in Pediatric Abcg2 Knock-Out Rats}},
year = {2014}
}
@article{Kochanek2008,
author = {Kochanek, Patrick M and Berger, Rachel P and Bayr, H{\"{u}}lya and Wagner, Amy K and Jenkins, Larry W and Clark, Robert SB},
doi = {10.1097/MCC.0b013e3282f57564},
issn = {1070-5295},
journal = {Current Opinion in Critical Care},
month = {apr},
number = {2},
pages = {135--141},
title = {{Biomarkers of primary and evolving damage in traumatic and ischemic brain injury: diagnosis, prognosis, probing mechanisms, and therapeutic decision making}},
url = {https://insights.ovid.com/crossref?an=00075198-200804000-00004},
volume = {14},
year = {2008}
}
@article{Mi2011,
abstract = {Dopamine is a catecholamine neurotransmitter and controls a variety of functions including cognition, emotion, locomotor activity, food intake, and endocrine system regulation in the central nervous system [1]. In the periphery, dopamine modulates cardiovascular and renal functions, hormone secretion, and gastrointestinal motility [1]. Several pathological conditions, such as Parkinson's disease (PD), schizophrenia, restless leg syndrome, and endocrine tumors, for example, pituitary adenomas have been linked to dysregulation of dopaminergic signal transmission [1,2]. The dopamine D2 receptor (DRD2) is one of the five different dopamine receptors that have been identified in humans, and shows high expression in both the pituitary gland and the central nervous system [3]. The dopamine receptors belong to the family of G protein coupled receptors. They are grouped into D1-like receptors, including DRD1 and DRD5, generally associated with stimulatory functions, and D2-like receptors, including DRD2, DRD3, and DRD4, generally associated with inhibitory functions [1]. The distinct subfamily of G protein coupled receptors, to which DRD2, DRD3, and DRD4 belong, was derived likely from gene duplication before the vertebrate expansion; these receptors are also known to share similar pharmacological profiles [4,5]. In},
archivePrefix = {arXiv},
arxivId = {NIHMS150003},
author = {Mi, H and Thomas, PD and Ring, HZ and Jiang, R and Sangkuhl, K and Klein, TE and Altman, RB},
doi = {10.1097/FPC.0b013e32833ee605},
eprint = {NIHMS150003},
file = {:Users/solomon/Documents/Mendeley Desktop/Mi et al. - 2011 - PharmGKB summary dopamine receptor D2.pdf:pdf},
isbn = {2156623929},
issn = {1744-6880},
journal = {Pharmacogenetics and genomics},
keywords = {colorectal tumor,hepatoma,hsv,immunotherapy,oncolytic therapy},
month = {jun},
number = {6},
pages = {350--6},
pmid = {20736885},
title = {{PharmGKB summary: dopamine receptor D2.}},
url = {http://linkinghub.elsevier.com/retrieve/pii/S0039606006007173 http://www.ncbi.nlm.nih.gov/pubmed/20736885 http://www.pubmedcentral.nih.gov/articlerender.fcgi?artid=PMC3091980},
volume = {21},
year = {2011}
}
@article{Liu2017,
abstract = {The ATP-binding cassette sub-family G member 2 (ABCG2) plays an important role in modulating drug disposition and endobiotic homeostasis. KO143 is a potent and relatively selective ABCG2 inhibitor. We found that the metabolic stability of KO143 was very poor in human liver microsomes (HLM). Our further studies illustrated that the tert-butyl ester group in KO143 can be rapidly hydrolyzed and removed by carboxylesterase 1. This metabolic pathway was confirmed as a major pathway of KO143 metabolism in both HLM and mice. K1 is an analog of KO143 without the ester group. We found that the metabolic stability of K1 was significantly improved in HLM when compared to KO143. These data suggest that the ester group in KO143 is the major cause of the poor metabolic stability of KO143. The data from this study can be used to guide the development of KO143 analogs with better metabolic properties.},
author = {Liu, Ke and Zhu, Junjie and Huang, Yixian and Li, Chaoyue and Lu, Jie and Sachar, Madhav and Li, Song and Ma, Xiaochao},
doi = {10.1016/j.dmpk.2017.02.003},
issn = {13474367},
journal = {Drug Metabolism and Pharmacokinetics},
month = {aug},
number = {4},
pages = {193--200},
publisher = {Elsevier},
title = {{Metabolism of KO143, an ABCG2 inhibitor}},
url = {https://www-sciencedirect-com.pitt.idm.oclc.org/science/article/pii/S1347436716300568?via{\%}3Dihub http://linkinghub.elsevier.com/retrieve/pii/S1347436716300568},
volume = {32},
year = {2017}
}
@article{Kandar2011,
abstract = {Oxidative stress has been proposed as one of the potential causes for infertility in men. Ascorbic acid and uric acid play important role in protection of spermatozoa against free radicals. A method for the simultaneous determination of ascorbic acid and uric acid in human seminal plasma using HPLC with UV detection and investigation their clinical significance as antioxidants protecting male germ cells against oxidative damage are described. Semen samples were obtained from consecutive male partners of couples presenting for a fertility evaluation. After liquefaction, the samples were centrifuged and the supernatants were diluted with dithiothreitol solution and after a filtration injected onto an analytical column. For the separation, a reverse-phase column MAG 1, 250. mm ?? 4.6 mm, Labiospher PSI 100 C18, 5??m, was used. The mixture of ethanol and 25. mmol/L sodium dihydrogenphosphate (2.5:97.5, v/v), pH 4.70 was used as a mobile phase. Analytical performance of this method is satisfactory for both ascorbic acid and uric acid: the intra-assay and inter-assay coefficients of variation were below 10{\%}. Quantitative recoveries from spiked seminal plasma were between 92.1 and 102.1{\%}. We have found no significant differences in both ascorbic acid and uric acid concentration between the smokers and non-smokers (351.0 ?? 237.9??mol/L and 323.7 ?? 99.5??mol/L vs 444.8 ?? 245.5??mol/L and 316.6 ?? 108.9??mol/L, p{\textgreater}0.05). This assay is a simple and reproducible HPLC method for the simultaneous measurement of ascorbic acid and uric acid in human seminal plasma. ?? 2011 Elsevier B.V.},
author = {Kanď{\'{a}}r, Roman and Dr{\'{a}}bkov{\'{a}}, Petra and Hampl, Radek},
doi = {10.1016/j.jchromb.2011.08.007},
file = {:Users/solomon/Documents/Mendeley Desktop/Kanď{\'{a}}r, Dr{\'{a}}bkov{\'{a}}, Hampl - 2011 - The determination of ascorbic acid and uric acid in human seminal plasma using an HPLC with UV dete.pdf:pdf},
issn = {15700232},
journal = {Journal of Chromatography B: Analytical Technologies in the Biomedical and Life Sciences},
keywords = {Ascorbic acid,Dithiothreitol,High-performance liquid chromatography with ultrav,Human seminal plasma,Protein precipitants,Uric acid},
number = {26},
pages = {2834--2839},
pmid = {21871848},
title = {{The determination of ascorbic acid and uric acid in human seminal plasma using an HPLC with UV detection}},
volume = {879},
year = {2011}
}
@article{VonGertten2005,
abstract = {BACKGROUND: Traumatic brain injury (TBI) initiates a complex sequence of destructive and neuroprotective cellular responses. The initial mechanical injury is followed by an extended time period of secondary brain damage. Due to the complicated pathological picture a better understanding of the molecular events occurring during this secondary phase of injury is needed. This study was aimed at analysing gene expression patterns following cerebral cortical contusion in rat using high throughput microarray technology with the goal of identifying genes involved in an early and in a more delayed phase of trauma, as genomic responses behind secondary mechanisms likely are time-dependent. RESULTS: Among the upregulated genes 1 day post injury, were transcription factors and genes involved in metabolism, e.g. STAT-3, C/EBP-delta and cytochrome p450. At 4 days post injury we observed increased gene expression of inflammatory factors, proteases and their inhibitors, like cathepsins, alpha-2-macroglobulin and C1q. Notably, genes with biological function clustered to immune response were significantly upregulated 4 days after injury, which was not found following 1 day. Osteopontin and one of its receptors, CD-44, were both upregulated showing a local mRNA- and immunoreactivity pattern in and around the injury site. Fewer genes had decreased expression both 1 and 4 days post injury and included genes implicated in transport, metabolism, signalling, and extra cellular matrix formation, e.g. vitronectin, neuroserpin and angiotensinogen. CONCLUSION: The different patterns of gene expression, with little overlap in genes, 1 and 4 days post injury showed time dependence in genomic responses to trauma. An early induction of factors involved in transcription could lead to the later inflammatory response with strongly upregulated CD-44 and osteopontin expression. An increased knowledge of genes regulating the pathological mechanisms in trauma will help to find future treatment targets. Since trauma is a risk factor for development of neurodegenerative disease, this knowledge may also reduce late negative effects.},
author = {von Gertten, Christina and {Flores Morales}, Amilcar and Holmin, Staffan and Mathiesen, Tiit and Nordqvist, Ann-Christin Sandberg},
doi = {10.1186/1471-2202-6-69},
file = {:Users/solomon/Documents/Mendeley Desktop/von Gertten et al. - 2005 - Genomic responses in rat cerebral cortex after traumatic brain injury.pdf:pdf},
issn = {1471-2202},
journal = {BMC neuroscience},
pages = {69},
pmid = {16318630},
title = {{Genomic responses in rat cerebral cortex after traumatic brain injury.}},
volume = {6},
year = {2005}
}
@article{Bailey-Dell2001,
abstract = {The breast cancer resistance protein (BCRP) gene, formally known as ATP-binding cassette transporter G2 (ABCG2) gene, encodes an ABC half transporter that causes resistance to certain cancer chemotherapeutic drugs when transfected and expressed in drug sensitive cancer cells. Here we report the organization of the BCRP gene, and the initial characterization of the BCRP promoter. We identified the genomic sequence of BCRP and its promoter by screening a human genomic lambda phage library, as well as a BAC library, and by searching the human genome database. The BCRP gene spans over 66 kb and consists of 16 exons and 15 introns. The exons range in size from 60 to 532 bp. The translational start site is found in the second exon. The first exon contains the majority of the 5' UTR. Promoter activity was characterized by a luciferase reporter assay using transient transfection of the human breast cancer cell line MCF7, and the human choriocarcinoma cell lines JAR, BeWo and JEG-3, which we find to have high endogenous expression of BCRP. The BCRP gene is transcribed by a TATA-less promoter with several putative Sp1 sites, which are downstream from a putative CpG island. The sequence 312 bp directly upstream from the BCRP transcriptional start site conferred basal promoter activity. The 5' region upstream of the basal promoter is characterized by both positive and negative regulatory domains.},
author = {Bailey-Dell, K J and Hassel, B and Doyle, L a and Ross, D D},
doi = {S0167478101002706 [pii]},
file = {:Users/solomon/Documents/Mendeley Desktop/Bailey-Dell et al. - 2001 - Promoter characterization and genomic organization of the human breast cancer resistance protein (ATP-bindin.pdf:pdf},
isbn = {0006-3002 (Print)$\backslash$n0006-3002 (Linking)},
issn = {0006-3002},
journal = {Biochim Biophys Acta},
keywords = {abc,atp-binding cassette transporter g2,bcrp,breast cancer resistance protein,certain cancer chemotherapeutic,choriocarcinoma,described atp-binding cassette,genomic organization,half transporter,is a recently,promoter region,sp1,that causes resistance to},
number = {3},
pages = {234--241},
pmid = {7},
title = {{Promoter characterization and genomic organization of the human breast cancer resistance protein (ATP-binding cassette transporter G2) gene.}},
url = {http://eutils.ncbi.nlm.nih.gov/entrez/eutils/elink.fcgi?dbfrom=pubmed{\&}id=11566359{\&}retmode=ref{\&}cmd=prlinks{\%}5Cnpapers2://publication/uuid/42E630F0-1C51-4149-86F3-0B823CE904AF},
volume = {1520},
year = {2001}
}
@article{Mega2008,
abstract = {Clopidogrel requires transformation into an active metabolite by cytochrome P-450 (CYP) enzymes for its antiplatelet effect. The genes encoding CYP enzymes are polymorphic, with common alleles conferring reduced function.},
author = {Mega, Jessica L and Mega, Jessica L and Close, Sandra L and Close, Sandra L and Wiviott, Stephen D and Wiviott, Stephen D and Shen, Lei and Shen, Lei and Hockett, Richard D and Hockett, Richard D and Brandt, John T and Brandt, John T and Walker, Joseph R and Walker, Joseph R and Antman, Elliott M and Antman, Elliott M and Macias, William and Macias, William and Braunwald, Eugene and Braunwald, Eugene and Sabatine, Marc S and Sabatine, Marc S},
doi = {10.1056/NEJMoa0809171},
isbn = {1533-4406 (Electronic) 0028-4793 (Linking)},
issn = {1533-4406},
journal = {Society},
keywords = {Acute Coronary Syndrome,Acute Coronary Syndrome: therapy,Adult,Angioplasty,Area Under Curve,Aryl Hydrocarbon Hydroxylases,Aryl Hydrocarbon Hydroxylases: genetics,Balloon,Cardiovascular Diseases,Cardiovascular Diseases: epidemiology,Cardiovascular Diseases: genetics,Combined Modality Therapy,Coronary,Female,Genetic,Genotype,Heterozygote,Humans,Male,Mutation,Platelet Aggregation,Platelet Aggregation Inhibitors,Platelet Aggregation Inhibitors: adverse effects,Platelet Aggregation Inhibitors: metabolism,Platelet Aggregation Inhibitors: pharmacology,Platelet Aggregation Inhibitors: therapeutic use,Platelet Aggregation: drug effects,Polymorphism,Randomized Controlled Trials as Topic,Stents,Thrombosis,Thrombosis: epidemiology,Thrombosis: genetics,Ticlopidine,Ticlopidine: adverse effects,Ticlopidine: analogs {\&} derivatives,Ticlopidine: metabolism,Ticlopidine: pharmacology,Ticlopidine: therapeutic use},
number = {4},
pages = {1--9},
pmid = {19106084},
title = {{Cytochrome P-450 Polymorphisms and Response to Clopidogrel}},
url = {http://www.ncbi.nlm.nih.gov/pubmed/19106084},
volume = {360},
year = {2008}
}
@article{Medina-Pulido2013,
abstract = {Under several adverse conditions, such as hypoxia or ischaemia, extracellular levels of adenosine are elevated because of increased energy demands and ATP metabolism. Because extracellular adenosine affects metabolism through G-protein-coupled receptors, its regulation is of high adaptive importance. CNT2 (concentrative nucleoside transporter 2) may play physiological roles beyond nucleoside salvage in brain as it does in other tissues. Even though nucleoside transport in brain has mostly been seen as being of equilibrative-type, in the present study, we prove that the rat phaeochromocytoma cell line PC12 shows a concentrative adenosine transport of CNT2-type when cells are differentiated to a neuronal phenotype by treatment with NGF (nerve growth factor). Differentiation of PC12 cells was also associated with the up-regulation of adenosine A1 receptors. Addition of adenosine receptor agonists to cell cultures increased CNT2-related activity by a mechanism consistent with A₁ and A2A receptor activation. The addition of adenosine to the culture medium also induced the phosphorylation of the intracellular regulatory kinase AMPK (AMP-activated protein kinase), with this effect being dependent upon adenosine transport. CNT2-related activity of differentiated PC12 cells was also dramatically down-regulated under hypoxic conditions. Interestingly, the analysis of nucleoside transporter expression after experimental focal ischaemia in rat brain showed that CNT2 expression was down-regulated in the infarcted tissue, with this effect somehow being restricted to other adenosine transporter proteins such as CNT3 and ENT1 (equilibrative nucleoside transporter 1). In summary, CNT2 is likely to modulate extracellular adenosine and cell energy balance in neuronal tissue.},
author = {Medina-Pulido, Lorena and Molina-Arcas, M{\'{i}}riam and Justicia, Carles and Soriano, Eduardo and Burgaya, Ferran and Planas, Anna M and Pastor-Anglada, Mar{\c{c}}al},
doi = {10.1042/BJ20130231},
file = {:Users/solomon/Documents/Mendeley Desktop/Medina-Pulido et al. - 2013 - Hypoxia and P1 receptor activation regulate the high-affinity concentrative adenosine transporter CNT2 in.pdf:pdf},
isbn = {1470-8728 (Electronic)$\backslash$r0264-6021 (Linking)},
issn = {1470-8728},
journal = {The Biochemical journal},
keywords = {Adenosine,Adenosine: metabolism,Animals,Carrier Proteins,Carrier Proteins: genetics,Carrier Proteins: metabolism,Cell Differentiation,Cell Hypoxia,Energy Metabolism,Gene Expression,Gene Expression Regulation,Infarction,Male,Membrane Transport Proteins,Membrane Transport Proteins: genetics,Membrane Transport Proteins: metabolism,Middle Cerebral Artery,Middle Cerebral Artery: metabolism,Neurons,Neurons: metabolism,PC12 Cells,Purinergic P1,Purinergic P1: metabolism,Rats,Receptors,Sprague-Dawley},
number = {3},
pages = {437--45},
pmid = {23819782},
title = {{Hypoxia and P1 receptor activation regulate the high-affinity concentrative adenosine transporter CNT2 in differentiated neuronal PC12 cells.}},
url = {http://www.biochemj.org/bj/454/bj4540437.htm{\%}5Cnhttp://www.biochemj.org/content/454/3/437.abstract},
volume = {454},
year = {2013}
}
@article{Stanek2012,
abstract = {To develop a benchmark measure of US physicians' level of knowledge and extent of use of pharmacogenomic testing, we conducted an anonymous, cross-sectional, fax-based, national survey. Of 397,832 physicians receiving the survey questionnaire, 10,303 (3{\%}) completed and returned it; the respondents were representative of the overall US physician population. The factors associated with the decision to test were evaluated using $\chi$(2) and multivariate logistic regression. Overall, 97.6{\%} of responding physicians agreed that genetic variations may influence drug response, but only 10.3{\%} felt adequately informed about pharmacogenomic testing. Only 12.9{\%} of physicians had ordered a test in the previous 6 months, and 26.4{\%} anticipated ordering a test in the next 6 months. Early and future adopters of testing were more likely to have received training in pharmacogenomics, but only 29.0{\%} of physicians overall had received any education in the field. Our findings highlight the need for more effective physician education on the clinical value, availability, and interpretation of pharmacogenomic tests.},
author = {Stanek, E J and Sanders, C L and Taber, K a Johansen and Khalid, M and Patel, A and Verbrugge, R R and Agatep, B C and Aubert, R E and Epstein, R S and Frueh, F W},
doi = {10.1038/clpt.2011.306},
file = {:Users/solomon/Documents/Mendeley Desktop/Stanek et al. - 2012 - Adoption of pharmacogenomic testing by US physicians results of a nationwide survey.pdf:pdf},
isbn = {0009-9236$\backslash$r1532-6535},
issn = {1532-6535},
journal = {Clinical pharmacology and therapeutics},
keywords = {Adult,Aged,Attitude of Health Personnel,Attitudes,Cross-Sectional Studies,Cross-Sectional Studies: methods,Data Collection,Data Collection: methods,Female,Genetic Testing,Genetic Testing: methods,Genetic Variation,Health Knowledge,Humans,Logistic Models,Male,Middle Aged,Multivariate Analysis,Pharmacogenetics,Physician's Practice Patterns,Practice,Questionnaires,United States,Young Adult},
number = {3},
pages = {450--458},
pmid = {22278335},
publisher = {Nature Publishing Group},
title = {{Adoption of pharmacogenomic testing by US physicians: results of a nationwide survey.}},
url = {http://www.ncbi.nlm.nih.gov/pubmed/22278335},
volume = {91},
year = {2012}
}
@article{Cherry2014,
author = {Cherry, Jonathan D and Olschowka, John A and O'Banion, M},
doi = {10.1186/1742-2094-11-98},
file = {:Users/solomon/Documents/Mendeley Desktop/Cherry, Olschowka, O'Banion - 2014 - Neuroinflammation and M2 microglia the good, the bad, and the inflamed.pdf:pdf},
isbn = {1742-2094 (Electronic)$\backslash$r1742-2094 (Linking)},
issn = {1742-2094},
journal = {Journal of Neuroinflammation},
keywords = {M2,alternative activation,m2,microglia,neuroinflammat,neuroinflammation},
number = {1},
pages = {98},
pmid = {24889886},
title = {{Neuroinflammation and M2 microglia: the good, the bad, and the inflamed}},
url = {http://jneuroinflammation.biomedcentral.com/articles/10.1186/1742-2094-11-98},
volume = {11},
year = {2014}
}
@article{Feig,
author = {Feig, Daniel I and Kang, Duk-Hee and Johnson, Richard J},
doi = {10.1056/NEJMra0800885},
file = {:Users/solomon/Documents/Mendeley Desktop/Feig, Kang, Johnson - 2008 - Uric Acid and Cardiovascular Risk.pdf:pdf},
issn = {0028-4793},
journal = {New England Journal of Medicine},
month = {oct},
number = {17},
pages = {1811--1821},
title = {{Uric Acid and Cardiovascular Risk}},
url = {https://www.ncbi.nlm.nih.gov/pmc/articles/PMC2684330/pdf/nihms109052.pdf http://www.nejm.org/doi/abs/10.1056/NEJMra0800885},
volume = {359},
year = {2008}
}
@article{Sivapackiam2016,
abstract = {Introduction: For stratification of chemotherapeutic choices, radiopharmaceuticals capable of imaging breast cancer resistance protein (BCRP/ABCG2)-mediated functional transport are desired. To accomplish this objective, Galmydar, a fluorescent and moderately hydrophobic Ga(III) cationic complex and its 67/68Ga-radiolabeled counterparts were interrogated in HEK293 cells stably transfected with BCRP and their WT counterparts transfected with empty vector. Additionally, the sensitivity and specificity of 68Ga-Galmydar to evaluate functional expression of BCRP at the blood-brain barrier (BBB) was investigated in gene-knockout mdr1a/1b(-/-) (double knockout, dKO) and mdr1a/1b(-/-)ABCG2(-/-) (triple knockout, tKO) mouse models. Methods: For radiotracer uptake assays and live cell fluorescence imaging, either 67Ga-Galmydar or its unlabeled counterpart was incubated in HEK293 cells transfected with BCRP (HEK293/BCRP) and their WT counterparts at 37 ??C under a continuous flux of CO2 (5{\%}) in the presence or absence of Ko143, a potent BCRP antagonist, and cellular uptake was measured to assess the sensitivity of Galmydar to probe BCRP-mediated functional transport activity in cellulo. For assessing the potential of Galmydar to enable diagnostic imaging of targeted tissues in vivo, the 67Ga-radiolabeled counterpart was incubated in either human serum albumin or human serum at 37 ??C and the percentage of unbound 67Ga-Galmydar was determined. To evaluate the sensitivity of 68Ga-Galmydar for molecular imaging of BCRP-mediated efflux activity in vivo, microPET/CT brain imaging was performed in dKO and tKO mice and their age-matched WT counterparts, 60 min post-intravenous injection. Results: 67Ga-Galmydar shows uptake profiles in HEK293 cells inversely proportional to BCRP expression, and antagonist (Ko143) induced accumulation in HEK293/BCRP cells, thus indicating target sensitivity and specificity. Furthermore, employing the fluorescent characteristics of Galmydar, optical imaging in HEK293/BCRP cells shows an excellent correlation with the radiotracer cellular accumulation data. 67Ga-Galmydar shows {\textgreater} 85{\%} unbound fraction and presence of parental compound in human serum. Finally, microPET/CT imaging shows higher retention of 68Ga-Galmydar in brains of dKO and tKO mice compared to their age-matched WT counterparts, 60 min post-intravenous tail-vein injection. Conclusions: Combined data indicate that Galmydar could provide a template scaffold for development of a PET tracer for imaging BCRP-mediated functional transport activity in vivo.},
author = {Sivapackiam, Jothilingam and Harpstrite, Scott E. and Prior, Julie L. and Mattingly, Stephen and Sharma, Vijay},
doi = {10.1016/j.nucmedbio.2015.12.001},
file = {:Users/solomon/Documents/Mendeley Desktop/Sivapackiam et al. - 2016 - 6768Galmydar A metalloprobe for monitoring breast cancer resistance protein (BCRP)-mediated functional trans.pdf:pdf},
isbn = {0969-8051},
issn = {18729614},
journal = {Nuclear Medicine and Biology},
keywords = {BCRP,Gallium,Metalloprobe},
number = {3},
pages = {191--197},
pmid = {26924499},
publisher = {Elsevier Inc.},
title = {{67/68Galmydar: A metalloprobe for monitoring breast cancer resistance protein (BCRP)-mediated functional transport activity}},
url = {http://dx.doi.org/10.1016/j.nucmedbio.2015.12.001},
volume = {43},
year = {2016}
}
@article{Ho,
abstract = {There is growing interest in applying machine learning methods to Electronic Medical Records (EMR). Across dif-ferent institutions, however, EMR quality can vary widely. This work investigated the impact of this disparity on the performance of three advanced machine learning algorithms: logistic regression, multilayer perceptron, and recur-rent neural network. The EMR disparity was emulated using different permutations of the EMR collected at Children's Hospital Los Angeles (CHLA) Pediatric Intensive Care Unit (PICU) and Cardiothoracic Intensive Care Unit (CTICU). The algorithms were trained using patients from the PICU to predict in-ICU mortality for patients in a held out set of PICU and CTICU patients. The disparate patient populations between the PICU and CTICU provide an estimate of generalization errors across different ICUs. We quantified and evaluated the generalization of these algorithms on varying EMR size, input types, and fidelity of data.},
author = {Ho, Long V and Ledbetter, David and Aczon, Melissa and Wetzel, Randall and The, Laura P and Whittier, Leland K},
file = {:Users/solomon/Documents/Mendeley Desktop/Ho et al. - Unknown - The Dependence of Machine Learning on Electronic Medical Record Quality.pdf:pdf},
title = {{The Dependence of Machine Learning on Electronic Medical Record Quality}},
url = {https://arxiv.org/pdf/1703.08251.pdf}
}
@article{Siddiqui2003,
abstract = {BACKGROUND One third of patients with epilepsy have drug-resistant epilepsy, which is associated with an increased risk of death and debilitating psychosocial consequences. Because this form is resistant to multiple antiepileptic drugs, the mode of resistance must be nonspecific, involving drug-efflux transporters such as ATP-binding cassette sub-family B member 1 (ABCB1, also known as MDR1 and P-glycoprotein 170). We hypothesized that the CC genotype at the ABCB1 C3435T polymorphism, which is associated with increased expression of the protein, influences the response to antiepileptic-drug treatment. METHODS ABCB1 3435 was genotyped in 315 patients with epilepsy, classified as drug-resistant in 200 and drug-responsive in 115, and 200 control subjects without epilepsy. Recently devised methods were used to control for population stratification, and linkage disequilibrium was calculated across the gene. RESULTS As compared with patients with drug-responsive epilepsy, patients with drug-resistant epilepsy were more likely to have the CC genotype at ABCB1 3435 than the TT genotype (odds ratio, 2.66; 95 percent confidence interval, 1.32 to 5.38; P=0.006). There was no genetic stratification between the two groups of patients. The polymorphism fell within an extensive block of linkage disequilibrium spanning much or all of the gene, implying that the polymorphism may not itself be causal but rather may be linked with the causal variant. CONCLUSIONS These pharmacogenomic results identify a genetic factor associated with resistance to antiepileptic drugs.},
author = {Siddiqui, A and Kerb, R and Weale, ME and Brinkmann, U and Smith, A and Goldstein, DB and Wood, NiW and Sisodiya, SM},
doi = {10.1056/NEJMoa021986},
file = {:Users/solomon/Documents/Mendeley Desktop/Siddiqui et al. - 2003 - Association of multidrug resistance in epilepsy with a polymorphism in the drug-transporter gene ABCB1.pdf:pdf},
isbn = {1533-4406},
issn = {1533-4406},
journal = {The New England journal of medicine},
number = {15},
pages = {1442--8},
pmid = {12686700},
title = {{Association of multidrug resistance in epilepsy with a polymorphism in the drug-transporter gene ABCB1.}},
url = {http://www.ncbi.nlm.nih.gov/pubmed/12686700},
volume = {348},
year = {2003}
}
@article{Oswald2007,
abstract = {Active transport across biological membranes represents a critical step in the disposition of many drugs. It is now well-established that different efflux and uptake transporters such as P-glycoprotein (P-gp), multidrug resistance-associated proteins (MRPs) or organic anion transporting polypeptides (OATPs) are involved in the overall disposition and efficacy of numerous compounds. These proteins are mainly expressed at physiological sites of drug absorption and elimination, thus leading to diminished absorption and/or increased transporter-facilitated excretion. Moreover, drug transporters are known to be of protective significance in blood-organ barriers. On the contrary, only little is known about the relevance of transporter function on drug levels within tissues and cellular compartments, i.e. the site of action for many substances. Moreover, the pharmacokinetic processing inside the cell is characterized by uptake, metabolism and elimination. It is gradually being recognized that active uptake and/or efflux transporters may modify target concentrations at the subcellular receptor sites which in turn may have an influence on drug effects. This review will summarize current knowledge about the impact of transporter proteins on drug availability within pharmacologically relevant cellular compartments and tissues as hepatocytes, enterocytes, different blood cell types, brain, and the heart with emphasis on the potential clinical significance of these transporters.},
author = {Oswald, S and Grube, M and Siegmund, W and Kroemer, H K},
doi = {10.1080/00498250701570251},
file = {:Users/solomon/Documents/Mendeley Desktop/Oswald et al. - 2007 - Transporter-mediated uptake into cellular compartments.pdf:pdf},
isbn = {0049825070157},
issn = {0049-8254},
journal = {Xenobiotica; the fate of foreign compounds in biological systems},
keywords = {blood cell transporters,brain transporters,cellular compartments,clinical significance,enterocyte transporters,heart transporters,hepatocyte transporters,tissue transporters,transporters,uptake},
number = {10-11},
pages = {1171--1195},
pmid = {17968742},
title = {{Transporter-mediated uptake into cellular compartments.}},
volume = {37},
year = {2007}
}
@article{Orford2009,
abstract = {The ATP-binding cassette (ABC) transporter 2 (ABCG2) is expressed by stem cells in many organs and in stem cells of solid tumors. These cells are isolated based on the side population (SP) phenotype, a Hoechst 3342 dye efflux property believed to be conferred by ABCG2. Because of the limitations of this approach we generated transgenic mice that express Nuclear GFP (GFPn) coupled to the Puromycin-resistance gene, under the control of ABCG2 promoter/enhancer sequences. We show that ABCG2 is expressed in neural progenitors of the developing forebrain and spinal cord and in embryonic and adult endothelial cells of the brain. Using the neurosphere assay, we isolated tripotent ABCG2-expressing neural stem cells from embryonic mouse brain. This transgenic line is a powerful tool for studying the expression of ABCG2 in many tissues and for performing functional studies in different experimental settings. ?? 2009 Elsevier Inc. All rights reserved.},
author = {Orford, Michael and Mean, Richard and Lapathitis, George and Genethliou, Nicholas and Panayiotou, Elena and Panayi, Helen and Malas, Stavros},
doi = {10.1016/j.bbrc.2009.04.089},
file = {:Users/solomon/Documents/Mendeley Desktop/Orford et al. - 2009 - Generation of an ABCG2GFPn-puro transgenic line - A tool to study ABCG2 expression in mice.pdf:pdf},
isbn = {1090-2104 (Electronic)$\backslash$r0006-291X (Linking)},
issn = {0006291X},
journal = {Biochemical and Biophysical Research Communications},
keywords = {ABCG2,Choroid plexus,Endothelial cells,Transgenic mice},
number = {2},
pages = {199--203},
pmid = {19393620},
publisher = {Elsevier Inc.},
title = {{Generation of an ABCG2GFPn-puro transgenic line - A tool to study ABCG2 expression in mice}},
url = {http://dx.doi.org/10.1016/j.bbrc.2009.04.089},
volume = {384},
year = {2009}
}
@article{Kawase2015,
abstract = {Hepatic transporters and metabolic enzymes affect drug pharmacokinetics. Limited information exists on the alteration in mRNA levels of hepatic transporters and metabolic enzymes with aging. We examined the effects of aging on the mRNA levels of representative hepatic drug transporters and metabolic enzymes by analyzing their levels in 10-, 30- and 50-week-old male and female rats. Levels of mRNA of drug transporters including multidrug resistance protein (Mdr)1a, multidrug resistance-associated protein (Mrp)2, breast cancer resistance protein (Bcrp) and organic anion-transporting polypeptide (Oatp)1a1, and the metabolic enzymes cytochrome P450 (CYP)3A1, CYP3A2 and UDP-glucuronosyltransferase (UGT)1A1 were analyzed using real-time reverse transcriptase polymerase chain reaction. The mRNA levels of transporters in male rats did not decrease with age, while the mRNA levels of Bcrp and Oatp1a1 in female rats decreased with age. The mRNA levels of CYP3A1 and CYP3A2 in male rats were higher than those in female rats. The mRNA levels of metabolic enzymes decreased with age in female but not male rats. In particular, the mRNA levels of UGT1A1 in 10-week-old female rats were higher than those in male rats. mRNA expression of hepatic transporters and metabolic enzymes are more susceptible to aging in female than male rats. The age-related decreases in the mRNA levels of Bcrp, Oatp1a1, CYP3A1 and CYP3A2 in female rats may affect the metabolism and transport of substrates. This study showed that aging affected the mRNA expression of hepatic transporters and metabolic enzymes in rats.},
author = {Kawase, Atsushi and Ito, Ayami and Yamada, Ayano and Iwaki, Masahiro},
doi = {10.1007/s13318-014-0208-7},
file = {:Users/solomon/Documents/Mendeley Desktop/Kawase et al. - 2015 - Age-related changes in mRNA levels of hepatic transporters, cytochrome P450 and UDP-glucuronosyltransferase in fe.pdf:pdf},
issn = {0378-7966},
journal = {European journal of drug metabolism and pharmacokinetics},
keywords = {Aging,Cytochrome P450,Liver,Transporter},
number = {2},
pages = {239--44},
pmid = {24899460},
publisher = {Springer International Publishing},
title = {{Age-related changes in mRNA levels of hepatic transporters, cytochrome P450 and UDP-glucuronosyltransferase in female rats.}},
url = {http://www.scopus.com/inward/record.url?eid=2-s2.0-84901732418{\&}partnerID=tZOtx3y1{\%}5Cnhttp://link.springer.com/10.1007/s13318-014-0208-7{\%}5Cnhttp://www.ncbi.nlm.nih.gov/pubmed/24899460},
volume = {40},
year = {2015}
}
@article{Ishikawa2011a,
abstract = {Photodynamic diagnosis (PDD) is a practical tool currently used in surgical operation of aggressive brain tumors, such as glioblastoma. PDD is achieved by a photon-induced physicochemical reaction which is induced by excitation of protoporphyrin IX (PpIX) exposed to light. Fluorescence-guided gross-total resection has recently been developed in PDD, where 5-aminolevulinic acid (ALA) or its ester is administered as the precursor of PpIX. ALA induces the accumulation of PpIX, a natural photo-sensitizer, in cancer cells. Recent studies provide evidence that adenosine triphosphate (ATP)-binding cassette (ABC) transporter ABCG2 plays a pivotal role in regulating the cellular accumulation of porphyrins in cancer cells and thereby affects the efficacy of PDD. Protein kinase inhibitors are suggested to potentially enhance the PDD efficacy by blocking ABCG2-mediated porphyrin efflux from cancer cells. It is of great interest to develop potent ABCG2-inhibitors that can be applied to PDD for brain tumor therapy. This review article addresses a pivotal role of human ABC transporter ABCG2 in PDD as well as a new approach of quantitative structure-activity relationship (QSAR) analysis to design potent ABCG2-inhibitors.},
author = {Ishikawa, Toshihisa and Takahashi, Kenkichi and Ikeda, Naokado and Kajimoto, Yoshinaga and Hagiya, Yuichiro and ichiro Ogura, Shun-ichiro and ichi Miyatake, Shin-ichi and Kuroiwa, Toshihiko},
doi = {10.3390/pharmaceutics3030615},
file = {:Users/solomon/Documents/Mendeley Desktop/Ishikawa et al. - 2011 - Transporter-mediated drug interaction strategy for 5-aminolevulinic acid (ALA)-based photodynamic diagnosis of.pdf:pdf},
isbn = {8145503922},
issn = {1999-4923},
journal = {Pharmaceutics},
keywords = {5-aminolevulinic acid (ALA),ABC transporter,ABCG2,Brain tumor,Glioma,Photodynamic diagnosis,Protein kinase inhibitor,brain tumor,glioma,photodynamic diagnosis,protein kinase inhibitor},
month = {sep},
number = {3},
pages = {615--635},
pmid = {27721322},
title = {{Transporter-mediated drug interaction strategy for 5-aminolevulinic acid (ALA)-based photodynamic diagnosis of malignant brain tumor: Molecular design of ABCG2 inhibitors}},
url = {http://www.ncbi.nlm.nih.gov/pubmed/27721322 http://www.mdpi.com/1999-4923/3/3/615},
volume = {3},
year = {2011}
}
@article{Johnson2012,
author = {Johnson, J A and Roden, D M and Lesko, L J and Ashley, E and Klein, T E and Shuldiner, A R},
doi = {10.1038/clpt.2012.21},
file = {:Users/solomon/Documents/Mendeley Desktop/Johnson et al. - 2012 - Clopidogrel A Case for Indication-Specific Pharmacogenetics.pdf:pdf},
issn = {0009-9236},
journal = {Clinical Pharmacology {\&} Therapeutics},
number = {5},
pages = {774--776},
title = {{Clopidogrel: A Case for Indication-Specific Pharmacogenetics}},
url = {http://doi.wiley.com/10.1038/clpt.2012.21},
volume = {91},
year = {2012}
}
@article{Adnan2016,
abstract = {A decision forest is an ensemble of decision trees, and it is often built to discover more patterns (i.e. logic rules) and predict/classify class values more accurately than a single decision tree. Existing decision forest algorithms are typically used for building huge numbers of decision trees, involving large memory and computational overhead, in order to achieve high accuracy. Generally, many of the trees do not contribute to improving the ensemble accuracy of a forest. As a result, ensemble pruning algorithms aim to get rid of those trees while generating a subforest in order to achieve higher (or comparable) ensemble accuracy than the original forest. The objectives are two fold: select as small number of trees as possible, and maintain the ensemble accuracy of the subforest as high as possible. An optimal subforest can be found by exhaustive search; however it is not practical for any standard-sized forest as the number of candidate subforests grows exponentially. In order to avoid the computational burden of an exhaustive search, many greedy and genetic algorithm-based subforest selection techniques have been proposed in literature. In this paper, we propose a subforest selection technique that achieves small size as well as high accuracy. We use a genetic algorithm where we carefully select high quality individual trees for the initial population of the genetic algorithm in order to improve the final output of the algorithm. Experiments are conducted on 20 data sets from the UCI Machine Learning Repository to compare the proposed technique with several existing state-of-the-art techniques. The results indicate that the proposed technique can select effective subforests which are significantly smaller than original forests while achieving better (or comparable) accuracy than the original forests.},
author = {Adnan, Md Nasim and Islam, Md Zahidul},
doi = {10.1016/j.knosys.2016.07.016},
file = {:Users/solomon/Documents/Mendeley Desktop/Adnan, Islam - 2016 - Optimizing the number of trees in a decision forest to discover a subforest with high ensemble accuracy using a ge.pdf:pdf},
issn = {09507051},
journal = {Knowledge-Based Systems},
month = {oct},
pages = {86--97},
publisher = {Elsevier},
title = {{Optimizing the number of trees in a decision forest to discover a subforest with high ensemble accuracy using a genetic algorithm}},
url = {https://www.sciencedirect.com/science/article/pii/S0950705116302301 http://linkinghub.elsevier.com/retrieve/pii/S0950705116302301},
volume = {110},
year = {2016}
}
@article{Qi2014,
abstract = {Glutathione-complexed [2Fe-2S] cluster is shown to significantly stimulate the ATPase activity of an ABCB7-type transporter in both solution and proteoliposome-bound forms (KD approximately 68 muM). The cluster is a likely natural substrate for this transporter, which has been implicated in cytosolic Fe-S cluster protein maturation. A possible substrate-binding site is identified on a new structural model for the active transporter.},
author = {Qi, W and Li, J and Cowan, J A},
doi = {10.1039/c3cc48239a},
file = {:Users/solomon/Documents/Mendeley Desktop/Qi, Li, Cowan - 2014 - A structural model for glutathione-complexed iron-sulfur cluster as a substrate for ABCB7-type transporters.pdf:pdf},
isbn = {1364-548X (Electronic)$\backslash$r1359-7345 (Linking)},
issn = {1364-548X},
journal = {Chem Commun (Camb)},
number = {29},
pages = {3795--3798},
pmid = {24584132},
title = {{A structural model for glutathione-complexed iron-sulfur cluster as a substrate for ABCB7-type transporters}},
url = {http://www.ncbi.nlm.nih.gov/pubmed/24584132},
volume = {50},
year = {2014}
}
@article{Hu2014,
abstract = {This study evaluated the developmental gene and protein expression of proton-coupled oligopeptide transporters (POTs: peptide transporter, PepT1 and PepT2; peptide-histidine transporter, PhT1 and PhT2) in different regions of rodent brain, and the age-dependent uptake of a POT substrate, glycylsarcosine (GlySar), in brain slices. Slices were obtained from cerebral cortex, cerebellum and hippocampus of wildtype and PepT2 null mice, and from rats at different ages. Gene and protein expression were determined by real-time PCR and immunoblot analyses. Brain slice uptakes of radiolabeled glycylsarcosine were determined in the absence and presence of excess unlabeled glycylsarcosine or l-histidine, the latter being an inhibitor of PhT1/2 but not PepT1/2. As PepT2 and PhT1 transcripts were abundantly expressed in all three regions of mouse brain, little to no expression was observed for PepT1 and PhT2. PhT1 protein was present in brain regions of adult but not neonatal mice and expression levels increased with age in rats. Glycylsarcosine uptake, inhibition and transporter dominance did not show regional brain or species differences. However, there were clear age-related differences in functional activity, with PepT2 dominating in neonatal mice and rats, and PhT1 dominating in adult rodents. These developmental changes may markedly impact the neural activity of both endogenous and exogenous (drug) peptides/mimetics. Developmental gene and protein expression of peptide transporters was evaluated in various regions of rodent brain, along with age-dependent uptake of dipeptide. We found marked changes in protein expression and functional activity of PhT1 and PepT2, the former predominating in adult and the latter in neonate. These developmental changes may markedly impact the neural activity of endogenous and exogenous peptides/mimetics.},
author = {Hu, Yongjun and Xie, Yehua and Keep, Richard F. and Smith, David E.},
doi = {10.1111/jnc.12687},
file = {:Users/solomon/Documents/Mendeley Desktop/Hu et al. - 2014 - Divergent developmental expression and function of the proton-coupled oligopeptide transporters PepT2 and PhT1 in reg.pdf:pdf},
isbn = {7346471431},
issn = {14714159},
journal = {Journal of Neurochemistry},
keywords = {PepT2,PhT1,brain,development,expression,function},
number = {6},
pages = {955--965},
pmid = {24548120},
title = {{Divergent developmental expression and function of the proton-coupled oligopeptide transporters PepT2 and PhT1 in regional brain slices of mouse and rat}},
volume = {129},
year = {2014}
}
@misc{Technologies2014,
author = {Technologies, Live},
booktitle = {Clinical Biochemistry},
file = {:Users/solomon/Documents/Mendeley Desktop/Technologies - 2014 - Amplex {\textregistered} Red Uric Acid Uricase Assay Kit ( A22181 ).pdf:pdf},
title = {{Amplex {\textregistered} Red Uric Acid / Uricase Assay Kit ( A22181 )}},
year = {2014}
}
@article{Bowman2010a,
abstract = {Oxidative damage is a consistent finding in a number of central nervous system (CNS) disorders. Uric acid (UA) is a potent hydrophilic antioxidant that is modified by diet and drug. Several lines of evidence suggest that plasma UA may modulate outcomes in neurologic disease, but little attention has been paid to CNS levels of UA. Our objective was to test the hypothesis that cerebrospinal fluid (CSF) UA is determined by plasma UA, modified by blood-brain barrier (BBB) integrity and associated with rate of cognitive decline in Alzheimer's disease (AD). Also, since UA and ascorbic acid may act as antioxidants for one another, we also explored a potential interaction between them in the brain. Thirty-two patients with mild to moderate AD (Mini-Mental Status Exam 19 +/- 5) participated in a longitudinal biomarker study for one year involving standardized clinical assessments. CSF and blood were collected at baseline for UA, ascorbic acid, and albumin. Cognitive measures were collected at baseline and again one year later. CSF UA was independent of age, gender, and AD severity. CSF and plasma UA were positively correlated (r=0.669, p=0.001) and BBB impairment was associated with higher CSF levels of UA (p=0.028). Neither plasma nor CSF UA reached significant association with rates of cognitive decline over 1 year. CSF UA and CSF ascorbic acid were positively correlated (r=0.388, p=0.001). The hypothesis that CSF UA is determined by plasma UA and BBB integrity is supported, as is the hypothesis that UA and ascorbic acid are associated in CSF but not plasma. Adequately powered prospective studies would help assess any role for UA in primary and secondary prevention of AD.},
author = {Bowman, Gene L. and Shannon, Jackilen and Frei, Balz and Kaye, Jeffrey A. and Quinn, Joseph F.},
doi = {10.3233/JAD-2010-1330},
file = {:Users/solomon/Documents/Mendeley Desktop/Bowman et al. - 2010 - Uric acid as a CNS antioxidant.pdf:pdf},
isbn = {1875-8908 (Electronic)$\backslash$r1387-2877 (Linking)},
issn = {13872877},
journal = {Journal of Alzheimer's Disease},
keywords = {Alzheimer's disease,Ascorbic acid,Blood-brain barrier,Cerebrospinal fluid,Uric acid},
number = {4},
pages = {1331--1336},
pmid = {20061611},
title = {{Uric acid as a CNS antioxidant}},
volume = {19},
year = {2010}
}
@article{Kikuchi2013,
abstract = {P-glycoprotein (P-gp) is expressed at the blood-brain barrier (BBB) and restricts the penetration of its substrates into the central nervous system (CNS). In vitro substrate assessment for P-gp is frequently used to predict the in vivo relevance of P-gp-mediated efflux at the BBB. We have conducted a comprehensive review of literature focusing on the in vitro-in vivo correlation of P-gp efflux ratio (ER), and demonstrated that in vitro substrates of P-gp are also in vivo substrates at the BBB. It was of note that the in vitro ER in the MDCK-MDR1 cell line from National Institutes of Health was found to be a better predictor of in vivo ER than that from Netherlands Cancer Institute, with r(2) values of 0.813 and 0.531, respectively. Recently, a research group proposed that 98{\%} of Biopharmaceutics Drug Disposition Classification System (BDDCS) class 1 drugs can penetrate the brain even when those compounds are shown as P-gp substrates in vitro. However, our data analysis suggested that in vitro ER can predict the in vivo brain penetration regardless of the class in BDDCS. Considering that very few marketed CNS drugs are in vivo substrates for P-gp, the in vitro substrate assessment of P-gp should be used in the early stages of drug discovery to select compounds that are most likely to penetrate the CNS to exert their pharmacologic action.},
author = {Kikuchi, Ryota and {De Morais}, Sonia M. and Kalvass, J. Cory},
doi = {10.1124/dmd.113.053868},
isbn = {1521-009X (Electronic)$\backslash$n0090-9556 (Linking)},
issn = {00909556},
journal = {Drug Metabolism and Disposition},
number = {12},
pages = {2012--2017},
pmid = {24009309},
title = {{In vitro P-glycoprotein efflux ratio can predict the in vivo brain penetration regardless of biopharmaceutics drug disposition classification system class}},
volume = {41},
year = {2013}
}
@article{Krishnamurthy2005,
abstract = {ABC (ATP-binding cassette) transporters have diverse roles in many cellular processes. These diverse roles require the presence of conserved membrane spanning domains and nucleotide binding domains. Bcrp (Abcg2) is a member of the ATP binding cassette family of plasma membrane transporters that was originally discovered for its ability to confer drug resistance in tumor cells. Subsequent studies showed Bcrp expression in normal tissues and high expression in primitive stem cells. Bcrp expression is induced under low oxygen conditions consistent with its high expression in tissues exposed to low oxygen environments. Moreover, Bcrp interacts with heme and other porphyrins. This finding and its regulation by hypoxia suggests it may play a role in protecting cells/tissue from protoporphyrin accumulation under hypoxia. These observations are strengthened by the fact that porphyrins accumulate in tissues of the Bcrp knockout mouse. It is possible that humans with loss of function Bcrp alleles may be more susceptible to porphyrin-induced phototoxicity. We propose that Bcrp plays a role in porphyrin homoeostasis and regulates survival under low oxygen conditions.},
author = {Krishnamurthy, Partha and Schuetz, John D.},
doi = {10.1007/s10534-005-3709-7},
isbn = {0966-0844 (Print)},
issn = {09660844},
journal = {BioMetals},
keywords = {ABC-transporter,BCRP,Heme,Hypoxia,Porphyrins},
number = {4},
pages = {349--358},
pmid = {16158227},
title = {{The ABC transporter Abcg2/Bcrp: Role in hypoxia mediated survival}},
volume = {18},
year = {2005}
}
@article{Verbelen2017,
abstract = {Pharmacogenetics (PGx) has the potential to personalize pharmaceutical treatments. Many relevant gene-drug associations have been discovered, but PGx guided treatment needs to be cost-effective as well as clinically beneficial to be incorporated into standard healthcare. Progress in this area can be assessed by reviewing economic evaluations to determine the cost-effectiveness of PGx testing versus standard treatment. We performed a review of economic evaluations for PGx associations listed in the US Food and Drug Administration (FDA) Table of Pharmacogenomic Biomarkers in Drug Labeling (http://www.fda.gov/Drugs/ScienceResearch/ResearchAreas/Pharmacogenetics/ucm083378.htm). We determined the proportion of evaluations that found PGx guided treatment to be cost-effective or dominant over the alternative strategies, and we estimated the impact on this proportion of removing the cost of genetic testing. Of the 130 PGx associations in the FDA table, 44 economic evaluations, relating to 10 drugs, were identified. Of these evaluations, 57{\%} drew conclusions in favour of PGx testing, of which 30{\%} were cost-effective and 27{\%} were dominant (cost-saving). If genetic information was freely available, 75{\%} of economic evaluations would support PGx guided treatment, of which 25{\%} would be cost-effective and 50{\%} would be dominant. Thus, PGx guided treatment can be a cost-effective and even cost-saving strategy. Having genetic information readily available in the clinical health record is a realistic future prospect, and would make more genetic tests economically worthwhile. However, few drugs with PGx associations have been studied and more economic evaluations are needed to underpin the uptake of genetic testing in clinical practice.},
author = {Verbelen, M. and Weale, M. E. and Lewis, C. M.},
doi = {10.1038/tpj.2017.21},
file = {:Users/solomon/Documents/Mendeley Desktop/Verbelen, Weale, Lewis - 2017 - Cost-effectiveness of pharmacogenetic-guided treatment Are we there yet.pdf:pdf},
issn = {14731150},
journal = {Pharmacogenomics Journal},
number = {5},
pages = {395--402},
pmid = {28607506},
publisher = {Nature Publishing Group},
title = {{Cost-effectiveness of pharmacogenetic-guided treatment: Are we there yet?}},
url = {http://dx.doi.org/10.1038/tpj.2017.21},
volume = {17},
year = {2017}
}
@article{Cui2014,
abstract = {Traumatic brain injury (TBI) is a leading cause of mortality and$\backslash$ndisability in children and young adults worldwide. Neurologic impairment$\backslash$nis caused by both immediate brain tissue disruption and post-injury$\backslash$ncellular and molecular events that worsen the primary neurologic insult.$\backslash$nThe beta-lactam antibiotic ceftriaxone (CTX) has been reported to induce$\backslash$nneuroprotection in animal models of diverse neurologic diseases via$\backslash$nup-regulation of GLT-1. However, no studies have addressed the$\backslash$nneuroprotective role of CTX in the setting of TBI, and whether the$\backslash$nmechanism is involved in the modulation of neuronal autophagy remains$\backslash$ntotally unclear. The present study was designed to determine the$\backslash$nhypothesis that administration of CTX could significantly enhance$\backslash$nfunctional recovery in a rat model of TBI and whether CTX treatment$\backslash$ncould up-regulate GLT-1 expression and suppress post-TBI neuronal$\backslash$nautophagy. The results demonstrated that daily treatment with CTX$\backslash$nattenuated TBI-induced brain edema and cognitive function deficits in$\backslash$nrats. GLT-1 is down-regulated following TBI and this phenomenon can be$\backslash$nreversed by treatment of CTX. In addition, we also found that CTX$\backslash$nsignificantly reduced autophagy marker protein, LC3 II, in hippocampus$\backslash$ncompared to the TBI group. These results suggest that CTX might provide$\backslash$na new therapeutic strategy for TBI and this protection might be$\backslash$nassociated with up-regulation of GLT-1 and suppression of neuronal$\backslash$nautophagy.},
author = {Cui, Changmeng and Cui, Ying and Gao, Junling and Sun, Liqian and Wang, Yongchao and Wang, Kaijie and Li, Ran and Tian, Yanxia and Song, Sixin and Cui, Jianzhong},
doi = {10.1007/s10072-013-1585-4},
file = {:Users/solomon/Documents/Mendeley Desktop/Cui et al. - 2014 - Neuroprotective effect of ceftriaxone in a rat model of traumatic brain injury.pdf:pdf},
issn = {15903478},
journal = {Neurological Sciences},
keywords = {Autophagy,Ceftriaxone,Functional recovery,GLT-1,Glutamate excitotoxicity,TBI},
number = {5},
pages = {695--700},
title = {{Neuroprotective effect of ceftriaxone in a rat model of traumatic brain injury}},
volume = {35},
year = {2014}
}
@article{Clark2017,
abstract = {Background},
author = {Clark, Robert S B and Empey, Philip E and {Lya Bayır}, H{\"{u}} and Rosario, Bedda L and Poloyac, Samuel M and Kochanek, Patrick M and Nolin, Thomas D and Au, Alicia K and Horvat, Christopher M and Wisniewski, Stephen R and Bell, Michael J},
doi = {10.1371/journal.pone.0180280},
file = {:Users/solomon/Documents/Mendeley Desktop/Clark et al. - 2017 - Phase I randomized clinical trial of N- acetylcysteine in combination with an adjuvant probenecid for treatment of.pdf:pdf},
isbn = {1111111111},
number = {November 2011},
pages = {1--16},
title = {{Phase I randomized clinical trial of N- acetylcysteine in combination with an adjuvant probenecid for treatment of severe traumatic brain injury in children}},
url = {http://journals.plos.org/plosone/article/file?id=10.1371/journal.pone.0180280{\&}type=printable},
year = {2017}
}
@article{Singh2010a,
author = {Singh, A. and Wu, H. and Zhang, P. and Happel, C. and Ma, J. and Biswal, S.},
doi = {10.1158/1535-7163.MCT-10-0108},
file = {:Users/solomon/Documents/Mendeley Desktop/Singh et al. - 2010 - Expression of ABCG2 (BCRP) Is Regulated by Nrf2 in Cancer Cells That Confers Side Population and Chemoresistance P.pdf:pdf},
issn = {1535-7163},
journal = {Molecular Cancer Therapeutics},
keywords = {abcg2,cancer stem cells,chemo-resistance,lung cancer,nrf2,rnai},
month = {aug},
number = {8},
pages = {2365--2376},
title = {{Expression of ABCG2 (BCRP) Is Regulated by Nrf2 in Cancer Cells That Confers Side Population and Chemoresistance Phenotype}},
url = {http://mct.aacrjournals.org/cgi/doi/10.1158/1535-7163.MCT-10-0108},
volume = {9},
year = {2010}
}
@article{Werner2007,
abstract = {The knowledge of the pathophysiology after traumatic head injury is necessary for adequate and patient-oriented treatment. As the primary insult, which represents the direct mechanical damage, cannot be therapeutically influenced, target of the treatment is the limitation of the secondary damage (delayed non-mechanical damage). It is influenced by changes in cerebral blood flow (hypo- and hyperperfusion), impairment of cerebrovascular autoregulation, cerebral metabolic dysfunction and inadequate cerebral oxygenation. Furthermore, excitotoxic cell damage and inflammation may lead to apoptotic and necrotic cell death. Understanding the multidimensional cascade of secondary brain injury offers differentiated therapeutic options.},
archivePrefix = {arXiv},
arxivId = {READ USE parts},
author = {Werner, C and Engelhard, K},
doi = {10.1093/bja/aem131},
eprint = {READ USE parts},
file = {:Users/solomon/Documents/Mendeley Desktop/Werner, Engelhard - 2007 - Pathophysiology of traumatic brain injury.pdf:pdf},
isbn = {0007-0912 (Print)},
issn = {00070912},
journal = {British Journal of Anaesthesia},
keywords = {Brain,Complications,Head,Inflammation,Pathophysiology,cerebral blood flow,trauma,vasospasm},
number = {1},
pages = {4--9},
pmid = {17573392},
title = {{Pathophysiology of traumatic brain injury}},
type = {Journal Article},
url = {https://www.ncbi.nlm.nih.gov/pubmed/17573392},
volume = {99},
year = {2007}
}
@article{Diaz-Arrastia2014,
abstract = {Despite substantial investments by government, philanthropic, and commercial sources over the past several decades, traumatic brain injury (TBI) remains an unmet medical need and a major source of disability and mortality in both developed and developing societies. The U.S. Department of Defense neurotrauma research portfolio contains more than 500 research projects funded at more than {\$}700 million and is aimed at developing interventions that mitigate the effects of trauma to the nervous system and lead to improved quality of life outcomes. A key area of this portfolio focuses on the need for effective pharmacological approaches for treating patients with TBI and its associated symptoms. The Neurotrauma Pharmacology Workgroup was established by the U.S. Army Medical Research and Materiel Command (USAMRMC) with the overarching goal of providing a strategic research plan for developing pharmacological treatments that improve clinical outcomes after TBI. To inform this plan, the Workgroup (a) assessed the current state of the science and ongoing research and (b) identified research gaps to inform future development of research priorities for the neurotrauma research portfolio. The Workgroup identified the six most critical research priority areas in the field of pharmacological treatment for persons with TBI. The priority areas represent parallel efforts needed to advance clinical care; each requires independent effort and sufficient investment. These priority areas will help the USAMRMC and other funding agencies strategically guide their research portfolios to ensure the development of effective pharmacological approaches for treating patients with TBI.},
author = {Diaz-Arrastia, R and Kochanek, PM and Bergold, P and Kenney, K and Marx, CE and Grimes, Col JB and Loh, LT C Y and Adam, LT C GE and Oskvig, D and Curley, KC and Salzer, W},
doi = {10.1089/neu.2013.3019},
file = {:Users/solomon/Documents/Mendeley Desktop/Diaz-Arrastia et al. - 2014 - Pharmacotherapy of traumatic brain injury state of the science and the road forward report of the Departme.pdf:pdf},
isbn = {0897-7151},
issn = {1557-9042 (Electronic) 0897-7151 (Linking)},
journal = {J Neurotrauma},
keywords = {animal studies,head trauma,human studies,pharmacology,traumatic brain injury},
number = {2},
pages = {135--158},
pmid = {23968241},
title = {{Pharmacotherapy of traumatic brain injury: state of the science and the road forward: report of the Department of Defense Neurotrauma Pharmacology Workgroup}},
type = {Journal Article},
url = {https://www.ncbi.nlm.nih.gov/pubmed/23968241 http://www.ncbi.nlm.nih.gov/pubmed/23968241{\%}5Cnhttp://www.pubmedcentral.nih.gov/articlerender.fcgi?artid=PMC3900003},
volume = {31},
year = {2014}
}
@article{Long1996,
author = {Long, D.A. and Ghosh, Kamalika and Moore, A.N. and Dixon, C.E. and Dash, P.K.},
file = {:Users/solomon/Documents/Mendeley Desktop/Long et al. - 1996 - Deferoxamine improves spatial memory performance following experimental brain injury in rats.pdf:pdf},
journal = {Brain research},
keywords = {brain injury,deferoxamine,free radical,spatial memory},
number = {1-2},
pages = {109--117},
title = {{Deferoxamine improves spatial memory performance following experimental brain injury in rats}},
url = {http://www.sciencedirect.com/science/article/pii/0006899395015000},
volume = {717},
year = {1996}
}
@article{To2013,
abstract = {BACKGROUND AND PURPOSE: Multidrug resistance (MDR), usually mediated by overexpression of efflux transporters such as P-gp, ABCG2 and/or MRP1, remains a major obstacle hindering successful cancer chemotherapy. There has been great interest in the development of inhibitors towards these transporters to circumvent resistance. However, since the inhibition of transporter is not specific to cancer cells, a decrease in the cytotoxic drug dosing may be needed to prevent excess toxicity, thus undermining the potential benefit brought about by a drug efflux inhibitor. The design of potent MDR modulators specific towards resistant cancer cells and devoid of drug-drug interactions will be needed to effect MDR reversal.$\backslash$n$\backslash$nEXPERIMENTAL APPROACH: Recent evidence suggests that the PTEN/PI3K/Akt pathway may be exploited to alter ABCG2 subcellular localization, thereby circumventing MDR. Three PPAR$\gamma$ agonists (telmisartan, pioglitazone and rosiglitazone) that have been used in the clinics were tested for their effect on the PTEN/PI3K/Akt pathway and possible reversal of ABCG2-mediated drug resistance.$\backslash$n$\backslash$nKEY RESULTS: The PPAR$\gamma$ agonists were found to be weak ABCG2 inhibitors by drug efflux assay. They were also shown to elevate the reduced PTEN expression in a resistant and ABCG2-overexpressing cell model, which inhibit the PI3K-Akt pathway and lead to the relocalization of ABCG2 from the plasma membrane to the cytoplasma, thus apparently circumventing the ABCG2-mediated MDR.$\backslash$n$\backslash$nCONCLUSIONS AND IMPLICATIONS: Since this PPAR$\gamma$/PTEN/PI3K/Akt pathway regulating ABCG2 is only functional in drug-resistant cancer cells with PTEN loss, the PPAR$\gamma$ agonists identified may represent promising agents targeting resistant cells for MDR reversal.},
author = {To, Kenneth K W and Tomlinson, Brian},
doi = {10.1111/bph.12367},
file = {:Users/solomon/Documents/Mendeley Desktop/To, Tomlinson - 2013 - Targeting the ABCG2-overexpressing multidrug resistant (MDR) cancer cells by PPAR$\gamma$ agonists.pdf:pdf},
isbn = {0007-1188},
issn = {00071188},
journal = {British Journal of Pharmacology},
keywords = {ABCG2,PPAR$\gamma$,multidrug resistance (MDR),phosphatase and tensin homologue deleted on chromo,pioglitazone,rosiglitazone,telmisartan},
number = {5},
pages = {1137--1151},
pmid = {24032744},
title = {{Targeting the ABCG2-overexpressing multidrug resistant (MDR) cancer cells by PPAR$\gamma$ agonists}},
volume = {170},
year = {2013}
}
@article{Thompson2015,
abstract = {BACKGROUND Head injury is a common event and can cause a spectrum of motor and cognition disabilities. A frequent complication is seizures. Antiepileptic drugs (AED) such as phenytoin are often used in clinical practice with the hopes of preventing post-traumatic epilepsy. Whether immediate medical intervention following head trauma with either AEDs or neuroprotective drugs can alter the process of epileptogenesis and lead to a more favorable outcome is currently unknown. This review attempted to address the effectiveness of these treatment interventions. This review updates and expands on the earlier Cochrane review. OBJECTIVES To compare the efficacy of antiepileptic drugs and neuroprotective agents with placebo, usual care or other pharmacologic agents for the prevention of post-traumatic epilepsy in people diagnosed with any severity of traumatic brain injury. SEARCH METHODS We searched The Cochrane Epilepsy Group's specialized register, CENTRAL, MEDLINE, ClinicalTrials.gov and World Health Organization International Clinical Trials Registry Platform (ICTRP) in January 2015. We searched EMBASE, Biological Abstracts and National Research Register in September 2014 and SCOPUS in December 2013. The Cochrane Epilepsy Group performed handsearches of relevant journals. SELECTION CRITERIA We included randomized controlled trials (RCTs) that include AEDs or neuroprotective agents compared with placebo, another pharmacologic agent or a usual care group. The outcomes measured included a seizure occurring within one week of trauma (early seizure), seizure occurring later than one week post-trauma (late seizure), mortality and any adverse events. DATA COLLECTION AND ANALYSIS Two review authors independently assessed study quality and extracted the data. We calculated risk ratios (RR) and 95{\%} confidence intervals (CI) for each outcome. We used random-effects models in the meta-analyses and performed pre-defined subgroup and sensitivity analyses. MAIN RESULTS This review included 10 RCTs (reported in 12 articles) consisting of 2326 participants The methodological quality of the studies varied. The type of intervention was separated into three categories; AED versus placebo or standard care, alternative neuroprotective agent versus placebo or standard care and AED versus other AED. Treatment with an AED (phenytoin or carbamazepine) decreased the risk of early seizure compared with placebo or standard care (RR 0.42, 95{\%} CI 0.23 to 0.73; very low quality evidence). There was no evidence of a difference in the risk of late seizure occurrence between AEDs and placebo or standard care (RR 0.91, 95{\%} CI 0.57 to 1.46; very low quality evidence). There was no evidence of a significant difference in all-cause mortality between AEDs and placebo or standard care (RR 1.08 95{\%} CI 0.79 to 1.46,very low quality of evidence). Only one study looked at other potentially neuroprotective agents (magnesium sulfate) compared with placebo. The risk ratios were: late seizure 1.07 (95{\%} CI 0.53 to 2.17) and all-cause mortality 1.20 (95{\%} CI 0.80 to 1.81). The risk ratio for occurrence of early seizure was not estimable.Two studies looked at comparison of two AEDs (levetiracetam, valproate) with phenytoin used as the main comparator in each study. The risk ratio for all-cause mortality was 0.53 (95{\%} CI 0.30 to 0.94). There was no evidence of treatment benefit of phenytoin compared with another AED for early seizures (RR 0.66, 95{\%} 0.20 to 2.12) or late seizures(RR 0.77, 95{\%} CI 0.46 to 1.30).Only two studies reported adverse events. The RR of any adverse event with AED compared with placebo was 1.65 (95{\%} CI 0.73 to 3.66; low quality evidence). There were insufficient data on adverse events in the other treatment comparisons. AUTHORS' CONCLUSIONS This review found low-quality evidence that early treatment with an AED compared with placebo or standard care reduced the risk of early post-traumatic seizures. There was no evidence to support a reduction in the risk of late seizures or mortality. There was insufficient evidence to make any conclusions regarding the effectiveness or safety of other neuroprotective agents compared with placebo or for the comparison of phenytoin, a traditional AED, with another AED.},
author = {Thompson, K and Pohlmann-Eden, B and Campbell, LA and Abel, H},
doi = {10.1002/14651858.CD009900.pub2},
issn = {1469-493X},
journal = {The Cochrane database of systematic reviews},
month = {aug},
number = {8},
pages = {CD009900},
pmid = {26259048},
title = {{Pharmacological treatments for preventing epilepsy following traumatic head injury.}},
url = {http://www.ncbi.nlm.nih.gov/pubmed/26259048},
year = {2015}
}
@article{Lodato2015,
abstract = {Neurons live for decades in a postmitotic state, their genomes susceptible to DNA damage. Here we survey the landscape of somatic single-nucleotide variants (SNVs) in the human brain. We identified thousands of somatic SNVs by single-cell sequencing of 36 neurons from the cerebral cortex of three normal individuals. Unlike germline and cancer SNVs, which are often caused by errors in DNA replication, neuronal mutations appear to reflect damage during active transcription. Somatic mutations create nested lineage trees, allowing them to be dated relative to developmental landmarks and revealing a polyclonal architecture of the human cerebral cortex. Thus, somatic mutations in the brain represent a durable and ongoing record of neuronal life history, from development through postmitotic function.},
archivePrefix = {arXiv},
arxivId = {15334406},
author = {Lodato, M. A. and Woodworth, M. B. and Lee, S. and Evrony, G. D. and Mehta, B. K. and Karger, A. and Chittenden, T. W. and D'Gama, A. M. and Cai, X. and Luquette, L. J. and Lee, E. and Park, P. J. and Walsh, C. A.},
doi = {10.1126/science.aab1785},
eprint = {15334406},
file = {:Users/solomon/Documents/Mendeley Desktop/Lodato et al. - 2015 - Somatic mutation in single human neurons tracks developmental and transcriptional history.pdf:pdf},
isbn = {1095-9203 (Electronic)$\backslash$r0036-8075 (Linking)},
issn = {0036-8075},
journal = {Science},
keywords = {Adolescent,Cell Lineage,Cerebral Cortex,Cerebral Cortex: cytology,Cerebral Cortex: growth {\&} development,DNA Mutational Analysis,DNA Replication,DNA Replication: genetics,Female,Genetic,Genetic Loci,Humans,Male,Mitosis,Mitosis: genetics,Mutation,Neurons,Neurons: cytology,Neurons: physiology,Polymorphism,Single Nucleotide,Single-Cell Analysis,Transcription},
number = {6256},
pages = {94--98},
pmid = {26430121},
title = {{Somatic mutation in single human neurons tracks developmental and transcriptional history}},
url = {http://www.ncbi.nlm.nih.gov/pubmed/26430121},
volume = {350},
year = {2015}
}
@article{Balding2006,
abstract = {Although genetic association studies have been with us for many years, even for the simplest analyses there is little consensus on the most appropriate statistical procedures. Here I give an overview of statistical approaches to population association studies, including preliminary analyses (Hardy-Weinberg equilibrium testing, inference of phase and missing data, and SNP tagging), and single-SNP and multipoint tests for association. My goal is to outline the key methods with a brief discussion of problems (population structure and multiple testing), avenues for solutions and some ongoing developments.},
author = {Balding, David J and Balding, David J},
doi = {10.1038/nrg1916},
file = {:Users/solomon/Documents/Mendeley Desktop/Balding, Balding - 2006 - A tutorial on statistical methods for population association studies.pdf:pdf},
isbn = {1471-0056},
issn = {1471-0056},
journal = {Nature reviews. Genetics},
keywords = {Genetics, Medical,Genetics, Medical: methods,Genetics, Population,Genetics, Population: methods,Genome, Human,Genome, Human: genetics,Genotype,Haplotypes,Humans,Polymorphism, Single Nucleotide,Polymorphism, Single Nucleotide: genetics,Statistics as Topic,Statistics as Topic: methods},
number = {10},
pages = {781--91},
pmid = {16983374},
title = {{A tutorial on statistical methods for population association studies.}},
url = {http://www.ncbi.nlm.nih.gov/pubmed/16983374},
volume = {7},
year = {2006}
}
@article{Zutz2009,
abstract = {ABC transporters represent one of the largest families of membrane proteins that are found in all three phyla of life. Mitochondria comprise up to four ABC systems, ABCB7/ATM1, ABCB10/MDL1, ABCB8 and ABCB6. These half-transporters, which assemble into homodimeric complexes, are involved in a number of key cellular processes, e.g. biogenesis of cytosolic iron-sulfur clusters, heme biosynthesis, iron homeostasis, multidrug resistance, and protection against oxidative stress. Here, we summarize recent advances and emerging themes in our understanding of how these ABC systems in the inner and outer mitochondrial membrane fulfill their functions in important (patho) physiological processes, including neurodegenerative and hematological disorders. ?? 2009 Elsevier B.V. All rights reserved.},
author = {Zutz, Ariane and Gompf, Simone and Sch{\"{a}}gger, Hermann and Tamp{\'{e}}, Robert},
doi = {10.1016/j.bbabio.2009.02.009},
file = {:Users/solomon/Documents/Mendeley Desktop/Zutz et al. - 2009 - Mitochondrial ABC proteins in health and disease.pdf:pdf},
isbn = {0006-3002; 0006-3002},
issn = {00052728},
journal = {Biochimica et Biophysica Acta - Bioenergetics},
keywords = {Heme biosynthesis,Intracellular transport,Iron homeostasis,Iron-sulfur cluster assembly,Mitochondrial targeting,Oxidative stress},
number = {6},
pages = {681--690},
pmid = {19248758},
publisher = {Elsevier B.V.},
title = {{Mitochondrial ABC proteins in health and disease}},
url = {http://dx.doi.org/10.1016/j.bbabio.2009.02.009},
volume = {1787},
year = {2009}
}
@article{Mills2013,
abstract = {Pharmacogenetic testing refers to a type of genetic test to predict a patient's likelihood to experience an adverse event or not respond to a given drug. Despite revision to several labels of commonly prescribed drugs regarding the impact of genetic variation, the use of this testing has been limited in many settings due to a number of factors. In the primary care setting, the limited office time as well as the limited knowledge and experience of primary care practitioners have likely attributed to the slow uptake of pharmacogenetic testing. This paper provides talking points for primary care physicians to discuss with patients when pharmacogenetic testing is warranted. As patients and physicians become more familiar and accepting of pharmacogenetic testing, it is anticipated that discussion time will be comparable to that of other clinical tests.},
author = {Mills, Rachel and Voora, Deepak and Peyser, Bruce and Haga, Susanne},
doi = {10.2147/PGPM.S50598},
file = {:Users/solomon/Documents/Mendeley Desktop/Mills et al. - 2013 - Delivering pharmacogenetic testing in a primary care setting.pdf:pdf},
issn = {1178-7066},
journal = {Pharmacogenomics and Personalized Medicine},
keywords = {patient education,pharmacogenetic testing,pharmacogenetics,primary care},
month = {sep},
pages = {105},
pmid = {24101877},
publisher = {Dove Press},
title = {{Delivering pharmacogenetic testing in a primary care setting}},
url = {http://www.ncbi.nlm.nih.gov/pubmed/24101877 http://www.pubmedcentral.nih.gov/articlerender.fcgi?artid=PMC3791676 http://www.dovepress.com/delivering-pharmacogenetic-testing-in-a-primary-care-setting-peer-reviewed-article-PGPM},
volume = {6},
year = {2013}
}
@article{Zhao2011,
abstract = {BACKGROUND AND PURPOSE: Brain iron overload plays a detrimental role in brain injury after intracerebral hemorrhage (ICH). A recent study found that minocycline acts as an iron chelator and reduces iron-induced neuronal death in vitro. The present study investigated if minocycline reduces iron overload after ICH and iron-induced brain injury in vivo. METHODS: This study was divided into four parts. (1) Rats with different sizes of ICH were euthanized 3 days later for serum total iron and brain edema determination. (2) Rats had an ICH treated with minocycline or vehicle. Serum iron, brain iron, and brain iron handling proteins were measured. (3) Rats had an intracaudate injection of either saline, iron, iron+minocycline or iron+macrophage/microglia inhibitory factor and were used for brain edema and neuronal death measurements. (4) Rats had an intracaudate injection of iron and were treated with minocycline. The brains were used for edema measurement. RESULTS: After ICH, serum total iron and brain non-heme iron increased and these changes were reduced by minocycline treatment. Minocycline also reduced ICH-induced upregulation of brain iron handling proteins and neuronal death. Intracaudate injection of iron caused brain edema, blood-brain barrier leakage and brain cell death, all of which were significantly reduced by co-injection with minocycline. CONCLUSIONS: The current study found that minocycline reduces iron overload after ICH and iron-induced brain injury. It is also well known minocycline is an inhibitor of microglial activation. Minocycline may be very useful for ICH patients because both iron accumulation and microglia activation contribute to brain damage following ICH. },
archivePrefix = {arXiv},
arxivId = {NIHMS150003},
author = {Zhao, Fan and Hua, Ya and He, Yangdong and Keep, Richard F. and Xi, Guohua},
doi = {10.1161/STROKEAHA.111.623926},
eprint = {NIHMS150003},
file = {:Users/solomon/Documents/Mendeley Desktop/Zhao et al. - 2011 - Minocycline-induced attenuation of iron overload and brain injury after experimental intracerebral hemorrhage.pdf:pdf},
isbn = {8585348585},
issn = {00392499},
journal = {Stroke},
keywords = {Brain edema,Cerebral hemorrhage,Iron,Minocycline},
number = {12},
pages = {3587--3593},
pmid = {3226873},
title = {{Minocycline-induced attenuation of iron overload and brain injury after experimental intracerebral hemorrhage}},
volume = {42},
year = {2011}
}
@article{Zhao2014,
author = {Zhao, Qin and Sun, Jianguo and Tao, Yifu and Wang, Shukui and Jiang, Chunzhi and Zhu, Yubing and Yu, Feng and Zhu, Junrong},
doi = {10.2217/pgs.14.22},
file = {:Users/solomon/Documents/Mendeley Desktop/Zhao et al. - 2014 - A logistic equation to determine the validity of tramadol from related gene polymorphisms and psychological factors.pdf:pdf},
issn = {1462-2416},
journal = {Pharmacogenomics},
month = {mar},
number = {4},
pages = {487--495},
title = {{A logistic equation to determine the validity of tramadol from related gene polymorphisms and psychological factors}},
url = {https://www-futuremedicine-com.pitt.idm.oclc.org/doi/pdf/10.2217/pgs.14.22 http://www.futuremedicine.com/doi/10.2217/pgs.14.22},
volume = {15},
year = {2014}
}
@article{Riches2009,
abstract = {Gout is a common rheumatic disease in humans which is characterized by elevation in serum uric acid levels, and deposition of uric acid crystals in the joint. Hyperuricaemia is the primary risk factor for the development of gout and primates have uniquely high levels of serum uric acid due to missense mutations in the uricase gene. Levels of serum uric acid are known to be highly heritable, and mutations in genes which encode enzymes in the purine salvage pathway have long been recognized as rare causes of gout. Until recently, however, little has been known about the genetic determinants of urate metabolism and susceptibility to gout in the general population. Over recent months, a series of large scale genome wide association studies have been performed which have shed new light on the genes which regulate serum uric acid levels and susceptibility to gout. Most of these genes seem to be involved in regulating the renal excretion of uric acid which underscores the importance of reduced urate excretion as opposed to increased endogenous production as a cause of gout. Further work will now be required to investigate the mechanisms by which these genetic variants regulate urate excretion and serum urate levels. However, it seems likely that the genes so far identified will represent new molecular targets for the design of drugs to enhance urate excretion and the genetic variants that predispose to gout might be of value as genetic markers of susceptibility to gout.},
author = {Riches, Philip L. and Wright, Alan F. and Ralston, Stuart H.},
doi = {10.1093/hmg/ddp369},
file = {:Users/solomon/Documents/Mendeley Desktop/Riches, Wright, Ralston - 2009 - Recent insights into the pathogenesis of hyperuricaemia and gout.pdf:pdf},
isbn = {1316511022},
issn = {09646906},
journal = {Human Molecular Genetics},
number = {R2},
pages = {177--184},
pmid = {19808794},
title = {{Recent insights into the pathogenesis of hyperuricaemia and gout}},
volume = {18},
year = {2009}
}
@article{Yamamoto2013,
author = {Yamamoto, Kazuko and Pepper-cunningham, Zachary A and Ferrari, Joseph D and Wilson, Andrew A and Jones, Matthew R and Quinton, Lee J and Mizgerd, Joseph P and Microbiology, Molecular and Sciences, Biomedical},
doi = {10.1165/rcmb.2012-0347OC},
file = {:Users/solomon/Documents/Mendeley Desktop/Yamamoto et al. - 2013 - Page 1 of 53.pdf:pdf},
isbn = {4072376329},
number = {412},
pages = {1--53},
title = {{Page 1 of 53}},
year = {2013}
}
@article{Wagner2012,
abstract = {BACKGROUND: Studies implicate single nucleotide polymorphism (SNP) rs17070145, a common T → C polymorphism on the KIBRA gene, in mediating differences in episodic memory. In healthy adults, T-allele carriers perform better than non-carriers on episodic memory measures. However, this association is reversed in adults with subjective memory complaints and populations vulnerable to memory deficits, a problem common in traumatic brain injury (TBI).$\backslash$n$\backslash$nMETHODS: This study assessed associations between variation in the KIBRA gene and cognitive function in 129 adults with severe TBI. In addition to other executive functioning and functional/global outcomes, the Buschke Selective Reminding Test (SRT), Rey-Osterrieth Complex Figure Test and California Verbal Learning Test-II (CVLT-II) were administered 6 and 12 months post-injury.$\backslash$n$\backslash$nRESULTS: T-allele non-carriers performed better than carriers on multiple episodic memory measures. At 6 months, T-allele non-carriers performed better for delayed recall measures on the SRT. At 12 months, T-allele non-carriers performed better on multiple SRT measures and on List-B learning with CVLT-II. No associations occurred with executive function or global outcome measures.$\backslash$n$\backslash$nCONCLUSION: These results suggest that rs17070145 T-allele effects are specific to episodic memory and support the hypothesis that associations between rs17070145 variation and memory are disparate between healthy and impaired populations.},
author = {Wagner, AK and Hatz, LE and Scanlon, JM and Niyonkuru, C and Miller, MA and Ricker, JH and Conley, YP and Ferrell, RE},
doi = {10.3109/02699052.2012.700089},
file = {:Users/solomon/Documents/Mendeley Desktop/Wagner et al. - 2012 - Association of KIBRA rs17070145 polymorphism and episodic memory in individuals with severe TBI.pdf:pdf},
isbn = {4126486666},
issn = {1362-301X},
journal = {Brain injury},
keywords = {Adolescent,Adult,Aged,Brain Injuries,Brain Injuries: psychology,Cognition,Cognition Disorders,Cognition Disorders: etiology,Cognition Disorders: genetics,Cognition Disorders: psychology,Episodic,Female,Genotype,Humans,Intracellular Signaling Peptides and Proteins,Intracellular Signaling Peptides and Proteins: gen,Intracellular Signaling Peptides and Proteins: met,Male,Memory,Middle Aged,Neuropsychological Tests,Phosphoproteins,Phosphoproteins: genetics,Phosphoproteins: metabolism,Polymorphism,Risk Factors,Single Nucleotide,Time Factors,Young Adult},
number = {13-14},
pages = {1658--69},
pmid = {22794909},
title = {{Association of KIBRA rs17070145 polymorphism and episodic memory in individuals with severe TBI.}},
url = {http://www.ncbi.nlm.nih.gov/pubmed/22794909},
volume = {26},
year = {2012}
}
@article{Romagnoli2017a,
author = {Romagnoli, Katrina M and Boyce, Richard D and Empey, Philip E and Ning, Yifan and Adams, Solomon and Hochheiser, Harry},
doi = {10.1093/jamia/ocx007},
file = {:Users/solomon/Documents/Mendeley Desktop/Romagnoli et al. - 2017 - Design and evaluation of a pharmacogenomics information resource for pharmacists.pdf:pdf},
issn = {1067-5027},
journal = {Journal of the American Medical Informatics Association},
keywords = {as pharmacogenomics-guided drug decision-making,background and significance,becomes more,clinical decision making,clinically actionable pharma-,common,evaluation,pharmacist,pharmacists seek trustworthy,pharmacogenomics,structured product labeling,usability},
number = {0},
pages = {1--10},
title = {{Design and evaluation of a pharmacogenomics information resource for pharmacists}},
url = {https://academic.oup.com/jamia/article/3053492/Design},
volume = {0},
year = {2017}
}
@article{Skrobik2013,
abstract = {BACKGROUND Delirium and sedative-induced coma are described as incremental manifestations of cerebral dysfunction. Both may be associated with sedative or opiate doses and pharmacokinetic or pharmacogenetic variables, such as drug plasma levels (exposure), drug metabolism, and/or their transport across the blood-brain barrier. OBJECTIVES To compare biological and drug treatment characteristics in patients with coma and/or delirium while in the ICU. PATIENTS AND MEASUREMENTS In 99 patients receiving IV fentanyl, midazolam, or both, we evaluated drug doses, covariates likely to influence drug effects (age, body mass index, and renal and hepatic dysfunction); delirium risk factors; concomitant administration of CYP3A and P-glycoprotein substrates/inhibitors; ABCB1, ABCG2, and CYP3A5 genetic polymorphisms; and fentanyl and midazolam plasma levels. Delirium and coma were evaluated daily. In patients with only coma (n=15), only delirium (n=7), and neither ever (n=14), we measured plasma levels of tumor necrosis factor-$\alpha$, interleukin (IL)-1$\beta$, IL-1RA, IL-6, IL-8, IL-10, IL-17,macrophage inflammatory protein-1$\beta$, and monocyte chemotactic protein-1. RESULTS Time to first coma was associated with fentanyl and midazolam doses (p=0.03 and p=0.01, respectively). The number of days in coma was associated with the number of days of coadministration of CYP3A inhibitors (r=0.30; p=0.006). Plasma levels of fentanyl were higher in patients with clinical coma (3.7±4.7 vs. 2.0±1.8 ng/mL, p=0.0001) as were midazolam plasma levels (1050±2232 vs. 168±249 ng/mL, p=0.0001). Delirium occurrence was unrelated to midazolam administration, cumulative doses, or serum levels. Days with delirium were associated with days of coadministration of P-glycoprotein inhibitor (r=0.35; p=0.0004). Delirious patients had higher levels of the inflammatory mediator IL-6 than comatose patients (129.3 vs. 35.0 pg/mL, p=0.05). CONCLUSIONS Coma is associated with fentanyl and midazolam exposure; delirium is unrelated to midazolam and may be linked to inflammatory status. These data suggest that iatrogenic coma and delirium are not mechanistically linked.},
author = {Skrobik, Y and Leger, C and Cossette, M and Michaud, V and Turgeon, J},
doi = {10.1097/CCM.0b013e318275d014},
file = {:Users/solomon/Documents/Mendeley Desktop/Skrobik et al. - 2013 - Factors predisposing to coma and delirium fentanyl and midazolam exposure CYP3A5, ABCB1, and ABCG2 genetic polym.pdf:pdf},
issn = {1530-0293},
journal = {Critical care medicine},
keywords = {coma,cyp3a4,delirium,inflammatory mediators},
month = {apr},
number = {4},
pages = {999--1008},
pmid = {23385102},
title = {{Factors predisposing to coma and delirium: fentanyl and midazolam exposure; CYP3A5, ABCB1, and ABCG2 genetic polymorphisms; and inflammatory factors.}},
url = {http://content.wkhealth.com/linkback/openurl?sid=WKPTLP:landingpage{\&}an=00003246-201304000-00008 http://www.ncbi.nlm.nih.gov/pubmed/23385102},
volume = {41},
year = {2013}
}
@article{Pirmohamed2013,
abstract = {BACKGROUND: The level of anticoagulation in response to a fixed-dose regimen of warfarin is difficult to predict during the initiation of therapy. We prospectively compared the effect of genotype-guided dosing with that of standard dosing on anticoagulation control in patients starting warfarin therapy.$\backslash$n$\backslash$nMETHODS: We conducted a multicenter, randomized, controlled trial involving patients with atrial fibrillation or venous thromboembolism. Genotyping for CYP2C9*2, CYP2C9*3, and VKORC1 (-1639G→A) was performed with the use of a point-of-care test. For patients assigned to the genotype-guided group, warfarin doses were prescribed according to pharmacogenetic-based algorithms for the first 5 days. Patients in the control (standard dosing) group received a 3-day loading-dose regimen. After the initiation period, the treatment of all patients was managed according to routine clinical practice. The primary outcome measure was the percentage of time in the therapeutic range of 2.0 to 3.0 for the international normalized ratio (INR) during the first 12 weeks after warfarin initiation.$\backslash$n$\backslash$nRESULTS: A total of 455 patients were recruited, with 227 randomly assigned to the genotype-guided group and 228 assigned to the control group. The mean percentage of time in the therapeutic range was 67.4{\%} in the genotype-guided group as compared with 60.3{\%} in the control group (adjusted difference, 7.0 percentage points; 95{\%} confidence interval, 3.3 to 10.6; P{\textless}0.001). There were significantly fewer incidences of excessive anticoagulation (INR ≥4.0) in the genotype-guided group. The median time to reach a therapeutic INR was 21 days in the genotype-guided group as compared with 29 days in the control group (P{\textless}0.001).$\backslash$n$\backslash$nCONCLUSIONS: Pharmacogenetic-based dosing was associated with a higher percentage of time in the therapeutic INR range than was standard dosing during the initiation of warfarin therapy. (Funded by the European Commission Seventh Framework Programme and others; ClinicalTrials.gov number, NCT01119300.).},
author = {Pirmohamed, Munir and Burnside, Girvan and Eriksson, Niclas and Jorgensen, Andrea L and Toh, Cheng Hock and Nicholson, Toby and Kesteven, Patrick and Christersson, Christina and Wahlstr{\"{o}}m, Bengt and Stafberg, Christina and Zhang, J Eunice and Leathart, Julian B and Kohnke, Hugo and {Maitland-van der Zee}, Anke H and Williamson, Paula R and Daly, Ann K and Avery, Peter and Kamali, Farhad and Wadelius, Mia},
doi = {10.1056/NEJMoa1311386},
file = {:Users/solomon/Documents/Mendeley Desktop/Pirmohamed et al. - 2013 - A randomized trial of genotype-guided dosing of warfarin.pdf:pdf},
isbn = {0028-4793},
issn = {1533-4406},
journal = {The New England journal of medicine},
keywords = {80 and over,Adult,Aged,Anticoagulants,Anticoagulants: administration {\&} dosage,Anticoagulants: adverse effects,Aryl Hydrocarbon Hydroxylases,Aryl Hydrocarbon Hydroxylases: genetics,Atrial Fibrillation,Atrial Fibrillation: drug therapy,Cytochrome P-450 CYP2C9,Female,Genotype,Hemorrhage,Hemorrhage: chemically induced,Humans,International Normalized Ratio,Kaplan-Meier Estimate,Male,Middle Aged,Pharmacogenetics,Venous Thromboembolism,Venous Thromboembolism: drug therapy,Vitamin K Epoxide Reductases,Vitamin K Epoxide Reductases: genetics,Warfarin,Warfarin: administration {\&} dosage,Warfarin: adverse effects},
number = {24},
pages = {2294--303},
pmid = {24251363},
title = {{A randomized trial of genotype-guided dosing of warfarin.}},
url = {http://www.ncbi.nlm.nih.gov/pubmed/24251363},
volume = {369},
year = {2013}
}
@article{Motsinger2007,
abstract = {The importance of gene–gene and gene–environment interactions in the underlying genetic architecture of common, complex phenotypes is gaining wide recognition in the field of pharmacogenomics. In epidemiological approaches to mapping genetic variants that predict drug response, it is important that researchers investigate potential epistatic interactions. In the current review, we discuss data-mining tools available in genetic epidemiology to detect such interactions and appropriate applications. We survey several classes of novel methods available and present an organized collection of successful applications in the literature. Finally, we provide guidance as to how to incorporate these novel methods into a genetic analysis. The overall goal of this paper is to aid researchers in developing an analysis plan that accounts for gene–gene and gene–environment in their own work.},
author = {Motsinger, Alison A and Ritchie, Marylyn D and Reif, David M},
doi = {10.2217/14622416.8.9.1229},
file = {:Users/solomon/Documents/Mendeley Desktop/Motsinger, Ritchie, Reif - 2007 - Novel methods for detecting epistasis in pharmacogenomics studies.pdf:pdf},
issn = {1462-2416},
journal = {Pharmacogenomics},
keywords = {data-mining,epistasis,gene–environment interactions,gene–gene interactions,novel computational methods,pharmacogenomics},
month = {sep},
number = {9},
pages = {1229--1241},
title = {{Novel methods for detecting epistasis in pharmacogenomics studies}},
url = {https://www-futuremedicine-com.pitt.idm.oclc.org/doi/pdf/10.2217/14622416.8.9.1229 http://www.futuremedicine.com/doi/10.2217/14622416.8.9.1229},
volume = {8},
year = {2007}
}
@article{Yan2015,
abstract = {Previous studies have shown that iron accumulation is involved in the pathogenesis of brain injury following subarachnoid hemorrhage (SAH) and chelation of iron reduced mortality and oxidative DNA damage. We previously reported that blockage of mitochondrial calcium uniporter (MCU) provided benefit in the early brain injury after experimental SAH. This study was undertaken to identify whether blockage of MCU could ameliorate iron accumulation-associated brain injury following SAH. Therefore, we used two reagents ruthenium red (RR) and spermine (Sper) to inhibit MCU. Sprague-Dawley (SD) rats were randomly divided into four groups including sham, SAH, SAH + RR, and SAH + Sper. Biochemical analysis and histological assays were performed. The results confirmed the iron accumulation in temporal lobe after SAH. Interestingly, blockage of MCU dramatically reduced the iron accumulation in this area. The mechanism was revealed that inhibition of MCU reversed the down-regulation of iron regulatory protein (IRP) 1/2 and increase of ferritin. Iron-sulfur cluster dependent-aconitase activity was partially conserved when MCU was blocked. In consistence with this and previous report, ROS levels were notably reduced and ATP supply was rescued; levels of cleaved caspase-3 dropped; and integrity of neurons in temporal lobe was protected. Taken together, our results indicated that blockage of MCU could alleviate iron accumulation and the associated injury following SAH. These findings suggest that the alteration of calcium and iron homeostasis be coupled and MCU be considered to be a therapeutic target for patients suffering from SAH.},
author = {Yan, Huiying and Hao, Shuangying and Sun, Xiaoyan and Zhang, Dingding and Gao, Xin and Yu, Zhuang and Li, Kuanyu and Hang, Chun Hua},
doi = {10.1016/j.bbrc.2014.12.073},
file = {:Users/solomon/Documents/Mendeley Desktop/Yan et al. - 2015 - Blockage of mitochondrial calcium uniporter prevents iron accumulation in a model of experimental subarachnoid hemor.pdf:pdf},
issn = {10902104},
journal = {Biochemical and Biophysical Research Communications},
keywords = {Brain injury,Iron accumulation,Mitochondrial calcium uniporter,Subarachnoid hemorrhage},
number = {4},
pages = {835--840},
publisher = {Elsevier Inc.},
title = {{Blockage of mitochondrial calcium uniporter prevents iron accumulation in a model of experimental subarachnoid hemorrhage}},
url = {http://dx.doi.org/10.1016/j.bbrc.2014.12.073},
volume = {456},
year = {2015}
}
@article{Wagner2014a,
abstract = {Dopamine (DA) neurotransmission influences cognition and recovery after traumatic brain injury (TBI). We explored whether functional genetic variants affecting the DA transporter (DAT) and D2 receptor (DRD2) impacted in vivo dopaminergic binding with positron emission tomography (PET) using [(11)C]$\beta$CFT and [(11)C]raclopride. We examined subjects with moderate/severe TBI (N=12) ∼1 year post injury and similarly matched healthy controls (N=13). The variable number of tandem repeat polymorphism within the DAT gene and the TaqI restriction fragment length polymorphism near the DRD2 gene were assessed. TBI subjects had age-adjusted DAT-binding reductions in the caudate, putamen, and ventral striatum, and modestly increased D2 binding in ventral striatum versus controls. Despite small sample sizes, multivariate analysis showed lower caudate and putamen DAT binding among DAT 9-allele carriers and DRD2 A2/A2 homozygotes with TBI versus controls with the same genotype. Among TBI subjects, 9-allele carriers had lower caudate and putamen binding than 10/10 homozygotes. This PET study suggests a hypodopaminergic environment and altered DRD2 autoreceptor DAT interactions that may influence DA transmission after TBI. Future work will relate these findings to cognitive performance; future studies are required to determine how DRD2/DAT1 genotype and DA-ligand binding are associated with neurostimulant response and TBI recovery.},
author = {Wagner, Amy K and Scanlon, Joelle M and Becker, Carl R and Ritter, Anne C and Niyonkuru, Christian and Dixon, Clifton E and Conley, Yvette P and Price, Julie C},
doi = {10.1038/jcbfm.2014.87},
file = {:Users/solomon/Documents/Mendeley Desktop/Wagner et al. - 2014 - The influence of genetic variants on striatal dopamine transporter and D2 receptor binding after TBI.pdf:pdf},
isbn = {1559-7016 (Electronic)$\backslash$r0271-678X (Linking)},
issn = {1559-7016},
journal = {Journal of cerebral blood flow and metabolism : official journal of the International Society of Cerebral Blood Flow and Metabolism},
keywords = {Age Factors,Brain Injuries,Brain Injuries: genetics,Brain Injuries: metabolism,Brain Injuries: radionuclide imaging,Case-Control Studies,Caudate Nucleus,Caudate Nucleus: metabolism,Caudate Nucleus: radionuclide imaging,Corpus Striatum,Corpus Striatum: metabolism,Corpus Striatum: radionuclide imaging,Dopamine,Dopamine D2,Dopamine D2: genetics,Dopamine Plasma Membrane Transport Proteins,Dopamine Plasma Membrane Transport Proteins: genet,Dopamine: metabolism,Glasgow Coma Scale,Humans,Magnetic Resonance Imaging,Multivariate Analysis,Polymorphism,Positron-Emission Tomography,Protein Binding,Putamen,Putamen: metabolism,Putamen: radionuclide imaging,Receptors,Restriction Fragment Length,Tandem Repeat Sequences,Tandem Repeat Sequences: genetics},
number = {8},
pages = {1328--39},
pmid = {24849661},
publisher = {Nature Publishing Group},
title = {{The influence of genetic variants on striatal dopamine transporter and D2 receptor binding after TBI.}},
url = {http://www.ncbi.nlm.nih.gov/pubmed/24849661},
volume = {34},
year = {2014}
}
@article{Alvarez-Lario2010a,
abstract = {Uric acid (UA) is the end product of purine metabolism in humans due to the loss of uricase activity by various mutations of its gene during the Miocene epoch, which led to humans having higher UA levels than other mammals. Furthermore, 90{\%} of UA filtered by the kidneys is reabsorbed, instead of being excreted. These facts suggest that evolution and physiology have not treated UA as a harmful waste product, but as something beneficial that has to be kept. This has led various researchers to think about the possible evolutionary advantages of the loss of uricase and the subsequent increase in UA levels. It has been argued that due to the powerful antioxidant activity of UA, the evolutionary benefit could be the increased life expectancy of hominids. For other authors, the loss of uricase and the increase in UA could be a mechanism to maintain blood pressure in times of very low salt ingestion. The oldest hypothesis associates the increase in UA with higher intelligence in humans. Finally, UA has protective effects against several neurodegenerative diseases, suggesting it could have interesting actions on neuronal development and function. These hypotheses are discussed from an evolutionary perspective and their clinical signifi-cance. UA has some obvious harmful effects, and some, not so well-known, beneficial effects as an antioxidant and neuroprotector.},
author = {{\'{A}}lvarez-Lario, Bonifacio and Macarr{\'{o}}n-Vicente, Jes{\'{u}}s and Alvarez-Lario, B. and Macarron-Vicente, J.},
doi = {10.1093/rheumatology/keq204},
file = {:Users/solomon/Documents/Mendeley Desktop/{\'{A}}lvarez-Lario et al. - 2010 - Uric acid and evolution.pdf:pdf},
isbn = {1462-0332 (Electronic)$\backslash$n1462-0324 (Linking)},
issn = {1462-0324},
journal = {Rheumatology},
keywords = {Antioxidants,Evolution,Gout,Hyperuricaemia,Neurodegenerative diseases,Neuroprotection,Oxidative stress,Salt ingestion,Uric acid,Uricase},
month = {nov},
number = {11},
pages = {2010--2015},
pmid = {20627967},
title = {{Uric acid and evolution}},
url = {https://watermark.silverchair.com/keq204.pdf?token=AQECAHi208BE49Ooan9kkhW{\_}Ercy7Dm3ZL{\_}9Cf3qfKAc485ysgAAAb8wggG7BgkqhkiG9w0BBwagggGsMIIBqAIBADCCAaEGCSqGSIb3DQEHATAeBglghkgBZQMEAS4wEQQMTlQk{\_}gZb7VUnfgCQAgEQgIIBcve5PEr11qWNT-ULXJ4y84sBwfgAs81UjEZymjciTbDgB6ub},
volume = {49},
year = {2010}
}
@article{Langemann2001,
abstract = {Objectives - Microdialysis has been extensively used to monitor brain metabolism in the extracellular fluid of patients with severe head injury, to detect the onset of secondary ischaemic damage. The aim was to investigate whether concentrations of uric and ascorbic acids were altered in such patients. Both these compounds play a part in free radical metabolism, which is accelerated after ischaemia and brain injury. Methods - Patients with aneurysm or bypass operations were monitored intraoperatively to assess concentrations in minimally disturbed tissue. Afterwards, 13 patients with severe head trauma were monitored for up to 13 days in the intensive care unit. Results - Intraoperatively, concentrations of both ascorbic and uric acids were significantly higher in the bypass group than in patients with aneurysm, which might be attributed to chronic ischaemic conditions caused by the unilateral occlusion of the carotid artery. In the patients with trauma, mean values of uric acid, varying between 6 $\mu$M and 180 $\mu$M, did not correlate with type of injury (contusion or diffuse) or duration of monitoring time. Patients who died had significantly higher concentrations of uric acid than those with a good outcome. Ascorbic acid could be detected only intermittently, probably due to technical problems. Concentrations of these two compounds could not be correlated with clinical findings during the course of monitoring. Conclusions - Although uric and ascorbic acids are influenced by ischaemic conditions - for example, in bypass patients, neither compound is suitable for monitoring for free radical activity after severe head injury. Patients with a bad outcome tended to have higher concentrations of uric acid.},
author = {Langemann, H and Feuerstein, T and Mendelowitsch, A and Gratzl, O},
doi = {10.1136/jnnp.71.2.169},
file = {:Users/solomon/Documents/Mendeley Desktop/Langemann et al. - 2001 - Microdialytical monitoring of uric and ascorbic acids in the brains of patients after severe brain injury and.pdf:pdf},
issn = {0022-3050},
journal = {Journal of Neurology Neurosurgery and Psychiatry},
keywords = {adult,aged,aneurysm,article,ascorbic acid,brain artery aneurysm,brain contusion,brain injury,brain ischemia,brain metabolism,bypass surgery,carotid artery obstruction,clinical article,controlled study,extracellular fluid,female,free radical,head injury,human,intensive care,male,microdialysis,microvascular surgery,patient monitoring,priority journal,uric acid},
number = {2},
pages = {169--174},
pmid = {11459887},
title = {{Microdialytical monitoring of uric and ascorbic acids in the brains of patients after severe brain injury and during neurovascular surgery}},
url = {http://www.embase.com/search/results?subaction=viewrecord{\&}from=export{\&}id=L32675140{\%}5Cnhttp://dx.doi.org/10.1136/jnnp.71.2.169},
volume = {71},
year = {2001}
}
@article{Crews2011,
abstract = {PURPOSE: The development and implementation of a pharmacist-managed clinical pharmacogenetics service are described.$\backslash$n$\backslash$nSUMMARY: A pharmacist-managed clinical pharmacogenetics service was designed and implemented at an academic specialty hospital to provide clinical pharmacogenetic testing for gene products important to the pharmacodynamics of medications used in the hospital's patients. A series of accredited educational seminars were conducted for our pharmacists to establish competencies in providing pharmacogenetic consults for the genes to be tested by the clinical pharmacogenetics service. The service was modeled after and integrated with an already-established clinical pharmacokinetics service. A steering committee was formed to evaluate the use of available tests, new evidence for implementation of additional tests, and other service quality metrics. All clinical pharmacogenetic test results are first reported to one of the pharmacists, who reviews the result and provides a written consultation. The consultation includes an interpretation of the result and recommendations for any indicated changes to therapy. In 2009, 136 clinical pharmacogenetic tests were performed. The service has been met with positive clinician feedback. The successful implementation of this service highlights the leadership role that pharmacists can take in moving pharmacogenetics from research to patient care.$\backslash$n$\backslash$nCONCLUSION: The development of and experience with a pharmacist-managed clinical pharmacogenetics service are described. The program's success has depended on collaboration between the clinical laboratory and pharmacists, and pharmacists' pharmacogenetic recommendations have been well accepted by prescribers.},
author = {Crews, Kristine R. and Cross, Shane J. and Mccormick, John N. and Baker, Donald K. and Molinelli, Alejandro R. and Richard, Mullins and Relling, Mary V. and Hoffman, James M.},
doi = {10.2146/ajhp100113},
file = {:Users/solomon/Documents/Mendeley Desktop/Crews et al. - 2011 - Development and implementation of a pharmacist-managed clinical pharmacogenetics service.pdf:pdf},
isbn = {1535-2900 (Electronic)$\backslash$r1079-2082 (Linking)},
issn = {10792082},
journal = {American Journal of Health-System Pharmacy},
keywords = {Clinical pharmacists,Clinical pharmacy,Hospitals,Pharmaceutical services,Pharmacogenetics},
number = {2},
pages = {143--150},
pmid = {21200062},
title = {{Development and implementation of a pharmacist-managed clinical pharmacogenetics service}},
volume = {68},
year = {2011}
}
@article{Mitsios2007,
abstract = {BACKGROUND: Altered gene expression is an important feature of ischemic cerebral injury and affects proteins of many functional classes. We have used microarrays to investigate the changes in gene expression at various times after middle cerebral artery occlusion in human and rat brain.$\backslash$n$\backslash$nRESULTS: Our results demonstrated a significant difference in the number of genes affected and the time-course of expression between the two cases. The total number of deregulated genes in the rat was 335 versus 126 in the human, while, of 393 overlapping genes between the two array sets, 184 were changed only in the rat and 36 in the human with a total of 41 genes deregulated in both cases. Interestingly, the mean fold changes were much higher in the human. The expression of novel genes, including p21-activated kinase 1 (PAK1), matrix metalloproteinase 11 (MMP11) and integrase interactor 1, was further analyzed by RT-PCR, Western blotting and immunohistochemistry. Strong neuronal staining was seen for PAK1 and MMP11.$\backslash$n$\backslash$nCONCLUSION: Our findings confirmed previous studies reporting that gene expression screening can detect known and unknown transcriptional features of stroke and highlight the importance of research using human brain tissue in the search for novel therapeutic agents.},
author = {Mitsios, Nick and Saka, Mohamad and Krupinski, Jerzy and Pennucci, Roberta and Sanfeliu, Coral and Wang, Qiuyu and Rubio, Francisco and Gaffney, John and Kumar, Pat and Kumar, Shant and Sullivan, Matthew and Slevin, Mark},
doi = {10.1186/1471-2202-8-93},
file = {:Users/solomon/Documents/Mendeley Desktop/Mitsios et al. - 2007 - A microarray study of gene and protein regulation in human and rat brain following middle cerebral artery occlus.pdf:pdf},
isbn = {1471-2202},
issn = {1471-2202},
journal = {BMC Neuroscience},
keywords = {Aged,Aged, 80 and over,Animals,Anoxia,Anoxia: etiology,Anoxia: mortality,Brain,Brain: metabolism,Brain: pathology,Cells, Cultured,Chromosomal Proteins, Non-Histone,Chromosomal Proteins, Non-Histone: genetics,Chromosomal Proteins, Non-Histone: metabolism,DNA-Binding Proteins,DNA-Binding Proteins: genetics,DNA-Binding Proteins: metabolism,Female,Fetus,Gene Expression Regulation,Gene Expression Regulation: physiology,Glucose,Glucose: deficiency,Humans,Infarction, Middle Cerebral Artery,Infarction, Middle Cerebral Artery: genetics,Infarction, Middle Cerebral Artery: metabolism,Infarction, Middle Cerebral Artery: pathology,Male,Matrix Metalloproteinase 11,Matrix Metalloproteinase 11: genetics,Matrix Metalloproteinase 11: metabolism,Middle Aged,Oligonucleotide Array Sequence Analysis,Oligonucleotide Array Sequence Analysis: methods,Rats,Rats, Sprague-Dawley,Reverse Transcriptase Polymerase Chain Reaction,Reverse Transcriptase Polymerase Chain Reaction: m,Stroke,Stroke: genetics,Stroke: metabolism,Stroke: pathology,Transcription Factors,Transcription Factors: genetics,Transcription Factors: metabolism,p21-Activated Kinases,p21-Activated Kinases: genetics,p21-Activated Kinases: metabolism},
number = {1},
pages = {93},
pmid = {17997827},
title = {{A microarray study of gene and protein regulation in human and rat brain following middle cerebral artery occlusion}},
url = {http://www.pubmedcentral.nih.gov/articlerender.fcgi?artid=2194693{\&}tool=pmcentrez{\&}rendertype=abstract},
volume = {8},
year = {2007}
}
@article{Allabi2005,
abstract = {The genetically polymorphic cytochrome P450 2C9 (CYP2C9) metabolizes many important drugs. Among them, phenytoin has been used as a probe to determine CYP2C9 phenotype by measuring the urinary excretion of its major metabolite, S-enantiomer of 5-(4-hydroxyphenyl)-5-phenylhydantoin (p-HPPH). Phenytoin pharmacokinetic is also dependent on the activity of CYP2C19 and p-glycoprotein (ABCB1). To determine the influence of CYP2C9, CYP2C19 and ABCB1 genetic polymorphisms on phenytoin metabolism in a Black population, 109 healthy Beninese subjects received a single 300 mg oral dose of phenytoin. Blood was drawn 4 h after drug intake and urine was collected during the first 8 h. Plasma phenytoin and urine S- and R-enantiomers of p-HPPH were determined by high-performance liquid chromatography. Urinary excretion of (S)-p-HPPH [defined as urinary volumex(S)-p-HPPH urinary concentration] and PMR (defined as the ratio of p-HPPH in urine to 4 h phenytoin plasma concentration), both markers of CYP2C9 activity, were used to determine the functional relevance of new variants of CYP2C9 (*5, *6, *8, *9 and *11) in this population. Plasma phenytoin concentration was significantly associated with ABCB1 haplotype/genotype (P=0.05, Kruskal-Wallis test) and levels increased significantly in the genotype order: wild-type, T3421A and Block-2 genotypes (P=0.015, Jonckheere-Terpstra test). Urinary excretion of (S)-p-HPPH and PMR were significantly associated with the CYP2C9 genotype (P=0.001, analysis of variance (ANOVA) and P{\textless}0.0001, Kruskal-Wallis test, respectively) and decreased in the order: CYP2C9*1/*1, CYP2C9*1/*9, CYP2C9*9/*9, CYP2C9*1/*8, CYP2C9*8/*9, CYP2C9*9/*11, CYP2C9*1/*5, CYP2C9*6/*9, CYP2C9*1/*6, CYP2C9*8/*11, CYP2C9*5/*8 and CYP2C9*5/*6 (P{\textless}0.001, Jonckheere-Terpstra test). A combined analysis of CYP2C9, 2C19 and ABCB1 revealed that only ABCB1 predicted phenytoin concentration at 4 h and explained 8{\%} of the variability (r=0.08, P=0.04). On the other hand, only CYP2C9 was predictive for the urinary excretion of (S)-p-HPPH and PMR (r=0.21, P=0.001 and r=0.25, P{\textless}0.001, respectively). Furthermore, significant relation was found between urinary excretion of (R)-p-HPPH and CYP2C9 genotype (P=0.035) and levels significantly increased in the genotype order: CYP2C9*1/*9, CYP2C9*1/*1, CYP2C9*9/*11, CYP2C9*1/*8 and CYP2C9*1/*5 (P{\textless}0.001, Jonckheere-Terpstra test). In summary, the present study demonstrates that, in a Black population, CYP2C9*5, *6, *8 and *11 variants, but not CYP2C9*9, are associated with a decreased phenytoin metabolism. The data also confirm the limited contribution of MDR1 gene to inter-individual phenytoin pharmacokinetic variation.},
author = {Allabi, AC and Gala, JL and Horsmans, Y},
issn = {1744-6872},
journal = {Pharmacogenetics and genomics},
month = {nov},
number = {11},
pages = {779--86},
pmid = {16220110},
title = {{CYP2C9, CYP2C19, ABCB1 (MDR1) genetic polymorphisms and phenytoin metabolism in a Black Beninese population.}},
url = {http://www.ncbi.nlm.nih.gov/pubmed/16220110},
volume = {15},
year = {2005}
}
@article{Philpott2010,
author = {Philpott, Sean and Baker, Robert},
doi = {10.1111/j.1467-8519.2010.01855.x},
file = {:Users/solomon/Documents/Mendeley Desktop/Philpott, Baker - 2010 - Why the avandia scandal proves big pharma needs stronger ethical standards.pdf:pdf},
isbn = {1467-8519 (Electronic)$\backslash$r0269-9702 (Linking)},
issn = {02699702},
journal = {Bioethics},
number = {8},
pages = {7--8},
pmid = {21050246},
title = {{Why the avandia scandal proves big pharma needs stronger ethical standards}},
volume = {24},
year = {2010}
}
@article{Sharma2006,
abstract = {A simple technique for cerebrospinal fluid (CSF) collection was developed in F-344 rats. Cell counts and total protein concentrations were evaluated to assess sample quality. While the 50 to 70 mu L samples of CSF collected on three different days showed a progressive decrease in the total erythrocyte and nucleated cell counts, no significant changes were observed in the total protein concentrations. Progressive decreases in the total erythrocyte count correlated positively with the decreases in volume of CSF collected. Our data suggest that collection of less than 50 mu L of CSF will give a better quality of CSF in F-344 rats. This is the first report of cellular and protein parameters in the CSF of F-344 rats.},
author = {Sharma, Alok K and Schultze, Albert E and Cooper, Dale M and Reams, Rachel Y and Jordan, William H and Snyder, Paul W},
doi = {10.1080/01926230600798609},
file = {:Users/solomon/Documents/Mendeley Desktop/Sharma et al. - 2006 - Development of a percutaneous cerebrospinal fluid collection technique in F-344 rats and evaluation of cell count.pdf:pdf},
isbn = {0192623060},
issn = {0192-6233},
journal = {Toxicologic pathology},
keywords = {Animals,Cell Count,Cerebrospinal Fluid,Cerebrospinal Fluid: chemistry,Cerebrospinal Fluid: cytology,Erythrocyte Count,Evaluation Studies as Topic,Inbred F344,Male,Proteins,Proteins: analysis,Rats,Specimen Handling,Specimen Handling: methods},
number = {4},
pages = {393--5},
pmid = {16844667},
title = {{Development of a percutaneous cerebrospinal fluid collection technique in F-344 rats and evaluation of cell counts and total protein concentrations.}},
url = {http://tpx.sagepub.com/content/34/4/393.short},
volume = {34},
year = {2006}
}
@article{Sheth2016,
abstract = {Background Preclinical models of stroke have shown that intravenous glyburide reduces brain swelling and improves survival. We assessed whether intravenous glyburide (RP-1127; glibenclamide) would safely reduce brain swelling, decrease the need for decompressive craniectomy, and improve clinical outcomes in patients presenting with a large hemispheric infarction. Methods For this double-blind, randomised, placebo-controlled phase 2 trial, we enrolled patients (aged 18???80 years) with a clinical diagnosis of large anterior circulation hemispheric infarction for less than 10 h and baseline diffusion-weighted MRI image lesion volume of 82???300 cm3 on MRI at 18 hospitals in the USA. We used web-based randomisation (1:1) to allocate patients to the placebo or intravenous glyburide group. Intravenous glyburide was given as a 0??13 mg bolus intravenous injection for the first 2 min, followed by an infusion of 0??16 mg/h for the first 6 h and then 0??11 mg/h for the remaining 66 h. The primary efficacy outcome was the proportion of patients who achieved a modified Rankin Scale (mRS) score of 0???4 at 90 days without undergoing decompressive craniectomy. Analysis was by per protocol. Safety analysis included all randomly assigned patients who received the study drug. This trial is registered with ClinicalTrials.gov, number NCT01794182. Findings Between May 3, 2013, and April 30, 2015, 86 patients were randomly assigned but enrolment was stopped because of funding reasons. The funder, principal investigators, site investigators, patients, imaging core, and outcomes personnel were masked to treatment. The per-protocol study population was 41 participants who received intravenous glyburide and 36 participants who received placebo. 17 (41{\%}) patients in the intravenous glyburide group and 14 (39{\%}) in the placebo group had an mRS score of 0???4 at 90 days without decompressive craniectomy (adjusted odds ratio 0??87, 95{\%} CI 0??32???2??32; p=0??77). Ten (23{\%}) of 44 participants in the intravenous glyburide group and ten (26{\%}) of 39 participants in the placebo group had cardiac events (p=0??76), and four of 20 had serious adverse events (two in the intravenous glyburide group and two in the placebo group, p=1??00). One cardiac death occurred in each group (p=1??00). Interpretation Intravenous glyburide was well tolerated in patients with large hemispheric stroke at risk for cerebral oedema. There was no difference in the composite primary outcome. Further study is warranted to assess the potential clinical benefit of a reduction in swelling by intravenous glyburide. Funding Remedy Pharmaceuticals.},
author = {Sheth, KN and Elm, JJ and Molyneaux, BJ and Hinson, H and Beslow, LA and Sze, GK and Ostwaldt, AC and del Zoppo, GJ and Simard, JM and Jacobson, S and Kimberly, WT},
doi = {10.1016/S1474-4422(16)30196-X},
file = {:Users/solomon/Documents/Mendeley Desktop/Sheth et al. - 2016 - Safety and efficacy of intravenous glyburide on brain swelling after large hemispheric infarction (GAMES-RP) a ran.pdf:pdf},
isbn = {1474-4465 (Electronic) 1474-4422 (Linking)},
issn = {14744465},
journal = {The Lancet Neurology},
number = {11},
pages = {1160--1169},
pmid = {27567243},
publisher = {Elsevier Ltd},
title = {{Safety and efficacy of intravenous glyburide on brain swelling after large hemispheric infarction (GAMES-RP): a randomised, double-blind, placebo-controlled phase 2 trial}},
url = {http://dx.doi.org/10.1016/S1474-4422(16)30196-X},
volume = {15},
year = {2016}
}
@article{Guyatt2012,
abstract = {BACKGROUND: The objective of this article is to summarize the published literature concerning the pharmacokinetics and pharmacodynamics of oral anticoagulant drugs that are currently available for clinical use and other aspects related to their management. METHODS: We carried out a standard review of published articles focusing on the laboratory and clinical characteristics of the vitamin K antagonists; the direct thrombin inhibitor, dabigatran etexilate; and the direct factor Xa inhibitor, rivaroxaban RESULTS: The antithrombotic effect of each oral anticoagulant drug, the interactions, and the monitoring of anticoagulation intensity are described in detail and discussed without providing specific recommendations. Moreover, we describe and discuss the clinical applications and optimal dosages of oral anticoagulant therapies, practical issues related to their initiation and monitoring, adverse events such as bleeding and other potential side effects, and available strategies for reversal. CONCLUSIONS: There is a large amount of evidence on laboratory and clinical characteristics of vitamin K antagonists. A growing body of evidence is becoming available on the first new oral anticoagulant drugs available for clinical use, dabigatran and rivaroxaban.},
author = {Guyatt, Gordon H. and Akl, Elie A. and Crowther, Mark and Gutterman, David D. and Sch{\"{u}}nemann, Holger J.},
doi = {10.1378/chest.1412S3},
file = {:Users/solomon/Documents/Mendeley Desktop/Guyatt et al. - 2012 - Executive summary Antithrombotic therapy and prevention of thrombosis, 9th ed American College of Chest Physician.pdf:pdf},
isbn = {0012-3692},
issn = {00123692},
journal = {Chest},
number = {2 SUPPL.},
pages = {7--47},
pmid = {22315257},
title = {{Executive summary: Antithrombotic therapy and prevention of thrombosis, 9th ed: American College of Chest Physicians evidence-based clinical practice guidelines}},
volume = {141},
year = {2012}
}
@article{DeVlaminck2014,
abstract = {Monitoring allograft health is an important component of posttransplant therapy. Endomyocardial biopsy is the current gold standard for cardiac allograft monitoring but is an expensive and invasive procedure. Proof of principle of a universal, noninvasive diagnostic method based on high-throughput screening of circulating cell-free donor-derived DNA (cfdDNA) was recently demonstrated in a small retrospective cohort. We present the results of a prospective cohort study (65 patients, 565 samples) that tested the utility of cfdDNA in measuring acute rejection after heart transplantation. Circulating cell-free DNA was purified from plasma and sequenced (mean depth, 1.2 giga-base pairs) to quantify the fraction of cfdDNA. Through a comparison with endomyocardial biopsy results, we demonstrate that cfdDNA enables diagnosis of acute rejection after heart transplantation, with an area under the receiver operating characteristic curve of 0.83 and sensitivity and specificity that are comparable to the intrinsic performance of the biopsy itself. This noninvasive genome transplant dynamics approach is a powerful and informative method for routine monitoring of allograft health without incurring the risk, discomfort, and expense of an invasive biopsy.},
archivePrefix = {arXiv},
arxivId = {NIHMS150003},
author = {{De Vlaminck}, I and Valantine, H A and Snyder, T M and Strehl, C and Cohen, G and Luikart, H and Neff, N F and Okamoto, J and Bernstein, D and Weisshaar, D and Quake, S R and Khush, K K},
doi = {10.1126/scitranslmed.3007803},
eprint = {NIHMS150003},
file = {:Users/solomon/Documents/Mendeley Desktop/De Vlaminck et al. - 2014 - Circulating cell-free DNA enables noninvasive diagnosis of heart transplant rejection.pdf:pdf},
isbn = {1946-6234$\backslash$r1946-6242},
issn = {1946-6234},
journal = {Science Translational Medicine},
keywords = {circulating DNA},
number = {241},
pages = {241ra77--241ra77},
pmid = {1000000221},
title = {{Circulating cell-free DNA enables noninvasive diagnosis of heart transplant rejection.}},
url = {http://stm.sciencemag.org/cgi/doi/10.1126/scitranslmed.3007803{\%}5Cnpapers3://publication/doi/10.1126/scitranslmed.3007803},
volume = {6},
year = {2014}
}
@article{Dehghan2008,
abstract = {BACKGROUND: Hyperuricaemia, a highly heritable trait, is a key risk factor for gout. We aimed to identify novel genes associated with serum uric acid concentration and gout. METHODS: Genome-wide association studies were done for serum uric acid in 7699 participants in the Framingham cohort and in 4148 participants in the Rotterdam cohort. Genome-wide significant single nucleotide polymorphisms (SNPs) were replicated in white (n=11 024) and black (n=3843) individuals who took part in the study of Atherosclerosis Risk in Communities (ARIC). The SNPs that reached genome-wide significant association with uric acid in either the Framingham cohort (p{\textless}5.0 x 10(-8)) or the Rotterdam cohort (p{\textless}1.0 x 10(-7)) were evaluated with gout. The results obtained in white participants were combined using meta-analysis. FINDINGS: Three loci in the Framingham cohort and two in the Rotterdam cohort showed genome-wide association with uric acid. Top SNPs in each locus were: missense rs16890979 in SLC2A9 (p=7.0 x 10(-168) and 2.9 x 10(-18) for white and black participants, respectively); missense rs2231142 in ABCG2 (p=2.5 x 10(-60) and 9.8 x 10(-4)), and rs1165205 in SLC17A3 (p=3.3 x 10(-26) and 0.33). All SNPs were direction-consistent with gout in white participants: rs16890979 (OR 0.59 per T allele, 95{\%} CI 0.52-0.68, p=7.0 x 10(-14)), rs2231142 (1.74, 1.51-1.99, p=3.3 x 10(-15)), and rs1165205 (0.85, 0.77-0.94, p=0.002). In black participants of the ARIC study, rs2231142 was direction-consistent with gout (1.71, 1.06-2.77, p=0.028). An additive genetic risk score of high-risk alleles at the three loci showed graded associations with uric acid (272-351 mumol/L in the Framingham cohort, 269-386 mumol/L in the Rotterdam cohort, and 303-426 mumol/L in white participants of the ARIC study) and gout (frequency 2-13{\%} in the Framingham cohort, 2-8{\%} in the Rotterdam cohort, and 1-18{\%} in white participants in the ARIC study). INTERPRETATION: We identified three genetic loci associated with uric acid concentration and gout. A score based on genes with a putative role in renal urate handling showed a substantial risk for gout.},
author = {Dehghan, Abbas and K{\"{o}}ttgen, Anna and Yang, Qiong and Hwang, S.-J. and Kao, W H Linda and Rivadeneira, Fernando and Boerwinkle, E and Levy, Daniel and Hofman, Albert and Astor, Brad C and Benjamin, Emelia J and van Duijn, C M and Witteman, Jacqueline C and Coresh, Josef and Fox, Caroline S and Duijn, Cornelia M Van and Witteman, Jacqueline C and Coresh, Josef and Fox, Caroline S},
doi = {10.1016/S0140-6736(08)61343-4.Association},
file = {:Users/solomon/Documents/Mendeley Desktop/Dehghan et al. - 2008 - Association of three genetic loci with uric acid concentration and risk of gout a genome-wide association study.pdf:pdf},
isbn = {0140-6736},
issn = {01406736},
journal = {Lancet},
keywords = {4-5,6-12 and X,Alleles,Atherosclerosis,Chromosomes,Cohort Studies,Female,Genes,Genetics,Genome,Genotype,Gout,Human,Humans,Hyperuricemia,Male,Middle Aged,Netherlands,Polymorphism,Population,Prevalence,Research,Risk,Risk Factors,SNP,Single Nucleotide,Uric Acid,blood,complications,epidemiology,etiology,genetics,metabolism,methods,single nucleotide polymorphism},
number = {9654},
pages = {1953--1961},
pmid = {18834626},
publisher = {Elsevier Ltd},
title = {{Association of three genetic loci with uric acid concentration and risk of gout: a genome-wide association study}},
url = {http://dx.doi.org/10.1016/S0140-6736(08)61343-4 http://www.ncbi.nlm.nih.gov/pubmed/18834626},
volume = {372},
year = {2008}
}
@article{Wagner2013,
abstract = {The molecular mechanisms underlying TBI pathophysiology and recovery are both complex and varied. Further, the pathology underlying many of the clinical sequelae observed in this population evolve over the acute injury period and encompass the subacute and chronic phases of recovery, supporting the contemporary concept that TBI is a chronic disease rather than a static insult from which limited recovery occurs. TBI related complications can also span from acute care to the very chronic stages of recovery that occur years after the initial trauma. Despite ongoing neurodegeneration, the TBI recovery period is also characterized by a propensity for neuroplasticity and rewiring through multiple mechanisms. This review summarizes key elements of acute pathophysiology, how they link to structural damage and ongoing degeneration, and how this process coincides with a permissive neuroplastic environment. The pathophysiology of selected TBI related complications is also discussed. Each of these concepts is studied through the lens of Rehabilomics, wherein an emphasis is placed on biomarker studies characterizing these pathophysiological mechanisms, and biomarker profiles are assessed in relation to multi-modal outcomes and susceptibility to rehabilitation relevant complications. In reviewing these concepts, implications for future research and theranostic principles for patient care are presented. {\textcopyright} 2012 Elsevier Ireland Ltd.},
author = {Wagner, AK and Zitelli, KT},
doi = {10.1016/j.pathophys.2012.02.007},
isbn = {0928-4680},
issn = {09284680},
journal = {Pathophysiology},
keywords = {Depression,Hypogonadism,Pathophysiology,Recovery,Rehabilomics,Seizures,TBI},
number = {1},
pages = {39--48},
pmid = {22444246},
publisher = {Elsevier Ireland Ltd},
title = {{A Rehabilomics focused perspective on molecular mechanisms underlying neurological injury, complications, and recovery after severe TBI}},
url = {http://dx.doi.org/10.1016/j.pathophys.2012.02.007},
volume = {20},
year = {2013}
}
@article{Chuang2010,
abstract = {Neuroglobin has shown rich neuroprotective effects against cerebral hypoxia, and therefore has the potential to impact outcomes after traumatic brain injury (TBI). However, to date an association between genetic variation within the human neuroglobin (NGB) gene and recovery post-TBI has not been reported. The purpose of this study was to explore the relationship between NGB genotypes and outcomes (as assessed by the Glasgow Outcome Scale [GOS], the Disability Rating Scale [DRS], and the Neurobehavioral Rating Scale-Revised [NRS-R]) after severe TBI. Genotyping using TaqMan allele discrimination for two tagging single nucleotide polymorphisms (tSNPs) that represent the two haplotype blocks for NGB (rs3783988 and rs10133981) was completed on DNA obtained from 196 Caucasian patients recovering from severe TBI. Patients were dichotomized based on the presence of the variant allele for each tSNP. Chi-square and Fisher's exact tests were used to compare characteristics between groups. Multivariate linear regression was used to examine NGB tSNPs and recovery from severe TBI. Subjects with the TT genotype (wild-type) for rs3783988 were more likely to have better GOS and DRS scores at 3, 6, 12, and 24 months, while rs10133981 genotype was not significantly related to functional outcome. After controlling for age, gender, and Glasgow Coma Scale (GCS) score, those subjects with the rs3783988 TT genotype had more than a 2.65-times greater likelihood of better functional outcomes than individuals with genotypes harboring a variant allele. Data suggest that the haplotype block represented by rs3783988 in NGB appears to influence recovery after severe TBI. Represented within this haplotype block of NGB is the region that codes for the oxygen-binding portion of NGB.},
author = {Chuang, Pei-Ying and Conley, Yvette P and Poloyac, Samuel M and Okonkwo, David O and Ren, Dianxu and Sherwood, Paula R and Hravnak, Marilyn and Alexander, Sheila a},
doi = {10.1089/neu.2009.1129},
file = {:Users/solomon/Documents/Mendeley Desktop/Chuang et al. - 2010 - Neuroglobin genetic polymorphisms and their relationship to functional outcomes after traumatic brain injury.pdf:pdf},
isbn = {1557-9042 (Electronic)$\backslash$r0897-7151 (Linking)},
issn = {1557-9042},
journal = {Journal of neurotrauma},
keywords = {Adolescent,Adult,Aged,Alleles,Brain Injuries,Brain Injuries: genetics,Brain Injuries: rehabilitation,Chi-Square Distribution,Disability Evaluation,Female,Genetic Association Studies,Genotype,Glasgow Outcome Scale,Globins,Globins: genetics,Haplotypes,Humans,Linear Models,Male,Middle Aged,Nerve Tissue Proteins,Nerve Tissue Proteins: genetics,Polymorphism,Recovery of Function,Recovery of Function: genetics,Single Nucleotide,Treatment Outcome},
number = {6},
pages = {999--1006},
pmid = {20345238},
title = {{Neuroglobin genetic polymorphisms and their relationship to functional outcomes after traumatic brain injury.}},
url = {http://www.pubmedcentral.nih.gov/articlerender.fcgi?artid=2943497{\&}tool=pmcentrez{\&}rendertype=abstract},
volume = {27},
year = {2010}
}
@article{Hung2011a,
author = {Hung, Chin-Chuan and Chiou, Mu-Han and Huang, Bo-Hau and Hsieh, Yow-Wen and Hsieh, Tsung-Jen and Huang, Chieh-Liang and Lane, Hsien-Yuan},
doi = {10.2217/pgs.11.96},
file = {:Users/solomon/Documents/Mendeley Desktop/Hung et al. - 2011 - Impact of genetic polymorphisms in ABCB1 , CYP2B6 , OPRM1 , ANKK1 and DRD2 genes on methadone therapy in Han Chines.pdf:pdf},
issn = {1462-2416},
journal = {Pharmacogenomics},
month = {nov},
number = {11},
pages = {1525--1533},
title = {{Impact of genetic polymorphisms in ABCB1 , CYP2B6 , OPRM1 , ANKK1 and DRD2 genes on methadone therapy in Han Chinese patients}},
url = {http://www.futuremedicine.com/doi/10.2217/pgs.11.96},
volume = {12},
year = {2011}
}
@article{Ben-Zvi2014,
abstract = {The central nervous system (CNS) requires a tightly controlled environment free of toxins and pathogens to provide the proper chemical composition for neural function. This environment is maintained by the 'blood-brain barrier' (BBB), which is composed of blood vessels whose endothelial cells display specialized tight junctions and extremely low rates of transcellular vesicular transport (transcytosis). In concert with pericytes and astrocytes, this unique brain endothelial physiological barrier seals the CNS and controls substance influx and efflux. Although BBB breakdown has recently been associated with initiation and perpetuation of various neurological disorders, an intact BBB is a major obstacle for drug delivery to the CNS. A limited understanding of the molecular mechanisms that control BBB formation has hindered our ability to manipulate the BBB in disease and therapy. Here we identify mechanisms governing the establishment of a functional BBB. First, using a novel tracer-injection method for embryos, we demonstrate spatiotemporal developmental profiles of BBB functionality and find that the mouse BBB becomes functional at embryonic day 15.5 (E15.5). We then screen for BBB-specific genes expressed during BBB formation, and find that major facilitator super family domain containing 2a (Mfsd2a) is selectively expressed in BBB-containing blood vessels in the CNS. Genetic ablation of Mfsd2a results in a leaky BBB from embryonic stages through to adulthood, but the normal patterning of vascular networks is maintained. Electron microscopy examination reveals a dramatic increase in CNS-endothelial-cell vesicular transcytosis in Mfsd2a(-/-) mice, without obvious tight-junction defects. Finally we show that Mfsd2a endothelial expression is regulated by pericytes to facilitate BBB integrity. These findings identify Mfsd2a as a key regulator of BBB function that may act by suppressing transcytosis in CNS endothelial cells. Furthermore, our findings may aid in efforts to develop therapeutic approaches for CNS drug delivery.},
archivePrefix = {arXiv},
arxivId = {NIHMS150003},
author = {Ben-Zvi, Ayal and Lacoste, Baptiste and Kur, Esther and Andreone, Benjamin J and Mayshar, Yoav and Yan, Han and Gu, Chenghua},
doi = {10.1038/nature13324},
eprint = {NIHMS150003},
file = {:Users/solomon/Documents/Mendeley Desktop/Ben-Zvi et al. - 2014 - Mfsd2a is critical for the formation and function of the blood-brain barrier.pdf:pdf},
isbn = {doi:10.1038/nature13324},
issn = {1476-4687},
journal = {Nature},
keywords = {Animals,Blood Vessels,Blood Vessels: metabolism,Blood-Brain Barrier,Blood-Brain Barrier: embryology,Blood-Brain Barrier: physiology,Cerebral Cortex,Cerebral Cortex: blood supply,Cerebral Cortex: embryology,Cerebral Cortex: metabolism,Drug Delivery Systems,Endothelial Cells,Endothelial Cells: metabolism,Female,Gene Expression Profiling,Male,Membrane Transport Proteins,Membrane Transport Proteins: deficiency,Membrane Transport Proteins: genetics,Membrane Transport Proteins: metabolism,Mice,Neovascularization, Physiologic,Pericytes,Pericytes: metabolism,Spatio-Temporal Analysis,Tight Junctions,Tight Junctions: metabolism,Tight Junctions: pathology,Transcytosis},
number = {7501},
pages = {507--11},
pmid = {24828040},
publisher = {Nature Publishing Group},
title = {{Mfsd2a is critical for the formation and function of the blood-brain barrier.}},
url = {http://dx.doi.org/10.1038/nature13324},
volume = {509},
year = {2014}
}
@article{Whirl-Carrillo2012,
abstract = {The Pharmacogenomics Knowledgebase (PharmGKB) is a resource that collects, curates, and disseminates information about the impact of human genetic variation on drug responses. It provides clinically relevant information, including dosing guidelines, annotated drug labels, and potentially actionable gene-drug associations and genotype-phenotype relationships. Curators assign levels of evidence to variant-drug associations using well-defined criteria based on careful literature review. Thus, PharmGKB is a useful source of high-quality information supporting personalized medicine-implementation projects.},
author = {Whirl-Carrillo, M and McDonagh, E M and Hebert, J M and Gong, L and Sangkuhl, K and Thorn, C F and Altman, R B and Klein, T E},
doi = {10.1038/clpt.2012.96},
file = {:Users/solomon/Documents/Mendeley Desktop/Whirl-Carrillo et al. - 2012 - Pharmacogenomics Knowledge for Personalized Medicine.pdf:pdf},
issn = {0009-9236},
journal = {Clinical Pharmacology {\&} Therapeutics},
month = {oct},
number = {4},
pages = {414--417},
pmid = {22992668},
publisher = {NIH Public Access},
title = {{Pharmacogenomics Knowledge for Personalized Medicine}},
url = {http://www.ncbi.nlm.nih.gov/pubmed/22992668 http://www.pubmedcentral.nih.gov/articlerender.fcgi?artid=PMC3660037 http://doi.wiley.com/10.1038/clpt.2012.96},
volume = {92},
year = {2012}
}
@article{Relling2013,
abstract = {Thiopurine methyltransferase (TPMT) activity exhibits monogenic co-dominant inheritance, with ethnic differences in the frequency of occurrence of variant alleles. With conventional thiopurine doses, homozygous TPMT-deficient patients ({\~{}}1 in 178 to 1 in 3,736 individuals with two nonfunctional TPMT alleles) experience severe myelosuppression, 30-60{\%} of individuals who are heterozygotes ({\~{}}3-14{\%} of the population) show moderate toxicity, and homozygous wild-type individuals ({\~{}}86-97{\%} of the population) show lower active thioguanine nucleolides and less myelosuppression. We provide dosing recommendations (updates at http://www.pharmgkb.org) for azathioprine, mercaptopurine (MP), and thioguanine based on TPMT genotype.},
author = {Relling, M V and Gardner, E E and Sandborn, W J and Schmiegelow, K and Pui, C-H and Yee, S W and Stein, C M and Carrillo, M and Evans, W E and Klein, T E and Hicks, J K and Schwab, M and Klein, T E},
doi = {10.1038/clpt.2013.4},
file = {:Users/solomon/Documents/Mendeley Desktop/Relling et al. - 2013 - Clinical Pharmacogenetics Implementation Consortium Guidelines for Thiopurine Methyltransferase Genotype and Thi.pdf:pdf},
issn = {0009-9236},
journal = {Clin. Pharmacol. Ther.},
month = {mar},
number = {3},
pages = {324--325},
pmid = {23422873},
publisher = {Wiley-Blackwell},
title = {{Clinical Pharmacogenetics Implementation Consortium Guidelines for Thiopurine Methyltransferase Genotype and Thiopurine Dosing: 2013 Update}},
url = {http://www.ncbi.nlm.nih.gov/pubmed/23422873 http://www.pubmedcentral.nih.gov/articlerender.fcgi?artid=PMC3604643 http://doi.wiley.com/10.1038/clpt.2013.4 http://www.ncbi.nlm.nih.gov/pubmed/21270794 http://www.pubmedcentral.nih.gov/articlerender.fcgi?artid},
volume = {93},
year = {2013}
}
@article{Tan2010,
abstract = {Breast cancer resistance protein (BCRP/ABCG2) is a membrane-bound efflux transporter important in cellular detoxification and multidrug resistance. Some aryl hydrocarbon receptor (AHR) agonists were reported to induce BCRP expression in human colon carcinoma cells. However, a direct involvement of AHR transcriptional regulation remains unexplored. In this study, we show that BCRP induction by AHR ligands occurs in human intestinal, liver, and mammary carcinoma cells and in primary colonocytes and hepatocytes. Increased BCRP transporter activity consistent with gene induction was also evident in the Caco2 subclone C2bbe1 cells. Using RNA interference and ectopic expression techniques to manipulate cellular AHR status, we confirmed AHR dependence of ABCG2 gene regulation. By gene promoter analysis, chromatin immunoprecipitation, and electrophoretic mobility shift assays, an active, proximal dioxin-response element at -194/-190 base pairs upstream of the transcription start site of the human ABCG2 gene was identified. Despite a common observation in human-derived cells, our in vitro and in vivo studies supported by phylogenetic footprinting analysis did not find that mouse Abcg2 is subject to AHR regulation. We conclude that AHR is a direct transcriptional regulator of human BCRP and provide an unprecedented role of AHR in cellular adaptive response and cytoprotection by up-regulating an important ATP-binding cassette efflux transporter.},
author = {Tan, Kah Poh and Wang, Bernice and Yang, Mingdong and Boutros, Paul C and Macaulay, Jane and Xu, Haibo and Chuang, Andrew I and Kosuge, Kazuhiro and Yamamoto, Mika and Takahashi, Shinichiro and Wu, Alex M L and Ross, Douglas D and Harper, Patricia a and Ito, Shinya},
doi = {10.1124/mol.110.065078.},
file = {:Users/solomon/Documents/Mendeley Desktop/Tan et al. - 2010 - Aryl Hydrocarbon Receptor Is a Transcriptional Activator of the Human Breast Cancer Resistance Protein ( BCRP ABCG2.pdf:pdf},
issn = {1521-0111},
journal = {Molecular Pharmacology},
keywords = {ATP-Binding Cassette Transporters,ATP-Binding Cassette Transporters: genetics,Animals,Aryl Hydrocarbon,Aryl Hydrocarbon: physiology,Base Sequence,Cell Line,DNA Primers,Female,Gene Expression Regulation,Humans,Immunohistochemistry,Mice,Neoplasm Proteins,Neoplasm Proteins: genetics,Phylogeny,Pregnancy,Receptors,Reverse Transcriptase Polymerase Chain Reaction,Trans-Activators,Trans-Activators: physiology,Tumor},
number = {2},
pages = {175--185},
pmid = {20460431},
title = {{Aryl Hydrocarbon Receptor Is a Transcriptional Activator of the Human Breast Cancer Resistance Protein ( BCRP / ABCG2 )}},
url = {http://www.ncbi.nlm.nih.gov/pubmed/20460431},
volume = {78},
year = {2010}
}
@article{Salottolo2014,
abstract = {IMPORTANCE: The Glasgow Coma Scale (GCS) is used frequently to define the extent of neurologic injury in patients with a traumatic brain injury (TBI). Whether age affects the predictive ability of the GCS for severity of TBI (determined by the Abbreviated Injury Scale [AIS] score) remains unknown.$\backslash$n$\backslash$nOBJECTIVE: To investigate the effect of age on the association between the GCS and anatomic TBI severity.$\backslash$n$\backslash$nDESIGN, SETTING, AND PARTICIPANTS: We examined all patients with a TBI, defined by diagnostic codes 850 to 854 from the International Classification of Diseases, Ninth Revision, Clinical Modification, who were admitted to 2 level I trauma centers from January 1, 2008, through December 31, 2012.$\backslash$n$\backslash$nEXPOSURES: We compared elderly (≥65 years) and younger (18-64 years) adults with TBI.$\backslash$n$\backslash$nMAIN OUTCOMES AND MEASURES: We examined differences by age in GCS category (defined by emergency department GCS as severe [3-8], moderate [9-12], or mild [13-15]) at each level of TBI severity (head AIS score, 1 [minor] to 5 [critical]). Cochran-Armitage $\chi$² trend tests and stepwise multivariate linear and logistic regression models were used.$\backslash$n$\backslash$nRESULTS: During the study period, 6710 patients had a TBI (aged {\textless}65 years, 73.17{\%}). Significant differences in GCS category by age occurred at each AIS score (P ≤ .01 for all). In particular, among patients with an AIS score of 5, most of the elderly patients (56.33{\%}) had a mild neurologic deficit (GCS score, 13-15), whereas most of the younger patients (63.28{\%}) had a severe neurologic deficit (GCS score, 3-8). After adjustment, the younger adults had increased odds of presenting with a severe neurologic deficit (GCS score, 3-8) at each of the following AIS scores: 1, 4.2 (95{\%} CI, 1.0-17.6; P = .05); 2, 2.0 (1.0-3.7; P = .04); 3, 2.0 (1.2-3.5; P = .01); 4, 4.6 (2.8-7.5; P {\textless} .001); and 5, 3.1 (2.1-4.6; P {\textless} .001). The interaction between age and GCS for anatomic TBI severity remained significant after adjustment (estimate, -0.11; P = .005).$\backslash$n$\backslash$nCONCLUSIONS AND RELEVANCE: Age affects the relationship between the GCS score and anatomic TBI severity. Elderly TBI patients have better GCS scores than younger TBI patients with similar TBI severity. These findings have implications for TBI outcomes research and for protocols and research selection criteria that use the GCS.},
author = {Salottolo, Kristin and {Stewart Levy}, A. and Slone, Denetta S. and Mains, Charles W. and Bar-Or, David},
doi = {10.1001/jamasurg.2014.13},
file = {:Users/solomon/Documents/Mendeley Desktop/Salottolo et al. - 2014 - The effect of age on glasgow coma scale score in patients with traumatic brain injury.pdf:pdf},
isbn = {2168-6262},
issn = {21686254},
journal = {JAMA Surgery},
number = {7},
pages = {727--734},
pmid = {24899145},
title = {{The effect of age on glasgow coma scale score in patients with traumatic brain injury}},
volume = {149},
year = {2014}
}
@article{Biswas2017,
abstract = {Background: Traumatic brain injury (TBI) is a much researched topic in medical health, which requires additional studies to understand various effects of demographic and geographic factors that can assist in developing the most effective treatments. Thousands of people of different ages are suffering from lifelong disabilities, either mild or severe, from TBI and the number is increasing. This study aims to increase our understanding of the effect of sex and age by applying five different statistical methods to evaluate the effect of these covariates on two independent TBI data sets representing patients from different geographical cohorts. A primary data was collected from Bangladesh and it was compared with CRASH (Corticosteroid Randomisation after Significant Head Injury) data, representing various countries around the world. Methods: The outcome variable for TBI considered in this paper is Glasgow Outcome Scale, which is a four point scale. It was converted to a binary outcome scale for fitting of Fisher's exact test, a test of proportions and a binary linear model. For analyzing ordinal outcomes, the proportional odds model and the sliding dichotomy model were fitted. As the sample size of the Bangladeshi data set was small, parametric bootstrapping was applied for the consistency of results. Results: Females were the worse sufferers of TBI compared to men, according to CRASH data set. The old (aged above 58 years) followed by adults (age 25 to 58) were the most vulnerable victims. Interaction effects concluded that old women tended to endure the worst outcomes of TBI. This conclusion came from the CRASH data set representing the world in general, whereas such effects were not present in the Bangladesh data set. Additional application of parametric bootstrapping for the smaller Bangladesh data set did not result into any significant outcome. Conclusion: The effect of gender and age could be stronger in some countries than others which is driving the significance in CRASH and was not found in Bangladesh. It reflects the necessity of incorporating geographic patterns as well as demographic features of patients while developing treatments and designing clinical trials.},
author = {Biswas, Raaj Kishore and Kabir, Enamul and King, Rachel},
doi = {10.1186/s13690-017-0211-y},
file = {:Users/solomon/Documents/Mendeley Desktop/Biswas, Kabir, King - 2017 - Effect of sex and age on traumatic brain injury A geographical comparative study.pdf:pdf},
issn = {20493258},
journal = {Archives of Public Health},
keywords = {Bangladesh,Glasgow outcome scale,Health geography,Ordinal outcome scale,Public health},
number = {1},
pages = {1--11},
publisher = {Archives of Public Health},
title = {{Effect of sex and age on traumatic brain injury: A geographical comparative study}},
volume = {75},
year = {2017}
}
@article{Borgna-pignatti2009,
author = {Borgna-pignatti, Caterina and Cappellini, Maria Domenica and Stefano, Piero De and Carlo, Giovanni and Vecchio, Del and Forni, Gian Luca and Gamberini, Maria Rita and Ghilardi, Roberta and Piga, Antonio and Romeo, Maria Antonietta and Dc, Washington and Cells, R E D and Zhao, Huaqing and Cnaan, Avital},
doi = {10.1182/blood-2005-07-2933},
file = {:Users/solomon/Documents/Mendeley Desktop/Borgna-pignatti et al. - 2009 - Cardiac morbidity and mortality in deferoxamine- or deferiprone-treated patients with thalassemia major.pdf:pdf},
journal = {Blood},
number = {9},
pages = {3733--3737},
title = {{Cardiac morbidity and mortality in deferoxamine- or deferiprone-treated patients with thalassemia major Cardiac morbidity and mortality in deferoxamine- or deferiprone-treated patients with thalassemia major}},
volume = {107},
year = {2009}
}
@article{Jennett1981,
abstract = {The nature of the neurological and mental disabilities resulting from severe head injuries are analysed in 150 patients. Mental handicap contributed more significantly to overall social disability than did neurological deficits. This social handicap is readily described by the Glasgow Outcome Scale, an extended version of which is described and compared with alternatives. Comments are made about the quality of life in disabled survivors.},
author = {Jennett, B and Snoek, J and Bond, M R and Brooks, N},
doi = {10.1136/jnnp.44.4.285},
file = {:Users/solomon/Documents/Mendeley Desktop/Jennett et al. - 1981 - Disability after severe head injury observations on the use of the Glasgow Outcome Scale.pdf:pdf},
isbn = {0022-3050 (Print)},
issn = {0022-3050},
journal = {Journal of neurology, neurosurgery, and psychiatry},
number = {4},
pages = {285--93},
pmid = {6453957},
title = {{Disability after severe head injury: observations on the use of the Glasgow Outcome Scale.}},
url = {http://www.ncbi.nlm.nih.gov/pubmed/6453957{\%}5Cnhttp://www.pubmedcentral.nih.gov/articlerender.fcgi?artid=PMC490949},
volume = {44},
year = {1981}
}
@article{Rasmussen-Torvik2014,
abstract = {We describe here the design and initial implementation of the eMERGE-PGx project. eMERGE-PGx, a partnership of the Electronic Medical Records and Genomics Network and the Pharmacogenomics Research Network, has three objectives: (i) to deploy PGRNseq, a next-generation sequencing platform assessing sequence variation in 84 proposed pharmacogenes, in nearly 9,000 patients likely to be prescribed drugs of interest in a 1- to 3-year time frame across several clinical sites; (ii) to integrate well-established clinically validated pharmacogenetic genotypes into the electronic health record with associated clinical decision support and to assess process and clinical outcomes of implementation; and (iii) to develop a repository of pharmacogenetic variants of unknown significance linked to a repository of electronic health record-based clinical phenotype data for ongoing pharmacogenomics discovery. We describe site-specific project implementation and anticipated products, including genetic variant and phenotype data repositories, novel variant association studies, clinical decision support modules, clinical and process outcomes, approaches to managing incidental findings, and patient and clinician education methods.},
author = {Rasmussen-Torvik, L J and Stallings, S C and Gordon, A S and Almoguera, B and Basford, M A and Bielinski, S J and Brautbar, A and Brilliant, M H and Carrell, D S and Connolly, J J and Crosslin, D R and Doheny, K F and Gallego, C J and Gottesman, O and Kim, D S and Leppig, K A and Li, R and Lin, S and Manzi, S and Mejia, A R and Pacheco, J A and Pan, V and Pathak, J and Perry, C L and Peterson, J F and Prows, C A and Ralston, J and Rasmussen, L V and Ritchie, M D and Sadhasivam, S and Scott, S A and Smith, M and Vega, A and Vinks, A A and Volpi, S and Wolf, W A and Bottinger, E and Chisholm, R L and Chute, C G and Haines, J L and Harley, J B and Keating, B and Holm, I A and Kullo, I J and Jarvik, G P and Larson, E B and Manolio, T and McCarty, C A and Nickerson, D A and Scherer, S E and Williams, M S and Roden, D M and Denny, J C},
doi = {10.1038/clpt.2014.137},
isbn = {1532-6535},
issn = {1532-6535},
journal = {Clinical pharmacology and therapeutics},
keywords = {Adolescent,Aged,Child,DNA,Databases,Drug Therapy,Electronic Health Records,Electronic Health Records: organization {\&} administ,Female,Genetic,Genetic Association Studies,Genetic Variation,Genotype,Humans,Knowledge Bases,Male,Middle Aged,Pharmacogenetics,Phenotype,Pilot Projects,Sequence Analysis,Young Adult},
number = {4},
pages = {482--9},
pmid = {24960519},
title = {{Design and anticipated outcomes of the eMERGE-PGx project: a multicenter pilot for preemptive pharmacogenomics in electronic health record systems.}},
url = {http://www.pubmedcentral.nih.gov/articlerender.fcgi?artid=4169732{\&}tool=pmcentrez{\&}rendertype=abstract},
volume = {96},
year = {2014}
}
@article{Brownstein2014,
abstract = {23andme has suspended marketing of health-related reports due to US Food and Drug Administration approval violations. This has fostered discussions on the actual risks associated with consumer use of these reports. In the case described below, rare genotypes for the gene encoding thiopurine methyltransferase (TPMT) were misinterpreted by a direct-to-consumer (DTC) company, and risk calculations for breast cancer were offered when accuracy was not possible from the available information. Politics aside, these examples illustrate risks associated with DTC genetic testing without professional interpretation.},
author = {Brownstein, C a and Margulies, D M and Manzi, S F},
doi = {10.1038/clpt.2014.60},
issn = {1532-6535},
journal = {Clinical pharmacology and therapeutics},
keywords = {Alleles,Breast Neoplasms,Breast Neoplasms: enzymology,Breast Neoplasms: genetics,Genetic Testing,Methyltransferases,Methyltransferases: genetics,Pharmacogenetics,Polymorphism,Risk Assessment,Single Nucleotide,Single Nucleotide: genetics,United States,United States Food and Drug Administration},
number = {6},
pages = {598--600},
pmid = {24714787},
publisher = {Nature Publishing Group},
title = {{Misinterpretation of TPMT by a DTC genetic testing company.}},
url = {http://www.ncbi.nlm.nih.gov/pubmed/24714787},
volume = {95},
year = {2014}
}
@article{Li2013,
abstract = {Hypoxia-inducible factor-1$\alpha$ (HIF-1$\alpha$), a well-identified hypoxia-related protein, is involved in regulating the biological functions of various cell types including neurons. The traditional biological function of HIF-1$\alpha$ is promoting the transcription of some pro-survival genes when exposing to low oxygen conditions. Meanwhile, some studies also point out that HIF-1$\alpha$ shows the detrimental role in several central nervous system (CNS) disorders. Up to now, the knowledge of HIF-1$\alpha$ function in CNS is still limited. To investigate whether HIF-1$\alpha$ is involved in CNS impairment and repair, we employed a traumatic brain injury model in adult rats. Upregulation of HIF-1$\alpha$ was observed in the injured brain cortex by western blot analysis and immunohistochemistry staining. Terminal deoxynucleotidyl transferase deoxy-UTP nick-end labeling (TUNEL) and 4',6-diamidino-2-phenylindole (DAPI) staining suggested that HIF-1$\alpha$ was relevant to neuronal apoptosis after brain injury. In addition, glutamate excitotoxic model of primary cortex neurons was introduced to further investigate the role of HIF-1$\alpha$ in neuronal apoptosis; the result implied HIF-1$\alpha$ was associated with the regulation of p53 and BNIP3 in the apoptotic neurons. Based on our data, we suggested that HIF-1$\alpha$ might play an important role in neuronal apoptosis after traumatic brain injury in rat, which might also provide a basis for the further study on its role in regulating the transcription of target genes in apoptotic neurons, and might gain a novel strategy for the clinical therapy for traumatic brain injury.},
author = {Li, Aihong and Sun, Xiaolei and Ni, Yaohui and Chen, Xin and Guo, Aisong},
doi = {10.1007/s12031-013-0084-7},
file = {:Users/solomon/Documents/Mendeley Desktop/Li et al. - 2013 - HIF-1$\alpha$ involves in neuronal apoptosis after traumatic brain injury in adult rats.pdf:pdf},
issn = {1559-1166},
journal = {Journal of molecular neuroscience : MN},
keywords = {hif-1 $\alpha$,neuronal,traumatic brain injury},
number = {3},
pages = {1052--62},
pmid = {23979836},
title = {{HIF-1$\alpha$ involves in neuronal apoptosis after traumatic brain injury in adult rats.}},
url = {http://www.ncbi.nlm.nih.gov/pubmed/23979836},
volume = {51},
year = {2013}
}
@article{Klein2017,
author = {Klein, Michelle E. and Parvez, Md Masud and Shin, Jae-Gook},
doi = {10.1016/j.xphs.2017.04.051},
issn = {00223549},
journal = {Journal of Pharmaceutical Sciences},
month = {sep},
number = {9},
pages = {2368--2379},
publisher = {Elsevier},
title = {{Clinical Implementation of Pharmacogenomics for Personalized Precision Medicine: Barriers and Solutions}},
url = {http://www.sciencedirect.com/science/article/pii/S0022354917303209 http://linkinghub.elsevier.com/retrieve/pii/S0022354917303209},
volume = {106},
year = {2017}
}
@article{Katzenberger2015,
abstract = {Traumatic brain injury (TBI) is a complex disorder that affects millions of people worldwide. The complexity of TBI partly stems from the fact that injuries to the brain instigate non-neurological injuries to other organs such as the intestine. Additionally, genetic variation is thought to play a large role in determining the nature and severity of non-neurological injuries. We recently reported that TBI in flies, as in humans, increases permeability of the intestinal epithelial barrier resulting in hyperglycemia and a higher risk of death. Furthermore, we demonstrated that genetic variation in flies is also pertinent to the complexity of non-neurological injuries following TBI. The goals of this review are to place our findings in the context of what is known about TBI-induced intestinal permeability from studies of TBI patients and rodent TBI models and to draw attention to how studies of the fly TBI model can provide unique insights that may facilitate diagnosis and treatment of TBI.},
author = {Katzenberger, Rebeccah J. and Ganetzky, Barry and Wassarman, David A.},
doi = {10.1080/19336934.2015.1085623},
file = {:Users/solomon/Documents/Mendeley Desktop/Katzenberger, Ganetzky, Wassarman - 2015 - The gut reaction to traumatic brain injury.pdf:pdf},
isbn = {1933-6942 (Electronic)$\backslash$r1933-6934 (Linking)},
issn = {19336942},
journal = {Fly},
keywords = {Bacteria,Drosophila,Glucose,Innate immune response,Intestine,Mortality,Septate junction,TBI,Tight junction,Traumatic brain injury},
number = {2},
pages = {68--74},
pmid = {26291482},
title = {{The gut reaction to traumatic brain injury}},
volume = {9},
year = {2015}
}
@article{Cooper2006,
abstract = {A simple HPLC method was developed and validated for the determination of uric acid (UA), xanthine (X) and hypoxanthine (HX) concentrations in human serum to support pharmacodynamic (PD) studies of a novel xanthine oxidase inhibitor during its clinical development. Serum proteins were removed by ultrafiltration. The hydrophilic analytes and the I.S. were eluted by 100{\%} aqueous phosphate buffer mobile phase. The hydrophobic matrix components (late peaks) were eluted with a step gradient of a higher organic mobile phase. Validation on linearity, sensitivity, precision, accuracy, stability, and robustness of the method for PD biomarkers (UA, X, and HX) was carried out in a similar manner to that for pharmacokinetic (PK) data where applicable. Issues of selectivity for endogenous biomarker analytes and individual concentration variations were addressed during method validation. Standards were prepared in analyte-free phosphate buffer. Quality control samples were prepared in control serum from individuals not dosed with the xanthine oxidase inhibitor. The method was simple and robust with good accuracy and precision for the measurement of serum UA, X, and HX concentrations. ?? 2006 Elsevier B.V. All rights reserved.},
author = {Cooper, Nancy and Khosravan, Reza and Erdmann, Carol and Fiene, John and Lee, Jean W.},
doi = {10.1016/j.jchromb.2006.02.060},
file = {:Users/solomon/Documents/Mendeley Desktop/Cooper et al. - 2006 - Quantification of uric acid, xanthine and hypoxanthine in human serum by HPLC for pharmacodynamic studies.pdf:pdf},
isbn = {1570-0232},
issn = {15700232},
journal = {Journal of Chromatography B: Analytical Technologies in the Biomedical and Life Sciences},
keywords = {HPLC,Human serum,Pharmacodynamic biomarkers,Uric acid,Xanthine oxidase inhibitor},
number = {1-2},
pages = {1--10},
pmid = {16631418},
title = {{Quantification of uric acid, xanthine and hypoxanthine in human serum by HPLC for pharmacodynamic studies}},
volume = {837},
year = {2006}
}
@article{Marciano2002,
abstract = {Traumatic brain injury (TBI) elicits a complex sequence of putative autodestructive and neuroprotective cellular cascades. It is hypothesized that the genomic responses of cells in the injured brain serve as the basis for these cascades. Traditional methods for analyzing differential gene expression following brain trauma demonstrate that immediate early genes, cytokines, transcription factors, and neurotrophic factors can all participate in the brain's active and directed response to injury, and may do so concurrently. It is this complexity and multiplicity of interrelated molecular mechanisms that has demanded new methods for comprehensive and parallel evaluation of putative as well as novel gene targets. Recent advances in DNA microarray technology have enabled the simultaneous evaluation of thousands of genes and the subsequent generation of massive amounts of biological data relevant to CNS injury. This emerging technology can serve to further current knowledge regarding recognized molecular cascades as well as to identify novel molecular mechanisms that occur throughout the post-traumatic period. The elucidation of the complex alterations in gene expression underlying the pathological sequelae following TBI is of central importance in the design of future therapeutic agents.},
author = {Marciano, Paolo G. and Eberwine, James H. and Ragupathi, Ramesh and Saatman, Kathryn E. and Meaney, David F. and McIntosh, Tracy K.},
doi = {10.1023/A:1020973308941},
file = {:Users/solomon/Documents/Mendeley Desktop/Marciano et al. - 2002 - Expression profiling following traumatic brain injury A review.pdf:pdf},
issn = {03643190},
journal = {Neurochemical Research},
keywords = {Cytokines,Gene expression,Immediate early genes,Microarray,Neurotrophic factors,Traumatic brain injury},
number = {10},
pages = {1147--1155},
pmid = {12462413},
title = {{Expression profiling following traumatic brain injury: A review}},
volume = {27},
year = {2002}
}
@article{Amaro2016,
abstract = {BACKGROUND AND PURPOSE: Identification of neuroprotective therapies in acute ischemic stroke is imperative. We report a predefined analysis of the URICO-ICTUS trial (Efficacy Study of Combined Treatment With Uric Acid and r-tPA in Acute Ischemic Stroke) assessing the efficacy of uric acid (UA) compared with placebo to prevent early ischemic worsening (EIW) and the relevance of collateral circulation. METHODS: URICO-ICTUS was a double-blind, placebo-controlled, phase 2b trial where a total of 411 patients treated with alteplase within 4.5 hours of stroke onset were randomized (1:1) to receive UA 1000 mg (n=211) or placebo (n=200) before the end of alteplase infusion. EIW defined an increment {\textgreater}/=4 points in the National Institutes of Health Stroke Scale score within 72 hours of treatment in the absence of hemorrhage or recurrent stroke. Logistic regression models assessed the interaction between therapy and the collateral circulation in 112 patients who had a pretreatment computed tomographic angiography. RESULTS: EIW occurred in 2 of 149 (1{\%}) patients with good outcome and 23 of 262 (9{\%}) patients with poor outcome (chi2; P=0.002). EIW occurred in 7 of 204 (3{\%}) patients treated with UA and in 18 of 200 (9{\%}) patients treated with placebo (chi2; P=0.01). There was a significant interaction between the efficacy of UA to prevent EIW and collaterals (P=0.029), with lower incidence in patients with good collaterals treated with UA compared with placebo (2{\%} versus 15{\%}, respectively; P=0.048). CONCLUSIONS: UA therapy may prevent EIW after acute stroke in thrombolysed patients. Optimal access of UA to its molecular targets through appropriate collaterals may modify the magnitude of the neuroprotective effect. CLINICAL TRIAL REGISTRATION: URL: http://www.clinicaltrials.gov. Unique identifier: NCT00860366.},
author = {Amaro, Sergio and Laredo, Carlos and Ren??, Arturo and Llull, Laura and Rudilosso, Salvatore and Obach, V??ctor and Urra, Xabier and Planas, Anna M. and Chamorro, ??ngel},
doi = {10.1161/STROKEAHA.116.014672},
file = {:Users/solomon/Documents/Mendeley Desktop/Amaro et al. - 2016 - Uric acid therapy prevents early ischemic stroke progression A tertiary analysis of the URICO-ICTUS trial (Efficac.pdf:pdf},
issn = {15244628},
journal = {Stroke},
keywords = {antioxidant,clinical trials,ischemic stroke,neuroprotection,treatment},
number = {11},
pages = {2874--2876},
pmid = {27758945},
title = {{Uric acid therapy prevents early ischemic stroke progression: A tertiary analysis of the URICO-ICTUS trial (Efficacy Study of Combined Treatment With Uric Acid and r-tPA in Acute Ischemic Stroke)}},
volume = {47},
year = {2016}
}
@article{Benet1979,
abstract = {None},
author = {Benet, Leslie Z and Galeazzi, Renato L},
doi = {10.1002/jps.2600680845},
file = {:Users/solomon/Documents/Mendeley Desktop/Benet, Galeazzi - 1979 - Noncompartmental determination of the steady‐state volume of distribution.pdf:pdf},
isbn = {0022-3549 (Print)},
issn = {15206017},
journal = {Journal of Pharmaceutical Sciences},
number = {2},
pages = {1071--1074},
pmid = {480170},
title = {{Noncompartmental determination of the steady‐state volume of distribution}},
volume = {73},
year = {1979}
}
@article{Kamper2013,
abstract = {Traumatic brain injury (TBI) refers to physical trauma to the brain that can lead to motor and cognitive dysfunctions. TBI is particularly serious in infants and young children, often leading to long-term functional impairments. Although clinical research is useful for quantifying and observing the effects of these injuries, few studies have empirically assessed the long-term effects of juvenile TBI (jTBI) on behavior and histology. After a controlled cortical impact delivered to postnatal 17. day old rats, functional abilities were measured after 3, 5, and 6. months using open field (activity levels), zero maze (anxiety-like behaviors), rotarod (sensorimotor abilities, coordination, and balance), and water maze (spatial learning and memory, swim speed, turn bias). Sensorimotor function was impaired for up to 6. months in jTBI animals, which showed no improvement from repeated test exposure. Although spatial learning was not impaired, spatial memory deficits were observed in jTBI animals starting at 3. months after injury. Magnetic resonance imaging and histological data revealed that the effects of jTBI were evolving for up to 6. months post-injury, with reduced cortical thickness, decreased corpus callosum area and CA1 neuronal cell death in jTBI animals distant to the impact site. These findings suggest that this model of jTBI produces long-term impairments comparable to those reported clinically. Although some deficits were stable over time, the variable nature of other deficits (e.g., memory) as well as changing properties of the lesion itself, suggest that the effects of a single jTBI produce a chronic brain disorder with long-term complications. ?? 2013 Elsevier Inc.},
author = {Kamper, Joel E. and Pop, Viorela and Fukuda, Andrew M. and Ajao, David O. and Hartman, Richard E. and Badaut, J??r??me},
doi = {10.1016/j.expneurol.2013.09.016},
isbn = {0014-4886},
issn = {00144886},
journal = {Experimental Neurology},
keywords = {Behavior,MRI,Neurodegeneration,Pediatric,Traumatic brain injury},
pages = {8--19},
pmid = {24076005},
publisher = {Elsevier Inc.},
title = {{Juvenile traumatic brain injury evolves into a chronic brain disorder: Behavioral and histological changes over 6months}},
url = {http://dx.doi.org/10.1016/j.expneurol.2013.09.016},
volume = {250},
year = {2013}
}
@article{Wray2005,
abstract = {The design and interpretation of genetic association studies depends on the relationship between the genotyped variants and the underlying functional variant, often parameterized as the squared correlation or r(2) measure of linkage disequilibrium between two loci. While it has long been recognized that placing a constraint on ther(2) between two loci also places a constraint on the difference in frequencies between the coupled alleles, this constraint has not been quantified. Here, quantification of this severe constraint is presented. For example, for r(2) {\textgreater}/= .8, the maximum difference in allele frequency is +/- .06 which occurs when one locus has allele frequency .5. For r(2) {\textgreater}/= .8 and allele frequency at one locus of .1, the maximum difference in allele frequency at the second locus is only +/- .02. The impact on the design and interpretation of association studies is discussed.},
author = {Wray, Naomi R},
doi = {10.1375/twin.8.2.87},
file = {:Users/solomon/Documents/Mendeley Desktop/Wray - 2005 - Allele frequencies and the r2 measure of linkage disequilibrium impact on design and interpretation of association studies.pdf:pdf},
isbn = {1832-4274 (Print)$\backslash$n1832-4274 (Linking)},
issn = {1832-4274},
journal = {Twin research and human genetics : the official journal of the International Society for Twin Studies},
keywords = {Algorithms,Chromosome Mapping,Data Interpretation,Gene Frequency,Gene Frequency: genetics,Genetic Markers,Genetic Markers: genetics,Genetic Variation,Genetic Variation: genetics,Genotype,Humans,Linkage Disequilibrium,Linkage Disequilibrium: genetics,Polymorphism,Research Design,Research Design: statistics {\&} numerical data,Single Nucleotide,Single Nucleotide: genetics,Statistical},
number = {2},
pages = {87--94},
pmid = {15901470},
title = {{Allele frequencies and the r2 measure of linkage disequilibrium: impact on design and interpretation of association studies.}},
url = {http://www.ncbi.nlm.nih.gov/pubmed/15901470},
volume = {8},
year = {2005}
}
@article{Balyan2017,
author = {Balyan, Rajiv and Mecoli, Marc and Venkatasubramanian, Raja and Chidambaran, Vidya and Kamos, Nichole and Clay, Smokey and Moore, David L and Mavi, Jagroop and Glover, Chris D and Szmuk, Peter and Vinks, Alexander and Sadhasivam, Senthilkumar},
doi = {10.2217/pgs-2016-0183},
file = {:Users/solomon/Documents/Mendeley Desktop/Balyan et al. - 2017 - CYP2D6 pharmacogenetic and oxycodone pharmacokinetic association study in pediatric surgical patients.pdf:pdf},
issn = {1462-2416},
journal = {Pharmacogenomics},
keywords = {cyp2d6,oxycodone,pediatrics,pharmacogenetics,pharmacokinetics},
month = {mar},
number = {4},
pages = {337--348},
title = {{CYP2D6 pharmacogenetic and oxycodone pharmacokinetic association study in pediatric surgical patients}},
url = {http://www.futuremedicine.com/doi/10.2217/pgs-2016-0183},
volume = {18},
year = {2017}
}
@article{Zhao2012,
abstract = {BACKGROUND: Accumulating evidence has demonstrated that over-expression of Neuroglobin (Ngb) is neuroprotective against hypoxic/ischemic brain injuries. In this study we tested the neuroprotective effects of Ngb over-expression against traumatic brain injury (TBI) in mice. RESULTS: Both Ngb over-expression transgenic (Ngb-Tg) and wild-type (WT) control mice were subjected to TBI induced by a controlled cortical impact (CCI) device. TBI significantly increased Ngb expression in the brains of both WT and Ngb-Tg mice, but Ngb-Tg mice had significantly higher Ngb protein levels at the pre-injury baseline and post-TBI. Production of oxidative tissue damage biomarker 3NT in the brain was significantly reduced in Ngb-Tg mice compared to WT controls at 6 hours after TBI. The traumatic brain lesion volume was significantly reduced in Ngb Tg mice compared to WT mice at 3 weeks after TBI; however, there were no significant differences in the recovery of sensorimotor and spatial memory functional deficits between Ngb-Tg and WT control mice for up to 3 weeks after TBI. CONCLUSION: Ngb over-expression reduced traumatic lesion volume, which might partially be achieved by decreasing oxidative stress.},
author = {Zhao, S and Yu, Z and Zhao, G and Xing, C and Hayakawa, K and Whalen, M J and Lok, J M and Lo, E H and Wang, X},
doi = {10.1186/1471-2202-13-67},
file = {:Users/solomon/Documents/Mendeley Desktop/Zhao et al. - 2012 - Neuroglobin-overexpression reduces traumatic brain lesion size in mice.pdf:pdf},
isbn = {1471-2202 (Electronic); 1471-2202 (Linking)},
issn = {1471-2202},
journal = {BMC Neurosci},
keywords = {*Seizures/etiology/metabolism/prevention {\&} control,Analysis of Variance,Animals,Brain Injuries/*complications/pathology,Cerebral Cortex/*metabolism,Disease Models, Animal,Gait Disorders, Neurologic/etiology,Gene Expression Regulation/genetics/*physiology,Globins/genetics/*metabolism,Maze Learning/physiology,Memory Disorders/etiology,Mice,Mice, Inbred C57BL,Mice, Transgenic,Nerve Tissue Proteins/genetics/*metabolism,Neuroprotective Agents/*therapeutic use,Space Perception/physiology,Tyrosine/analogs {\&} derivatives/metabolism},
number = {1},
pages = {67},
pmid = {22703519},
publisher = {BMC Neuroscience},
title = {{Neuroglobin-overexpression reduces traumatic brain lesion size in mice}},
url = {http://www.ncbi.nlm.nih.gov/entrez/query.fcgi?cmd=Retrieve{\&}db=pubmed{\&}dopt=Abstract{\&}list{\_}uids=22703519{\&}query{\_}hl=1http://www.biomedcentral.com/content/pdf/1471-2202-13-67.pdf 全文链接},
volume = {13},
year = {2012}
}
@article{Hori2004,
abstract = {The purpose of the present study was to clarify the expression, transport properties and regulation of ATP-binding cassette G2 (ABCG2) transporter at the rat blood-brain barrier (BBB). The rat homologue of ABCG2 (rABCG2) was cloned from rat brain capillary fraction. In rABCG2-transfected HEK293 cells, rABCG2 was detected as a glycoprotein complex bridged by disulfide bonds, possibly a homodimer. The protein transported mitoxantrone and BODIPY-prazosin. In rat brain capillary fraction, rABCG2 protein was also detected as a glycosylated and disulfide-linked complex. Immunohistochemical analysis revealed that rABCG2 was localized mainly on the luminal side of rat brain capillaries, suggesting that rABCG2 is involved in brain-to-blood efflux transport. For the regulation study, conditionally immortalized rat brain capillary endothelial (TR-BBB13), astrocyte (TR-AST4) and pericyte (TR-PCT1) cell lines were used as an in vitro BBB model. Following treatment of TR-BBB13 cells with conditioned medium of TR-AST4 cells, the Ko143 (an ABCG2-specific inhibitor)-sensitive transport activity and rABCG2 mRNA level were significantly increased, whereas conditioned medium of TR-PCT1 cells had no effect. These results suggest that rat brain capillaries express functional rABCG2 protein and that the transport activity of the protein is up-regulated by astrocyte-derived soluble factor(s) concomitantly with the induction of rABCG2 mRNA.},
author = {Hori, Satoko and Ohtsuki, Sumio and Tachikawa, Masanori and Kimura, Norihisa and Kondo, Tetsu and Watanabe, Masahiko and Nakashima, Emi and Terasaki, Tetsuya},
doi = {10.1111/j.1471-4159.2004.02537.x},
file = {:Users/solomon/Documents/Mendeley Desktop/Hori et al. - 2004 - Functional expression of rat ABCG2 on the luminal side of brain capillaries and its enhancement by astrocyte-derive.pdf:pdf},
isbn = {0022-3042 (Print)$\backslash$n0022-3042 (Linking)},
issn = {00223042},
journal = {Journal of Neurochemistry},
keywords = {ABCG2,Astrocyte,Blood-brain barrier,In vitro BBB model},
number = {3},
pages = {526--536},
pmid = {15255930},
title = {{Functional expression of rat ABCG2 on the luminal side of brain capillaries and its enhancement by astrocyte-derived soluble factor(s)}},
volume = {90},
year = {2004}
}
@article{Surgeons.2007,
abstract = {First lines: TRAUMATIC BRAIN INJURY (TBI) is a major cause of disability, death, and economic cost to our society. One of the central concepts that emerged from research is that all neurological damage from TBI does not occur at the moment of impact, but evolves over the ensuing hours and days. Furthermore, improved outcome results when these secondary, delayed insults, resulting in reduced cerebral perfusion to the injured brain, are prevented or respond to treatment. This is reflected in the progressive and significant reduction in severe TBI mortality from 50{\%} to 35{\%} to 25{\%} and lower over the last 30 years, even when adjusted for injury severity, age and other admission prognostic parameters.1 This trend in reduced mortality and improved outcomes from TBI has been subsequent to the use of evidence-based protocols that emphasize monitoring and maintaining adequate cerebral perfusion.2,3},
author = {Surgeons., Brain Trauma Foundation; American Association of Neurological Surgeons; Congress of Neurological},
doi = {10.1089/neu.2007.9990},
file = {:Users/solomon/Documents/Mendeley Desktop/Surgeons. - 2007 - Guidelines for the Management of Severe Traumatic Brain Injury 3rd Edition.pdf:pdf},
isbn = {087977151F},
issn = {08977151},
journal = {Journal of neurosurgery},
keywords = {Practice Guideline,Severe Traumatic Brain Injury,Traumatic Brain Injury,Traumatic Brain Injury: therapy},
number = {212},
pages = {S1--106},
pmid = {17511534},
title = {{Guidelines for the Management of Severe Traumatic Brain Injury 3rd Edition}},
url = {http://www.ncbi.nlm.nih.gov/pubmed/17511534},
volume = {24, Suppl },
year = {2007}
}
@article{Carden1987,
abstract = {Monoclonal antibodies (mAbs) to rat neurofilament (NF) proteins NF-L, NF-M, and NF-H were used to examine the developmental programs of NF expression in rat embryos. The ability of these mAbs to recognize differentially phosphorylated states of NF-M and NF-H (Lee et al., 1987, the preceding paper) was exploited in order to examine the temporal and spatial patterns of NF phosphorylation during early neuronal development in vivo. NF proteins were first detected on the twelfth day postfertilization (E12) using NF-L- or NF-M-specific mAbs. By E13, the coexpression of NF-L and NF-M was widespread, reflecting dramatic increases of immunoreactivity to both subunits. Partial phosphorylation, denoted P[+], of NF-M was already present in perikarya and neurites of E12 neurons. Extensively phosphorylated, or P[+++], isoforms of NF-M appeared in E13 axons, thereby establishing a proximodistal gradient of NF phosphorylation during the earliest phase of NF expression. Immunoblots of tissue homogenates revealed that most NF-M of E13 embryos exists in a partially phosphorylated, or P[+], isoform. Unequivocal staining for NF-H first appeared at E15, a time at which NF-L and NF-M had already attained their adult patterns of immunocytochemical staining. Levels of NF-H were extremely low at E15 but could be detected in all of its differentially phosphorylated states, i.e., nonphosphorylated P[-], partly P[+], and highly P[+++] phosphorylated isoforms. P[+++] isoforms of NF-H were restricted to the distal portions of E15 axons, although staining of more proximal axons, like those in adult, was noted by E17. Immunoblots of E17 embryos revealed most NF-H as P[-] and P[+] isoforms. Quantities of immunoreactive NF-H increased very slowly and remained well below those of NF-M and NF-L for several weeks beyond birth. These results show that sequential forms of NFs are expressed by developing and maturing neurons throughout the nervous system. An "immature" form of NFs, composed of NF-M and NF-L, appears to function in establishing the neuronal phenotype and in initiating and maintaining neurite outgrowth. Addition of NF-H confers a "mature" state to the NF. This delayed expression of NF-H is a slow and graduated process that coincides in time with the stabilization of neuronal circuitries and may be important in modulating axonal events, such as the slowing of cytoskeletal transport and the growth of axonal caliber.},
author = {Carden, M J and Trojanowski, J Q and Schlaepfer, W W and Lee, V M},
file = {:Users/solomon/Documents/Mendeley Desktop/Carden et al. - 1987 - Two-stage expression of neurofilament polypeptides during rat neurogenesis with early establishment of adult phos.pdf:pdf},
isbn = {0270-6474 (Print)$\backslash$n0270-6474 (Linking)},
issn = {0270-6474},
journal = {The Journal of neuroscience : the official journal of the Society for Neuroscience},
number = {11},
pages = {3489--504},
pmid = {3119790},
title = {{Two-stage expression of neurofilament polypeptides during rat neurogenesis with early establishment of adult phosphorylation patterns.}},
url = {http://www.ncbi.nlm.nih.gov/pubmed/3119790},
volume = {7},
year = {1987}
}
@article{Hajj2017,
abstract = {Despite increased attention on assessment and management, pain remains the most persistent symptom in patients with cancer, in particular in end-of-life settings, with detrimental impact on their quality-of-life (QOL). We conducted this study to evaluate the added value of determining some genetic and non-genetic factors to optimize cancer pain treatment. Eighty-nine patients were included in the study for the evaluation of palliative cancer pain management. The regression analysis showed that age, OPRM1 single nucleotide polymorphism (SNP), as well as the duration of morphine treatment were significantly associated with morphine doses at 24 h (given by infusion pump; p = 0.043, 0.029, and {\textless}0.001, respectively). The mean doses of morphine decreased with age but increased with the duration of morphine treatment. In addition, patients with AG genotype c.118A{\textgreater}G OPRM1 needed a higher dose of morphine than AA patients. Moreover, metastases, OPRM1 SNP, age, and gender were significantly associated with the QOL in our population. In particular, AA patients for OPRM1 SNP had significantly lower cognitive function than AG patients, a result not previously reported in the literature. These findings could help increase the effectiveness of morphine treatment and enhance the QOL of patients in regards to personalized medicine.},
author = {Hajj, Aline and Halepian, Lucine and Osta, Nada and Chahine, Georges and Kattan, Joseph and {Rabbaa Khabbaz}, Lydia},
doi = {10.3390/ijms18040669},
file = {:Users/solomon/Documents/Mendeley Desktop/Hajj et al. - 2017 - OPRM1 c.118AG Polymorphism and Duration of Morphine Treatment Associated with Morphine Doses and Quality-of-Life in.pdf:pdf},
issn = {1422-0067},
journal = {Int. J. Mol. Sci.},
keywords = {ABCB1,COMT,OPRM1,cancer,morphine,pain,pharmacogenetics,polymorphism,quality-of-life},
month = {mar},
number = {5},
pages = {669},
pmid = {28346387},
publisher = {Multidisciplinary Digital Publishing Institute (MDPI)},
title = {{OPRM1 c.118A{\textgreater}G Polymorphism and Duration of Morphine Treatment Associated with Morphine Doses and Quality-of-Life in Palliative Cancer Pain Settings}},
url = {http://www.ncbi.nlm.nih.gov/pubmed/28346387 http://www.pubmedcentral.nih.gov/articlerender.fcgi?artid=PMC5412267 http://www.mdpi.com/1422-0067/18/4/669},
volume = {18},
year = {2017}
}
@article{Lazarowski2007,
abstract = {It is estimated 20-25{\%} of the epileptic patients fails to achieve good control with the different antiepileptic drugs (AEDs) treatments, developing refractory epilepsy (RE). Discovered first in cancer, the activity of P-glycoprotein (P-gp) and others ABC transporters as multidrug-resistance-associated proteins (MRPs) and breast cancer resistant protein (BCRP) are directly related with the refractoriness. We have observed the overexpression of these all transporters in the brain of patients with RE, and according with other authors, all these data suggests an active drug efflux from brain. Both constitutive and seizure induced brain P-gp overexpression was also suggested. As confirmation of these clinical evidences, different models of experimental epilepsy have demonstrated P-gp overexpression on blood brain barrier (BBB) and brain parenchyma cells, as astrocytes and neurons. In our model, early P-pg detection in vessel-related cells and later additional P-gp detection in neurons, correlated with the gradual loss of protective effect of phenytoin. The progressive neuronal P-gp expression, depending on intensity and time-constancy of seizure-injury, was in agreement with the development of "P-gp-positive seizure-axis" proposed by Kwan {\&} Brodie, who also showed that the development of RE directly correlated with the number and frequency of seizures before initiation of drug therapy. P-gp expression in excretory organs suggests that P-gp have a central role in drug elimination. Persistent low levels of AEDs in plasma and P-gp brain overexpression in several RE pediatric patients were reported. We also observed in adult RE patients, an increased liver clearance of 99mTc-hexakis-2-methoxyisobutylisonitrile (99mTc-MIBI) (a P-gp substrate), and the surgically treated cases showed P-gp brain overexpression. These results suggest the systemic hyperactivity of P-gp in RE patients, including brain P-gp over-expression should be suspected when persistent subtherapeutic levels of AEDs in plasma are detected. P-gp neuronal expression described in both clinical and experimental reports indicates that additional mechanisms could be operative from seizure-affected P-gp-positive neurons, due to AEDs targets are expressed at membrane level. An alternative mechanism was demonstrated in P-gp-expressed cells that exhibit lower membrane potential (Deltapsi(0)=-10 to -20) compared to normal physiological Deltapsi(0) of -60 mV. Under this situation and irrespective to the P-gp pharmacoresistant property or type of drug treatment selected, P-gp-expressed neurons could increase their sensitivity to new seizures perhaps as an epileptogenic mechanism. The understanding of properties of these ABC transporters can offer new tools for better selection of more effective preventive or therapeutic strategies and avoid the invasive surgical treatments for RE.},
author = {Lazarowski, Alberto and Czornyj, Liliana and Lubienieki, Fabiana and Girardi, Elena and Vazquez, Silvia and D'Giano, Carlos},
doi = {10.1111/j.1528-1167.2007.01302.x},
file = {:Users/solomon/Documents/Mendeley Desktop/Lazarowski et al. - 2007 - ABC transporters during epilepsy and mechanisms underlying multidrug resistance in refractory epilepsy.pdf:pdf},
isbn = {0013-9580 (Print)$\backslash$n0013-9580 (Linking)},
issn = {00139580},
journal = {Epilepsia},
keywords = {99mTc-MIBI,BCRP,MDR,MVP,Refractory epilepsy},
number = {SUPPL. 5},
pages = {140--149},
pmid = {17910594},
title = {{ABC transporters during epilepsy and mechanisms underlying multidrug resistance in refractory epilepsy}},
volume = {48},
year = {2007}
}
@article{Potts2006,
abstract = {Traumatic brain injury (TBI) is the leading cause of morbidity and mortality among children and both clinical and experimental data reveal that the immature brain is unique in its response and vulnerability to TBI compared to the adult brain. Current therapies for pediatric TBI focus on physiologic derangements and are based primarily on adult data. However, it is now evident that secondary biochemical perturbations play an important role in the pathobiology of pediatric TBI and may provide specific therapeutic targets for the treatment of the head-injured child. In this review, we discuss three specific components of the secondary pathogenesis of pediatric TBI-- inflammation, oxidative injury, and iron-induced damage-- and potential therapeutic strategies associated with each. The inflammatory response in the immature brain is more robust than in the adult and characterized by greater disruption of the blood-brain barrier and elaboration of cytokines. The immature brain also has a muted response to oxidative stress compared to the adult due to inadequate expression of certain antioxidant molecules. In addition, the developing brain is less able to detoxify free iron after TBI-induced hemorrhage and cell death. These processes thus provide potential therapeutic targets that may be tailored to pediatric TBI, including anti-inflammatory agents such as minocycline, antioxidants such as glutathione peroxidase, and the iron chelator deferoxamine.},
author = {Potts, Mathew B and Koh, Seong-Eun and Whetstone, William D and Walker, Breset A and Yoneyama, Tomoko and Claus, Catherine P and Manvelyan, Hovhannes M and Noble-Haeusslein, Linda J},
doi = {10.1016/j.nurx.2006.01.006},
file = {:Users/solomon/Documents/Mendeley Desktop/Potts et al. - 2006 - Traumatic injury to the immature brain inflammation, oxidative injury, and iron-mediated damage as potential thera.pdf:pdf},
isbn = {1545-5343 (Print) 1545-5343 (Linking)},
issn = {1545-5343},
journal = {NeuroRx : the journal of the American Society for Experimental NeuroTherapeutics},
keywords = {Animals,Brain,Brain Injuries,Brain Injuries: complications,Brain Injuries: metabolism,Brain Injuries: pathology,Brain Injuries: therapy,Brain: growth {\&} development,Child,Encephalitis,Encephalitis: etiology,Encephalitis: therapy,Humans,Iron,Iron: metabolism,Oxidative Stress},
number = {2},
pages = {143--53},
pmid = {16554253},
title = {{Traumatic injury to the immature brain: inflammation, oxidative injury, and iron-mediated damage as potential therapeutic targets.}},
url = {http://www.sciencedirect.com/science/article/pii/S1545534306000186},
volume = {3},
year = {2006}
}
@article{Zappaterra2013,
abstract = {The CSF is a complex fluid with a dynamically varying proteome throughout development and in adulthood. During embryonic development, the nascent CSF differentiates from the amniotic fluid upon closure of the anterior neural tube. CSF volume then increases over subsequent days as the neuroepithelial progenitor cells lining the ventricles and the choroid plexus generate CSF. The embryonic CSF contacts the apical, ventricular surface of the neural stem cells of the developing brain and spinal cord. CSF provides crucial fluid pressure for the expansion of the developing brain and distributes important growth promoting factors to neural progenitor cells in a temporally-specific manner. To investigate the function of the CSF, it is important to isolate pure samples of embryonic CSF without contamination from blood or the developing telencephalic tissue. Here, we describe a technique to isolate relatively pure samples of ventricular embryonic CSF that can be used for a wide range of experimental assays including mass spectrometry, protein electrophoresis, and cell and primary explant culture. We demonstrate how to dissect and culture cortical explants on porous polycarbonate membranes in order to grow developing cortical tissue with reduced volumes of media or CSF. With this method, experiments can be performed using CSF from varying ages or conditions to investigate the biological activity of the CSF proteome on target cells.},
author = {Zappaterra, Mauro W and LaMantia, Anthony S and Walsh, Christopher A and Lehtinen, Maria K},
doi = {10.3791/50333},
file = {:Users/solomon/Documents/Mendeley Desktop/Zappaterra et al. - 2013 - Isolation of cerebrospinal fluid from rodent embryos for use with dissected cerebral cortical explants.pdf:pdf},
isbn = {1940-087X (Electronic)$\backslash$r1940-087X (Linking)},
issn = {1940-087X},
journal = {Journal of visualized experiments : JoVE},
keywords = {11,2013,3,anatomy,animal model,biomedical,brain,cellular biology,cerebral cortex,cerebral cortical explant,cerebrospinal fluid,csf,culture,date published,developmental biology,engineering,isolation,issue 73,medicine,mouse,neural stem cells,neurobiology,neuroscience,nscs,physiology,stem cell biology,stem cells,surgery,tissue,ventricular embryonic csf},
number = {73},
pages = {e50333},
pmid = {23524481},
title = {{Isolation of cerebrospinal fluid from rodent embryos for use with dissected cerebral cortical explants.}},
url = {http://www.ncbi.nlm.nih.gov/pubmed/23524481{\%}5Cnhttp://www.pubmedcentral.nih.gov/articlerender.fcgi?artid=PMC3635544},
year = {2013}
}
@article{Cresci2014,
abstract = {BACKGROUND Clopidogrel is recommended after acute myocardial infarction but has variable efficacy and safety, in part related to the effect of cytochrome P450 (CYP) polymorphisms on its metabolism. The effect of CYP polymorphisms on cardiovascular events among clopidogrel-treated patients after acute myocardial infarction remains controversial, and no studies to date have investigated the association of CYP variants with outcomes in black patients. METHODS AND RESULTS Subjects (2732: 2062 whites; 670 blacks) hospitalized with acute myocardial infarction enrolled in the prospective, multicenter TRIUMPH study were genotyped for CYP polymorphisms. The majority of whites (79{\%}) and blacks (64.4{\%}) were discharged on clopidogrel. Among whites, carriers of the loss-of-function CYP2C19*2 allele had significantly increased 1-year mortality (adjusted hazards ratio [HR]: 1.70; confidence interval [CI]: 1.01-2.86; P=0.046) and a trend toward increased rate of recurrent MI (adjusted HR: 2.10; CI: 0.95-4.63; P=0.066). Among blacks, increased 1-year mortality was associated with the gain-of-function CYP2C19*17 allele (adjusted HR for *1/*17 versus *1/*1: 2.02; CI: 0.92-4.44; *17/*17 versus *1/*1: 8.97; CI: 3.34-24.10; P{\textless}0.0001) and the CYP1A2*1C allele (adjusted HR for *1/*1C versus *1/*1: 1.89; CI: 0.85-4.22; *1C/*1C versus *1/*1: 4.96; CI: 1.69-14.56; P=0.014). Bleeding events were significantly more common among black carriers of CYP2C19*17 or CYP1A2*1C. CONCLUSIONS Both loss-of-function and gain-of-function CYP polymorphisms affecting clopidogrel metabolism are associated with increased mortality among clopidogrel-treated patients after acute myocardial infarction; the specific polymorphism and the putative mechanism vary according to race.},
author = {Cresci, Sharon and Depta, Jeremiah P. and Lenzini, Petra A. and Li, Allie Y. and Lanfear, David E. and Province, Michael A. and Spertus, John A. and Bach, Richard G.},
doi = {10.1161/CIRCGENETICS.113.000303},
file = {:Users/solomon/Documents/Mendeley Desktop/Cresci et al. - 2014 - Cytochrome p450 gene variants, race, and mortality among clopidogrel-treated patients after acute myocardial infa.pdf:pdf},
issn = {19423268},
journal = {Circulation: Cardiovascular Genetics},
keywords = {Clopidogrel,Genetic variation,Mortality,Myocardial infarction},
number = {3},
pages = {277--286},
pmid = {24762860},
title = {{Cytochrome p450 gene variants, race, and mortality among clopidogrel-treated patients after acute myocardial infarction}},
volume = {7},
year = {2014}
}
@article{Walker2013,
abstract = {Traumatic brain injury (TBI) results in significant disability due to cognitive deficits particularly in attention, learning and memory, and higher-order executive functions. The role of TBI in chronic neurodegeneration and the development of neurodegenerative diseases including Alzheimer's disease (AD), Parkinson's disease (PD), Amyotrophic Lateral Sclerosis (ALS) and most recently chronic traumatic encephalopathy (CTE) is of particular importance. However, despite significant effort very few therapeutic options exist to prevent or reverse cognitive impairment following TBI. In this review, we present experimental evidence of the known secondary injury mechanisms which contribute to neuronal cell loss, axonal injury, and synaptic dysfunction and hence cognitive impairment both acutely and chronically following TBI. In particular we focus on the mechanisms linking TBI to the development of two forms of dementia: AD and CTE. We provide evidence of potential molecular mechanisms involved in modulating A$\beta$ and Tau following TBI and provide evidence of the role of these mechanisms in AD pathology. Additionally we propose a mechanism by which A$\beta$ generated as a direct result of TBI is capable of exacerbating secondary injury mechanisms thereby establishing a neurotoxic cascade that leads to chronic neurodegeneration.},
author = {Walker, KR and Tesco, G},
doi = {10.3389/fnagi.2013.00029},
file = {:Users/solomon/Documents/Mendeley Desktop/Walker, Tesco - 2013 - Molecular mechanisms of cognitive dysfunction following traumatic brain injury.pdf:pdf},
isbn = {1663-4365},
issn = {16634365},
journal = {Frontiers in Aging Neuroscience},
keywords = {Alzheimer's,A$\beta$,CTE,Cognition,TBI,Tau},
number = {JUL},
pages = {1--25},
pmid = {23847533},
title = {{Molecular mechanisms of cognitive dysfunction following traumatic brain injury}},
volume = {5},
year = {2013}
}
@article{Empey2010,
author = {Empey, PE},
doi = {10.1097/CCM.0b013e3181de09f8},
file = {:Users/solomon/Documents/Mendeley Desktop/Empey - 2010 - Genetic predisposition to adverse drug reactions in the intensive care unit.pdf:pdf},
issn = {1530-0293 (Electronic) 0090-3493 (Linking)},
journal = {Crit Care Med},
number = {6 Suppl},
pages = {S106--16},
title = {{Genetic predisposition to adverse drug reactions in the intensive care unit}},
type = {Journal Article},
url = {https://www.ncbi.nlm.nih.gov/pubmed/20502164},
volume = {38},
year = {2010}
}
@article{Tavazzi2005,
abstract = {Objectives: To set up a novel simple, sensitive, and reliable ion-pairing HPLC method for the synchronous separation of several purines, pyrimidines, N-acetylated amino acids, and dicarboxylic acids for the chemical diagnosis and screening of inborn errors of metabolism (IEM). Design and methods: The separation was set up using a Hypersil C-18, 5-??m particle size, 250 ?? 4.6 mm column, and a step gradient using two buffers and tetrabutylammonium hydroxide as the pairing reagent. A highly sensitive diode array UV detector was set up at a wavelength between 200 and 300 nm that revealed purines and pyrimidines at 260 nm and other compounds at 206 nm. Results: Compounds were determined in the plasma of 15 healthy adults, in the urine of 50 healthy subjects (1-3 years, 4-6 years, 8-10 years, 12-18 years, 25-35 years), and in 10 non-pathological amniotic fluid samples. To assess the validity of the chemical diagnosis of IEM, plasma and urine samples were analyzed in patients affected by Canavan disease (n = 10; mean age 4.6 ?? 2.3). Low plasma levels of N-acetylaspartate (16.96 ?? 19.57 ??mol/L plasma; not detectable in healthy adults) and dramatically high urinary N-acetylaspartate concentrations (1872.03 ?? 631.86 ??mol/mmol creatinine; 450 times higher than that which was observed in age-matched controls) were recorded. Neither N-acetylglutamate nor N-acetylaspartylglutamate could be detected in the plasma or urine of controls or patients with Canavan disease. Conclusions: The results demonstrate the suitability of the present ion-pairing HPLC separation with UV detection of cytosine, cytidine, creatinine, uracil, uridine, ??-pseudouridine, adenine, 3-methyladenine, hypoxanthine, xanthine, xanthosine, inosine, guanosine, ascorbic acid, thymine, thymidine, uric acid, 1-methyluric acid, orotic acid, N-acetylaspartate, N-acetylglutamate, N-acetylaspartylglutamate, malonic acid, methylmalonic acid, GSH, and GSSG as a reliable method for the prenatal and neonatal chemical diagnosis and screening of IEM using biological fluids. ?? 2005 The Canadian Society of Clinical Chemists. All rights reserved.},
author = {Tavazzi, Barbara and Lazzarino, Giuseppe and Leone, Paola and Amorini, Angela Maria and Bellia, Francesco and Janson, Christopher G. and {Di Pietro}, Valentina and Ceccarelli, Lia and Donzelli, Sonia and Francis, Jeremy S. and Giardina, Bruno},
doi = {10.1016/j.clinbiochem.2005.08.002},
file = {:Users/solomon/Documents/Mendeley Desktop/Tavazzi et al. - 2005 - Simultaneous high performance liquid chromatographic separation of purines, pyrimidines, N-acetylated amino acid.pdf:pdf},
isbn = {0009-9120},
issn = {00099120},
journal = {Clinical Biochemistry},
keywords = {Biological fluids,Canavan disease,Chemical diagnosis,Dicarboxylic acids,HPLC,Inborn errors of metabolism,N-acetylated amino acids,Prenatal and neonatal screening,Purines,Pyrimidines},
number = {11},
pages = {997--1008},
pmid = {16139832},
title = {{Simultaneous high performance liquid chromatographic separation of purines, pyrimidines, N-acetylated amino acids, and dicarboxylic acids for the chemical diagnosis of inborn errors of metabolism}},
volume = {38},
year = {2005}
}
@article{Richet1995,
abstract = {At the end of the 18th century, as soon as modern chemistry was created, dedicated physicians tried to apply it to medicine. A rewarding field was that of urinary lithiasis. Stones offered a sufficient amount of a relatively pure chemical present in the body. Indeed, urine and the solidified matter was within the analytical grasp of the existing techniques. The first step was made by Scheele in Sweden who identified uric acid in calculi and normal human urine. During the following thirty years, Fourcroy and Vauquelin in Paris and Wollaston, Pearson, Marcet and Prout in London identified the various salts, uric acid, urate, various phosphates, oxalate, calcium, ammonium and magnesium forming current calculi. The conditions of their solubility in vitro were described. Even rare components such as cystine and xanthine were unraveled in London. The impetus given by these studies offered a good start to clinical chemistry in general and to the understanding of urinary lithiasis in particular. This led to the discovery of the corresponding solutes in the urine of such patients. Unfortunately, during the following decades nothing was added to these chemical investigations, and most of what had been acquired was forgotten. Research was concentrated on clinical pathology. The implementation of chemistry to medicine had to wait.},
author = {Richet, G},
doi = {10.1038/ki.1995.364},
file = {:Users/solomon/Documents/Mendeley Desktop/Richet - 1995 - The chemistry of urinary stones around 1800 a first in clinical chemistry.pdf:pdf},
issn = {0085-2538 (Print)},
journal = {Kidney international},
keywords = {Chemistry, Clinical,History, 18th Century,Humans,Urinary Bladder Calculi,Urinary Calculi,chemistry,history},
number = {3},
pages = {876--886},
pmid = {7474678},
title = {{The chemistry of urinary stones around 1800: a first in clinical chemistry.}},
volume = {48},
year = {1995}
}
@article{Dougall2015,
abstract = {BACKGROUND Traumatic brain injury (TBI) is a major cause of chronic disability. Worldwide, it is the leading cause of disability in the under 40s, resulting in severe disability in some 150 to 200 million people per annum. In addition to mood and behavioural problems, cognition-particularly memory, attention and executive function-are commonly impaired by TBI. Cognitive problems following TBI are one of the most important factors in determining people's subjective well-being and their quality of life. Drugs are widely used in an attempt to improve cognitive functions. Whilst cholinergic agents in TBI have been reviewed, there has not yet been a systematic review or meta-analysis of the effect on chronic cognitive problems of all centrally acting pharmacological agents. OBJECTIVES To assess the effects of centrally acting pharmacological agents for treatment of chronic cognitive impairment subsequent to traumatic brain injury in adults. SEARCH METHODS We searched ALOIS-the Cochrane Dementia and Cognitive Improvement Group's Specialised Register-on 16 November 2013, 23 February 2013, 20 January 2014, and 30 December 2014 using the terms: traumatic OR TBI OR "brain injury" OR "brain injuries" OR TBIs OR "axonal injury" OR "axonal injuries". ALOIS contains records of clinical trials identified from monthly searches of a number of major healthcare databases, numerous trial registries and grey literature sources. Supplementary searches were also performed in MEDLINE, EMBASE, PsycINFO, The Cochrane Library, CINAHL, LILACs, ClinicalTrials.gov, the World Health Organization (WHO) Portal (ICTRP) and Web of Science with conference proceedings. SELECTION CRITERIA We included randomised controlled trials (RCTs) assessing the effectiveness of any one centrally acting pharmacological agent that affects one or more of the main neurotransmitter systems in people with chronic traumatic brain injury; and there had to be a minimum of 12 months between the injury and entry into the trial. DATA COLLECTION AND ANALYSIS Two review authors examined titles and abstracts of citations obtained from the search. Relevant articles were retrieved for further assessment. A bibliographic search of relevant papers was conducted. We extracted data using a standardised tool, which included data on the incidence of adverse effects. Where necessary we requested additional unpublished data from study authors. Risk of bias was assessed by a single author. MAIN RESULTS Only four studies met the criteria for inclusion, with a total of 274 participants. Four pharmacological agents were investigated: modafinil (51 participants); (-)-OSU6162, a monoamine stabiliser (12 participants of which six had a TBI); atomoxetine (60 participants); and rivastigmine (157 participants). A meta-analysis could not be performed due to the small number and heterogeneity of the studies.All studies examined cognitive performance, with the majority of the psychometric sub-tests showing no difference between treatment and placebo (n = 274, very low quality evidence). For (-)-OSU6162 modest superiority over placebo was demonstrated on three measures, but markedly inferior performance on another. Rivastigmine was better than placebo on one primary measure, and a single cognitive outcome in a secondary analysis of a subgroup with more severe memory impairment at baseline. The study of modafinil assessed clinical global improvement (n = 51, low quality evidence), and did not find any difference between treatment and placebo. Safety, as measured by adverse events, was reported by all studies (n = 274, very low quality evidence), with significantly more nausea reported by participants who received rivastigmine compared to placebo. There were no other differences in safety between treatment and placebo. No studies reported any deaths. AUTHORS' CONCLUSIONS There is insufficient evidence to determine whether pharmacological treatment is effective in chronic cognitive impairment in TBI. Whilst there is a positive finding for rivastigmine on one primary measure, all other primary measures were not better than placebo. The positive findings for (-)-OSU6162 are interpreted cautiously as the study was small (n = 6). For modafinil and atomoxetine no positive effects were found. All four drugs appear to be relatively well tolerated, although evidence is sparse.},
author = {Dougall, D and Poole, No and Agrawal, N},
doi = {10.1002/14651858.CD009221.pub2},
file = {:Users/solomon/Documents/Mendeley Desktop/Dougall, Poole, Agrawal - 2015 - Pharmacotherapy for chronic cognitive impairment in traumatic brain injury.pdf:pdf},
issn = {1469-493X},
journal = {The Cochrane database of systematic reviews},
month = {dec},
number = {12},
pages = {CD009221},
pmid = {26624881},
title = {{Pharmacotherapy for chronic cognitive impairment in traumatic brain injury.}},
url = {http://www.ncbi.nlm.nih.gov/pubmed/26624881},
year = {2015}
}
@article{Zweckberger2014,
abstract = {Following traumatic brain injury (TBI) SUR1-regulated NCCa-ATP (SUR1/TRPM4) channels are transcriptionally up-regulated in ischemic astrocytes, neurons, and capillaries. ATP depletion results in depolarization and opening of the channel leading to cytotoxic edema. Glibenclamide is an inhibitor of SUR-1 and, thus, might prevent cytotoxic edema and secondary brain damage following TBI.Anesthetized adult Sprague-Dawley rats underwent parietal craniotomy and were subjected to controlled cortical impact injury (CCI). Glibenclamide was administered as a bolus injection 15. min after CCI injury and continuously via osmotic pumps throughout 7. days. In an acute trial (180. min) mean arterial blood pressure, heart rate, intracranial pressure, encephalographic activity, and cerebral metabolism were monitored. Brain water content was assessed gravimetrically 24. h after CCI injury and contusion volumes were measured by MRI scanning technique at 8. h, 24. h, 72. h, and 7. d post injury. Throughout the entire time of observation neurological function was quantified using the "beam-walking" test.Glibenclamide-treated animals showed a significant reduction in the development of brain tissue water content(80.47{\%}??0.37{\%} (glibenclamide) vs. 80.83{\%}??0.44{\%} (control); p{\textless}0.05; n=14). Contusion sizes increased continuously within 72h following CCI injury, but glibenclamide-treated animals had significantly smaller volumes at any time-points, like 172.53??38.74mm3 (glibenclamide) vs. 299.20??64.02mm3 (control) (p{\textless}0.01; n=10; 24h) or 211.10??41.03mm3 (glibenclamide) vs. 309.76??19.45mm3 (control) (p{\textless}0.05; n=10; 72h), respectively. An effect on acute parameters, however, could not be detected, most likely because of the up-regulation of the channel within 3-6h after injury. Furthermore, there was no significant effect on motor function assessed by the beam-walking test throughout 7days.In accordance to these results and the available literature, glibenclamide seems to have promising potency in the treatment of TBI. ?? 2014 IBRO.},
author = {Zweckberger, K. and Hackenberg, K. and Jung, C. S. and Hertle, D. N. and Kiening, K. L. and Unterberg, A. W. and Sakowitz, O. W.},
doi = {10.1016/j.neuroscience.2014.04.040},
file = {:Users/solomon/Documents/Mendeley Desktop/Zweckberger et al. - 2014 - Glibenclamide reduces secondary brain damage after experimental traumatic brain injury.pdf:pdf},
isbn = {6221563631},
issn = {18737544},
journal = {Neuroscience},
keywords = {Brain edema,Cerebral metabolism,Epileptic seizures,Glibenclamide,Secondary brain damage,Traumatic brain injury},
pages = {199--206},
pmid = {24792709},
publisher = {IBRO},
title = {{Glibenclamide reduces secondary brain damage after experimental traumatic brain injury}},
url = {http://dx.doi.org/10.1016/j.neuroscience.2014.04.040},
volume = {272},
year = {2014}
}
@article{Sastre2015,
abstract = {BACKGROUND Genetic factors are known to influence individual differences in pain and sensitivity to analgesics. Different genetic polymorphisms in opioid-metabolizing enzymes that can affect the analgesic response to opioids have been proposed. This study investigates a possible difference in the response to postoperative buprenorphine analgesia related to the presence of different isoforms (cytosine or thymine substitution at nucleotide 802) of UGT2B7 gene. METHODS Transdermal buprenorphine was administered to 91 patients who underwent muscle-sparing thoracotomy. UGT2B7 polymorphism at locus C802T (His268Tyr) was detected using a PCR Taqman-based procedure. The severity of postoperative pain at rest and during coughing or deep inspiration was assessed by visual analog scale score after surgery. Hospital stay and perioperative opioid consumption were collected. RESULTS Genotype frequencies were 18.4{\%} for UGT2B7*1/*1, 52.9{\%} for UGT2B7*1/*2, and 28.7{\%} for UGT2B7*2/*2. VAS pain scores at rest were statistically similar among the groups except at 24, 60, and 120 hours (UGT2B7*2/*2 genotype showing higher pain scores). Patients with the UGT2B7*2/*2 genotype showed higher VAS scores triggered by coughing after the 48 hours (P {\textless}0.05). In addition, patients with this genotype reported a higher prevalence of severe pain after 48 postoperative hours (P {\textless}0.05). Thirty-eight percent of patients carrying genotype UGT2B7*2/*2 experienced severe pain in a final survey vs. 17{\%} in the group with UGT2B7*1/*1 (P = 0.36). CONCLUSIONS The presence of the SNP 802C{\textgreater}T UGT2B7 (UGT2B7*2/*2) is associated with a worse analgesic response to transdermal buprenorphine in the postoperative period of thoracic surgery.},
author = {Sastre, Jos{\'{e}} A and Varela, Gonzalo and L{\'{o}}pez, M{\'{o}}nica and Muriel, Clemente and Gonz{\'{a}}lez-Sarmiento, Rogelio},
doi = {10.1111/papr.12152},
issn = {15307085},
journal = {Pain Pract.},
keywords = {analgesia,buprenorphine,human UGT2B7,single-nucleotide polymorphisms,thoracic surgery},
month = {jan},
number = {1},
pages = {22--30},
pmid = {24256307},
title = {{Influence of Uridine Diphosphate-Glucuronyltransferase 2B7 ( UGT2B7 ) Variants on Postoperative Buprenorphine Analgesia}},
url = {http://www.ncbi.nlm.nih.gov/pubmed/24256307 http://doi.wiley.com/10.1111/papr.12152},
volume = {15},
year = {2015}
}
@article{Mei2015,
author = {Mei, Yang and Wang, Sai-Ying and Li, Yang and Yi, Shuang-Qiang and Wang, Chun-Yan and Yang, Mi and Duan, Kai-Ming},
doi = {10.1002/jcph.405},
file = {:Users/solomon/Documents/Mendeley Desktop/Mei et al. - 2015 - Role of SLCO1B1, ABCB1, and CHRNA1 gene polymorphisms on the efficacy of rocuronium in Chinese patients.pdf:pdf},
issn = {00912700},
journal = {The Journal of Clinical Pharmacology},
keywords = {abcb1,be a widespread clinical,chrna1,genetic polymorphism,nondepolarizing skeletal muscle relaxant,pharmacogenetics,rocuronium is considered to,slco1b1},
number = {3},
pages = {261--268},
title = {{Role of SLCO1B1, ABCB1, and CHRNA1 gene polymorphisms on the efficacy of rocuronium in Chinese patients}},
url = {http://doi.wiley.com/10.1002/jcph.405},
volume = {55},
year = {2015}
}
@article{Korenaga2005,
abstract = {Breast cancer resistance protein (BCRP), the second member of the ATP-binding cassette membrane transporter family, has a single nucleotide polymorphism, C421A (resulting in Q141K), that is of functional importance. Our aim was to explore the relationship between this polymorphism of the BCRP gene and the risk of renal cell carcinoma (RCC) development. For a case-control study, DNA samples from 200 nonpapillary RCC patients and 200 healthy control subjects were analyzed using the TaqMan technique. The genotypic frequencies of the BCRP C421A polymorphism were compared between RCC patients and control subjects. The frequency of the C/C genotype was significantly higher in RCC patients than in control subjects (age- and gender-adjusted OR = 1.96, 95{\%} CI 1.32-2.93). No associations were observed between the BCRP C421A polymorphism and clinicopathologic or epidemiologic factors, including age, gender, tumor grade, stage, cigarette smoking, family history of cancer and body mass index. Carriers with the C/C genotype of the BCRP C421A polymorphism are at risk of developing nonpapillary RCC. These data suggest that BCRP is a candidate RCC susceptibility gene.},
author = {Korenaga, Yoshihito and Naito, Katsusuke and Okayama, Naoko and Hirata, Hiroshi and Suehiro, Yutaka and Hamanaka, Yuichiro and Matsuyama, Hideyasu and Hinoda, Yuji},
doi = {10.1002/ijc.21187},
file = {:Users/solomon/Documents/Mendeley Desktop/Korenaga et al. - 2005 - Association of the BCRP C421A polymorphism with nonpapillary renal cell carcinoma.pdf:pdf},
isbn = {0020-7136},
issn = {00207136},
journal = {International Journal of Cancer},
keywords = {Breast cancer resistance protein,Renal cell carcinoma,Single nucleotide polymorphism},
number = {3},
pages = {431--434},
pmid = {15906349},
title = {{Association of the BCRP C421A polymorphism with nonpapillary renal cell carcinoma}},
volume = {117},
year = {2005}
}
@article{Grubs2014,
abstract = {Research using standard psychological mea-sures has not revealed a dramatic psychological reaction to receipt of predictive genetic test results. Qualitative research, however, indicates that there are subtle psycho-social sequelae of receiving such results that should be considered by individuals and their clinicians prior to testing. Among these more subtle, but burdensome seque-lae of testing are (i) changed social relationships (including relationships with clinicians), (ii) the impact of test results on important life decisions (e.g., career and reproductive choices), and (iii) altered self-concept among those tested. This review examines these more subtle sequelae, as well as the inadequacy of existing quantitative measures for assessing their impact, and discusses the relevance of such sequelae for decision-making about predictive genetic testing.},
author = {Grubs, Robin E and Parker, Lisa S and Hamilton, Rebekah},
doi = {10.1007/s40142-014-0053-7},
file = {:Users/solomon/Documents/Mendeley Desktop/Grubs, Parker, Hamilton - 2014 - Subtle Psychosocial Sequelae of Genetic Test Results.pdf:pdf},
issn = {2167-4876},
journal = {Curr Genet Med Rep},
keywords = {Ethics {\'{A}},Genetic testing {\'{A}},Huntington disease,Predictive testing {\'{A}},Psychosocial concerns {\'{A}},Qualitative research {\'{A}} BRCA {\'{A}}},
pages = {242--249},
title = {{Subtle Psychosocial Sequelae of Genetic Test Results}},
year = {2014}
}
@article{Cornelius2013,
abstract = {SIGNIFICANCE: A vast amount of circumstantial evidence implicates high energy oxidants and oxidative stress as mediators of secondary damage associated with traumatic brain injury. The excessive production of reactive oxygen species due to excitotoxicity and exhaustion of the endogenous antioxidant system induces peroxidation of cellular and vascular structures, protein oxidation, cleavage of DNA, and inhibition of the mitochondrial electron transport chain.{\$}\backslash{\$}n{\$}\backslash{\$}nRECENT ADVANCES: Different integrated responses exist in the brain to detect oxidative stress, which is controlled by several genes termed vitagens. Vitagens encode for cytoprotective heat shock proteins, and thioredoxin and sirtuins.{\$}\backslash{\$}n{\$}\backslash{\$}nCRITICAL ISSUES AND FUTURE DIRECTIONS: This article discusses selected aspects of secondary brain injury after trauma and outlines key mechanisms associated with toxicity, oxidative stress, inflammation, and necrosis. Finally, this review discusses the role of different oxidants and presents potential clinically relevant molecular targets that could be harnessed to treat secondary injury associated with brain trauma.},
author = {Cornelius, C and Crupi, R and Calabrese, V and Graziano, A and Milone, P and Pennisi, G and Radak, Z and Calabrese, EJ J and Cuzzocrea, S},
doi = {10.1089/ars.2012.4981},
file = {:Users/solomon/Documents/Mendeley Desktop/Cornelius et al. - 2013 - Traumatic brain injury oxidative stress and neuroprotection.pdf:pdf},
isbn = {1557-7716 (Electronic){\$}\backslash{\$}r1523-0864 (Linking)},
issn = {1557-7716},
journal = {Antioxid. Redox Signal.},
number = {8},
pages = {836--853},
pmid = {23547621},
title = {{Traumatic brain injury: oxidative stress and neuroprotection.}},
url = {http://www.ncbi.nlm.nih.gov/pubmed/23547621},
volume = {19},
year = {2013}
}
@article{Robertson2009,
abstract = {Levels of multidrug efflux transporter P-glycoprotein (P-gp) on endothelial cells lining brain blood vessels are important for limiting access of many compounds to the brain. In vivo studies have indicated that ischaemia-reperfusion that generates reactive oxygen species also increases P-gp levels in brain endothelial cells. To investigate possible mechanisms, in vitro studies were performed on immortalised (GPNT) and primary rat brain endothelial cells. Exposure to hydrogen peroxide (200 microM) resulted in intracellular oxidative stress as detected from higher levels of dichlorofluorescein fluorescence and raised levels of P-gp protein, mdr1a and mdr1b transcripts and, in GPNT cells, increased mdr1a and mdr1b promoter activity. The P-gp protein increases were abolished by pre-treatment with polyethylene glycol-catalase and were curtailed by co-culture with primary rat astrocytes. Exposure of GPNT cells to 6 h hypoxia followed by 24 h reoxygenation produced less intracellular oxidative stress as judged from smaller increments in dichlorofluorescein fluorescence but still resulted in raised levels of P-gp protein, an effect partially abolished by pre-treatment with polyethylene glycol-catalase. However, transcript levels and promoter activities were not significantly increased. These data suggest that hydrogen peroxide contributes to P-gp up-regulation following hypoxia-reoxygenation but the underlying mechanisms of its actions differ from those occurring after direct hydrogen peroxide application.},
author = {Robertson, Samantha J. and Kania, Katarzyna D. and Hladky, Stephen B. and Barrand, Margery A.},
doi = {10.1111/j.1471-4159.2009.06306.x},
file = {:Users/solomon/Documents/Mendeley Desktop/Robertson et al. - 2009 - P-glycoprotein expression in immortalised rat brain endothelial cells Comparisons following exogenously applie.pdf:pdf},
isbn = {1471-4159 (Electronic)$\backslash$r0022-3042 (Linking)},
issn = {00223042},
journal = {Journal of Neurochemistry},
keywords = {Astrocyte-endothelial interactions,Brain endothelial cell,Hydrogen peroxide,Hypoxia-reoxygenation,Oxidative stress,P-glycoprotein expression},
number = {1},
pages = {132--141},
pmid = {19656260},
title = {{P-glycoprotein expression in immortalised rat brain endothelial cells: Comparisons following exogenously applied hydrogen peroxide and after hypoxia-reoxygenation}},
volume = {111},
year = {2009}
}
@article{Ramsey2014a,
abstract = {Cholesterol reduction from statin therapy has been one of the greatest public health successes in modern medicine. Simvastatin is among the most commonly used prescription medications. A non-synonymous coding single-nucleotide polymorphism (SNP), rs4149056, in SLCO1B1 markedly increases systemic exposure to simvastatin and the risk of muscle toxicity. This guideline explores the relationship between rs4149056 (c.521T{\textgreater}C, p.V174A) and clinical outcome for all statins. The strength of the evidence is high for myopathy with simvastatin. We limit our recommendations accordingly.},
author = {Ramsey, L B and Johnson, S G and Caudle, K E and Haidar, C E and Voora, D and Wilke, R A and Maxwell, W D and McLeod, H L and Krauss, R M and Roden, D M and Feng, Q and Cooper-DeHoff, R M and Gong, L and Klein, T E and Wadelius, M and Niemi, M},
doi = {10.1038/clpt.2014.125},
file = {:Users/solomon/Documents/Mendeley Desktop/Ramsey et al. - 2014 - The Clinical Pharmacogenetics Implementation Consortium Guideline for SLCO1B1 and Simvastatin-Induced Myopathy(3).pdf:pdf},
issn = {0009-9236},
journal = {Clinical Pharmacology {\&} Therapeutics},
month = {oct},
number = {4},
pages = {423--428},
pmid = {22617227},
publisher = {Wiley-Blackwell},
title = {{The Clinical Pharmacogenetics Implementation Consortium Guideline for SLCO1B1 and Simvastatin-Induced Myopathy: 2014 Update}},
url = {http://www.ncbi.nlm.nih.gov/pubmed/22617227 http://www.pubmedcentral.nih.gov/articlerender.fcgi?artid=PMC3384438 http://doi.wiley.com/10.1038/clpt.2012.57 http://doi.wiley.com/10.1038/clpt.2014.125},
volume = {96},
year = {2014}
}
@article{Carrano2014,
abstract = {a b s t r a c t Alzheimer's disease (AD) is the most common form of dementia and marked by deposition of amyloid-b (Ab) within the brain. Alterations of Ab transporters at the neurovasculature may play a role in the disease process. We investigated the expression of ABC transporters P-glycoprotein (P-gp) and breast cancer related protein (BCRP) in non-neurologic controls, AD, and severe capillary cerebral amyloid angiopathy (capCAA) cases, which are characterized by deposition of Ab within cerebral capillaries. Our data show that microvascular expression of P-gp and BCRP is strikingly decreased in capCAA-affected vessels but not in AD and control samples. Messenger RNA levels of P-gp, but not of BCRP, were downregulated in brain endothelial cells on exposure to oligomeric Ab42, but not fibrillar Ab42 or Ab40. Coincubating Ab42 together with clusterin, an amyloid-associated protein highly expressed in capCAA-affected vessels, strongly reduced levels of P-gp. In conclusion, accumulation of Ab, in combination with clusterin, within and around cerebral capillaries, may further aggravate the disease process in AD by affecting P-gp expression. Loss of P-gp expression or activity may serve as a selective biomarker for ongoing capCAA.},
author = {Carrano, Anna and Snkhchyan, Hripsime and Kooij, Gijs and van der Pol, Susanne and van Horssen, Jack and Veerhuis, Robert and Hoozemans, Jeroen and Rozemuller, Annemieke and de Vries, Helga E.},
doi = {10.1016/j.neurobiolaging.2013.09.015},
file = {:Users/solomon/Documents/Mendeley Desktop/Carrano et al. - 2014 - ATP-binding cassette transporters P-glycoprotein and breast cancer related protein are reduced in capillary cere.pdf:pdf},
issn = {01974580},
journal = {Neurobiology of Aging},
keywords = {ABC transporters,Alzheimer's disease,BCRP,Blood-brain barrier,Cerebral amyloid angiopathy,P-gp},
month = {mar},
number = {3},
pages = {565--575},
title = {{ATP-binding cassette transporters P-glycoprotein and breast cancer related protein are reduced in capillary cerebral amyloid angiopathy}},
url = {https://ac-els-cdn-com.pitt.idm.oclc.org/S019745801300417X/1-s2.0-S019745801300417X-main.pdf?{\_}tid=88706664-02d7-11e8-bc1c-00000aacb360{\&}acdnat=1516998717{\_}68129319bb2455332ef0d1c8bf6b69bd http://linkinghub.elsevier.com/retrieve/pii/S019745801300417X},
volume = {35},
year = {2014}
}
@article{Barbujani2013,
abstract = {Understanding how and why humans are biologically different is indispensable to get oriented in the ever-growing body of genomic data. Here we discuss the evidence based on which we can confidently state that humans are the least genetically variable primate, both when individuals and when populations are compared, and that each individual genome can be regarded as a mosaic of fragments of different origins. Each population is somewhat different from any other population, and there are geographical patterns in that variation. These patterns clearly indicate an African origin for our species, and keep a record of the main demographic changes accompanying the peopling of the whole planet. However, only a minimal fraction of alleles, and a small fraction of combinations of alleles along the chromosome, is restricted to a single geographical region (and even less so to a single population), and diversity between members of the same population is very large. The small genomic differences between populations and the extensive allele sharing across continents explain why historical attempts to identify, once and for good, major biological groups in humans have always failed. Nevertheless, racial categorization is all but gone, especially in clinical studies. We argue that racial labels may not only obscure important differences between patients but also that they have become positively useless now that cheap and reliable methods for genotyping are making it possible to pursue the development of truly personalized medicine.},
author = {Barbujani, G. and Ghirotto, S. and Tassi, F.},
doi = {10.1111/tan.12165},
file = {:Users/solomon/Documents/Mendeley Desktop/Barbujani, Ghirotto, Tassi - 2013 - Nine things to remember about human genome diversity.pdf:pdf},
isbn = {0001-2815},
issn = {00012815},
journal = {Tissue Antigens},
keywords = {Admixture,FST,Gene flow,Genetic drift,Human origins,Human races,Pharmacogenomics,Population structure},
number = {3},
pages = {155--164},
pmid = {24032721},
title = {{Nine things to remember about human genome diversity}},
volume = {82},
year = {2013}
}
@article{Li2015,
author = {Li, Y and Jackson, KA and Slon, B and Hardy, JR and Franco, M and William, L and Poon, P and Coller, JK and Hutchinson, MR and Currow, DC and Somogyi, AA},
doi = {10.1111/bcp.12614},
file = {:Users/solomon/Documents/Mendeley Desktop/Li et al. - 2015 - CYP2B66 allele and age substantially reduce steady-state ketamine clearance in chronic pain patients impact on advers.pdf:pdf},
issn = {1365-2125 (Electronic) 0306-5251 (Linking)},
journal = {Br J Clin Pharmacol},
number = {2},
pages = {276--284},
title = {{CYP2B6*6 allele and age substantially reduce steady-state ketamine clearance in chronic pain patients: impact on adverse effects}},
type = {Journal Article},
url = {https://www.ncbi.nlm.nih.gov/pubmed/25702819},
volume = {80},
year = {2015}
}
@article{Kontoghiorghes1990,
author = {Kontoghiorghes, G J and Goddard, J G and Bartlett, A N and Sheppard, L},
file = {:Users/solomon/Documents/Mendeley Desktop/Kontoghiorghes et al. - 1990 - Pharmacokinetic studies in humans with the oral iron chelator 1,2-dimethyl-3-hydroxypyrid-4-one.pdf:pdf},
journal = {Clinincal Pharmacology},
pages = {255--261},
title = {{Pharmacokinetic studies in humans with the oral iron chelator 1,2-dimethyl-3-hydroxypyrid-4-one.}},
volume = {48},
year = {1990}
}
@article{Verhoef2013,
abstract = {BACKGROUND Observational evidence suggests that the use of a genotype-guided dosing algorithm may increase the effectiveness and safety of acenocoumarol and phenprocoumon therapy. METHODS We conducted two single-blind, randomized trials comparing a genotype-guided dosing algorithm that included clinical variables and genotyping for CYP2C9 and VKORC1 with a dosing algorithm that included only clinical variables, for the initiation of acenocoumarol or phenprocoumon treatment in patients with atrial fibrillation or venous thromboembolism. The primary outcome was the percentage of time in the target range for the international normalized ratio (INR; target range, 2.0 to 3.0) in the 12-week period after the initiation of therapy. Owing to low enrollment, the two trials were combined for analysis. The primary outcome was assessed in patients who remained in the trial for at least 10 weeks. RESULTS A total of 548 patients were enrolled (273 patients in the genotype-guided group and 275 in the control group). The follow-up was at least 10 weeks for 239 patients in the genotype-guided group and 245 in the control group. The percentage of time in the therapeutic INR range was 61.6{\%} for patients receiving genotype-guided dosing and 60.2{\%} for those receiving clinically guided dosing (P=0.52). There were no significant differences between the two groups for several secondary outcomes. The percentage of time in the therapeutic range during the first 4 weeks after the initiation of treatment in the two groups was 52.8{\%} and 47.5{\%} (P=0.02), respectively. There were no significant differences with respect to the incidence of bleeding or thromboembolic events. CONCLUSIONS Genotype-guided dosing of acenocoumarol or phenprocoumon did not improve the percentage of time in the therapeutic INR range during the 12 weeks after the initiation of therapy. (Funded by the European Commission Seventh Framework Programme and others; EU-PACT ClinicalTrials.gov numbers, NCT01119261 and NCT01119274.).},
author = {Verhoef, Talitha I and Ragia, Georgia and de Boer, Anthonius and Barallon, Rita and Kolovou, Genovefa and Kolovou, Vana and Konstantinides, Stavros and {Le Cessie}, Saskia and Maltezos, Efstratios and van der Meer, Felix J M and Redekop, William K and Remkes, Mary and Rosendaal, Frits R and van Schie, Rianne M F and Tavridou, Anna and Tziakas, Dimitrios and Wadelius, Mia and Manolopoulos, Vangelis G and {Maitland-van der Zee}, Anke H and {EU-PACT Group}},
doi = {10.1056/NEJMoa1311388},
file = {:Users/solomon/Documents/Mendeley Desktop/Verhoef et al. - 2013 - A randomized trial of genotype-guided dosing of acenocoumarol and phenprocoumon.pdf:pdf},
isbn = {0028-4793},
issn = {1533-4406},
journal = {The New England journal of medicine},
number = {24},
pages = {2304--12},
pmid = {24251360},
title = {{A randomized trial of genotype-guided dosing of acenocoumarol and phenprocoumon.}},
url = {http://www.ncbi.nlm.nih.gov/pubmed/24251360},
volume = {369},
year = {2013}
}
@article{Aiello1995,
abstract = {Brain tissue is metabolically expensive, but there is no signifi- cant correlation between relative basal metabolic rate and rela- The present paper was sumitted in final form i5 VI 94. tive brain size in humans and other encephalized mammals. The expensive-tissue hypothesis suggests that the metabolic require- ments of relatively large brains are offset by a corresponding reduction of the gut. The splanchnic organs (liver and gastro- Much of the work that has been done on encephalization intestinal tract) are as metabolically expensive as brains, and the in humans and other primates has been oriented toward gut is the only one of the metabolically expensive organs in the why questions-why different primate taxa have differ- human body that is markedly small in relation to body size. Gut size is highly correlated with diet, and relatively small guts ent relative brain sizes or why the human line has un- are compatible only with high-quality, easy-to-digest food. The dergone such a phenomenal increase in brain size during often-cited relationship between diet and relative brain size is the past 2 million years. Hypotheses that have been put more properly viewed as a relationship between relative brain forward to answer these questions primarily invoke size and relative gut size, the latter being determined by dietary quality. No matter what is selecting for relatively large brains in socio-ecological factors such as group size (Aiello and humans and other primates, they cannot be achieved without a Dunbar I993), social (or Machiavellian) intelligence shift to a high-quality diet unless there is a rise in the metabolic (Byrne and Whiten I988), or complexity of foraging strat- rate. Therefore the incorporation of increasingly greater amounts egy (Milton I979, Parker and Gibson I979, Clutton- of animal products into the diet was essential in the evolution of Brock and Harvey I980, Gibson I986, MacNab and Eisen- the large human brain},
author = {Aiello, Leslie C and Wheeler, Peter and Chivers, David},
doi = {10.1086/204350},
file = {:Users/solomon/Documents/Mendeley Desktop/Aiello, Wheeler, Chivers - 1995 - The Expensive-Tissue Hypothesis The Brain and the Digestive Evolution '.pdf:pdf},
isbn = {0011-3204},
issn = {0011-3204},
journal = {Current Anthropology},
number = {2},
pages = {199--221},
pmid = {16730368},
title = {{The Expensive-Tissue Hypothesis The Brain and the Digestive Evolution '}},
volume = {36},
year = {1995}
}
@article{Xie2008,
abstract = {Membrane transport proteins are integral membrane proteins and considered as potential drug targets. Activity assay of transport proteins is essential for developing drugs to target these proteins. Major issues related to activity assessment of transport proteins include availability of transporters, transport activity of transporters, and interactions between ligands and transporters. Researchers need to consider the physiological status of proteins (bound in lipid membranes or purified), availability and specificity of substrates, and the purpose of the activity assay (screening, identifying, or comparing substrates and inhibitors) before choosing appropriate assay strategies and techniques. Transport proteins bound in vesicular membranes can be assayed for transporting substrate across membranes by means of uptake assay or entrance counterflow assay. Alternatively, transport proteins can be assayed for interactions with ligands by using techniques such as isothermal titration calorimetry, nuclear magnetic resonance spectroscopy, or surface plasmon resonance. Other methods and techniques such as fluorometry, scintillation proximity assay, electrophysiological assay, or stopped-flow assay could also be used for activity assay of transport proteins. In this paper the major strategies and techniques for activity assessment of membrane transport proteins are reviewed.},
author = {Xie, Hao},
doi = {10.1111/j.1745-7270.2008.00400.x},
isbn = {1745-7270 (Electronic)$\backslash$r1672-9145 (Linking)},
issn = {16729145},
journal = {Acta Biochimica et Biophysica Sinica},
keywords = {Activity assay,Membrane transport protein,Protein-ligand interaction},
number = {4},
pages = {269--277},
pmid = {18401524},
title = {{Activity assay of membrane transport proteins}},
volume = {40},
year = {2008}
}
@article{Vahakangas2009,
abstract = {Studies on the increasing number of transporters found in the placental barrier are gaining momentum, because of their tissue-specific expression, significance in physiology and disease, and the possible utilization of the emerging knowledge in pharmacology. In the placenta, both syncytiotrophoblast and fetal capillary endothelium express transporters. Fetal exposure is determined by the net effect of combination of transporters, their nature and localization in relation to placental cells and their substrate specificity. Although the significance of placental transporters on human fetal drug exposure is almost an unstudied field so far, their potential use to design drugs that do not cross the placenta is already being pursued. It is thus of interest to review the existing knowledge of human placental transporters. Transporters in all groups which take part in drug transport are found in human placenta. Especially, ATP-binding cassette transporters ABCG2/breast cancer resistance protein, ABCB1/P-glycoprotein and ABCC2/MRP2 are all expressed at the apical surface of syncytiotrophoblast facing maternal blood and are putatively important protective proteins both for placental tissue and the fetus, because they are efflux transporters and their substrates include many drugs and also environmental chemicals. Such protective effect has been shown in animals, but these results cannot be directly extrapolated to humans due to interspecies differences in placental structure and function. Experimental models utilizing human placental tissue, especially human placental perfusion, offer valuable possibilities, which have been insufficiently studied so far.},
author = {V{\"{a}}h{\"{a}}kangas, Kirsi and Myllynen, P{\"{a}}ivi},
doi = {10.1111/j.1476-5381.2009.00336.x},
file = {:Users/solomon/Documents/Mendeley Desktop/V{\"{a}}h{\"{a}}kangas, Myllynen - 2009 - Drug transporters in the human blood-placental barrier.pdf:pdf},
isbn = {1476-5381},
issn = {00071188},
journal = {British Journal of Pharmacology},
keywords = {ABC transporters,Breast cancer resistance protein,Monoamine transporters,Organic anion transporters,Organic cation transporters,P-glycoprotein,Syncytiotrophoblast},
number = {3},
pages = {665--678},
pmid = {19788499},
title = {{Drug transporters in the human blood-placental barrier}},
volume = {158},
year = {2009}
}
@article{Kolias2013,
abstract = {Traumatic brain injury (TBI) remains a major public health problem. This review aims to present the principles upon which modern TBI management should be based. The early management phase aims to achieve haemodynamic stability, limit secondary insults (eg hypotension, hypoxia), obtain accurate neurological assessment and appropriately select patients for further investigation. Since 2003, the mainstay of risk stratification in the UK emergency departments has been a system of triage based on clinical assessment, which then dictates the need for a CT scan of the head. For patients with acute subdural or extradural haematomas, time from clinical deterioration to operation should be kept to a minimum, as it can affect their outcome. In addition, it is increasingly recognised that patients with severe and moderate TBI should be managed in neuroscience centres, regardless of the need for neurosurgical intervention. The monitoring and treatment of raised intracranial pressure is paramount for maintaining cerebral blood supply and oxygen delivery in patients with severe TBI. Decompressive craniectomy and therapeutic hypothermia are the subject of ongoing international multi-centre randomised trials. TBI is associated with a number of complications, some of which require specialist referral. Patients with post-concussion syndrome can be helped by supportive management in the context of a multi-disciplinary neurotrauma clinic and by patient support groups. Specialist neurorehabilitation after TBI is important for improving outcome.},
author = {Kolias, Angelos G and Guilfoyle, Mathew R and Helmy, Adel and Allanson, Judith and Hutchinson, Peter J},
doi = {10.1136/practneurol-2012-000268},
isbn = {1474-7758},
issn = {1474-7766},
journal = {Practical neurology},
keywords = {Brain Injuries,Brain Injuries: diagnosis,Brain Injuries: physiopathology,Brain Injuries: therapy,Decompressive Craniectomy,Disease Management,Humans,Intensive Care,Intracranial Hemorrhage,Tomography,Traumatic,Traumatic: etiology,X-Ray Computed},
number = {4},
pages = {228--35},
pmid = {23487823},
title = {{Traumatic brain injury in adults.}},
url = {http://www.ncbi.nlm.nih.gov/pubmed/23487823},
volume = {13},
year = {2013}
}
@article{Simard2010,
abstract = {Microvascular failure largely underlies the damaging secondary events that accompany traumatic brain injury (TBI). Changes in capillary permeability result in the extravasation of extracellular fluid, inflammatory cells, and blood, thereby producing cerebral edema, inflammation, and progressive secondary hemorrhage (PSH). Recent work in rat models of TBI and stroke have implicated 2 ion transport proteins expressed in brain endothelial cells as critical mediators of edema formation: the constitutively expressed Na(+)-K(+)-2Cl(-) cotransporter, NKCC1, and the trauma/ischemia-induced SUR1-regulated NC(Ca-ATP) (SUR1/TRPM4) channel. Whereas NKCC1 function requires adenosine 5'-triphosphate (ATP), activation of SUR1/TRPM4 occurs only after ATP depletion. This opposite dependence on intracellular ATP levels implies that one or the other mechanism will activate/deactivate as ATP concentrations rise and fall during periods of ischemia/reperfusion, resulting in continuous edema formation regardless of cellular energy status. Moreover, with critical ATP depletion, sustained opening of SUR1/TRPM4 channels results in the oncotic death of endothelial cells, leading to capillary fragmentation and PSH. Bumetanide and glibenclamide are 2 well-characterized, safe, FDA-approved drugs that inhibit NKCC1 and the SUR1/TRPM4 channel, respectively. When used alone, these drugs have provided documented beneficial effects in animal models of TBI- and ischemiaassociated cerebral edema and PSH. Given the mechanistic and temporal differences by which NKCC1 and the SUR1/TRPM4 channel contribute to the pathophysiological mechanisms of these events, combination therapy with bumetanide and glibenclamide may yield critical synergy in preventing injury-associated capillary failure.},
author = {Simard, JM and Kahle, KT and Gerzanich, V},
doi = {10.3171/2009.11.JNS081052},
file = {:Users/solomon/Documents/Mendeley Desktop/Simard, Kahle, Gerzanich - 2010 - Molecular mechanisms of microvascular failure in central nervous system injury--synergistic roles of N.pdf:pdf},
isbn = {1933-0693 (Electronic)$\backslash$r0022-3085 (Linking)},
issn = {0022-3085},
journal = {J Neurosurg},
keywords = {ATP-Binding Cassette Transporters/*metabolism,Animals,Brain Injuries/*metabolism,Brain/blood supply/metabolism,Drug/*metabolism,Humans,Inwardly Rectifying/*metabolis,Member 2,Microvessels/*metabolism,Models,Neurological,Potassium Channels,Receptors,Sodium-Potassium-Chloride Symporters/*metabolism,Solute Carrier Family 12,Sulfonylurea Receptors,TRPM Cation Channels/*metabolism},
number = {3},
pages = {622--629},
pmid = {20035575},
title = {{Molecular mechanisms of microvascular failure in central nervous system injury--synergistic roles of NKCC1 and SUR1/TRPM4}},
url = {http://www.ncbi.nlm.nih.gov/pubmed/20035575{\%}5Cnhttp://www.ncbi.nlm.nih.gov/pmc/articles/PMC3032917/pdf/nihms268500.pdf},
volume = {113},
year = {2010}
}
@article{Erny2017a,
author = {Erny, Daniel and Prinz, Marco},
doi = {10.1038/nature21910},
file = {:Users/solomon/Documents/Mendeley Desktop/Erny, Prinz - 2017 - Microbiology Gut microbes augment neurodegeneration.pdf:pdf},
issn = {0028-0836},
journal = {Nature},
month = {apr},
number = {7650},
pages = {304--305},
title = {{Microbiology: Gut microbes augment neurodegeneration}},
url = {http://www.nature.com/doifinder/10.1038/nature21910},
volume = {544},
year = {2017}
}
@article{Bowman2008,
abstract = {OBJECTIVES: The goals were to describe trends in pediatric traumatic brain injury hospitalizations in the United States and to provide national benchmarks for state and regional comparisons. METHODS: Analysis of existing data (1991-2005) from the Nationwide Inpatient Sample, the largest longitudinal, all-payer, inpatient care database in the United States, was performed. Children 0 to 19 years of age were included. Annual rates of traumatic brain injury-related hospitalizations, stratified according to age, gender, severity of traumatic brain injury, and outcome, were determined. RESULTS: From 1991 to 2005, the estimated annual incidence rate of pediatric hospitalizations associated with traumatic brain injury decreased 39{\%}, from 119.4 to 72.7 hospitalizations per 100,000. The rates decreased for all age groups and for both boys and girls, although the rate for boys remained consistently higher at each time point. Fatal hospitalization rates decreased from 3.5 deaths per 100,000 in 1991-1993 to 2.8 deaths per 100,000 in 2003-2005. The rate of mild traumatic brain injury hospitalizations accounted for most of the overall decrease, whereas nonfatal hospitalization rates for moderate and severe traumatic brain injuries remained relatively unchanged. CONCLUSIONS: Although pediatric hospitalization rates for mild traumatic brain injuries have decreased over the past 15 years, rates for moderate and severe traumatic brain injuries are relatively unchanged. Our study provides national estimates of pediatric traumatic brain injury hospitalizations that can be used as benchmarks to increase injury prevention effectiveness through targeting of effective strategies.},
author = {Bowman, Stephen M and Bird, Tommy M and Aitken, Mary E and Tilford, John M},
doi = {10.1542/peds.2007-3511},
file = {:Users/solomon/Documents/Mendeley Desktop/Bowman et al. - 2008 - Trends in hospitalizations associated with pediatric traumatic brain injuries.pdf:pdf},
isbn = {1098-4275 (Electronic) 0031-4005 (Linking)},
issn = {0031-4005},
journal = {Pediatrics},
keywords = {Accidents,Adolescent,Adult,Brain Injuries/ epidemiology/prevention {\&} control,Child,Head Protective Devices,Hospital Mortality,Hospitalization/ trends,Humans,Infant,Injury Severity Score,Preschool,Traffic/statistics {\&} numerical data,United States/epidemiology,hospitalizations,traumatic brain injury},
number = {5},
pages = {988--993},
pmid = {18977977},
title = {{Trends in hospitalizations associated with pediatric traumatic brain injuries}},
volume = {122},
year = {2008}
}
@article{Park2014,
abstract = {The sorting of an individual transmembrane (TM) segment of multi-spanning membrane proteins by the TIM23 complex in the mitochondrial inner membrane is poorly understood. Using the Mgm1 fusion approach, we attempted to assess the membrane insertion of individual TM segments of Mdl1p and Mdl2p, mitochondrial ABC transporters. Although these transporters share high sequence similarity, our results show that their membrane sorting patterns differ and that specific residues in TM domains strongly influence membrane insertion or translocation. These data imply that TIM23-mediated membrane insertion highly depends on the TM domain sequence context.},
author = {Park, Kwangjin and Jung, Sung Jun and Kim, Hyeonseong and Kim, Hyun},
doi = {10.1016/j.febslet.2014.08.001},
file = {:Users/solomon/Documents/Mendeley Desktop/Park et al. - 2014 - Mode of membrane insertion of individual transmembrane segments in Mdl1 and Mdl2, multi-spanning mitochondrial ABC.pdf:pdf},
issn = {18733468},
journal = {FEBS Letters},
keywords = {Conservative sorting,Import,Mgm1,Mitochondria,Stop-transfer,TIM23},
number = {18},
pages = {3445--3453},
pmid = {25128661},
publisher = {Federation of European Biochemical Societies},
title = {{Mode of membrane insertion of individual transmembrane segments in Mdl1 and Mdl2, multi-spanning mitochondrial ABC transporters}},
url = {http://dx.doi.org/10.1016/j.febslet.2014.08.001},
volume = {588},
year = {2014}
}
@article{Miller2010a,
abstract = {ATP-binding cassette (ABC) transporters are important selective elements of the blood-brain barrier. They line the luminal plasma membrane of the brain capillary endothelium, facing the vascular space, and both protect the central nervous system from entry of neurotoxicants and limit the access of therapeutic drugs to the brain parenchyma. Recent studies highlight the multiple signaling pathways through which the expression and activity of P-glycoprotein and other ABC transporters are modulated in response to xenobiotics, stress and disease. The results show that increased transporter expression occurs in response to signals that activate specific transcription factors, including pregnane-X receptor, constitutive androstane receptor, nuclear factor-??B and activator protein-1, and that reduced transporter activity occurs rapidly and reversibly in response to signaling through Src kinase, protein kinase C and estrogen receptors. A detailed understanding of such regulation can provide the basis for improved neuroprotection and enhanced therapeutic drug delivery to the brain. ?? 2010.},
author = {Miller, David S.},
doi = {10.1016/j.tips.2010.03.003},
file = {:Users/solomon/Documents/Mendeley Desktop/Miller - 2010 - Regulation of P-glycoprotein and other ABC drug transporters at the blood-brain barrier.pdf:pdf},
isbn = {1873-3735 (Electronic)$\backslash$r0165-6147 (Linking)},
issn = {01656147},
journal = {Trends in Pharmacological Sciences},
number = {6},
pages = {246--254},
pmid = {20417575},
publisher = {Elsevier Ltd},
title = {{Regulation of P-glycoprotein and other ABC drug transporters at the blood-brain barrier}},
url = {http://dx.doi.org/10.1016/j.tips.2010.03.003},
volume = {31},
year = {2010}
}
@article{Shah2015,
abstract = {Phenoconversion is a phenomenon that converts genotypic extensive metabolizers (EM) into phenotypic poor metabolizers (PM) of drugs, thereby modifying their clinical response to that of genotypic PMs. Phenoconversion, usually resulting from non-genetic extrinsic factors, has significant impact on analysis and interpretation of genotype-focused clinical outcome association studies and in clinical situations. The high phenotypic variability or genotype-phenotype mismatch, frequently observed within the genotypic EM population, means that the actual number of phenotypic PM subjects may be greater since many genotypic EMs would be phenotypically PMs. If the phenoconverted population with genotype-phenotype mismatch, most extensively studied for CYP2D6, is as large as the evidence suggests, there is a real risk that genotype-focused association studies, typically correlating only the genotype with clinical outcomes, may miss clinically strong pharmacogenetic associations; thus compromising any potential for advancing the prospects of personalized medicine. Phenotyping of subjects in genotype-based association studies should be routine. This review discusses potential approaches to rectify some of the current shortcomings and proposes a new nomenclature to categorise study populations. Even with strong and reliable data associating patients' genotype with clinical outcome(s), there are problems clinically in applying this knowledge into routine pharmacotherapy because of potential genotype-phenotype mismatch. Drug-induced phenoconversion during routine clinical practice remains a major public health issue. Therefore, principle challenges confronting personalized medicine, which would need to be addressed, include identifying (a) drugs that are susceptible to phenoconversion, (b) co-medications that can cause phenoconversion and (c) dosage amendments that need to be applied during and following phenoconversion.},
author = {Shah, Rashmi R. and Smith, Robert L.},
doi = {10.1111/bcp.12441},
isbn = {0306-5251},
issn = {13652125},
journal = {British Journal of Clinical Pharmacology},
keywords = {genotype-phenotype mismatch,personalized medicine,pharmacogenetic association studies,pharmacogenetics,phenoconversion,phenotyping probes},
number = {2},
pages = {222--240},
pmid = {24913012},
title = {{Addressing phenoconversion: The Achilles' heel of personalized medicine}},
volume = {79},
year = {2015}
}
@article{Consortium2010,
abstract = {Membrane transporters can be major determinants of the pharmacokinetic, safety and efficacy profiles of drugs. This presents several key questions for drug development, including which transporters are clinically important in drug absorption and disposition, and which in vitro methods are suitable for studying drug interactions with these transporters. In addition, what criteria should trigger follow-up clinical studies, and which clinical studies should be conducted if needed. In this article, we provide the recommendations of the International Transporter Consortium on these issues, and present decision trees that are intended to help guide clinical studies on the currently recognized most important drug transporter interactions. The recommendations are generally intended to support clinical development and filing of a new drug application. Overall, it is advised that the timing of transporter investigations should be driven by efficacy, safety and clinical trial enrolment questions (for example, exclusion and inclusion criteria), as well as a need for further understanding of the absorption, distribution, metabolism and excretion properties of the drug molecule, and information required for drug labelling.},
author = {Consortium, International Transporter and Giacomini, Kathleen and Huang, Shiew-Mei and Tweedie, Donald and Benet, Leslie and Brouwer, Kim and Chu, Xiaoyan and Dahlin, Amber and Evers, Raymond and Fischer, Volker and Hillgren, Kathleen and Hoffmaster, Keith and Ishikawa, Toshihisa and Keppler, Dietrich and Kim, Richard and Lee, Caroline and Niemi, Mikko and Polli, Joseph and Sugiyama, Yuichi and Swaan, Peter and Ware, Joseph and Wright, Stephen and Yee, Sook and Zamek-Gliszczynski, Maciej and Zhang, Lei},
doi = {10.1038/nrd3028},
file = {:Users/solomon/Documents/Mendeley Desktop/Consortium et al. - 2010 - Membrane transporters in drug development.pdf:pdf},
isbn = {1474-1784 (Electronic)$\backslash$n1474-1776 (Linking)},
issn = {1474-1776 VN  - readcube.com},
journal = {Nature reviews. Drug discovery},
number = {3},
pages = {215--236},
pmid = {20190787},
title = {{Membrane transporters in drug development.}},
url = {http://dx.doi.org/10.1038/nrd3028{\%}5Cn/Users/hsinfang/Box Documents/Default Sync Folder/2013 Spring rotation/Uptake Assay/Papers/Membrane Transporters in Drug Development.pdf},
volume = {9},
year = {2010}
}
@article{Winner2015,
abstract = {OBJECTIVES The objective of this project was to determine pharmacy cost savings and improvement in adherence based on a combinatorial pharmacogenomic test (CPGx ) in patients who had switched or added a new psychiatric medication after having failed monotherapy for their psychiatric disorder. RESEARCH DESIGN AND METHODS The prospective project compared 1 year pharmacy claims between a GeneSight CPGx guided cohort and a propensity-matched control group. Patients were project eligible if they augmented or switched to a different antidepressant or antipsychotic medication within the previous 90 days. Following the medication switch or augmentation, pharmacogenomic (PGx) testing was offered to each patient's treating clinician. Pharmacy claims were extracted from the Medco pharmacy claims database for each patient (n = 2168) for 1 year following testing and compared to a 5-to-1 propensity-matched treatment as usual (TAU), standard of care control group (n = 10,880). MAIN OUTCOME MEASURES Total pharmacy spend per member per year; adherence. RESULTS Patients who received PGx testing saved {\$}1035.60 in total medication costs (both CNS and non-CNS medications) over 1 year compared to the non-tested standard of care cohort (p = 0.007). PGx testing improved adherence compared to standard of care (\DeltaPDCCPGx = 0.11 vs \DeltaPDCTAU = -0.01; p {\textless} 0.0001). Pharmacy cost savings averaged {\$}2774.53 for patients who were changed to a CPGx congruent medication regimen, compared to those who were not (p {\textless} 0.0001). CONCLUSIONS PGx testing provides significant 'real world' cost savings, while simultaneously improving adherence in a difficult to treat psychiatric population. Limitations of this study include the lack of therapeutic efficacy follow-up data and possible confounding due to matching only on demographic and psychiatric variables.},
author = {Winner, Joel G. and Carhart, Joseph M. and Altar, C. Anthony and Goldfarb, Seth and Allen, Josiah D. and Lavezzari, Gabriela and Parsons, Kelly K. and Marshak, Andrew G. and Garavaglia, Susan and Dechairo, Bryan M.},
doi = {10.1185/03007995.2015.1063483},
file = {:Users/solomon/Documents/Mendeley Desktop/Winner et al. - 2015 - Combinatorial pharmacogenomic guidance for psychiatric medications reduces overall pharmacy costs in a 1 year pro.pdf:pdf},
isbn = {0300-7995},
issn = {0300-7995},
journal = {Current Medical Research and Opinion},
number = {9},
pages = {1633--1643},
pmid = {26086890},
title = {{Combinatorial pharmacogenomic guidance for psychiatric medications reduces overall pharmacy costs in a 1 year prospective evaluation}},
url = {http://www.tandfonline.com/doi/full/10.1185/03007995.2015.1063483},
volume = {31},
year = {2015}
}
@article{Cowie2017,
abstract = {Electronic health records (EHRs) provide opportunities to enhance patient care, embed performance measures in clinical practice, and facilitate clinical research. Concerns have been raised about the increasing recruitment challenges in trials, burdensome and obtrusive data collection, and uncertain generalizability of the results. Leveraging electronic health records to counterbalance these trends is an area of intense interest. The initial applications of electronic health records, as the primary data source is envisioned for observational studies, embedded pragmatic or post-marketing registry-based randomized studies, or comparative effectiveness studies. Advancing this approach to randomized clinical trials, electronic health records may potentially be used to assess study feasibility, to facilitate patient recruitment, and streamline data collection at baseline and follow-up. Ensuring data security and privacy, overcoming the challenges associated with linking diverse systems and maintaining infrastructure for repeat use of high quality data, are some of the challenges associated with using electronic health records in clinical research. Collaboration between academia, industry, regulatory bodies, policy makers, patients, and electronic health record vendors is critical for the greater use of electronic health records in clinical research. This manuscript identifies the key steps required to advance the role of electronic health records in cardiovascular clinical research.},
author = {Cowie, Martin R and Blomster, Juuso I and Curtis, Lesley H and Duclaux, Sylvie and Ford, Ian and Fritz, Fleur and Goldman, Samantha and Janmohamed, Salim and Kreuzer, J{\"{o}}rg and Leenay, Mark and Michel, Alexander and Ong, Seleen and Pell, Jill P and Southworth, Mary Ross and Stough, Wendy Gattis and Thoenes, Martin and Zannad, Faiez and Zalewski, Andrew},
doi = {10.1007/s00392-016-1025-6},
file = {:Users/solomon/Documents/Mendeley Desktop/Cowie et al. - 2017 - Electronic health records to facilitate clinical research.pdf:pdf},
issn = {1861-0684},
journal = {Clinical Research in Cardiology},
keywords = {Cardiovascular diseases,Clinical trials as topic,Electronic health records,Pragmatic clinical trials as topic},
month = {jan},
number = {1},
pages = {1--9},
pmid = {27557678},
publisher = {Springer},
title = {{Electronic health records to facilitate clinical research}},
url = {http://www.ncbi.nlm.nih.gov/pubmed/27557678 http://www.pubmedcentral.nih.gov/articlerender.fcgi?artid=PMC5226988 http://link.springer.com/10.1007/s00392-016-1025-6},
volume = {106},
year = {2017}
}
@article{Shin2013,
abstract = {Experimental investigations into the effects of traumatic brain injury (TBI) have demonstrated significant alterations in dopaminergic systems. Dopaminergic fibers originating within the substantia nigra and ventral tegmental area (VTA) are important for reward learning, addiction, movement, and behavior. However, little is known about the effect of TBI on substantia nigra and VTA function. Environmental enrichment (EE) has been shown to improve functional outcome after TBI, and a number of studies suggest that it may exert some benefits via dopaminergic signaling. To better understand the role of dopamine in chronic TBI pathophysiology and the effect of EE, we examined the mRNA expression profile within the substantia nigra and VTA at 4 weeks post-injury. Specifically, three comparisons were made: 1) TBI versus sham, 2) sham+EE versus sham+standard (STD) housing, and 3) TBI+EE versus TBI+STD. There were differential expressions of 25, 4, and 40 genes in these comparisons, respectively. Chronic alterations in genes post-injury within the substantia nigra and VTA included genes important for cellular membrane homeostasis and transcription. EE-induced gene alterations after TBI included genes important for signal transduction, in particular calcium signaling pathways, membrane homeostasis, and metabolism. Elucidation of these alterations in gene expression within the substantia nigra and VTA provides new insights into chronic changes in dopamine signaling post-TBI, and the potential role of EE in TBI rehabilitation.},
author = {Shin, S S and Bales, J W and Yan, H Q and Kline, A E and Wagner, A K and Lyons-Weiler, J and Dixon, C E},
doi = {10.1089/neu.2012.2462},
file = {:Users/solomon/Documents/Mendeley Desktop/Shin et al. - 2013 - The effect of environmental enrichment on substantia nigra gene expression after traumatic brain injury in rats.pdf:pdf},
isbn = {1557-9042 (Electronic)$\backslash$r0897-7151 (Linking)},
issn = {0897-7151},
journal = {J Neurotrauma},
keywords = {*Transcriptome,Animals,Blotting, Western,Brain Injuries/*genetics/physiopathology/rehabilit,Calcium Signaling/physiology,Disease Models, Animal,Dopamine/biosynthesis/genetics,Environment,Housing, Animal,In Situ Hybridization,Male,Motor Activity/physiology,Oligonucleotide Array Sequence Analysis,RNA, Messenger/analysis,Rats,Rats, Sprague-Dawley,Real-Time Polymerase Chain Reaction,Reverse Transcriptase Polymerase Chain Reaction,Substantia Nigra/*physiopathology},
number = {4},
pages = {259--270},
pmid = {23094804},
title = {{The effect of environmental enrichment on substantia nigra gene expression after traumatic brain injury in rats}},
url = {http://www.ncbi.nlm.nih.gov/pubmed/23094804},
volume = {30},
year = {2013}
}
@article{Ebell2004,
abstract = {A large number of taxonomies are used to rate the quality of an individual study and the strength of a recommendation based on a body of evidence. We have developed a new grading scale that will be used by several family medicine and primary care journals (required or optional), with the goal of allowing readers to learn one taxonomy that will apply to many sources of evidence. Our scale is called the Strength of Recommendation Taxonomy. It addresses the quality, quantity, and consistency of evidence and allows authors to rate individual studies or bodies of evidence. The taxonomy is built around the information mastery framework, which emphasizes the use of patient-oriented outcomes that measure changes in morbidity or mortality. An A-level recommendation is based on consistent and good-quality patient-oriented evidence; a B-level recommendation is based on inconsistent or limited-quality patient-oriented evidence; and a C-level recommendation is based on consensus, usual practice, opinion, disease-oriented evidence, or case series for studies of diagnosis, treatment, prevention, or screening. Levels of evidence from 1 to 3 for individual studies also are defined. We hope that consistent use of this taxonomy will improve the ability of authors and readers to communicate about the translation of research into practice.},
author = {Ebell, Mark H. and Siwek, Jay and Weiss, Barry D. and Woolf, Steven H. and Susman, Jeffrey and Ewigman, Bernard and Bowman, Marjorie},
doi = {10.3122/jabfm.17.1.59},
file = {:Users/solomon/Documents/Mendeley Desktop/Ebell et al. - 2004 - Strength of Recommendation Taxonomy (SORT) A Patient-Centered Approach to Grading Evidence in the Medical Literatu.pdf:pdf},
isbn = {10.3122/jabfm.17.1.59},
issn = {0002838X},
journal = {American Family Physician},
number = {3},
pages = {548--556},
pmid = {14971837},
title = {{Strength of Recommendation Taxonomy (SORT): A Patient-Centered Approach to Grading Evidence in the Medical Literature}},
volume = {69},
year = {2004}
}
@article{Higashikuni2010,
abstract = {To clarify the impact of breast cancer resistance protein 1 (BCRP1)/ATP-binding cassette transporter subfamily G member 2 (ABCG2) expression on cardiac repair after myocardial infarction (MI).},
author = {Higashikuni, Yasutomi and Sainz, Julie and Nakamura, Kazuto and Takaoka, Minoru and Enomoto, Soichiro and Iwata, Hiroshi and Sahara, Makoto and Tanaka, Kimie and Koibuchi, Nobutaka and Ito, Sumito and Kusuhara, Hiroyuki and Sugiyama, Yuichi and Hirata, Yasunobu and Nagai, Ryozo and Sata, Masataka},
doi = {10.1161/ATVBAHA.110.211755},
file = {:Users/solomon/Documents/Mendeley Desktop/Higashikuni et al. - 2010 - The ATP-binding cassette transporter BCRP1ABCG2 plays a pivotal role in cardiac repair after myocardial infa.pdf:pdf},
issn = {10795642},
journal = {Arteriosclerosis, Thrombosis, and Vascular Biology},
keywords = {BCRP1/ABCG2,angiogenesis,myocardial infarction,pathology,remodeling},
number = {11},
pages = {2128--2135},
pmid = {20829509},
title = {{The ATP-binding cassette transporter BCRP1/ABCG2 plays a pivotal role in cardiac repair after myocardial infarction via modulation of microvascular endothelial cell survival and function}},
volume = {30},
year = {2010}
}
@article{Natale2003,
abstract = {Brain trauma is a major cause of morbidity and mortality, both in adult and pediatric populations. Much of the functional deficit derives from delayed cell death resulting from induction of neurotoxic factors that overwhelm endogenous neuroprotective responses. To identify the potential molecular mechanisms underlying such delayed responses, we compared gene expression patterns using high-density oligonucleotide arrays at 4, 8, 24, and 72 h after moderate levels of lateral fluid percussion-induced brain injury in rats and lateral controlled cortical impact injury in mice (a total of 47 profiles). Expression of 82 genes in 12 functional categories was significantly changed in both species after trauma. The largest number of gene expression changes were found in the functional groups related to inflammation (17{\%}), transcription regulation (16{\%}), and cell adhesion/extracellular matrix (15{\%}). Fifty percent of genes similarly altered across models had not been previously implicated in traumatic brain injury. Of particular interest were expression changes in genes linked to neurodegeneration, such as ATF3 and lysosomal membrane glycoprotein 2, and to neuroprotection including lipocortin 1, calponin 3, gelsolin, Id-1, and p45 NF-E2. Gene expression profiling across species and models may help identify candidate molecular pathways induced by brain injury, some of which may provide novel targets for therapeutic intervention.},
author = {Natale, Joanne E and Ahmed, Farid and Cernak, Ibolja and Stoica, Bogdan and Faden, Alan I},
doi = {10.1089/089771503770195777},
file = {:Users/solomon/Documents/Mendeley Desktop/Natale et al. - 2003 - Gene expression profile changes are commonly modulated across models and species after traumatic brain injury.pdf:pdf},
isbn = {0897-7151 (Print)$\backslash$r0897-7151 (Linking)},
issn = {0897-7151},
journal = {Journal of neurotrauma},
keywords = {Animal,Animals,Brain Injuries,Brain Injuries/drug therapy/ genetics/metabolism,Brain Injuries: drug therapy,Brain Injuries: genetics,Brain Injuries: metabolism,Disease Models,Gene Expression Profiling,Gene Expression Profiling/ methods,Gene Expression Profiling: methods,Inbred C57BL,Male,Mice,Oligonucleotide Array Sequence Analysis,Oligonucleotide Array Sequence Analysis/ methods,Oligonucleotide Array Sequence Analysis: methods,Rats,Species Specificity,Sprague-Dawley},
number = {10},
pages = {907--927},
pmid = {14588109},
title = {{Gene expression profile changes are commonly modulated across models and species after traumatic brain injury.}},
url = {http://www.ncbi.nlm.nih.gov/pubmed/14588109},
volume = {20},
year = {2003}
}
@article{Bratton2007,
author = {Bratton, SL and Chestnut, RM and Ghajar, J and {McConnell Hammond}, FF and Harris, OA and Hartl, R and Manley, GT and Nemecek, A and Newell, DW and Rosenthal, G and Schouten, J and Shutter, L and Timmons, SD and Ullman, JS and Videtta, W and Wilberger, JE and Wright, DW},
doi = {10.1089/neu.2007.9985},
file = {:Users/solomon/Documents/Mendeley Desktop/Bratton et al. - 2007 - XI. Anesthetics, Analgesics, and Sedatives.pdf:pdf},
issn = {0897-7151},
journal = {Journal of Neurotrauma},
month = {may},
number = {supplement 1},
pages = {S--71--S--76},
title = {{XI. Anesthetics, Analgesics, and Sedatives}},
url = {http://www.liebertonline.com/doi/abs/10.1089/neu.2007.9985},
volume = {24},
year = {2007}
}
@article{Rosenblat2017,
author = {Rosenblat, Joshua D. and Lee, Yena and McIntyre, Roger S.},
doi = {10.4088/JCP.15r10583},
issn = {0160-6689},
journal = {The Journal of Clinical Psychiatry},
month = {jun},
number = {06},
pages = {720--729},
title = {{Does Pharmacogenomic Testing Improve Clinical Outcomes for Major Depressive Disorder?}},
url = {http://www.psychiatrist.com/jcp/article/pages/2017/v78n06/v78n0614.aspx},
volume = {78},
year = {2017}
}
@article{Wagner2014,
author = {Wagner, AK K and Sowa, G},
doi = {10.1097/PHM.0000000000000114},
file = {:Users/solomon/Documents/Mendeley Desktop/Wagner, Sowa - 2014 - Rehabilomics research a model for translational rehabilitation and comparative effectiveness rehabilitation resear.pdf:pdf},
isbn = {0000000000000},
issn = {1537-7385},
journal = {Am. J. Phys. Med. Rehabil.},
keywords = {biomarkers,comparative effectiveness,evidence-based medicine,personalized medicine,rehabilitation,rehabilomics research,research},
month = {oct},
number = {10},
pages = {913--916},
pmid = {24901762},
title = {{Rehabilomics research: a model for translational rehabilitation and comparative effectiveness rehabilitation research.}},
url = {http://content.wkhealth.com/linkback/openurl?sid=WKPTLP:landingpage{\&}an=00002060-201410000-00010 http://www.ncbi.nlm.nih.gov/pubmed/24901762 http://content.wkhealth.com/linkback/openurl?sid=WKPTLP:landingpage{\%}7B{\&}{\%}7Dan=00002060-201410000-00010},
volume = {93},
year = {2014}
}
@article{Caudle2014,
author = {Caudle, KE and Rettie, AE and Whirl-Carrillo, M and Smith, LH and Mintzer, S and Lee, MT and Klein, TE and Callaghan, JT},
doi = {10.1038/clpt.2014.159},
file = {:Users/solomon/Documents/Mendeley Desktop/Caudle et al. - 2014 - Clinical pharmacogenetics implementation consortium guidelines for CYP2C9 and HLA-B genotypes and phenytoin dosin.pdf:pdf},
issn = {1532-6535 (Electronic) 0009-9236 (Linking)},
journal = {Clin Pharmacol Ther},
number = {5},
pages = {542--548},
title = {{Clinical pharmacogenetics implementation consortium guidelines for CYP2C9 and HLA-B genotypes and phenytoin dosing}},
type = {Journal Article},
url = {https://www.ncbi.nlm.nih.gov/pubmed/25099164},
volume = {96},
year = {2014}
}
@article{Kapoor2013,
author = {Kapoor, Shailendra},
doi = {10.1007/s10072-013-1441-6},
file = {:Users/solomon/Documents/Mendeley Desktop/Kapoor - 2013 - Deferoxamine Emerging, new neuro-protective benefits.pdf:pdf},
isbn = {1007201314416},
issn = {15901874},
journal = {Neurological Sciences},
number = {11},
pages = {2069--2070},
title = {{Deferoxamine: Emerging, new neuro-protective benefits}},
volume = {34},
year = {2013}
}
@article{DeFerra1985,
abstract = {We have isolated cDNA clones encoding the four different forms of mouse myelin basic protein (MBP) and have analyzed the structure of the MBP gene. The three larger forms of MBP differ from the smallest by the inclusion of either or both of two short amino acid sequences at positions 57 and 124 of the smallest protein. The mouse genome contains a single MBP gene comprised of seven exons. The two amino acid sequences present only in the larger MBPs are encoded by separate exons. Furthermore, all exons in the coding region begin or end in complete codons so that alternative splicing does not alter the reading frame. We conclude that the four forms of this myelin protein are encoded in separate mRNAs, each derived by a simple alternative splicing of the primary MBP gene transcript. Comparison of the amino acid sequence encoded by each exon with a recent model of the secondary structure of MBP suggests that each of the seven exons encodes one or two of the predicted structural motifs of the protein. ?? 1985.},
author = {de Ferra, Francesca and Engh, Helen and Hudson, Lynn and Kamholz, John and Puckett, Carmie and Molineaux, Susan and Lazzarini, Robert A.},
doi = {10.1016/0092-8674(85)90245-4},
file = {:Users/solomon/Documents/Mendeley Desktop/de Ferra et al. - 1985 - Alternative splicing accounts for the four forms of myelin basic protein.pdf:pdf},
isbn = {0092-8674 (Print)$\backslash$r0092-8674 (Linking)},
issn = {00928674},
journal = {Cell},
number = {3 PART 2},
pages = {721--727},
pmid = {2416470},
title = {{Alternative splicing accounts for the four forms of myelin basic protein}},
volume = {43},
year = {1985}
}
@article{Eng2000,
abstract = {It is now well established that the glial fibrillary acidic protein (GFAP) is the principal 8-9 nm intermediate filament in mature astrocytes of the central nervous system (CNS). Over a decade ago, the value of GFAP as a prototype antigen in nervous tissue identification and as a standard marker for fundamental and applied research at an interdisciplinary level was recognized (Raine, 135). As a member of the cytoskeletal protein family, GFAP is thought to be important in modulating astrocyte motility and shape by providing structural stability to astrocytic processes. In the CNS of higher vertebrates, following injury, either as a result of trauma, disease, genetic disorders, or chemical insult, astrocytes become reactive and respond in a typical manner, termed astrogliosis. Astrogliosis is characterized by rapid synthesis of GFAP and is demonstrated by increase in protein content or by immunostaining with GFAP antibody. In addition to the major application of GFAP antisera for routine use in astrocyte identification in the CNS, the molecular cloning of the mouse gene in 1985 has opened a new and rich realm for GFAP studies. These include antisense, null mice, and numerous promoter studies. Studies showing that mice lacking GFAP are hypersensitive to cervical spinal cord injury caused by sudden acceleration of the head have provided more direct evidence for a structural role of GFAP. While the structural function of GFAP has become more acceptable, the use of GFAP antibodies and promoters continue to be valuable in studying CNS injury, disease, and development.},
author = {Eng, Lawrence F and Ghirnikar, Roopa S and Lee, Yuen L},
doi = {10.1023/A:1007677003387},
file = {:Users/solomon/Documents/Mendeley Desktop/Eng, Ghirnikar, Lee - 2000 - Glial Fibrillary Acidic Protein GFAP-Thirty-One Years (1969-2000).pdf:pdf},
isbn = {0364-3190 (Print)$\backslash$n0364-3190 (Linking)},
issn = {0364-3190},
journal = {Neurochemical Research},
keywords = {astrogliosis,cns intermediate filaments,gfap},
number = {9-10},
pages = {1439--1451},
pmid = {11059815},
title = {{Glial Fibrillary Acidic Protein : GFAP-Thirty-One Years (1969-2000)}},
volume = {25},
year = {2000}
}
@article{Ayton2014,
abstract = {Traumatic brain injury (TBI) is in part complicated by pro-oxidant iron elevation independent of brain hemorrhage. Ceruloplasmin (CP) and ??-amyloid protein precursor (APP) are known neuroprotective proteins that reduce oxidative damage through iron regulation. We surveyed iron, CP, and APP in brain tissue from control and TBI-affected patients who were stratified according to time of death following injury. We observed CP and APP induction after TBI accompanying iron accumulation. Elevated APP and CP expression was also observed in a mouse model of focal cortical contusion injury concomitant with iron elevation. To determine if changes in APP or CP were neuroprotective we employed the same TBI model on APP-/- and CP-/- mice and found that both exhibited exaggerated infarct volume and iron accumulation postinjury. Evidence supports a regulatory role of both proteins in defence against iron-induced oxidative damage after TBI, which presents as a tractable therapeutic target. ?? 2014 Elsevier Inc.},
author = {Ayton, Scott and Zhang, Moses and Roberts, Blaine R. and Lam, Linh Q. and Lind, Monica and McLean, Catriona and Bush, Ashley I. and Frugier, Tony and Crack, Peter J. and Duce, James A.},
doi = {10.1016/j.freeradbiomed.2014.01.041},
file = {:Users/solomon/Documents/Mendeley Desktop/Ayton et al. - 2014 - Ceruloplasmin and -amyloid precursor protein confer neuroprotection in traumatic brain injury and lower neuronal i.pdf:pdf},
isbn = {1873-4596 (Electronic)$\backslash$r0891-5849 (Linking)},
issn = {08915849},
journal = {Free Radical Biology and Medicine},
keywords = {Amyloid precursor protein,Ceruloplasmin,Free radicals,Iron,Mouse models,Traumatic brain injury},
pages = {331--337},
pmid = {24509156},
publisher = {Elsevier},
title = {{Ceruloplasmin and ??-amyloid precursor protein confer neuroprotection in traumatic brain injury and lower neuronal iron}},
url = {http://dx.doi.org/10.1016/j.freeradbiomed.2014.01.041},
volume = {69},
year = {2014}
}
@article{Babikian2011,
abstract = {Traumatic brain injury (TBI), one of the most frequent causes of neurologic and neurobehavioral morbidity in the pediatric population, can result in lifelong challenges not only for patients, but also for their families. Survivors of a brain injury experienced during childhood—when the brain is undergoing a period of rapid development—frequently experience unique challenges as the consequences of their injuries are overlaid on normal developmental changes. Experimental studies have significantly advanced our understanding of the mechanisms and underlying molecular underpinnings of the injury response and recovery process following a TBI in the developing brain. In this paper, normal and TBI-related alterations in growth, development and metabolism are comprehensively reviewed in the postweanling/juvenile age range in the rat (postnatal days 21–60). As part of this review, TBI-related changes in gene expression are presented, with a focus on the injury-induced alterations related to cerebral growth and metabolism, and discussed in the context of existing literature related to physiological and behavioral responses to experimental TBI. Increasing evidence from the existing literature and from our own gene microarray data indicates that molecular responses related to growth, development and metabolism may play a particularly important role in the injury response and the recovery trajectory following developmental TBI. While gene expression analysis shows many of these changes occur at the level of transcription, a comprehensive review of other studies suggests that the control of metabolic substrates may preferentially be regulated through changes in transporters and enzymatic activity. The interrelation between cellular metabolism and activity-dependent neuroplasticity shows great promise as an area for future study for an optimal translation of experimental data to clinical TBI, with the ultimate goal of guiding therapeutic interventions. (PsycINFO Database Record (c) 2012 APA, all rights reserved). (journal abstract)},
author = {Babikian, Talin and Prins, Mayumi L. and Cai, Yan and Barkhoudarian, Garni and Hartonian, Ivet and Hovda, David A. and Giza, Christopher C.},
doi = {10.1159/000320667},
file = {:Users/solomon/Documents/Mendeley Desktop/Babikian et al. - 2011 - Molecular and physiological responses to juvenile traumatic brain injury Focus on growth and metabolism.pdf:pdf},
isbn = {0378-5866},
issn = {03785866},
journal = {Developmental Neuroscience},
keywords = {Gene expression,Growth,Growth factor,Metabolism,Oxidative stress,Traumatic brain injury, experimental,Traumatic brain injury, juvenile},
number = {5-6},
pages = {431--441},
pmid = {21071915},
title = {{Molecular and physiological responses to juvenile traumatic brain injury: Focus on growth and metabolism}},
volume = {32},
year = {2011}
}
@article{Anderson1999,
abstract = {Nucleoside transport processes may play a role in regulating endogenous levels of the inhibitory neuromodulator adenosine in brain. The cDNAs encoding species homologues of one member of the equilibrative nucleoside transporter (ENT) gene family have recently been isolated from rat (rENT1) and human (hENT1) tissues. The current study used RT-PCR, northern blot, in situ hybridization, and [3H]nitrobenzylthioinosine autoradiography to determine the distribution of mRNA and protein for ENT1 in rat and human brain. Northern blot analysis indicated that hENT1 mRNA is widely distributed in adult human brain. 35S-labeled sense and antisense riboprobes, transcribed from a 153-bp segment of rENT1, were hybridized to fresh frozen coronal sections from adult rat brain and revealed widespread rENT1 mRNA in pyramidal neurons of the hippocampus, granule neurons of the dentate gyrus, Purkinje and granule neurons of the cerebellum, and cortical and striatal neurons. Regional localization in rat brain was confirmed by RT-PCR. Thus, ENT1 mRNA has a wide cellular and regional distribution in brain, indicating that this nucleoside transporter subtype may be important in regulating intra- and extracellular levels of adenosine in brain.},
author = {Anderson, Christopher M. and Xiong, Wei and Geiger, Jonathan D. and Young, James D. and Cass, Carol E. and Baldwin, Stephen A. and Parkinson, Fiona E.},
doi = {10.1046/j.1471-4159.1999.0730867.x},
file = {:Users/solomon/Documents/Mendeley Desktop/Anderson et al. - 1999 - Distribution of equilibrative, nitrobenzylthioinosine-sensitive nucleoside transporters (ENT1) in brain.pdf:pdf},
isbn = {0022-3042 (Print)$\backslash$r0022-3042 (Linking)},
issn = {00223042},
journal = {Journal of Neurochemistry},
keywords = {Adenosine,Adenosine transport,ENT1,Es nucleoside transport,In situ hybridization,Nitrobenzylthioinosine,Nucleoside transport,RT-PCR},
number = {2},
pages = {867--873},
pmid = {10428086},
title = {{Distribution of equilibrative, nitrobenzylthioinosine-sensitive nucleoside transporters (ENT1) in brain}},
volume = {73},
year = {1999}
}
@article{Thomas2009,
abstract = {Mitochondrial dysfunction and reactive oxygen species (ROS) are often implicated in diseases involving oxidative stress and elevated iron. As mitochondria produce ATP by oxidative phosphorylation, ROS by-products are generated from the electron transport chain. Although superoxide and hydrogen peroxide have been thoroughly investigated, little evidence documents hydroxyl radical (HO(*)) production in mitochondria. In order to determine whether HO(*) is generated under oxidative stress conditions by a Fenton-type mechanism, bovine heart submitochondrial particles were examined for HO(*) in the presence and absence of iron ligands, antioxidant enzymes and HO(*) scavengers. HO(*) was measured as 2,3- and 2,5-dihydroxybenzoic acid (DHBA), using HPLC with electrochemical detection. The iron ligand desferrioxamine significantly decreased DHBAs, indicating that HO(*) generation required iron redox-cycling. In addition, results from exogenous SOD and catalase, exogenous hydrogen peroxide, and HO(*)-scavenger studies support a Fenton-type reaction mechanism. The results indicate that increased HO(*) levels occur in mitochondria under oxidative stress and that the HO(*) levels can be modulated with antioxidant enzymes and iron ligands. Our findings together with reports on iron accumulation in degenerative diseases highlight the importance of developing mitochondrial-targeted antioxidants for the therapeutic intervention of diseases associated with mitochondrial dysfunction and oxidative stress.},
author = {Thomas, Carin and Mackey, Melissa M and Diaz, Amy A and Cox, David P},
doi = {10.1179/135100009X392566},
file = {:Users/solomon/Documents/Mendeley Desktop/Thomas et al. - 2009 - Hydroxyl radical is produced via the Fenton reaction in submitochondrial particles under oxidative stress Implica.pdf:pdf},
isbn = {5099632815},
issn = {1743-2928},
journal = {Redox report : communications in free radical research},
keywords = {Animals,Catechols,Catechols: chemistry,Cattle,Chromatography,Heart,Heart: metabolism,High Pressure Liquid,Hydroxybenzoates,Hydroxyl Radical,Hydroxyl Radical: chemistry,Hydroxyl Radical: metabolism,Mitochondria,Oxidative Stress,Oxidative Stress: physiology,Reactive Oxygen Species,Reactive Oxygen Species: metabolism,Submitochondrial Particles,Submitochondrial Particles: metabolism},
number = {3},
pages = {102--8},
pmid = {19490751},
title = {{Hydroxyl radical is produced via the Fenton reaction in submitochondrial particles under oxidative stress: Implications for diseases associated with iron accumulation}},
url = {http://www.ncbi.nlm.nih.gov/pubmed/19490751},
volume = {14},
year = {2009}
}
@article{Yee2010,
author = {Yee, SW and Chen, L and Giacomini, KM},
doi = {10.2217/pgs.10.22},
file = {:Users/solomon/Documents/Mendeley Desktop/Yee, Chen, Giacomini - 2010 - Pharmacogenomics of membrane transporters past, present and future.pdf:pdf},
issn = {1462-2416},
journal = {Pharmacogenomics},
keywords = {coding variant,genome-wide association study,membrane,noncoding variant,pharmacogenomics,polymorphism,transporter},
month = {apr},
number = {4},
pages = {475--479},
pmid = {20350125},
title = {{Pharmacogenomics of membrane transporters: past, present and future}},
url = {http://www.futuremedicine.com/doi/10.2217/pgs.10.22},
volume = {11},
year = {2010}
}
@book{Tobergte2013a,
abstract = {applicability for this approach.},
archivePrefix = {arXiv},
arxivId = {arXiv:1011.1669v3},
author = {Tobergte, David R. and Curtis, Shirley},
booktitle = {Journal of Chemical Information and Modeling},
doi = {10.1017/CBO9781107415324.004},
eprint = {arXiv:1011.1669v3},
file = {:Users/solomon/Documents/Mendeley Desktop/Tobergte, Curtis - 2013 - Statistics for biology and health.pdf:pdf},
isbn = {9788578110796},
issn = {1098-6596},
keywords = {icle},
number = {9},
pages = {1689--1699},
pmid = {25246403},
title = {{Statistics for biology and health}},
volume = {53},
year = {2013}
}
@article{Gislefoss2008,
author = {Gislefoss, Randi E. and Grimsrud, Tom K. and M{\o}rkrid, Lars},
doi = {10.1080/00365510701809235},
issn = {0036-5513},
journal = {Scandinavian Journal of Clinical and Laboratory Investigation},
month = {jan},
number = {5},
pages = {402--409},
title = {{Long‐term stability of serum components in the Janus Serum Bank}},
url = {http://www.tandfonline.com/doi/full/10.1080/00365510701809235},
volume = {68},
year = {2008}
}
@article{Chiabrando2014,
abstract = {Heme (iron-protoporphyrin IX) is an essential co-factor involved in multiple biological processes: oxygen transport and storage, electron transfer, drug and steroid metabolism, signal transduction, and micro RNA processing. However, excess free-heme is highly toxic due to its ability to promote oxidative stress and lipid peroxidation, thus leading to membrane injury and, ultimately, apoptosis. Thus, heme metabolism needs to be finely regulated. Intracellular heme amount is controlled at multiple levels: synthesis, utilization by hemoproteins, degradation and both intracellular and intercellular trafficking. This review focuses on recent findings highlighting the importance of controlling intracellular heme levels to counteract heme-induced oxidative stress. The contributions of heme scavenging from the extracellular environment, heme synthesis and incorporation into hemoproteins, heme catabolism and heme transport in maintaining adequate intracellular heme content are discussed. Particular attention is put on the recently described mechanisms of heme trafficking through the plasma membrane mediated by specific heme importers and exporters. Finally, the involvement of genes orchestrating heme metabolism in several pathological conditions is illustrated and new therapeutic approaches aimed at controlling heme metabolism are discussed.},
author = {Chiabrando, Deborah and Vinchi, Francesca and Fiorito, Veronica and Mercurio, Sonia and Tolosano, Emanuela},
doi = {10.3389/fphar.2014.00061},
file = {:Users/solomon/Documents/Mendeley Desktop/Chiabrando et al. - 2014 - Heme in pathophysiology A matter of scavenging, metabolism and trafficking across cell membranes.pdf:pdf},
isbn = {1663-9812},
issn = {16639812},
journal = {Frontiers in Pharmacology},
keywords = {ABCG2,FLVCR1,FLVCR2,HCP1/PCFT,HO-1,Hemopexin},
number = {April},
pages = {1--24},
pmid = {24782769},
title = {{Heme in pathophysiology: A matter of scavenging, metabolism and trafficking across cell membranes}},
volume = {5},
year = {2014}
}
@article{Cordell2002,
author = {Cordell, H. J.},
doi = {10.1093/hmg/11.20.2463},
file = {:Users/solomon/Documents/Mendeley Desktop/Cordell - 2002 - Epistasis what it means, what it doesn't mean, and statistical methods to detect it in humans.pdf:pdf},
issn = {14602083},
journal = {Human Molecular Genetics},
month = {oct},
number = {20},
pages = {2463--2468},
publisher = {Oxford University Press},
title = {{Epistasis: what it means, what it doesn't mean, and statistical methods to detect it in humans}},
url = {https://academic.oup.com/hmg/article-lookup/doi/10.1093/hmg/11.20.2463},
volume = {11},
year = {2002}
}
@article{Bushnell2014,
abstract = {PURPOSE: The aim of this statement is to summarize data on stroke risk factors that are unique to and more common in women than men and to expand on the data provided in prior stroke guidelines and cardiovascular prevention guidelines for women. This guideline focuses on the risk factors unique to women, such as reproductive factors, and those that are more common in women, including migraine with aura, obesity, metabolic syndrome, and atrial fibrillation. METHODS: Writing group members were nominated by the committee chair on the basis of their previous work in relevant topic areas and were approved by the American Heart Association (AHA) Stroke Council's Scientific Statement Oversight Committee and the AHA's Manuscript Oversight Committee. The panel reviewed relevant articles on adults using computerized searches of the medical literature through May 15, 2013. The evidence is organized within the context of the AHA framework and is classified according to the joint AHA/American College of Cardiology and supplementary AHA Stroke Council methods of classifying the level of certainty and the class and level of evidence. The document underwent extensive AHA internal peer review, Stroke Council Leadership review, and Scientific Statements Oversight Committee review before consideration and approval by the AHA Science Advisory and Coordinating Committee. RESULTS: We provide current evidence, research gaps, and recommendations on risk of stroke related to preeclampsia, oral contraceptives, menopause, and hormone replacement, as well as those risk factors more common in women, such as obesity/metabolic syndrome, atrial fibrillation, and migraine with aura. CONCLUSIONS: To more accurately reflect the risk of stroke in women across the lifespan, as well as the clear gaps in current risk scores, we believe a female-specific stroke risk score is warranted.},
author = {Bushnell, Cheryl and McCullough, Louise D. and Awad, Issam A. and Chireau, Monique V. and Fedder, Wende N. and Furie, Karen L. and Howard, Virginia J. and Lichtman, Judith H. and Lisabeth, Lynda D. and Pi??a, Ileana L. and Reeves, Mathew J. and Rexrode, Kathryn M. and Saposnik, Gustavo and Singh, Vineeta and Towfighi, Amytis and Vaccarino, Viola and Walters, Matthew R.},
doi = {10.1161/01.str.0000442009.06663.48},
isbn = {0039-2499, 1524-4628},
issn = {15244628},
journal = {Stroke},
keywords = {AHA Scientific Statements,Atrial fibrillation,Hormone replacement therapy,Menopause,Metabolic syndrome X,Preeclampsia/eclampsia,Sex differences,Stroke},
number = {5},
pages = {1545--1588},
pmid = {24503673},
title = {{Guidelines for the prevention of stroke in women: A statement for healthcare professionals from the American heart association/American stroke association}},
volume = {45},
year = {2014}
}
@article{Beschorner2007,
abstract = {UNLABELLED:Glutamate-mediated excitotoxicity is known to cause secondary brain damage following stroke and traumatic brain injury (TBI). However, clinical trials using NMDA antagonists failed. Thus, glial excitatory amino acid transporters (EAATs) might be a promising target for therapeutic intervention.$\backslash$n$\backslash$nMETHODS AND RESULTS:We examined expression of EAAT1 (GLAST) and EAAT2 (Glt-1) in 36 TBI cases by immunohistochemistry. Cortical expression of both EAATs decreased rapidly and widespread throughout the brain (in lesional, adjacent and remote areas) following TBI. In the white matter numbers of EAAT1+ parenchymal cells increased 39-fold within 24h (p{\textless}0.001) and remained markedly elevated till later stages in the lesion (90-fold, p{\textless}0.01) and in peri-lesional regions (86-fold, p{\textless}0.01). In contrast, EAAT2+ parenchymal cells and EAAT1+ or EAAT2+ perivascular cells did not increase significantly. Within the first days following TBI mainly activated microglia and thereafter mainly reactive astrocytes expressed EAAT1. Perivascular monocytes and foamy macrophages lacked EAAT1 immunoreactivity. We conclude that following TBI i) loss of cortical EAATs contributes to secondary brain damage, ii) glial EAAT1 expression reflects a potential neuroprotective function of microglia and astrocytes, iii) microglial EAAT1 expression is restricted to an early stage of activation, iv) blood-derived monocytes do not express EAAT1 and v) pharmacological modification of glial EAAT expression might further limit neuronal damage.},
author = {Beschorner, Rudi and Dietz, K. and Schauer, N. and Mittelbronn, M. and Schluesener, H. J. and Trautmann, K. and Meyermann, R. and Simon, P.},
doi = {10.14670/HH-22.515},
file = {:Users/solomon/Documents/Mendeley Desktop/Beschorner et al. - 2007 - Expression of EAAT1 reflects a possible neuroprotective function of reactive astrocytes and activated microgl.pdf:pdf},
isbn = {1699-5848 (Electronic)$\backslash$n0213-3911 (Linking)},
issn = {02133911},
journal = {Histology and Histopathology},
keywords = {Excitatory amino acid transporters,Microglial activation,Neuroprotection,Traumatic brain injury},
number = {4-6},
pages = {515--526},
pmid = {17330806},
title = {{Expression of EAAT1 reflects a possible neuroprotective function of reactive astrocytes and activated microglia following human traumatic brain injury}},
volume = {22},
year = {2007}
}
@article{Cheng2017,
abstract = {Background/Purpose: Recent studies suggest that hyperuricemia is a potential risk factor for cardiovascular disease (CVD). Hyperuricemia is highly heritable and is associated with sex and body weight. Previous genome-wide association studies have found that the ABCG2 single nucleotide polymorphism (SNP) rs2231142 is an important genetic factor for increased uric acid (UA) levels, and the degree of association between rs2231142 and hyperuricemia is affected by both sex and ethnicity. This investigation aimed to analyze the association be-tween ABCG2 polymorphisms and UA levels, as well as their interactions with sex and obesity in Taiwanese. Methods: Two genetic polymorphisms around the ABCG2 gene were genotyped in 459 patients. Results: After adjusting for clinical covariates, the rs2231142 SNP was found significantly asso-ciated with UA levels using a dominant inheritance model. Patients carrying the rs2231142-A allele had a higher frequency of hyperuricemia than those with the rs2231142-CC allele. Sub-group analysis revealed an association of rs2231142 with UA levels in male or obese patients, and there was no association in nonobese female patients.},
author = {Cheng, Shih-Tsung and Wu, Semon and Su, Cheng-Wen and Teng, Ming-Sheng and Hsu, Lung-An and Ko, Yu-Lin},
doi = {10.1016/j.jfma.2015.12.002},
file = {:Users/solomon/Documents/Mendeley Desktop/Cheng et al. - 2017 - Association of ABCG2 rs2231142-A allele and serum uric acid levels in male and obese individuals in a Han Taiwanes.pdf:pdf},
issn = {09296646},
journal = {Journal of the Formosan Medical Association},
keywords = {obesity,polymorphism,sex,single nucleotide,uric acid},
month = {jan},
number = {1},
pages = {18--23},
title = {{Association of ABCG2 rs2231142-A allele and serum uric acid levels in male and obese individuals in a Han Taiwanese population}},
url = {http://dx.doi.org/10.1016/j.jfma.2015.12.002 www.jfma-onl http://linkinghub.elsevier.com/retrieve/pii/S0929664615004064},
volume = {116},
year = {2017}
}
@article{McDevitt2015,
abstract = {OBJECTIVE: To determine genetic variability within the N-methyl-D-aspartate receptor 2A sub-unit (GRIN2A) gene promoter and its association with concussion recovery time. The hypothesis tested was that there would be a difference in allele and/or genotype distribution between two groups of athletes with normal and prolonged recovery. METHODS: DNA was extracted from saliva collected from a total of 87 athletes with a physician-diagnosed concussion. The (GT) variable number tandem repeats (VNTR) within the promoter region of GRIN2A was genotyped. The long (L) allele was an allele with {\textgreater}/=25 repeats and the short (S) allele was an allele with {\textless}25 repeats in the GT tract. Participants' recovery time was followed prospectively and was categorized as normal ({\textless}/=60 days) or prolonged ({\textgreater}60 days). RESULTS: LL carriers were 6-times more likely to recover longer than 60 days following the concussive event (p = 0.0433) when compared to SS carriers. Additionally, L allele carriers were found more frequently in the prolonged recovery group (p = 0.048). CONCLUSION: Determining genetic influence on concussion recovery will aid in future development of genetic counselling. The clinical relevance of genotyping athletes could improve management of athletes who experience concussion injuries.},
author = {McDevitt, Jane and Tierney, Ryan Timothy and Phillips, Jacqueline and Gaughan, John P and Torg, Joseph S and Krynetskiy, Evgeny},
doi = {10.3109/02699052.2015.1075252},
file = {:Users/solomon/Documents/Mendeley Desktop/McDevitt et al. - 2015 - Association between GRIN2A promoter polymorphism and recovery from concussion.pdf:pdf},
isbn = {0269-9052},
issn = {1362-301X (Electronic)},
journal = {Brain injury},
keywords = {brain injury,calcium,nr2a sub-unit},
number = {13-14},
pages = {1674--1681},
pmid = {26502998},
title = {{Association between GRIN2A promoter polymorphism and recovery from concussion.}},
volume = {29},
year = {2015}
}
@article{Topolovec-Vranic2014,
abstract = {Background: Little empiric research has investigated the interrelationship between homelessness and traumatic brain injury. The objectives of this study were to determine the rate, mechanisms and associated outcomes of traumatic brain injury among men in an urban homeless shelter. Methods: We recruited participants from an urban men's shelter in Toronto, Ontario. Researchers administered the Brain Injury Screening Questionnaire, a semistructured interview screening tool for brain injury. Demographic information and detailed histories of brain injuries were obtained. Participants with positive and negative screening results were compared, and the rates and mechanisms of injury were analyzed by age group. Results: A total of 111 men (mean age 54.2 ± standard deviation 11.5 yr; range 27–81 yr) participated. Nearly half (50 [45{\%}]) of the respondents had a positive screening result for traumatic brain injury. Of these, 73{\%} (35/48) reported experiencing their first injury before adulthood ({\textless} 18 yr), and 87{\%} (40/46) reported a first injury before the onset of homelessness. Among those with a positive screening result, 33 (66{\%}) reported sustaining at least one traumatic brain injury by assault, 22 (44{\%}) by sports or another recre- ational activity, 21 (42{\%}) by motor vehicle collision and 21 (42{\%}) by a fall. A positive screening result was significantly associated with a lifetime history of arrest or mental illness and a parental history of substance abuse. Interpretation: Multiple mechanisms contributed to high rates of traumatic brain injury within a sample of homeless men. Assault was the most common mechanism, with sports and recreation, motor vehicle collisions and falls also being reported frequently by the par- ticipants. Injury commonly predated the onset of homelessness, with most participants experiencing their first injury in childhood. Additional research is needed to understand the complex interactions among homelessness, traumatic brain injury, mental illness and substance use.},
archivePrefix = {arXiv},
arxivId = {NIHMS150003},
author = {Topolovec-Vranic, J. and Ennis, N. and Howatt, M. and Ouchterlony, D. and Michalak, a. and Masanic, C. and Colantonio, a. and Hwang, S. W. and Kontos, P. and Stergiopoulos, V. and Cusimano, M. D.},
doi = {10.9778/cmajo.20130046},
eprint = {NIHMS150003},
file = {:Users/solomon/Documents/Mendeley Desktop/Topolovec-Vranic et al. - 2014 - Traumatic brain injury among men in an urban homeless shelter observational study of rates and mechanis.pdf:pdf},
isbn = {2291-0026 (Electronic)$\backslash$r2291-0026 (Linking)},
issn = {2291-0026},
journal = {CMAJ Open},
number = {2},
pages = {E69--E76},
pmid = {25077132},
title = {{Traumatic brain injury among men in an urban homeless shelter: observational study of rates and mechanisms of injury}},
url = {http://cmajopen.ca/cgi/doi/10.9778/cmajo.20130046},
volume = {2},
year = {2014}
}
@article{Zhang2013,
abstract = {This study investigated whether deferoxamine (DFO), an iron chelator attenuates iron-induced toxicity in rats with traumatic brain injury. In this study, three groups of Sprague–Dawley rats (sham, injury and DFO groups) were examined. Rats were killed on day 28 after Morris water maze testing and brains perfused for either non-heme brain binding or hemosiderin staining. Western blotting was used to measure protein levels of ferritin, transferrin and transient receptor potential canonical channel 6 (TRPC6). In TBI rats, there was a significant increase in brain iron on day 28, ferritin L, ferritin H, transferrin and TRPC6 levels were all significantly elevated post-TB1. There were also deficits in spatial learning and memory; however, DFO administration attenuated these effects in TBI rats supporting the notion that DFO may reduce brain injury accentuated by iron overload. (PsycINFO Database Record (c) 2016 APA, all rights reserved)},
author = {Zhang, Lijun and Hu, Rong and Li, Mei and Li, Fei and Meng, Hui and Zhu, Gang and Lin, Jiangkai and Feng, Hua},
doi = {10.1007/s10072-012-1090-1},
file = {:Users/solomon/Documents/Mendeley Desktop/Zhang et al. - 2013 - Deferoxamine attenuates iron-induced long-term neurotoxicity in rats with traumatic brain injury.pdf:pdf},
isbn = {1590-3478 (Electronic)$\backslash$r1590-1874 (Linking)},
issn = {15901874},
journal = {Neurological Sciences},
keywords = {Deferoxamine,Iron,Spatial learning and memory,TRPC6,Traumatic brain injury},
number = {5},
pages = {639--645},
pmid = {22538758},
title = {{Deferoxamine attenuates iron-induced long-term neurotoxicity in rats with traumatic brain injury}},
volume = {34},
year = {2013}
}
@article{Shojo2010,
abstract = {Traumatic brain injury (TBI) causes massive brain damage. However, the secondary injury and temporal sequence of events with multiple mechanisms after the insult has not been elucidated. Here, we examined the occurrence of apoptosis and a causal relationship between inflammation and apoptosis in the TBI brain. Following a lateral moderate fluid percussion injury model of TBI in adult rats, microarray analyses detected apparent changes in the expression levels of apoptosis-related genes which revealed time-dependent expression patterns for 23 genes in the lateral cortex. The upregulated 23 genes included inflammatory cytokines such as interleukin 1 (IL-1) ??, IL-1??, and tumor necrotic factor (TNF) which immediately increased at 3 h following the injury. Time-dependent gene expression profile analyses showed that apoptosis was subsequently induced following inflammation. These results taken together suggested changes in expression of apoptosis-related genes may be associated with inflammatory response. Accompanying this surge of cell death genes after TBI was a neurostructural pathologic hallmark of apoptosis characterized by leakage of cytochrome c into cytoplasm, DNA fragmentation and apoptotic cells in the lateral cortex of the impacted hemisphere. Caspase-3 positive cells in the TBI brain were initially sporadic after 3 h, but these apoptotic cells subsequently increased and populated the cerebral cortex at 6 and 12 h, and gradually reached a plateau by 48 h. Interestingly, the expression profile of CD68 macrophage labeled cells closely resembled that of apoptotic cells after TBI, including the role of inflammatory signaling pathway in the progression of apoptotic cell death. These results taken together suggest that TBI induced upregulation of apoptosis-related genes, concomitant with the detection of apoptotic brain pathology during the 3-48 h post-injury period, which may be likely mediated by inflammation. Therapies designed at abrogating apoptosis and/or inflammation may prove effective when initiated at this subacute TBI phase. ?? 2010 IBRO.},
author = {Shojo, H. and Kaneko, Y. and Mabuchi, T. and Kibayashi, K. and Adachi, N. and Borlongan, C. V.},
doi = {10.1016/j.neuroscience.2010.10.018},
file = {:Users/solomon/Documents/Mendeley Desktop/Shojo et al. - 2010 - Genetic and histologic evidence implicates role of inflammation in traumatic brain injury-induced apoptosis in the.pdf:pdf},
isbn = {8155273954},
issn = {03064522},
journal = {Neuroscience},
keywords = {Apoptosis-related genes,Brain injury model,Gene expression,Head trauma,Inflammation},
number = {4},
pages = {1273--1282},
pmid = {20950674},
publisher = {Elsevier Inc.},
title = {{Genetic and histologic evidence implicates role of inflammation in traumatic brain injury-induced apoptosis in the rat cerebral cortex following moderate fluid percussion injury}},
url = {http://dx.doi.org/10.1016/j.neuroscience.2010.10.018},
volume = {171},
year = {2010}
}
@article{Dekker2014,
abstract = {BACKGROUND: We have previously shown that addition of valproic acid (VPA; a histone deacetylase inhibitor) to hetastarch (Hextend [HEX]) resuscitation significantly decreases lesion size in a swine model of traumatic brain injury (TBI) and hemorrhagic shock (HS). However, the precise mechanisms have not been well defined. As VPA is a transcriptional modulator, the aim of this study was to investigate its effect on brain gene expression profiles. METHODS: Swinewere subjected to controlled TBI and HS (40{\%} blood volume), kept in shock for 2 hours, and resuscitated with HEX or HEX + VPA (n = 5 per group). Following 6 hours of observation, brain RNAwas isolated, and gene expression profiles were measured using a Porcine Gene ST 1.1 microarray (Affymetrix, Santa Clara, CA). Pathway analysis was done using network analysis tools Gene Ontology, Ingenuity PathwayAnalysis, and Parametric Gene SetEnrichmentAnalysis.Real-timepolymerase chain reaction was used to verify the key microarray findings. RESULTS: A total of 1,668 probe sets mapping to 370 known genes were differentially expressed between the HEX and HEX + VPA groups. Expression of apoptotic genes differed between groups, and biologic function analysis predicted a significant downregulation of apoptosis (p = 1.29? 10j12), cell death (p = 8.46?10j12), and necrosis (p = 9.07? 10j11). Pathway analysis indicated a significant modulation of pathways involved in cell signaling, dendritic cell response, and the complement system. CONCLUSION: This is the first high-throughput analysis of cerebral gene profiling following TBI + HS. It shows that treatment with VPA significantly alters early transcription of pathways related to cell survival,which may explain its neuroprotective effects.},
author = {Dekker, Simone E. and Bambakidis, Ted and Sillesen, Martin and Liu, Baoling and Johnson, Craig N. and Jin, Guang and Li, Yongqing and Alam, Hasan B.},
doi = {10.1097/TA.0000000000000345},
file = {:Users/solomon/Documents/Mendeley Desktop/Dekker et al. - 2014 - Effect of pharmacologic resuscitation on the brain gene expression profiles in a swine model of traumatic brain i.pdf:pdf},
isbn = {0000000000000},
issn = {2163-0755},
journal = {Journal of Trauma and Acute Care Surgery},
keywords = {adults and,and mortality in young,apoptosis,brain,genes,hemorrhage,injury,is one of the,leading causes,morbidity,of disability,pigs,raumatic brain injury,tbi},
number = {6},
pages = {906--912},
title = {{Effect of pharmacologic resuscitation on the brain gene expression profiles in a swine model of traumatic brain injury and hemorrhage}},
url = {http://content.wkhealth.com/linkback/openurl?sid=WKPTLP:landingpage{\&}an=01586154-201412000-00016},
volume = {77},
year = {2014}
}
@article{Wagner2010,
abstract = {Globally, traumatic brain injury (TBI) is a leading cause of death and persistent disability. Although improvements in standard of care have reduced the overall mortality rates associated with this disease, there is a paucity of effective neuroprotective therapies. However, some rehabilitation focused strategies have shown promise with enhancing neurorecovery. One major challenge in identifying effective therapies is that TBI is an inherently heterogeneous disease. Despite many patients having similar injury factors and clinical care after their TBI, recovery and outcomes can be very different. This commentary discusses the value of treatment effectiveness research that also incorporates theranostic principles of individualized care. Key to this concept is the utilization of state-of-the-art biomarker platforms and technologies that can identify relevant molecular or physiological fingerprints that can assist rehabilitation practitioners in delineating long term outcome that can be linked to plasticity, treatment response, and natural recovery. This commentary proposes a unique concept of "Rehabilomics" as a field of study involving the systematic collection and study of rehabilitation relevant phenotypes, in conjunction with a transdisciplinary evaluation of biomarkers, in order to better understand the biology, function, prognosis, complications, treatments, adaptation, and recovery for persons with disabilities. Specific Rehabilomics applications to TBI research, as well as relevance to the National Institutes of Health Roadmap, are discussed.},
author = {Wagner, AK},
doi = {R33102398 [pii]},
isbn = {1973-9095 (Electronic)$\backslash$n1973-9087 (Linking)},
issn = {19739087},
journal = {European Journal of Physical and Rehabilitation Medicine},
keywords = {Brain injuries - Rehabilitation - Biological marke},
number = {4},
pages = {549--555},
pmid = {21224787},
title = {{TBI translational rehabilitation research in the 21st Century: Exploring a Rehabilomics research model}},
volume = {46},
year = {2010}
}
@article{Kelly2004,
author = {Kelly, G and McNally, T and Thepaut, J},
file = {:Users/solomon/Documents/Mendeley Desktop/Kelly, McNally, Thepaut - 2004 - OSE of all main data types in the ECMWF operational system.pdf:pdf},
journal = {Third WMO Workshop on the Impact of},
pages = {2--6},
title = {{OSE of all main data types in the ECMWF operational system}},
url = {http://scholar.google.com/scholar?hl=en{\&}btnG=Search{\&}q=intitle:ose{\#}4},
year = {2004}
}
@article{Macalou2016,
abstract = {The ATP-binding cassette (ABC) transporters of class G display a different domain organisation than P-glycoprotein/ABCB1 and bacterial homologues with a nucleotide-binding domain preceding the transmembrane domain. The linker region connecting these domains is unique and its function and structure cannot be predicted. Sequence analysis revealed that the human ABCG2 linker contains a LSGGE sequence, homologous to the canonical C-motif/ABC signature present in all ABC nucleotide- binding domains. Predictions of disorder and of secondary structures indicated that this C2-sequence was highly mobile and located between an a-helix and a loop similarly to the C-motif. Point mutations of the two first residues of the C2-sequence fully abolished the transport-coupled ATPase activity, and led to the complete loss of cell resistance to mitoxantrone. The interaction with potent, selective and non-competitive, ABCG2 inhibitors was also significantly an important mechanistic role for the C2-sequence of the ABCG2 linker region in ATP binding and/or hydrolysis coupled to drug efflux.},
author = {Macalou, S. and Robey, R. W. and {Jabor Gozzi}, G. and Shukla, S. and Grosjean, I. and Hegedus, T. and Ambudkar, S. V. and Bates, S. E. and {Di Pietro}, A.},
doi = {10.1007/s00018-015-2118-5},
file = {:Users/solomon/Documents/Mendeley Desktop/Macalou et al. - 2016 - The linker region of breast cancer resistance protein ABCG2 is critical for coupling of ATP-dependent drug trans.pdf:pdf},
issn = {14209071},
journal = {Cellular and Molecular Life Sciences},
keywords = {ABC transporter,ATP hydrolysis,Breast cancer resistance protein/ABCG2,C motif/ABC signature,Drug efflux coupling,Specific sequence},
number = {9},
pages = {1927--1937},
pmid = {26708291},
publisher = {Springer International Publishing},
title = {{The linker region of breast cancer resistance protein ABCG2 is critical for coupling of ATP-dependent drug transport}},
volume = {73},
year = {2016}
}
@article{Lin2002,
author = {Lin, Y. S.},
doi = {10.1124/mol.62.1.162},
file = {:Users/solomon/Documents/Mendeley Desktop/Lin - 2002 - Co-Regulation of CYP3A4 and CYP3A5 and Contribution to Hepatic and Intestinal Midazolam Metabolism.pdf:pdf},
issn = {0026895X},
journal = {Molecular Pharmacology},
number = {1},
pages = {162--172},
title = {{Co-Regulation of CYP3A4 and CYP3A5 and Contribution to Hepatic and Intestinal Midazolam Metabolism}},
url = {http://molpharm.aspetjournals.org/cgi/doi/10.1124/mol.62.1.162},
volume = {62},
year = {2002}
}
@article{Eloranta2012,
abstract = {Organic anion-transporting polypeptide 1A2 (OATP1A2) (gene symbol, SLCO1A2) mediates cellular uptake of a wide range of endogenous substrates, as well as drugs and xenobiotics. OATP1A2 is expressed in several tissues, including apical membranes of small intestinal epithelial cells. Given its role in intestinal drug absorption, a detailed analysis of the mechanisms that regulate SLCO1A2 gene expression is potentially of great pharmacological relevance. We show here that treatment of human intestine-derived Caco-2 cells with vitamin D(3) markedly increased endogenous OATP1A2 mRNA and protein levels. Suppression of endogenous vitamin D receptor (VDR) expression with siRNAs significantly reduced this induction. Two alternative promoter regions exist in genomic databases for the SLCO1A2 gene. One putative VDR response element (VDRE) that was predicted to interact efficiently with VDR-retinoid X receptor $\alpha$ (RXR$\alpha$) was identified in silico within SLCO1A2 promoter variant 1. This VDRE served as a strong binding site for the recombinant VDR-RXR$\alpha$ heterodimers in vitro and was potently activated by VDR in the presence of vitamin D(3) in heterologous promoter assays. In reporter assays using native promoter constructs, SLCO1A2 promoter variant 1 was strongly induced by VDR, and site-directed mutagenesis of a single VDRE within this region abolished this activation. Native VDR-RXR$\alpha$ also interacted with this element both in vitro and in living cells. We showed that expression of the SLCO1A2 gene is induced by vitamin D(3) at the transcriptional level through the VDR. Our results suggest that pharmacological administration of vitamin D(3) may allow modulation of intestinal absorption of OATP1A2 transport substrates.},
author = {Eloranta, Jyrki J and Hiller, Christian and Juttner, M. and Kullak-Ublick, Gerd A},
doi = {10.1124/mol.112.077909},
issn = {1521-0111},
journal = {Molecular Pharmacology},
month = {jul},
number = {1},
pages = {37--46},
pmid = {22474172},
publisher = {American Society for Pharmacology and Experimental Therapeutics},
title = {{The SLCO1A2 Gene, Encoding Human Organic Anion-Transporting Polypeptide 1A2, Is Transactivated by the Vitamin D Receptor}},
url = {http://www.ncbi.nlm.nih.gov/pubmed/22474172 http://molpharm.aspetjournals.org/cgi/doi/10.1124/mol.112.077909},
volume = {82},
year = {2012}
}
@article{Kalsotra2003,
author = {Kalsotra, Auinash and Turman, Cheri M. and Dash, Pramod K. and Strobel, Henry W},
doi = {10.1089/089771503322686139},
file = {:Users/solomon/Documents/Mendeley Desktop/Kalsotra et al. - 2003 - Differential Effects of Traumatic Brain Injury on the Cytochrome P450 System A Perspective into Hepatic and Ren.pdf:pdf},
issn = {0897-7151},
journal = {Journal of Neurotrauma},
keywords = {cyp450 drug substrates,inflammation and injury,leukotriene b 4,tissue-specific response},
month = {dec},
number = {12},
pages = {1339--1350},
title = {{Differential Effects of Traumatic Brain Injury on the Cytochrome P450 System: A Perspective into Hepatic and Renal Drug Metabolism}},
url = {http://www.liebertpub.com/doi/10.1089/089771503322686139},
volume = {20},
year = {2003}
}
@article{Anderson2009,
author = {Anderson, GD and Temkin, NR and Dikmen, SS and Diaz-Arrastia, R and Machamer, JE and Farhrenbruch, C and Miller, JW and Sadrzadeh, SMH},
doi = {10.1016/j.yebeh.2009.08.025},
file = {:Users/solomon/Documents/Mendeley Desktop/Anderson et al. - 2009 - Haptoglobin phenotype and apolipoprotein E polymorphism Relationship to posttraumatic seizures and neuropsychol.pdf:pdf},
issn = {15255050},
journal = {Epilepsy Behav.},
keywords = {apolipoprotein e,haptoglobin,post-traumatic seizures,traumatic brain injury},
month = {nov},
number = {3},
pages = {501--506},
title = {{Haptoglobin phenotype and apolipoprotein E polymorphism: Relationship to posttraumatic seizures and neuropsychological functioning after traumatic brain injury}},
url = {http://linkinghub.elsevier.com/retrieve/pii/S1525505009004879},
volume = {16},
year = {2009}
}
@techreport{CentersforDiseaseControlandPrevention2015,
address = {Atlanta, GA},
author = {{Centers for Disease Control and Prevention}},
file = {:Users/solomon/Documents/Mendeley Desktop/Centers for Disease Control and Prevention - 2015 - Report to Congress on Traumatic Brain Injury in the United States Epidemiology and R.pdf:pdf},
institution = {National Center for Injury Prevention and Control},
title = {{Report to Congress on Traumatic Brain Injury in the United States: Epidemiology and Rehabilitation}},
year = {2015}
}
@article{Schumacher2012,
abstract = {BACKGROUND: ATP-binding cassette (ABC) transporters are essential regulators of organismic homeostasis, and are particularly important in protecting the body from potentially harmful exogenous substances. Recently, an increasing number of in vitro observations have indicated a functional role of ABC transporters in the differentiation and maintenance of stem cells. Therefore, we sought to determine brain-related phenotypic changes in animals lacking the expression of distinct ABC transporters (ABCB1, ABCG2 or ABCC1). METHODOLOGY AND PRINCIPAL FINDINGS: Analyzing adult neurogenesis in ABC transporter-deficient animals in vivo and neuronal stem/progenitor cells in vitro resulted in complex findings. In vivo, the differentiation of neuronal progenitors was hindered in ABC transporter-deficient mice (ABCB1(0/0)) as evidenced by lowered numbers of doublecortin(+) (-36{\%}) and calretinin(+) (-37{\%}) cells. In vitro, we confirmed that this finding is not connected to the functional loss of single neural stem/progenitor cells (NSPCs). Furthermore, assessment of activity, exploratory behavior, and anxiety levels revealed behavioral alterations in ABCB1(0/0) and ABCC1(0/0) mice, whereas ABCG2(0/0) mice were mostly unaffected. CONCLUSION AND SIGNIFICANCE: Our data show that single ABC transporter-deficiency does not necessarily impair neuronal progenitor homeostasis on the single NSPC level, as suggested by previous studies. However, loss of distinct ABC transporters impacts global brain homeostasis with far ranging consequences, leading to impaired neurogenic functions in vivo and even to distinct behavioral phenotypes. In addition to the known role of ABC transporters in proteopathies such as Parkinson's disease and Alzheimer's disease, our data highlight the importance of understanding the general function of ABC transporters for the brain's homeostasis and the regeneration potential.},
author = {Schumacher, Toni and Krohn, Markus and Hofrichter, Jacqueline and Lange, Cathleen and Stenzel, Jan and Steffen, Johannes and Dunkelmann, Tina and Paarmann, Kristin and Fr{\"{o}}hlich, Christina and Uecker, Annekathrin and Plath, Anne Sophie and Sommer, Alexandra and Br{\"{u}}ning, Thomas and Heinze, Hans Jochen and Pahnke, Jens},
doi = {10.1371/journal.pone.0035613},
file = {:Users/solomon/Documents/Mendeley Desktop/Schumacher et al. - 2012 - ABC transporters B1, C1 and G2 differentially regulate neuroregeneration in mice.pdf:pdf},
isbn = {1932-6203 (Electronic)$\backslash$n1932-6203 (Linking)},
issn = {19326203},
journal = {PLoS ONE},
number = {4},
pmid = {22545122},
title = {{ABC transporters B1, C1 and G2 differentially regulate neuroregeneration in mice}},
volume = {7},
year = {2012}
}
@article{Barr2011,
abstract = {Several clinical trials have failed to demonstrate a significant effect on outcome following human traumatic brain injury (TBI) despite promising results obtained in preclinical animal studies. These failures may be due in part to a misinterpretation of the findings obtained in preclinical animal models of TBI, a misunderstanding of the complexity of the human response to TBI, limited knowledge about the biological pathways that interact to contribute to good and bad outcomes after brain injury, and the effects of genomic variability and environment on individual recovery. Recent publications suggest that data obtained from gene expression profiling studies of complex neurological diseases such as stroke, multiple sclerosis (MS), Alzheimer's and Parkinson's may contribute to a more informed understanding of what affects outcome following TBI. These data may help to bridge the gap between successful preclinical studies and negative clinical trials in humans to reveal novel targets for therapy. Gene expression profiling has the capability to identify biomarkers associated with response to TBI, elucidate complex genetic interactions that may play a role in outcome following TBI, and reveal biological pathways related to brain health. This review highlights the current state of the literature on gene expression profiling for neurological disease and discusses its ability to aid in unraveling the variable human response to TBI and the potential for it to offer treatment strategies in an area where we currently have limited therapeutic options primarily based on supportive care.},
author = {Barr, Taura L and Alexander, Sheila and Conley, Yvette},
doi = {10.1177/1099800410385671},
file = {:Users/solomon/Documents/Mendeley Desktop/Barr, Alexander, Conley - 2011 - Gene expression profiling for discovery of novel targets in human traumatic brain injury.pdf:pdf},
isbn = {1099800410385},
issn = {1552-4175},
journal = {Biological Research for Nursing},
keywords = {Brain Injuries,Brain Injuries: drug therapy,Brain Injuries: genetics,Drug Discovery,Gene Expression Profiling,Humans,Oligonucleotide Array Sequence Analysis},
number = {2},
pages = {140--53},
pmid = {21112922},
title = {{Gene expression profiling for discovery of novel targets in human traumatic brain injury.}},
url = {http://www.ncbi.nlm.nih.gov/pubmed/21112922},
volume = {13},
year = {2011}
}
@article{Yang2014,
abstract = {ATP-binding cassette (ABC) transporters are implicated in a diverse range of physiological and pathophysiological processes, such as cholesterol and lipid transportation and multidrug resistance. Despite the considerable efforts made in understanding of the cellular function of ABC proteins, the regulation mechanism of this type of protein is still poorly defined. Here we report the identification and functional characterization of a novel ATPase protein, protein associated with ABC transporters (PAAT), in humans. PAAT contains a nucleotide-binding domain (NBD)-like domain and a signal for intramitochondrial sorting. We showed that PAAT is localized in both the cytoplasm and the mitochondria and has an intrinsic ATPase activity. PAAT physically interacts with the 3 known mitochondrial inner membrane ABC proteins, ABCB7, ABCB8, and ABCB10, but not ABCB1, ABCB6, or ABCG2, and functionally regulates the transport of ferric nutrients and heme biosynthesis. Significantly, PAAT deficiency promotes cell death, reduces mitochondrial potential, and sensitizes mitochondria to oxidative stress-induced DNA damages. Our experiments revealed that PAAT is a novel ATPase and a trans-regulator of mitochondrial ABC transporters that plays an important role in the maintenance of mitochondrial homeostasis and cell survival.-Yang, X., Yang, J., Li, L., Sun, L., Yi, X., Han, X., Si, W., Yan, R., Chen, Z., Xie, G., Li, W., Shang, Y., Liang, J. PAAT, a novel ATPase and trans-regulator of mitochondrial ABC transporters, is critically involved in the maintenance of mitochondrial homeostasis.},
author = {Yang, Xiaohan and Yang, Jianguo and Li, Lei and Sun, Luyang and Yi, Xia and Han, Xiao and Si, Wenzhe and Yan, Ruorong and Chen, Zhe and Xie, Guojia and Li, Wanjin and Shang, Yongfeng and Liang, Jing},
doi = {10.1096/fj.14-254045},
file = {:Users/solomon/Documents/Mendeley Desktop/Yang et al. - 2014 - PAAT, a novel ATPase and trans-regulator of mitochondrial ABC transporters, is critically involved in the maintenan.pdf:pdf},
isbn = {1530-6860 (Electronic)$\backslash$r0892-6638 (Linking)},
issn = {15306860},
journal = {FASEB Journal},
keywords = {Cell death,Iron metabolism,NBD domain,Oxidative stress},
number = {11},
pages = {4821--4834},
pmid = {25063848},
title = {{PAAT, a novel ATPase and trans-regulator of mitochondrial ABC transporters, is critically involved in the maintenance of mitochondrial homeostasis}},
volume = {28},
year = {2014}
}
@article{Chen2017a,
abstract = {Targeted next-generation sequencing (NGS) has been widely used as a cost-effective way to identify the genetic basis of human disorders. Copy number variations (CNVs) contribute significantly to human genomic variability, some of which can lead to disease. However, effective detection of CNVs from targeted capture sequencing data remains challenging. Here we present SeqCNV, a novel CNV calling method designed to use capture NGS data. SeqCNV extracts the read depth information and utilizes the maximum penalized likelihood estimation (MPLE) model to identify the copy number ratio and CNV boundary. We applied SeqCNV to both bacterial artificial clone (BAC) and human patient NGS data to identify CNVs. These CNVs were validated by array comparative genomic hybridization (aCGH). SeqCNV is able to robustly identify CNVs of different size using capture NGS data. Compared with other CNV-calling methods, SeqCNV shows a significant improvement in both sensitivity and specificity.},
author = {Chen, Yong and Zhao, Li and Wang, Yi and Cao, Ming and Gelowani, Violet and Xu, Mingchu and Agrawal, Smriti A. and Li, Yumei and Daiger, Stephen P. and Gibbs, Richard and Wang, Fei and Chen, Rui},
doi = {10.1186/s12859-017-1566-3},
file = {:Users/solomon/Documents/Mendeley Desktop/Chen et al. - 2017 - SeqCNV A novel method for identification of copy number variations in targeted next-generation sequencing data.pdf:pdf},
issn = {14712105},
journal = {BMC Bioinformatics},
keywords = {Copy number variation,Maximum penalized likelihood estimation,Next-generation sequencing},
number = {1},
pages = {1--9},
pmid = {28253855},
publisher = {BMC Bioinformatics},
title = {{SeqCNV: A novel method for identification of copy number variations in targeted next-generation sequencing data}},
volume = {18},
year = {2017}
}
@article{JohansenTaber2014,
abstract = {BACKGROUND: The use of pharmacogenomic testing in the clinical setting has the potential to improve the safety and effectiveness of drug therapy, yet studies have revealed that physicians lack knowledge about the topic of pharmacogenomics, and are not prepared to implement it in the clinical setting. This study further explores the pharmacogenomic knowledge deficit and educational resource needs among physicians.MATERIALS AND METHODS: Surveys of primary care physicians, cardiologists, and psychiatrists were conducted.RESULTS: Few physicians reported familiarity with the topic of pharmacogenomics, but more reported confidence in their knowledge about the influence of genetics on drug therapy. Only a small minority had undergone formal training in pharmacogenomics, and a majority reported being unsure what type of pharmacogenomic tests were appropriate to order for the clinical situation. Respondents indicated that an ideal pharmacogenomic educational resource should be electronic and include such components as how to interpret pharmacogenomic test results, recommendations for prescribing, population subgroups most likely to be affected, and contact information for laboratories offering pharmacogenomic testing.CONCLUSION: Physicians continue to demonstrate pharmacogenomic knowledge gaps, and are unsure about how to use pharmacogenomic testing in clinical practice. Educational resources that are clinically oriented and easily accessible are preferred by physicians, and may best support appropriate clinical implementation of pharmacogenomics},
author = {{Johansen Taber}, Katherine A. and Dickinson, Barry D.},
doi = {10.2147/PGPM.S63715},
file = {:Users/solomon/Documents/Mendeley Desktop/Johansen Taber, Dickinson - 2014 - Pharmacogenomic knowledge gaps and educational resource needs among physicians in selected specialtie.pdf:pdf},
isbn = {1178-7066},
issn = {11787066},
journal = {Pharmacogenomics and Personalized Medicine},
keywords = {Drug response,Educational resource,Knowledge gap,Pharmacogenomics,drug response,educational resource,knowledge gap,pharmacogenomics},
month = {jul},
number = {1},
pages = {145},
pmid = {25045280},
publisher = {Dove Press},
title = {{Pharmacogenomic knowledge gaps and educational resource needs among physicians in selected specialties}},
url = {http://www.ncbi.nlm.nih.gov/pubmed/25045280 http://www.pubmedcentral.nih.gov/articlerender.fcgi?artid=PMC4100727 http://www.dovepress.com/pharmacogenomic-knowledge-gaps-and-educational-resource-needs-among-ph-peer-reviewed-article-PGPM},
volume = {7},
year = {2014}
}
@article{Stroupe2013,
abstract = {Approximately 15{\%} of casualties in the Afghanistan (Operation Enduring Freedom [OEF]) and Iraq (Operation Iraqi Freedom [OIF]) conflicts received mild traumatic brain injury (TBI). To identify Veterans who may benefit from treatment, the Department of Veterans Affairs (VA) implemented a national clinical reminder in 2007 to screen for TBI. Veterans who screen positive are referred for a comprehensive TBI evaluation. We conducted a national retrospective study of OIF/OEF Veterans receiving care at VA facilities between 2007 and 2008. We examined the association of the TBI screen with healthcare costs over a 12 mo period following the initial evaluation. Of the Veterans, 164,438 met inclusion criteria: 31,627 screened positive, 118,545 screened negative, and 14,266 received no TBI screening. Total healthcare costs of Veterans who screened positive, screened negative, or had no TBI screening were {\$}9,610, {\$}5,184, and {\$}3,399, respectively (p {\textless} 0.001). Understanding these healthcare utilization and cost patterns will assist policy-makers to address the ongoing and future healthcare needs of these returning Veterans. [ABSTRACT FROM AUTHOR]},
author = {Stroupe, Kevin T and Smith, Bridget M and Hogan, Timothy P and {St. Andre}, Justin R and Pape, Theresa and Steiner, Monica L and Proescher, Eric and Huo, Zhiping and Evans, Charlesnika T},
doi = {10.1682/JRRD.2012.06.0107},
file = {:Users/solomon/Documents/Mendeley Desktop/Stroupe et al. - 2013 - Healthcare utilization and costs of Veterans screened and assessed for traumatic brain injury.pdf:pdf},
isbn = {0748-7711},
issn = {07487711},
journal = {Journal of Rehabilitation Research {\&} Development},
keywords = {ANALYSIS of variance,BRAIN -- Wounds {\&} injuries,BRAIN -- Wounds {\&} injuries -- Prevention,CHI-squared test,COMORBIDITY,CONFIDENCE intervals,DATA analysis,DATA analysis -- Software,DATABASES,DESCRIPTIVE statistics,EPIDEMIOLOGY,INFORMATION storage {\&} retrieval systems -- Medicin,LONGITUDINAL method,MEDICAL care -- Utilization,MEDICAL care costs,MENTAL health services,PATIENTS,PRIMARY health care,PROBABILITY theory,RESEARCH -- Finance,RETROSPECTIVE studies,SOCIOECONOMIC factors,UNITED States,VETERANS,VETERANS' hospitals},
number = {8},
pages = {1047--1068},
pmid = {24458891},
title = {{Healthcare utilization and costs of Veterans screened and assessed for traumatic brain injury.}},
url = {10.1682/JRRD.2012.06.0107{\%}5Cnhttp://offcampus.lib.washington.edu/login?url=http://search.ebscohost.com/login.aspx?direct=true{\&}db=a9h{\&}AN=93279747{\&}site=ehost-live},
volume = {50},
year = {2013}
}
@article{Bellin2014,
archivePrefix = {arXiv},
arxivId = {NIHMS150003},
author = {Bellin, Melena D. and Gelrud, Andres and Arreaza-Rubin, Guillermo and Dunn, Ty B. and Humar, Abhinav and Morgan, Katherine A. and Naziruddin, Bashoo and Rastellini, Cristiana and Rickels, Michael R. and Schwarzenberg, Sarah J. and Andersen, Dana K.},
doi = {10.1097/MPA.0000000000000236},
eprint = {NIHMS150003},
file = {:Users/solomon/Documents/Mendeley Desktop/Bellin et al. - 2014 - Total Pancreatectomy With Islet Autotransplantation.pdf:pdf},
isbn = {0000000000000},
issn = {0885-3177},
journal = {Pancreas},
month = {nov},
number = {8},
pages = {1163--1171},
pmid = {1000000221},
title = {{Total Pancreatectomy With Islet Autotransplantation}},
url = {https://insights.ovid.com/crossref?an=00006676-201411000-00007},
volume = {43},
year = {2014}
}
@article{Livak2001,
abstract = {The two most commonly used methods to analyze data from real-time, quantitative},
archivePrefix = {arXiv},
arxivId = {1003.3921v1},
author = {Livak, K J and Schmittgen, T D},
doi = {10.1006/meth.2001.1262},
eprint = {1003.3921v1},
isbn = {1046-2023},
issn = {1046-2023 (Print)1046-2023},
journal = {Methods},
keywords = {quantitative polymerase chain reaction,reaction,real-time polymerase chain reaction,relative quantification,reserve transcription combined with,reverse transcription polymerase chain,taq man,the polymer-},
pages = {402--408},
pmid = {11846609},
title = {{Analysis of relative gene expression data using real-time quantitative PCR and}},
volume = {25},
year = {2001}
}
@article{Bombardier2010,
abstract = {CONTEXT Uncertainties exist about the rates, predictors, and outcomes of major depressive disorder (MDD) among individuals with traumatic brain injury (TBI). OBJECTIVE To describe MDD-related rates, predictors, outcomes, and treatment during the first year after TBI. DESIGN Cohort from June 2001 through March 2005 followed up by structured telephone interviews at months 1 through 6, 8, 10, and 12 (data collection ending February 2006). SETTING Harborview Medical Center, a level I trauma center in Seattle, Washington. PARTICIPANTS Five hundred fifty-nine consecutively hospitalized adults with complicated mild to severe TBI. MAIN OUTCOME MEASURES The Patient Health Questionnaire (PHQ) depression and anxiety modules were administered at each assessment and the European Quality of Life measure was given at 12 months. RESULTS Two hundred ninety-seven of 559 patients (53.1{\%}) met criteria for MDD at least once in the follow-up period. Point prevalences ranged between 31{\%} at 1 month and 21{\%} at 6 months. In a multivariate model, risk of MDD after TBI was associated with MDD at the time of injury (risk ratio [RR], 1.62; 95{\%} confidence interval [CI], 1.37-1.91), history of MDD prior to injury (but not at the time of injury) (RR, 1.54; 95{\%} CI, 1.31-1.82), age (RR, 0.61; 95{\%} CI, 0.44-0.83 for {\textgreater} or = 60 years vs 18-29 years), and lifetime alcohol dependence (RR, 1.34; 95{\%} CI, 1.14-1.57). Those with MDD were more likely to report comorbid anxiety disorders after TBI than those without MDD (60{\%} vs 7{\%}; RR, 8.77; 95{\%} CI, 5.56-13.83). Only 44{\%} of those with MDD received antidepressants or counseling. After adjusting for predictors of MDD, persons with MDD reported lower quality of life at 1 year compared with the nondepressed group. CONCLUSIONS Among a cohort of patients hospitalized for TBI, 53.1{\%} met criteria for MDD during the first year after TBI. Major depressive disorder was associated with history of MDD and was an independent predictor of poorer health-related quality of life.},
author = {Bombardier, CH and Fann, JR and Temkin, NR and Esselman, PC and Barber, J and Dikmen, SS},
doi = {10.1001/jama.2010.599},
file = {:Users/solomon/Documents/Mendeley Desktop/Bombardier et al. - 2010 - Rates of major depressive disorder and clinical outcomes following traumatic brain injury.pdf:pdf},
issn = {1538-3598},
journal = {JAMA},
month = {may},
number = {19},
pages = {1938--45},
pmid = {20483970},
title = {{Rates of major depressive disorder and clinical outcomes following traumatic brain injury.}},
url = {http://jama.jamanetwork.com/article.aspx?articleid=185892 http://jama.jamanetwork.com/article.aspx?doi=10.1001/jama.2010.599 http://www.ncbi.nlm.nih.gov/pubmed/20483970 http://www.pubmedcentral.nih.gov/articlerender.fcgi?artid=PMC3090293},
volume = {303},
year = {2010}
}
@article{Kochanek2006,
author = {Kochanek, PM and Vagni, VA and Janesko, KL and Washington, CB and Crumrine, PK and Garman, RH and Jenkins, LW and Clark, RSB and Homanics, GE and Dixon, CE and Schnermann, J and Jackson, EK},
doi = {10.1038/sj.jcbfm.9600218},
file = {:Users/solomon/Documents/Mendeley Desktop/Kochanek et al. - 2006 - Adenosine A1 receptor knockout mice develop lethal status epilepticus after experimental traumatic brain injury.pdf:pdf},
issn = {0271-678X (Print) 0271-678X (Linking)},
journal = {J Cereb Blood Flow Metab},
number = {4},
pages = {565--575},
title = {{Adenosine A1 receptor knockout mice develop lethal status epilepticus after experimental traumatic brain injury}},
type = {Journal Article},
url = {https://www.ncbi.nlm.nih.gov/pubmed/16121125},
volume = {26},
year = {2006}
}
@article{Nagashima2013,
abstract = {Powerful array-based single-nucleotide polymorphism-typing platforms have recently heralded a new era in which genome-wide studies are conducted with increasing frequency. A genetic polymorphism associated with population pharmacokinetics (PK) is typically analyzed using nonlinear mixed-effect models (NLMM). Applying NLMM to large-scale data, such as those generated by genome-wide studies, raises several issues related to the assumption of random effects as follows: (i) computation time: it takes a long time to compute the marginal likelihood; (ii) convergence of iterative calculation: an adaptive Gauss-Hermite quadrature is generally used to estimate NLMM; however, iterative calculations may not converge in complex models; and (iii) random-effects misspecification leads to slightly inflated type-I error rates. As an alternative effective approach to resolving these issues, in this article, we propose a generalized estimating equation (GEE) approach for analyzing population PK data. In general, GEE analysis does not account for interindividual variability in PK parameters; therefore, the usual GEE estimators cannot be interpreted straightforwardly, and their validities have not been justified. Here, we propose valid inference methods for using GEE even under conditions of interindividual variability and provide theoretical justifications of the proposed GEE estimators for population PK data. In numerical evaluations by simulations, the proposed GEE approach exhibited high computational speed and stability relative to the NLMM approach. Furthermore, the NLMM analysis was sensitive to the misspecification of the random-effects distribution, and the proposed GEE inference is valid for any distributional form. We provided an illustration by using data from a genome-wide pharmacogenomic study of an anticancer drug.},
author = {Nagashima, Kengo and Sato, Yasunori and Noma, Hisashi and Hamada, Chikuma},
doi = {10.1002/sim.5895},
file = {:Users/solomon/Documents/Mendeley Desktop/Nagashima et al. - 2013 - An efficient and robust method for analyzing population pharmacokinetic data in genome-wide pharmacogenomic st.pdf:pdf},
issn = {10970258},
journal = {Statistics in Medicine},
keywords = {Gene screening,Generalized estimating equations,Genome-wide study,Misspecified models,Population pharmacokinetic data},
number = {27},
pages = {4838--4858},
pmid = {23852468},
title = {{An efficient and robust method for analyzing population pharmacokinetic data in genome-wide pharmacogenomic studies: A generalized estimating equation approach}},
volume = {32},
year = {2013}
}
@article{Tang2016,
abstract = {Aquaporin-4 (AQP4) is a family member of water-channel proteins and is dominantly expressed in the foot process of glial cells surrounding capillaries. The predominant expression at the boundaries between cerebral parenchyma and major fluid compartments suggests the function of aquaporin-4 in water transfer into and out of the brain parenchyma. Accumulating evidences have suggested that the dysregulation of aquaporin-4 relates to the brain edema resulting from a variety of neuro-disorders, such as ischemic or hemorrhagic stroke, trauma, etc. During edema formation in the brain, aquaporin-4 has been shown to contribute to the astrocytic swelling, while in the resolution phase, it has been seen to facilitate the reabsorption of extracellular fluid. In addition, aquaporin-4-deficient mice are protected from cytotoxic edema produced by water intoxication and brain ischemia. However, aquaporin-4 deletion exacerbates vasogenic edema in the brain of different pathological disorders. Recently, our published data showed that the upregulation of aquaporin-4 in astrocytes probably contributes to the transition from cytotoxic edema to vasogenic edema. In this review, apart from the traditional knowledge, we also introduce our latest findings about the effects of mesenchymal stem cells (MSCs) and microRNA-29b on aquaporin-4, which could provide powerful intervention tools targeting aquaporin-4.},
author = {Tang, G and Yang, GY},
doi = {10.3390/ijms17101413},
file = {:Users/solomon/Documents/Mendeley Desktop/Tang, Yang - 2016 - Aquaporin-4 A Potential Therapeutic Target for Cerebral Edema.pdf:pdf},
issn = {1422-0067},
journal = {International journal of molecular sciences},
keywords = {aquaporin-4,brain edema,therapeutic target,water channel},
month = {sep},
number = {10},
pages = {1--11},
pmid = {27690011},
title = {{Aquaporin-4: A Potential Therapeutic Target for Cerebral Edema.}},
url = {http://www.ncbi.nlm.nih.gov/pubmed/27690011 http://www.pubmedcentral.nih.gov/articlerender.fcgi?artid=PMC5085613},
volume = {17},
year = {2016}
}
@article{Narayan1982,
abstract = {The authors have analyzed their experience with intracranial pressure (ICP) monitoring in 207 patients over a 4-year period. Patients with either high-density or low-density lesions on computerized tomography (CT) at admission had a high incidence (53{\%} to 63{\%}) of intracranial hypertension (ICP persistently over 20 mm Hg). In contrast, patients with normal CT scans at admission had a relatively low incidence of ICP elevation (13{\%}). Among these patients, three features were found to be strongly associated with the development of intracranial hypertension: 1) age over 40 years; 2) systolic blood pressure under 90 mm Hg; and 3) motor posturing — unilateral or bilateral. When two or more of these features were noted at admission, the incidence of intracranial hypertension was 60{\%}, as compared to 4{\%} when only one, or none, of these features were present. Thus, the patients at high risk for developing intracranial hypertension after severe head injury are those with abnormal CT scans at admission, and those with normal CT scans who demonstrate two or more of the above-mentioned adverse features. Based on these criteria, only 16{\%} of this series of patients with normal CT scans would have qualified for monitoring. In addition to the three clinical features noted above, multimodality evoked potential (MEP) studies were also found to be strong predictors of ICP elevation in the normal CT scan group, with a 75{\%} incidence of intracranial hypertension in patients with disseminated deficits. There was no statistically significant correlation between the Glasgow Coma Scale score, eye movements, pupillary reaction, hypoxia, or anemia at admission and subsequent ICP elevation in the group with normal CT scans. In this series, an intraventricular catheter was used as the sole monitoring device in 91{\%} of the cases. In the remaining 9{\%}, subarachnoid screws were employed, either alone, or upon failure of the ventriculostomy. While no mortality was directly ascribed to the monitoring process, there was a 7.7{\%} complication rate (infection 6.3{\%} + hemorrhage 1.4{\%}). Eighty-five percent of the infections occurred in patients who had been monitored for 5 days or more, while no infections were noted in those monitored for less than 3 days. Used judiciously, this technique can be valuable in the monitoring and treatment of the brain-injured patient.},
author = {Narayan, RK and Kishore, PR and Becker, DP and Ward, JD and Enas, GG and Greenberg, RP and {Domingues Da Silva}, A and Lipper, MH and Choi, SC and Mayhall, CG and Lutz, HA and Young, H F},
doi = {10.3171/jns.1982.56.5.0650},
file = {:Users/solomon/Documents/Mendeley Desktop/Narayan et al. - 1982 - Intracranial pressure to monitor or not to monitor A review of our experience with severe head injury.pdf:pdf},
isbn = {0022-3085 (Print)$\backslash$r0022-3085 (Linking)},
issn = {0022-3085},
journal = {Journal of neurosurgery},
number = {5},
pages = {650--659},
pmid = {7069477},
title = {{Intracranial pressure: to monitor or not to monitor? A review of our experience with severe head injury.}},
volume = {56},
year = {1982}
}
@article{Dunnenberger2015,
abstract = {Although the field of pharmacogenetics has existed for decades, practioners have been slow to implement pharmacogenetic testing in clinical care. Numerous publications describe the barriers to clinical implementation of pharmacogenetics. Recently, several freely available resources have been developed to help address these barriers. In this review, we discuss current programs that use preemptive genotyping to optimize the pharmacotherapy of patients. Array-based preemptive testing includes a large number of relevant pharmacogenes that impact multiple high-risk drugs. Using a preemptive approach allows genotyping results to be available prior to any prescribing decision so that genomic variation may be considered as an inherent patient characteristic in the planning of therapy. This review describes the common elements among programs that have implemented preemptive genotyping and highlights key processes for implementation, including clinical decision support.},
author = {Dunnenberger, H M and Crews, K R and Hoffman, J M and Caudle, K E and Broeckel, U and Howard, S C and Hunkler, R J and Klein, T E and Evans, W E and Relling, M V},
doi = {10.1146/annurev-pharmtox-010814-124835},
file = {:Users/solomon/Documents/Mendeley Desktop/Dunnenberger et al. - 2015 - Preemptive clinical pharmacogenetics implementation current programs in five US medical centers.pdf:pdf},
isbn = {1545-4304 (Electronic)$\backslash$r0362-1642 (Linking)},
issn = {1545-4304},
journal = {Annu Rev Pharmacol Toxicol},
keywords = {Academic Medical Centers Decision Support Techniqu},
pages = {89--106},
pmid = {25292429},
title = {{Preemptive clinical pharmacogenetics implementation: current programs in five US medical centers}},
url = {http://www.ncbi.nlm.nih.gov/pubmed/25292429},
volume = {55},
year = {2015}
}
@article{Levin1987,
abstract = {To investigate the inter-rater reliability and validity of the Neurobehavioural Rating Scale at various stages of recovery after hospitalisation for closed head injury, we studied 101 head trauma patients who had no antecedent neuropsychiatric disorder. The results demonstrated satisfactory inter-rater reliability and showed that the Neurobehavioural Rating Scale reflects both the severity and chronicity of closed head injury. A principal components analysis revealed four factors which were differentially related to severity of head injury and the presence of a frontal lobe mass lesion. Although our findings provide support for utilising clinical ratings of behaviour to investigate sequelae of head injury, extension of this technique to other settings is necessary to evaluate the distinctiveness of the neurobehavioural profile of closed head injury as compared with other aetiologies of brain damage.},
author = {Levin, H S and High, W M and Goethe, K E and Sisson, R a and Overall, J E and Rhoades, H M and Eisenberg, H M and Kalisky, Z and Gary, H E},
doi = {10.1136/jnnp.50.2.183},
file = {:Users/solomon/Documents/Mendeley Desktop/Levin et al. - 1987 - The neurobehavioural rating scale assessment of the behavioural sequelae of head injury by the clinician.pdf:pdf},
isbn = {0022-3050 (Print)$\backslash$r0022-3050 (Linking)},
issn = {0022-3050},
journal = {Journal of neurology, neurosurgery, and psychiatry},
keywords = {Adolescent,Adult,Brain Concussion,Brain Concussion: diagnosis,Brain Concussion: psychology,Brain Damage, Chronic,Brain Damage, Chronic: diagnosis,Brain Damage, Chronic: psychology,Cerebral Hemorrhage,Cerebral Hemorrhage: diagnosis,Delirium, Dementia, Amnestic, Cognitive Disorders,Delirium, Dementia, Amnestic, Cognitive Disorders:,Female,Follow-Up Studies,Frontal Lobe,Frontal Lobe: injuries,Humans,Male,Middle Aged,Neuropsychological Tests,Psychometrics},
number = {2},
pages = {183--93},
pmid = {3572433},
title = {{The neurobehavioural rating scale: assessment of the behavioural sequelae of head injury by the clinician.}},
url = {http://www.pubmedcentral.nih.gov/articlerender.fcgi?artid=1031490{\&}tool=pmcentrez{\&}rendertype=abstract},
volume = {50},
year = {1987}
}
@article{Klotz1994,
author = {Klotz, Irving M and Scott, Winfield and Johnson, Samuel},
file = {:Users/solomon/Documents/Mendeley Desktop/Klotz, Scott, Johnson - 1994 - Gout and Genius A Chemical Connection.pdf:pdf},
issn = {00219584},
keywords = {General Public,History / Philosophy},
number = {12},
pages = {1015--1018},
title = {{Gout and Genius : A Chemical Connection ?}},
volume = {71},
year = {1994}
}
@article{Ammar2015,
author = {Ammar, Ron and Paton, Tara A. and Torti, Dax and Shlien, Adam and Bader, Gary D.},
doi = {10.12688/f1000research.6037.1},
file = {:Users/solomon/Documents/Mendeley Desktop/Ammar et al. - 2015 - Long read nanopore sequencing for detection of HLA and CYP2D6 variants and haplotypes.pdf:pdf},
issn = {2046-1402},
journal = {F1000Research},
month = {jan},
number = {May},
title = {{Long read nanopore sequencing for detection of HLA and CYP2D6 variants and haplotypes}},
url = {http://f1000research.com/articles/4-17/v1},
year = {2015}
}
@article{Johnson2013a,
abstract = {The University of Florida and Shands Hospital recently launched a genomic medicine program focused on the clinical implementation of pharmacogenetics called the Personalized Medicine Program. We focus on a pre-emptive, chip-based genotyping approach that is cost effective, while providing experience that will be useful as genomic medicine moves towards genome sequence data for patients becoming available. The Personalized Medicine Program includes a regulatory body that is responsible for ensuring that evidence-based examples are moved to clinical implementation, and relies on clinical decision support tools to provide healthcare providers with guidance on use of the genetic information. The pilot implementation was with CYP2C19-clopidogrel and future plans include expansion to additional pharmacogenetic examples, along with aiding in implementation in other health systems across Florida.},
author = {Johnson, Julie a and Elsey, Amanda R and Clare-Salzler, Michael J and Nessl, David and Conlon, Michael and Nelson, David R},
doi = {10.2217/pgs.13.59},
file = {:Users/solomon/Documents/Mendeley Desktop/Johnson et al. - 2013 - Institutional profile University of Florida and Shands Hospital Personalized Medicine Program clinical implement.pdf:pdf},
isbn = {1462-2416},
issn = {1744-8042},
journal = {Pharmacogenomics},
keywords = {Cost-Benefit Analysis,DNA,DNA: methods,Delivery of Health Care,Delivery of Health Care: economics,Delivery of Health Care: organization {\&} administra,Delivery of Health Care: standards,Florida,Genome,Genotype,Health Personnel,Hospitals,Human,Humans,Individualized Medicine,Individualized Medicine: economics,Individualized Medicine: methods,Individualized Medicine: standards,Pharmacogenetics,Pharmacogenetics: economics,Pharmacogenetics: methods,Pharmacogenetics: organization {\&} administration,Pharmacogenetics: standards,Sequence Analysis,Universities},
number = {7},
pages = {723--6},
pmid = {23651020},
title = {{Institutional profile: University of Florida and Shands Hospital Personalized Medicine Program: clinical implementation of pharmacogenetics.}},
url = {http://www.pubmedcentral.nih.gov/articlerender.fcgi?artid=3684075{\&}tool=pmcentrez{\&}rendertype=abstract},
volume = {14},
year = {2013}
}
@article{Macedo2013,
abstract = {OBJECTIVE: To evaluate the relationship between brain damage biomarkers and mortality in the intensive care unit (ICU).$\backslash$n$\backslash$nMETHODS: The sample comprised 70 patients admitted to an ICU. Blood samples were collected from all patients on ICU admission, and levels of S100$\beta$ and neuron-specific enolase (NSE) were determined by ELISA.$\backslash$n$\backslash$nRESULTS: Acute Physiologic and Chronic Health Evaluation (APACHE II) score was associated with mortality, but NSE and S100$\beta$ were not associated with this outcome. In contrast, S100$\beta$ levels were significantly higher in delirious and non-delirious patients who required mechanical ventilation during ICU stay.$\backslash$n$\backslash$nCONCLUSION: Levels of brain biomarkers at the time of ICU admission did not predict mortality in critically ill patients.},
author = {Macedo, Roberta Candal De and Tomasi, Cristiane Damiani and Giombelli, Vinicius Ren{\'{e}} and Alves, Sarah Cascaes and Bristot, Maria De Lourdes Ugioni and Locks, Maria Fernanda Topanotti and Petronilho, Fabr{\'{i}}cia and Grandi, Carmen and Quevedo, Jo{\~{a}}o and Dal-Pizzol, Felipe and Ritter, Cristiane},
doi = {10.1590/1516-4446-2012-0835},
file = {:Users/solomon/Documents/Mendeley Desktop/Macedo et al. - 2013 - Lack of association of S100$\beta$ and neuron-specific enolase with mortality in critically ill patients.pdf:pdf},
isbn = {1516444620},
issn = {1809-452X},
journal = {Revista brasileira de psiquiatria (S{\~{a}}o Paulo, Brazil : 1999)},
keywords = {APACHE,Biological Markers,Biological Markers: blood,Brain Injuries,Brain Injuries: blood,Brain Injuries: mortality,Case-Control Studies,Critical Illness,Critical Illness: mortality,Delirium,Delirium: blood,Enzyme-Linked Immunospot Assay,Female,Humans,Intensive Care Units,Male,Middle Aged,Phosphopyruvate Hydratase,Phosphopyruvate Hydratase: blood,Predictive Value of Tests,Prospective Studies,S100 Calcium Binding Protein beta Subunit,S100 Calcium Binding Protein beta Subunit: blood},
number = {49},
pages = {267--70},
pmid = {24142088},
title = {{Lack of association of S100$\beta$ and neuron-specific enolase with mortality in critically ill patients.}},
url = {http://www.scopus.com/inward/record.url?eid=2-s2.0-84885106215{\&}partnerID=tZOtx3y1},
volume = {35},
year = {2013}
}
@article{Xinhua2006,
author = {Xinhua, D.},
doi = {10.1373/clinchem.2006.077669},
file = {:Users/solomon/Documents/Mendeley Desktop/Xinhua - 2006 - Preparation of Uric Acid Standard Stock Solution.pdf:pdf},
issn = {0009-9147},
journal = {Clinical Chemistry},
month = {sep},
number = {11},
pages = {2117--2118},
pmid = {16873299},
title = {{Preparation of Uric Acid Standard Stock Solution}},
url = {http://www.clinchem.org/cgi/doi/10.1373/clinchem.2006.077669},
volume = {52},
year = {2006}
}
@article{Anderson2012,
author = {Anderson, Jeffrey L and Antman, Elliott M and Bliden, Kevin and Cannon, Christopher P and Gurbel, Paul and Horne, Benjamin D and Steg, P Gabriel and Sabatine, Marc S},
file = {:Users/solomon/Documents/Mendeley Desktop/Anderson et al. - 2012 - Among Patients Treated With Clopidogrel Predominantly for PCI.pdf:pdf},
journal = {October},
number = {16Anderson, J. L., Antman, E. M., Bliden, K., Cannon, C. P., Gurbel, P., Horne, B. D., {\ldots} Sabatine, M. S. (2012). Among Patients Treated With Clopidogrel Predominantly for PCI. October, 304(16), 1821–1830.},
pages = {1821--1830},
title = {{Among Patients Treated With Clopidogrel Predominantly for PCI}},
volume = {304},
year = {2012}
}
@article{Dennis2012,
author = {Dennis, Megan Y. and Nuttle, Xander and Sudmant, Peter H. and Antonacci, Francesca and Graves, Tina A. and Nefedov, Mikhail and Rosenfeld, Jill A. and Sajjadian, Saba and Malig, Maika and Kotkiewicz, Holland and Curry, Cynthia J. and Shafer, Susan and Shaffer, Lisa G. and de Jong, Pieter J. and Wilson, Richard K. and Eichler, Evan E.},
doi = {10.1016/j.cell.2012.03.033},
file = {:Users/solomon/Documents/Mendeley Desktop/Dennis et al. - 2012 - Evolution of Human-Specific Neural SRGAP2 Genes by Incomplete Segmental Duplication.pdf:pdf},
issn = {00928674},
journal = {Cell},
month = {may},
number = {4},
pages = {912--922},
title = {{Evolution of Human-Specific Neural SRGAP2 Genes by Incomplete Segmental Duplication}},
url = {http://linkinghub.elsevier.com/retrieve/pii/S0092867412004618},
volume = {149},
year = {2012}
}
@article{Tsaousides2009,
abstract = {Cognitive rehabilitation refers to a set of interventions that aim to improve a person's ability to perform cognitive tasks by retraining previously learned skills and teaching compensatory strategies. Cognitive rehabilitation begins with a thorough neuropsychological assessment to identify cognitive strengths and weaknesses and the degree of change in cognitive ability following a brain injury. The conclusions of the assessment are used to formulate appropriate treatment plans. Common interventions for improvements in attention, memory, and executive function, as well as the nature of comprehensive programs, which combine treatment modalities, are reviewed. Cognitive rehabilitation is effective for mild-to-severe injuries and beneficial at any time post-injury. Sufficient evidence exists supporting the efficacy and effectiveness of cognitive rehabilitation, which has become the treatment of choice for cognitive impairments and leads to improvements in cognitive and psychosocial functioning.},
archivePrefix = {arXiv},
arxivId = {NIHMS150003},
author = {Tsaousides, Theodore and Gordon, Wayne a},
doi = {10.1002/MSJ},
eprint = {NIHMS150003},
isbn = {1931-7581},
issn = {1931-7581},
journal = {The Mount Sinai journal of medicine, New York},
keywords = {adaptive immunity,autoimmunity,diabetes,genetics in type 1,innate immunity,type},
number = {2},
pages = {173--181},
pmid = {19306374},
title = {{Cognitive rehabilitation following traumatic brain injury: assessment to treatment.}},
volume = {76},
year = {2009}
}
@article{Dardiotis2014a,
abstract = {Abstract Accumulating evidence suggests that the extent of brain injury and the clinical outcome after traumatic brain injury (TBI) are modulated, to some degree, by genetic variants. Aquaporin-4 (AQP4) is the predominant water channel in the central nervous system and plays a critical role in controlling the water content of brain cells and the development of brain edema after TBI. We sought to investigate the influence of the AQP4 gene region on patient outcome after TBI by genotyping tag single nucleotide polymorphisms (SNPs) along AQP4 gene. A total of 363 patients with TBI (19.6{\%} female) were prospectively evaluated. Data including the Glasgow Coma Scale (GCS) scores at admission, the presence of intracranial hemorrhage, and the 6-month Glasgow Outcome Scale (GOS) scores were collected. Seven tag SNPs across the AQP4 gene were identified based on the HapMap data. Using logistic regression analyses, SNPs and haplotypes were tested for associations with 6-month GOS after adjusting for age, GCS score, and sex. Significant associations with TBI outcome were detected for rs3763043 (OR [95{\%} confidence interval (CI)]: 5.15 [1.60-16.5], p=0.006, for recessive model), rs3875089 (OR [95{\%} CI]: 0.18 [0.07-0.50] p=0.0009, for allele difference model), and a common haplotype of AQP4 tag SNPs (OR [95{\%} CI]: 2.94, [1.34-6.36], p=0.0065). AQP4 tag SNPs were not found to influence the initial severity of TBI or the presence of intracranial hemorrhages. In conclusion, the present study provides evidence for possible involvement of genetic variations in AQP4 gene in the functional outcome of patients with TBI.},
author = {Dardiotis, Efthimios and Paterakis, Konstantinos and Tsivgoulis, Georgios and Tsintou, Magdalini and Hadjigeorgiou, Georgios M GF Georgios F GM and Dardioti, Maria and Grigoriadis, Savas and Simeonidou, Constantina and Komnos, Apostolos and Kapsalaki, Eftychia and Fountas, Kostas and Hadjigeorgiou, Georgios M GF Georgios F GM},
doi = {10.1089/neu.2014.3347},
file = {:Users/solomon/Documents/Mendeley Desktop/Dardiotis et al. - 2014 - AQP4 Tag Single Nucleotide Polymorphisms in Patients with Traumatic Brain Injury.pdf:pdf;:Users/solomon/Documents/Mendeley Desktop/Dardiotis et al. - 2014 - AQP4 Tag Single Nucleotide Polymorphisms in Patients with Traumatic Brain Injury(2).pdf:pdf},
issn = {1557-9042},
journal = {Journal of neurotrauma},
keywords = {aqp4,genetics,polymorphism,tag snps,traumatic brain injury},
number = {23},
pages = {1920--1926},
pmid = {24999750},
title = {{AQP4 Tag Single Nucleotide Polymorphisms in Patients with Traumatic Brain Injury.}},
url = {http://www.ncbi.nlm.nih.gov/pubmed/24999750},
volume = {31},
year = {2014}
}
@article{Tu2002,
abstract = {The endoplasmic reticulum (ER) supports disulfide formation through an essential protein relay involving Ero1p and protein disulfide isomerase (PDI). We find that in addition to having a tightly associated flavin adenine dinucleotide (FAD) moiety, yeast Ero1p is highly responsive to small changes in physiological levels of free FAD. This sensitivity underlies the dependence of oxidative protein folding on cellular FAD levels. FAD is synthesized in the cytosol but can readily enter the ER lumen and promote Ero1p-catalyzed oxidation. Ero1p then uses molecular oxygen as its preferred terminal electron acceptor. Thus Ero1p directly couples disulfide formation to the consumption of molecular oxygen, but its activity is modulated by free lumenal FAD levels, potentially linking disulfide formation to a cell's nutritional or metabolic status.},
author = {Tu, Benjamin P. and Weissman, Jonathan S.},
doi = {10.1016/S1097-2765(02)00696-2},
file = {:Users/solomon/Documents/Mendeley Desktop/Tu, Weissman - 2002 - The FAD- and O2-dependent reaction cycle of Ero1-mediated oxidative protein folding in the endoplasmic reticulum.pdf:pdf},
isbn = {1097-2765 (Print)$\backslash$r1097-2765 (Linking)},
issn = {10972765},
journal = {Molecular Cell},
number = {5},
pages = {983--994},
pmid = {12453408},
title = {{The FAD- and O2-dependent reaction cycle of Ero1-mediated oxidative protein folding in the endoplasmic reticulum}},
volume = {10},
year = {2002}
}
@article{Lala-Tabbert2016,
author = {Lala-Tabbert, Neena and Fu, Dechen and Wiper-Bergeron, Nadine},
doi = {10.5966/sctm.2015-0169},
file = {:Users/solomon/Documents/Mendeley Desktop/Lala-Tabbert, Fu, Wiper-Bergeron - 2016 - Induction of CCAATEnhancer-Binding Protein $\beta$ Expression With the Phosphodiesterase Inhibitor.pdf:pdf},
isbn = {2010080432347},
issn = {21576564},
journal = {STEM CELLS Translational Medicine},
keywords = {angiogenesis x chemotaxis,cytokines x,human mesenchymal stem cells,hypoxia x secretome x,x near anoxia and},
month = {apr},
number = {4},
pages = {500--510},
title = {{Induction of CCAAT/Enhancer-Binding Protein $\beta$ Expression With the Phosphodiesterase Inhibitor Isobutylmethylxanthine Improves Myoblast Engraftment Into Dystrophic Muscle}},
url = {http://doi.wiley.com/10.5966/sctm.2014-0180 http://doi.wiley.com/10.5966/sctm.2015-0169},
volume = {5},
year = {2016}
}
@article{Bell2017,
abstract = {5-hydroxytryptamine type 3 (5-HT3 ) receptor antagonists are used in the prevention of chemotherapy- induced, radiation-induced and postoperative nausea and vomiting. CYP2D6 polymorphisms can influence the metabolism of some of these drugs (i.e. ondansetron and tropisetron) thereby affecting drug efficacy. We summarize evidence from the published literature supporting these associations and provide therapeutic recommendations for ondansetron and tropisetron based on CYP2D6 genotype (updates at www.pharmgkb.org). This article is protected by copyright. All rights reserved.},
author = {Bell, G C and Caudle, K E and Whirl-Carrillo, M and Gordon, R J and Hikino, K and Prows, C A and Gaedigk, A and Agundez, J A G and Sadhasivam, S and Klein, T E and Schwab, M},
doi = {10.1002/cpt.598},
file = {:Users/solomon/Documents/Mendeley Desktop/Bell et al. - 2017 - Clinical Pharmacogenetics Implementation Consortium (CPIC) guideline for CYP2D6 genotype and use of ondansetron and.pdf:pdf},
issn = {15326535},
journal = {Clin. Pharmacol. Ther.},
number = {2},
pages = {213--218},
pmid = {28002639},
title = {{Clinical Pharmacogenetics Implementation Consortium (CPIC) guideline for CYP2D6 genotype and use of ondansetron and tropisetron}},
volume = {102},
year = {2017}
}
@article{Alluri2015,
abstract = {Traumatic brain injury is a serious cause of morbidity and mortality worldwide. After traumatic brain injury, the blood-brain barrier, the protective barrier between the brain and the intravascular compartment, becomes dysfunctional, leading to leakage of proteins, fluid, and transmigration of immune cells. As this leakage has profound clinical implications, including edema formation, elevated intracranial pressure and decreased perfusion pressure, much interest has been paid to better understanding the mechanisms responsible for these events. Various molecular pathways and numerous mediators have been found to be involved in the intricate process of regulating blood-brain barrier permeability following traumatic brain injury. This review provides an update to the existing knowledge about the various pathophysiological pathways and advancements in the field of blood-brain barrier dysfunction and hyperpermeability following traumatic brain injury, including the role of various tight junction proteins involved in blood-brain barrier integrity and regulation. We also address pitfalls of existing systems and propose strategies to improve the various debilitating functional deficits caused by this progressive epidemic.},
author = {Alluri, Himakarnika and Wiggins-Dohlvik, Katie and Davis, Matthew L. and Huang, Jason H. and Tharakan, Binu},
doi = {10.1007/s11011-015-9651-7},
file = {:Users/solomon/Documents/Mendeley Desktop/Alluri et al. - 2015 - Blood–brain barrier dysfunction following traumatic brain injury.pdf:pdf},
issn = {15737365},
journal = {Metabolic Brain Disease},
keywords = {Blood–brain barrier,Brain edema,Central nervous system trauma,Traumatic brain injury,Vascular permeability},
number = {5},
pages = {1093--1104},
pmid = {25624154},
title = {{Blood–brain barrier dysfunction following traumatic brain injury}},
volume = {30},
year = {2015}
}
@article{Kurowski2016,
author = {Kurowski, BG and Backeljauw, B and Zang, H and Zhang, N and Martin, LJ and Pilipenko, V and Yeates, K and Taylor, HG and Wade, S},
doi = {10.1097/HTR.0000000000000162},
file = {:Users/solomon/Documents/Mendeley Desktop/Kurowski et al. - 2016 - Influence of Catechol-O-methyltransferase on Executive Functioning Longitudinally After Early Childhood Traumat.pdf:pdf},
issn = {1550-509X (Electronic) 0885-9701 (Linking)},
journal = {J Head Trauma Rehabil},
number = {3},
pages = {E1--9},
title = {{Influence of Catechol-O-methyltransferase on Executive Functioning Longitudinally After Early Childhood Traumatic Brain Injury: Preliminary Findings}},
type = {Journal Article},
url = {https://www.ncbi.nlm.nih.gov/pubmed/26394291},
volume = {31},
year = {2016}
}
@article{Bauer2016,
abstract = {ABCB1 and ABCG2 work together at the blood-brain barrier (BBB) to limit brain distribution of dual ABCB1/ABCG2 substrates. In this pilot study we used PET to assess brain distribution of two model ABCB1/ABCG2 substrates ([(11) C]elacridar and [(11) C]tariquidar) in healthy subjects without (c.421CC) or with (c.421CA) the ABCG2 single-nucleotide polymorphism (SNP) c.421C{\textgreater}A. Subjects underwent PET scans under conditions when ABCB1 and ABCG2 were functional and during ABCB1 inhibition with high-dose tariquidar. In contrast to the ABCB1-selective substrate (R)-[(11) C]verapamil, [(11) C]elacridar and [(11) C]tariquidar showed only moderate increases in brain distribution during ABCB1 inhibition. This provides evidence for a functional interplay between ABCB1 and ABCG2 at the human BBB and suggests that both ABCB1 and ABCG2 need to be inhibited to achieve substantial increases in brain distribution of dual ABCB1/ABCG2 substrates. During ABCB1 inhibition c.421CA subjects had significantly higher increases in [(11) C]tariquidar brain distribution than c.421CC subjects pointing to impaired cerebral ABCG2 function. This article is protected by copyright. All rights reserved.},
author = {Bauer, M. and R{\"{o}}mermann, K. and Karch, R. and Wulkersdorfer, B. and Stanek, J. and Philippe, C. and Maier-Salamon, A. and Haslacher, H. and Jungbauer, C. and Wadsak, W. and J{\"{a}}ger, W. and L{\"{o}}scher, W. and Hacker, M. and Zeitlinger, M. and Langer, O.},
doi = {10.1002/cpt.362},
file = {:Users/solomon/Documents/Mendeley Desktop/Bauer et al. - 2016 - Pilot PET Study to Assess the Functional Interplay Between ABCB1 and ABCG2 at the Human Blood–Brain Barrier.pdf:pdf},
isbn = {0009-9236},
issn = {15326535},
journal = {Clinical Pharmacology and Therapeutics},
pages = {131--141},
pmid = {26940368},
title = {{Pilot PET Study to Assess the Functional Interplay Between ABCB1 and ABCG2 at the Human Blood–Brain Barrier}},
year = {2016}
}
@article{Pike2001,
abstract = {Although a number of increased CSF proteins have been correlated with brain damage and outcome after traumatic brain injury (TBI), a major limitation of currently tested biomarkers is a lack of specificity for defining neuropathological cascades. Identification of surrogate biomarkers that are elevated in CSF in response to brain injury and that offer insight into one or more pathological neurochemical events will provide critical information for appropriate administration of therapeutic compounds for treatment of TBI patients. Non-erythroid alpha II-spectrin is a cytoskeletal protein that is a substrate of both calpain and caspase-3 cysteine proteases. As we have previously demonstrated, cleavage of alpha II-spectrin by calpain and caspase-3 results in accumulation of protease-specific spectrin breakdown products (SBDPs) that can be used to monitor the magnitude and temporal duration of protease activation. However, accumulation of alpha II-spectrin and alpha II-SBDPs in CSF after TBI has never been examined. Following a moderate level (2.0 mm) of controlled cortical impact TBI in rodents, native alpha II-spectrin protein was decreased in brain tissue and increased in CSF from 24 h to 72 h after injury. In addition, calpain-specific SBDPs were observed to increase in both brain and CSF after injury. Increases in the calpain-specific 145 kDa SBDP in CSF were 244{\%}, 530{\%} and 665{\%} of sham-injured control animals at 24 h, 48 h and 72 h after TBI, respectively. The caspase-3-specific SBDP was observed to increase in CSF in some animals but to a lesser degree. Importantly, levels of these proteins were undetectable in CSF of uninjured control rats. These results indicate that detection of alpha II-spectrin and alpha II-SBDPs is a powerful discriminator of outcome and protease activation after TBI. In accord with our previous studies, results also indicate that calpain may be a more important effector of cell death after moderate TBI than caspase-3.},
author = {Pike, B R and Flint, J and Dutta, S and Johnson, E and Wang, K K and Hayes, R L},
doi = {10.1046/j.1471-4159.2001.00510.x},
file = {:Users/solomon/Documents/Mendeley Desktop/Pike et al. - 2001 - Accumulation of non-erythroid alpha II-spectrin and calpain-cleaved alpha II-spectrin breakdown products in cerebro.pdf:pdf},
isbn = {0022-3042 (Print)$\backslash$r0022-3042 (Linking)},
issn = {0022-3042},
journal = {Journal of neurochemistry},
keywords = {1297,1306,2001,78,brain injury,calpain,caspase-3,cell death,cerebrospinal,in the united,j,neurochem,spectrin,tbi,the incidence of traumatic,traumatic brain injury,uid},
number = {6},
pages = {1297--1306},
pmid = {11579138},
title = {{Accumulation of non-erythroid alpha II-spectrin and calpain-cleaved alpha II-spectrin breakdown products in cerebrospinal fluid after traumatic brain injury in rats.}},
volume = {78},
year = {2001}
}
@article{Coffman1997,
abstract = {A human UDP-glucuronosyltransferase (UGT) catalyzing the glucuronidation of morphine has been identified. A full length cDNA was isolated from a human liver cDNA library and found to be identical to the UGT2B7 form having a tyrosine at position 288. This cDNA was transfected into HK 293 cells, and stable expression was achieved. Cell homogenates and membrane preparations from HK 293 cells expressing UGT2B7 catalyzed the glucuronidation of morphine and other clinically significant opioid agonists, antagonists, and partial agonists. UGT2B7 catalyzed morphine glucuronidation at the 3- and 6-hydroxy positions and also mediated the formation of codeine-6-glucuronide from codeine. This represents the first demonstration of a UGT capable of catalyzing the glucuronidation of both the 3- and 6-positions of opioids. Since humans excrete morphine-3-glucuronide and morphine-6-glucuronide after morphine administration, it is likely that UGT2B7 is a major isoform in humans responsible for the metabolism of this important drug and its surrogates.},
author = {Coffman, B L and Rios, G R and King, C D and Tephly, T R},
doi = {10.1016/j.bmc.2005.05.002},
file = {:Users/solomon/Documents/Mendeley Desktop/Coffman et al. - 1997 - Human UGT2B7 catalyzes morphine glucuronidation.pdf:pdf},
isbn = {0968-0896},
issn = {0090-9556},
journal = {Drug metabolism and disposition: the biological fate of chemicals},
month = {jan},
number = {1},
pages = {1--4},
pmid = {9010622},
title = {{Human UGT2B7 catalyzes morphine glucuronidation.}},
url = {http://linkinghub.elsevier.com/retrieve/pii/S0024320597011065 http://www.ncbi.nlm.nih.gov/pubmed/15951183 http://www.ncbi.nlm.nih.gov/pubmed/9010622},
volume = {25},
year = {1997}
}
@article{Ogura2015,
abstract = {The prevalence of hyperuricemia/gout increases with aging. However, the effect of aging on function for excretion of uric acid to out of the body has not been clarified. We found that ileal uric acid clearance in middle-aged rats (11-12 months) was decreased compared with that in young rats (2 months). In middle-aged rats, xanthine oxidase (XO) activity in the ileum was significantly higher than that in young rats. Inosine-induced reactive oxygen species (ROS), which are derived from XO, also decreased ileal uric acid clearance. ROS derived from XO decreased the active homodimer level of breast cancer resistance protein (BCRP), which is a uric acid efflux transporter, in the ileum. Pre-administration of allopurinol recovered the BCRP homodimer level, resulting in the recovering ileal uric acid clearance. Moreover, we investigated the effects of ROS derived from XO on BCRP homodimer level directly in Caco-2 cells using hypoxanthine. Treatment with hypoxanthine decreased BCRP homodimer level. Treatment with hypoxanthine induced mitochondrial dysfunction, suggesting that the decreasing BCRP homodimer level might be caused by mitochondrial dysfunction. In conclusion, ROS derived from XO decrease BCRP homodimer level, resulting in suppression of function for uric acid excretion to the ileal lumen. ROS derived from XO may cause the suppression of function of the ileum for the excretion of uric acid with aging. The results of our study provide a new insight into the causes of increasing hyperuricemia/gout prevalence with aging.},
author = {Ogura, Jiro and Kuwayama, Kaori and Sasaki, Shunichi and Kaneko, Chihiro and Koizumi, Takahiro and Yabe, Keisuke and Tsujimoto, Takashi and Takeno, Reiko and Takaya, Atsushi and Kobayashi, Masaki and Yamaguchi, Hiroaki and Iseki, Ken},
doi = {10.1016/j.bcp.2015.06.021},
file = {:Users/solomon/Documents/Mendeley Desktop/Ogura et al. - 2015 - Reactive oxygen species derived from xanthine oxidase interrupt dimerization of breast cancer resistance protein,.pdf:pdf},
isbn = {0006-2952},
issn = {18732968},
journal = {Biochemical Pharmacology},
keywords = {Aging,Breast cancer resistance protein,Hyperuricemia,Reactive oxygen species,Uric acid excretion},
number = {1},
pages = {89--98},
pmid = {26119820},
publisher = {Elsevier Inc.},
title = {{Reactive oxygen species derived from xanthine oxidase interrupt dimerization of breast cancer resistance protein, resulting in suppression of uric acid excretion to the intestinal lumen}},
url = {http://dx.doi.org/10.1016/j.bcp.2015.06.021},
volume = {97},
year = {2015}
}
@article{Sanchez-Aguilar2013,
abstract = {{\textcopyright}AANS, 2013 T raumaTic brain injury continues to be a worldwide health problem with devastating familial, social, and economic consequences. 7 The recovery phase includes many societal difficulties and loss of productiv-ity. The available interventions do not clearly improve long-term functional outcomes after TBI. Thus, the man-agement of TBI remains largely supportive, directed to-ward control of cerebral edema and intracranial hyperten-sion. 24 The structural analogs of b-hydroxy-b-methylglu-taryl coenzyme A (HMG CoA) reductase (statins) have been assessed in the treatment of aneurysmal subarach-noid hemorrhage and spontaneous intracerebral hemor-rhage in humans (for a review, see the article by Tapia-P{\'{e}}rez et al. Object. The favorable effect of statin treatment after traumatic brain injury (TBI) has been shown in animal stud-ies and is probably true in humans as well. The objective of this study was to determine whether acute statin treatment following TBI could reduce inflammatory cytokines and improve functional outcomes in humans. Methods. The authors performed a double-blind randomized clinical trial in patients with moderate to severe TBI. Exclusion criteria were as follows: prior severe disability; use of modifiers of statin metabolism; multisystem trauma; prior use of mannitol, barbiturates, corticosteroids, or calcium channel blockers; isolated brainstem lesions; allergy to statins; previous hepatopathy or myopathy; previous treatment at another clinic; and pregnancy. Patients were randomly selected to receive 20 mg of rosuvastatin or placebo for 10 days. The main goal was to determine the effect of rosuvastatin on plasma levels of tumor necrosis factor–a, interleukin (IL)–1b, IL-6, and IL-10 after 72 hours of TBI. Amnesia, disorientation, and disability were assessed 3 and 6 months after TBI. Results. Thirty-six patients were analyzed according to intention-to-treat analysis; 19 patients received rosuva-statin and 17 received placebo. The best-fit mixed model showed a significant effect of rosuvastatin on the reduction of tumor necrosis factor–a levels (p = 0.004). Rosuvastatin treatment did not appear to affect the levels of IL-1b, IL-6, and IL-10. The treatment was associated with a reduction in disability scores (p = 0.03), indicating a favorable functional outcome. Life-threatening adverse effects were not observed. Conclusions. The authors' data suggest that statins may induce an antiinflammatory effect and may promote recovery after TBI. The role of statins in TBI therapy should be confirmed in larger clinical trials. Clinical trial reg-istration no.: NCT00990028. Key WorDS • traumatic brain injury • rosuvastatin • cytokine • disability 669 Abbreviations used in this paper: ALT = alanine transaminase; AST = aspartate transaminase; CK = creatine kinase; DRS = Dis-ability Rating Scale; GCS = Glasgow Coma Scale; GOAT = Galves-ton Orientation Amnesia Test; IL = interleukin; TBI = traumatic brain injury; TCDB = Traumatic Coma Data Bank; TNFa = tumor necrosis factor–a. * Drs. S{\'{a}}nchez-Aguilar and Tapia-P{\'{e}}rez contributed equally to this work.},
