## Supplementary material for "Temporal Profile of Transporter mRNA Expression in the Brain after Traumatic Brain Injury in Developing Rats": Archive containing raw data, r functions, and manuscript (Rmd): TBI_Gene_Expression_Submission.pdf

### Running Title Page

**Running Title:** Pediatric Neurotrauma Associated Transporter Expression

**Corresponding Author:**
Philip E. Empey, PharmD, PhD
Department of Pharmacy and Therapeutics, School of Pharmacy
University of Pittsburgh
205 Salk Pavilion
335 Sutherland Drive
Pittsburgh, PA 15213


#### Figure Legends

**Figure 1**

Baseline transporter expression (PND-17) in kidney, liver, choroid plexus, hippocampus, and cortex. Error bars show standard deviation.

**Figure 2**

Ontogeny of transporters in cortex, hippocampus, choroid plexus, liver, and kidney. Point color indicates statistical significance (FDR < 0.05).

**Figure 1**

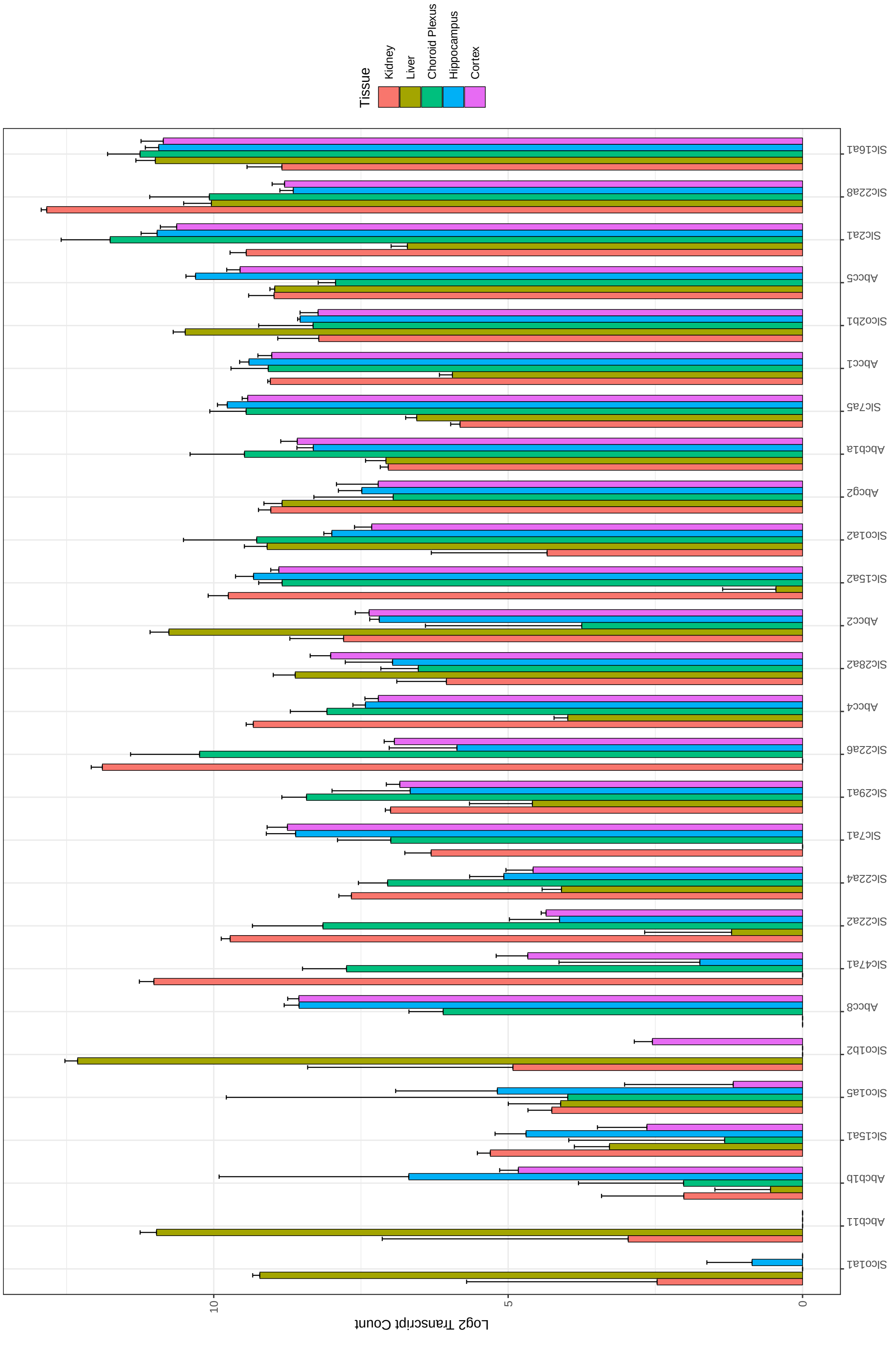
Tissue
Kidney
Liver
Choroid Plexus
Hippocampus
Cortex
Slc16a1
Slc22a8
Slc2a1
Abcc5
Slco2b1
Abcc1
Slc7a5
Abcb1a
Abcg2
Slco1a2
Slc15a2
Abcc2
Slc28a2
Abcc4
Slc22a6
Slc29a1
Slc7a1
Slc22a4
Slc22a2
Slc47a1
Abcc8
Slco1b2
Slco1a5
Slc15a1
Abcb1b
Abcb11
Slco1a1
10
5
0
Log2 Transcript Count

#### Figure 2

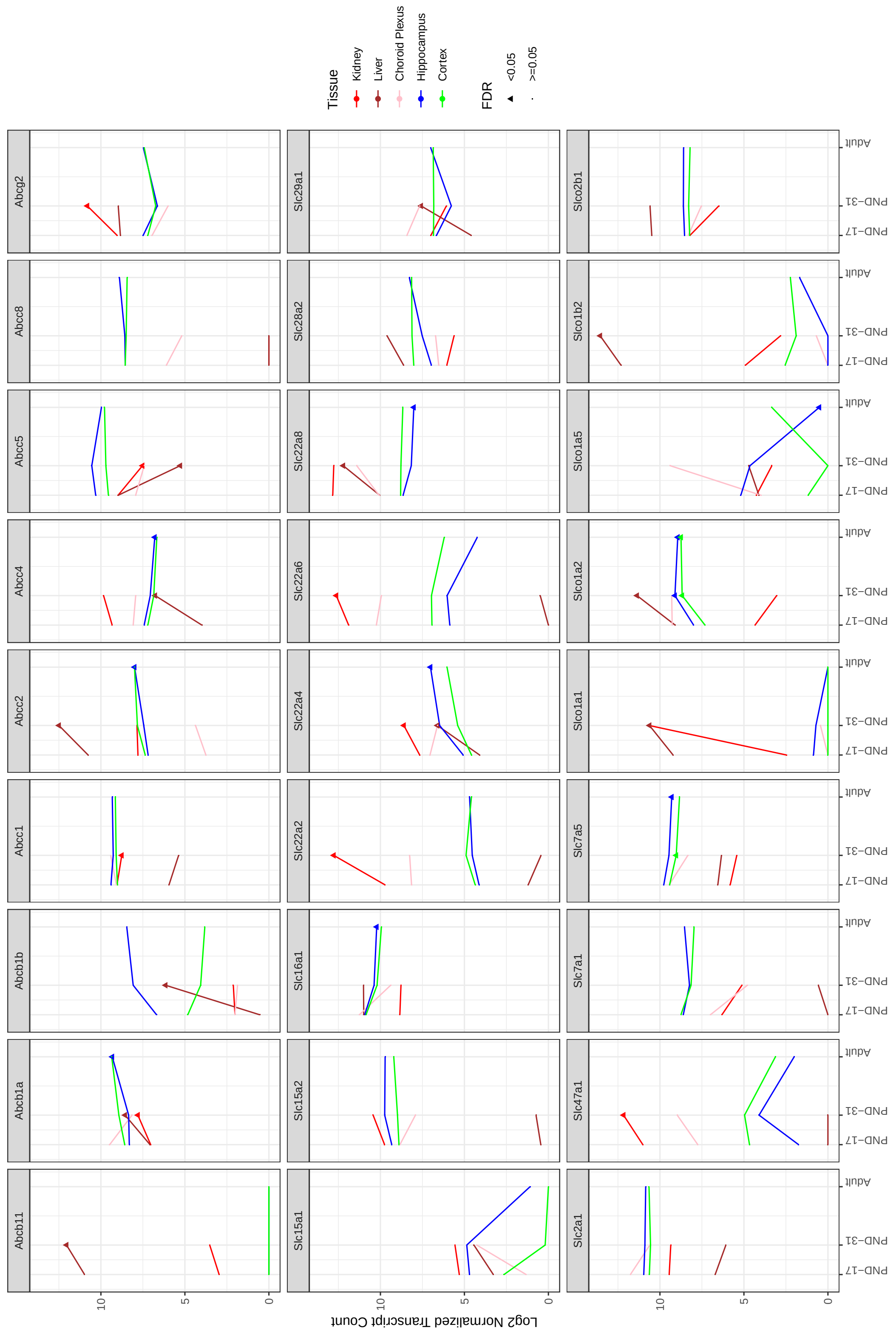

Abcb11
Abcb1a
Abcb1b
Abcc1
Abcc2
Abcc4
Abcc5
Abcc8
Abcg2
Slc15a1
Slc15a2
Slc16a1
Slc22a2
Slc22a4
Slc22a6
Slc22a8
Slc28a2
Slc29a1
Slc2a1
Slc47a1
Slc7a1
Slc7a5
Slco1a1
Slco1a2
Slco1a5
Slco1b2
Slco2b1
Log2 Normalized Transcript Count
10
5
0
PND-17
PND-31
Adult
Tissue
Kidney
Liver
Choroid Plexus
Hippocampus
Cortex
FDR
<0.05
>=0.05

**Table 1** Genes Included on Expression Panel

| Rat Gene | Human Gene | Common Name |
|---|---|---|
| | Transporters | |
| *Abcb11* | *ABCB11* | BSEP |
| *Abcb1a* | *ABCB1* | MDR1/PGP |
| *Abcb1b* | *ABCB1* | MDR1/PGP |
| *Abcc1* | *ABCC1* | MRP1 |
| *Abcc2* | *ABCC2* | MRP2 |
| *Abcc4* | *ABCC4* | MRP4 |
| *Abcc5* | *ABCC5* | MRP5 |
| *Abcc8* | *ABCC8* | SUR1 |
| *Abcg2* | *ABCG2* | BCRP |
| *Slc15a1* | *SLC15A1* | PEPT1 |
| *Slc15a2* | *SLC15A2* | PEPT2 |
| *Slc16a1* | *SLC16A1* | MCT1 |
| *Slc22a2* | *SLC22A2* | OCT2 |
| *Slc22a4* | *SLC22A4* | OCTN1 |
| *Slc22a6* | *SLC22A6* | OAT1 |
| *Slc22a8* | *SLC22A8* | OAT3 |
| *Slc28a2* | *SLC28A2* | CNT2 |
| *Slc29a1* | *SLC29A1* | ENT1 |
| *Slc2a1* | *SLC2A1* | GLUT1 |
| *Slc47a1* | *SLC47A1* | MATE1 |
| *Slc7a1* | *SLC7A1* | CAT1 |
| *Slc7a5* | *SLC7A5* | LAT1 |
| *Slco1a1* | — | OATP |
| *Slco1a2* | — | OATP |
| *Slco1a5* | *SLCO1A2* | OATP1A2 |
| *Slco1b2* | *SLCO1B3* | OATP8 |
| *Slco2b1* | *SLCO2B1* | OATP2B1 |
| | Transcription Factors | |
| *Hif1a* | *HIF1-A* | HIF1-A |
| *Nfe2l2* | *NFE2L2* | NRF2 |
| *Il-6* | *IL-6* | IL-6 |
| *Nr1i2* | *NR1I2* | PXR |
| | Biomarkers | |
| *Gfap* | *GFAP* | GFAP |
| *Vim* | *VIM* | VIM |
| *Icam1* | *ICAM1* | CD54 |
| *Ngb* | *NGB* | NGB |
| *Edn1* | *EDN1* | ET1 |

**Table 2** Expression Changes in the Ipsilateral Cortex Post TBI

| | | 3 Hours | | | 12 Hours | | | 24 Hours | | | 72 Hours | | | 1 Week | | | 2 Weeks | | |
|---|---|---|---|---|---|---|---|---|---|---|---|---|---|---|---|---|---|---|---|
| | EML* | FC | SD | FDR | FC | SD | FDR | FC | SD | FDR | FC | SD | FDR | FC | SD | FDR | FC | SD | FDR |
| *Abcb11* | ↔ ↑ ↑ | 0.000 | 0.000 | 1.000 | 0.170 | 0.481 | 0.459 | 1.149 | 1.099 | 0.066 | 1.836 | 1.306 | **0.018** | 1.357 | 1.215 | **0.041** | 0.769 | 1.157 | 0.189 |
| *Abcb1a* | ↔ ↔ ↑ | -0.187 | 0.391 | 0.559 | -0.324 | 0.378 | 0.110 | -0.145 | 0.251 | 0.645 | 0.242 | 0.142 | 0.110 | 0.415 | 0.137 | **0.030** | 0.275 | 0.215 | 0.281 |
| *Abcb1b* | ↑ ↑ ↔ | 0.820 | 0.218 | **0.020** | 0.834 | 0.535 | **0.019** | 0.507 | 0.231 | **0.007** | 2.077 | 0.345 | **0.000** | 1.550 | 0.631 | 0.058 | 0.954 | 0.615 | 0.241 |
| *Abcc1* | ↑ ↑ ↔ | -0.180 | 0.146 | 0.320 | 0.090 | 0.205 | 0.388 | 0.321 | 0.089 | **0.002** | 0.421 | 0.133 | **0.012** | 0.269 | 0.246 | 0.067 | 0.189 | 0.217 | 0.393 |
| *Abcc2* | ↓ ↔ ↔ | -0.384 | 0.163 | **0.004** | -0.505 | 0.481 | 0.060 | -0.561 | 0.372 | **0.036** | 0.050 | 0.146 | 0.658 | 0.051 | 0.048 | 0.616 | -0.224 | 0.292 | 0.280 |
| *Abcc4* | ↔ ↑ ↑ | 0.186 | 0.260 | 0.385 | 0.156 | 0.322 | 0.362 | 0.260 | 0.167 | 0.198 | 0.775 | 0.028 | **0.004** | 0.597 | 0.137 | **0.004** | 0.474 | 0.268 | 0.259 |
| *Abcc5* | ↔ ↔ ↔ | -0.098 | 0.221 | 0.634 | 0.112 | 0.153 | 0.567 | 0.214 | 0.201 | 0.225 | 0.298 | 0.110 | 0.058 | 0.164 | 0.187 | 0.162 | -0.009 | 0.295 | 0.965 |
| *Abcc8* | ↓ ↓ ↔ | 0.267 | 0.087 | **0.037** | 0.025 | 0.281 | 0.898 | -0.333 | 0.087 | **0.006** | -0.231 | 0.156 | **0.018** | -0.170 | 0.134 | 0.315 | -0.284 | 0.258 | 0.356 |
| *Abcg2* | ↔ ↔ ↔ | -0.507 | 0.483 | 0.315 | -0.618 | 0.147 | 0.152 | -0.320 | 0.412 | 0.367 | -0.267 | 0.212 | 0.205 | 0.074 | 0.351 | 0.816 | -0.007 | 0.233 | 0.973 |
| *Edn1* | ↑ ↔ ↔ | 1.166 | 0.290 | **0.003** | 1.232 | 0.377 | **0.001** | 0.823 | 0.267 | 0.093 | 0.300 | 0.388 | 0.385 | 0.611 | 0.494 | 0.203 | -0.064 | 0.208 | 0.616 |
| *Gfap* | ↑ ↑ ↑ | 0.411 | 0.177 | 0.088 | 1.768 | 0.232 | **0.000** | 2.156 | 0.630 | **0.019** | 2.699 | 0.278 | **0.001** | 2.495 | 0.418 | **0.004** | 1.834 | 0.459 | **0.049** |
| *Hif1a* | ↑ ↑ ↔ | 0.247 | 0.067 | **0.040** | 0.753 | 0.277 | **0.001** | 1.028 | 0.264 | **0.000** | 0.277 | 0.146 | **0.013** | 0.234 | 0.094 | 0.106 | 0.276 | 0.314 | 0.227 |
| *Icam1* | ↑ ↑ ↑ | 1.951 | 0.549 | **0.018** | 2.123 | 0.172 | **0.000** | 1.784 | 0.938 | **0.005** | 1.430 | 0.257 | **0.001** | 1.111 | 0.279 | **0.025** | 0.869 | 0.924 | 0.203 |
| *Il6* | ↑ ↔ ↔ | 4.377 | 1.694 | **0.038** | 4.996 | 0.967 | **0.000** | 2.077 | 1.849 | **0.041** | 1.065 | 1.326 | 0.115 | 1.230 | 1.245 | 0.064 | 0.872 | 2.022 | 0.372 |
| *Nfe2l2* | ↑ ↑ ↑ | 0.677 | 0.091 | **0.003** | 0.988 | 0.115 | **0.003** | 0.456 | 0.056 | **0.028** | 1.179 | 0.177 | **0.002** | 1.019 | 0.310 | **0.001** | 0.791 | 0.416 | **0.018** |
| *Ngb* | ↓ ↔ ↔ | -0.139 | 0.251 | 0.448 | -0.841 | 0.449 | **0.005** | -1.588 | 0.172 | **0.001** | 0.143 | 0.271 | 0.580 | 0.096 | 0.562 | 0.835 | -0.379 | 0.259 | 0.237 |
| *Nr1i2* | ↔ ↔ ↔ | 0.000 | 0.000 | 1.000 | 0.105 | 0.298 | 0.459 | 0.000 | 0.000 | 1.000 | 0.268 | 0.791 | 0.567 | 0.085 | 0.535 | 0.816 | 0.306 | 0.866 | 0.459 |
| *Slc15a1* | ↔ ↔ ↔ | 0.241 | 0.486 | 0.733 | 0.058 | 0.315 | 0.915 | -0.172 | 1.357 | 0.887 | -1.023 | 1.293 | 0.165 | -1.224 | 0.860 | 0.265 | 0.086 | 0.906 | 0.915 |
| *Slc15a2* | ↓ ↑ ↑ | -0.190 | 0.243 | 0.328 | -0.932 | 0.247 | **0.000** | -1.155 | 0.382 | **0.001** | 0.677 | 0.303 | **0.004** | 0.809 | 0.182 | **0.010** | 0.497 | 0.109 | **0.023** |
| *Slc16a1* | ↓ ↑ ↑ | -0.042 | 0.129 | 0.772 | -0.448 | 0.186 | **0.003** | -0.296 | 0.132 | **0.025** | 0.524 | 0.112 | **0.034** | 0.485 | 0.173 | **0.003** | 0.586 | 0.317 | **0.046** |
| *Slc22a2* | ↔ ↔ ↔ | -0.145 | 0.399 | 0.742 | -0.524 | 0.671 | 0.190 | 0.027 | 0.346 | 0.925 | 0.248 | 0.214 | 0.469 | 0.456 | 0.082 | 0.203 | 0.021 | 0.405 | 0.965 |
| *Slc22a4* | ↑ ↑ ↑ | 0.711 | 0.833 | 0.356 | 1.200 | 0.149 | **0.016** | 1.416 | 0.413 | **0.007** | 2.316 | 0.340 | **0.003** | 1.513 | 0.250 | **0.004** | 1.687 | 1.022 | 0.100 |
| *Slc22a6* | ↓ ↔ ↔ | -0.525 | 0.535 | 0.424 | -2.825 | 1.898 | **0.015** | -1.602 | 0.913 | **0.008** | -0.060 | 0.216 | 0.894 | 1.452 | 0.770 | 0.088 | 0.505 | 0.874 | 0.559 |
| *Slc22a8* | ↓ ↔ ↔ | -0.208 | 0.422 | 0.540 | -1.157 | 0.448 | **0.001** | -1.493 | 0.288 | **0.000** | -0.428 | 0.246 | 0.051 | 0.205 | 0.111 | 0.540 | -0.141 | 0.239 | 0.695 |
| *Slc28a2* | ↓ ↔ ↔ | -0.543 | 0.070 | **0.028** | -0.634 | 0.145 | **0.018** | -0.066 | 0.458 | 0.888 | 0.508 | 0.304 | 0.110 | -0.080 | 0.357 | 0.645 | -0.290 | 0.471 | 0.459 |
| *Slc29a1* | ↓ ↑ ↔ | -0.261 | 0.115 | 0.115 | -0.678 | 0.160 | **0.001** | 0.248 | 0.185 | 0.189 | 0.591 | 0.162 | **0.000** | 0.378 | 0.199 | 0.104 | 0.252 | 0.437 | 0.461 |
| *Slc2a1* | ↑ ↔ ↔ | 0.394 | 0.201 | 0.084 | 0.789 | 0.305 | **0.001** | 0.705 | 0.546 | **0.028** | 0.085 | 0.167 | 0.616 | 0.301 | 0.156 | 0.131 | 0.392 | 0.259 | 0.067 |
| *Slc47a1* | ↓ ↔ ↔ | -2.303 | 1.935 | **0.034** | -0.690 | 0.447 | 0.444 | -1.280 | 1.193 | 0.201 | -0.570 | 0.479 | 0.196 | 0.921 | 0.397 | 0.259 | 0.435 | 1.015 | 0.634 |
| *Slc7a1* | ↔ ↔ ↔ | 0.076 | 0.095 | 0.676 | -0.053 | 0.135 | 0.686 | -0.157 | 0.136 | 0.259 | -0.064 | 0.054 | 0.726 | -0.007 | 0.048 | 0.965 | 0.061 | 0.051 | 0.695 |
| *Slc7a5* | ↑ ↔ ↑ | 0.229 | 0.060 | 0.160 | 0.672 | 0.128 | **0.001** | 0.599 | 0.104 | **0.000** | 0.210 | 0.132 | 0.066 | 0.398 | 0.053 | **0.001** | 0.459 | 0.055 | **0.012** |
| *Slco1a1* | ↔ ↔ ↔ | 0.000 | 0.000 | 1.000 | 0.000 | 0.000 | 1.000 | 0.000 | 0.000 | 1.000 | 0.000 | 0.000 | 1.000 | 0.000 | 0.000 | 1.000 | 0.000 | 0.000 | 1.000 |
| *Slco1a2* | ↓ ↔ ↔ | -0.479 | 0.373 | 0.266 | -0.889 | 0.412 | **0.004** | -0.647 | 0.193 | 0.189 | -0.127 | 0.211 | 0.487 | 0.533 | 0.370 | 0.152 | 0.308 | 0.324 | 0.283 |
| *Slco1a5* | ↔ ↔ ↔ | -0.696 | 1.885 | 0.676 | -1.346 | 1.555 | 0.281 | 0.105 | 0.381 | 0.600 | 0.699 | 1.604 | 0.367 | -0.516 | 1.411 | 0.793 | -0.055 | 0.680 | 0.979 |
| *Slco1b2* | ↔ ↔ ↓ | 0.327 | 0.426 | 0.751 | -0.517 | 0.198 | 0.540 | 0.318 | 1.147 | 0.793 | 0.254 | 0.605 | 0.759 | -1.715 | 0.469 | **0.013** | -0.155 | 0.629 | 0.888 |
| *Slco2b1* | ↓ ↑ ↑ | -0.268 | 0.326 | 0.362 | -0.637 | 0.238 | **0.001** | -0.490 | 0.177 | **0.002** | 0.716 | 0.316 | **0.004** | 1.203 | 0.146 | **0.001** | 0.712 | 0.203 | 0.127 |
| *Vim* | ↑ ↑ ↑ | 0.216 | 0.203 | 0.446 | 1.130 | 0.375 | **0.001** | 2.230 | 0.091 | **0.004** | 3.459 | 0.412 | **0.002** | 2.607 | 0.400 | **0.000** | 1.689 | 0.277 | 0.115 |

EML*: Early (3-24 hours), Mid (72 hours), Late (7-14 Days); ↑: expression increased, ↓: expression decreased, ↔: expression did not change significantly

**Table 3** Expression Changes in the Ipsilateral Hippocampus Post TBI

| | | 3 Hours | | | 12 Hours | | | 24 Hours | | | 72 Hours | | | 1 Week | | | 2 Weeks | | |
|---|---|---|---|---|---|---|---|---|---|---|---|---|---|---|---|---|---|---|---|
| | EML* | FC | SD | FDR | FC | SD | FDR | FC | SD | FDR | FC | SD | FDR | FC | SD | FDR | FC | SD | FDR |
| *Abcb11* | ↔ ↔ ↔ | 0.000 | 0.000 | 1.000 | 0.000 | 0.000 | 1.000 | 0.000 | 0.000 | 1.000 | 0.809 | 1.009 | 0.114 | 0.000 | 0.000 | 1.000 | 0.000 | 0.000 | 1.000 |
| *Abcb1a* | ↓ ↔ ↔ | -0.634 | 0.124 | **0.010** | -0.812 | 0.330 | **0.001** | -0.220 | 0.186 | 0.108 | 0.030 | 0.233 | 0.890 | 0.039 | 0.257 | 0.846 | -0.039 | 0.153 | 0.853 |
| *Abcb1b* | ↓ ↔ ↔ | -0.541 | 1.302 | 0.541 | -1.034 | 0.353 | **0.000** | -1.326 | 0.269 | **0.000** | 0.084 | 0.872 | 0.895 | -0.194 | 0.078 | 0.396 | -0.151 | 0.185 | 0.554 |
| *Abcc1* | ↔ ↔ ↔ | -0.114 | 0.204 | 0.555 | -0.285 | 0.056 | 0.068 | 0.022 | 0.154 | 0.845 | 0.192 | 0.171 | 0.292 | 0.282 | 0.277 | 0.055 | -0.004 | 0.054 | 0.963 |
| *Abcc2* | ↓ ↔ ↔ | -0.331 | 0.187 | 0.060 | -0.490 | 0.473 | 0.053 | -0.592 | 0.401 | **0.020** | -0.336 | 0.324 | 0.161 | -0.036 | 0.119 | 0.853 | 0.122 | 0.071 | 0.555 |
| *Abcc4* | ↑ ↑ ↑ | -0.227 | 0.319 | 0.432 | -0.034 | 0.126 | 0.888 | 0.399 | 0.220 | **0.016** | 1.060 | 0.059 | **0.006** | 0.692 | 0.190 | **0.001** | 0.281 | 0.347 | 0.360 |
| *Abcc5* | ↔ ↔ ↔ | -0.069 | 0.110 | 0.362 | -0.064 | 0.095 | 0.451 | 0.011 | 0.098 | 0.895 | 0.069 | 0.149 | 0.555 | -0.070 | 0.149 | 0.431 | -0.054 | 0.104 | 0.555 |
| *Abcc8* | ↓ ↓ ↓ | 0.167 | 0.189 | 0.188 | -0.154 | 0.109 | 0.188 | -0.713 | 0.280 | **0.044** | -0.761 | 0.293 | **0.011** | -0.495 | 0.227 | **0.002** | -0.343 | 0.432 | 0.212 |
| *Abcg2* | ↓ ↔ ↔ | -0.918 | 0.263 | **0.018** | -0.924 | 0.458 | 0.089 | -0.468 | 0.127 | **0.017** | -0.405 | 0.261 | 0.162 | 0.056 | 0.072 | 0.888 | -0.328 | 0.253 | 0.374 |
| *Edn1* | ↑ ↔ ↔ | 1.256 | 0.179 | **0.006** | 0.239 | 0.427 | 0.550 | 0.838 | 0.356 | **0.004** | 0.124 | 0.412 | 0.676 | 0.150 | 0.356 | 0.529 | -0.222 | 0.394 | 0.583 |
| *Gfap* | ↑ ↑ ↑ | 0.704 | 0.351 | **0.008** | 2.038 | 0.225 | **0.000** | 2.351 | 0.277 | **0.001** | 2.836 | 0.276 | **0.001** | 1.916 | 0.367 | **0.000** | 1.539 | 0.495 | **0.005** |
| *Hif1a* | ↑ ↑ ↑ | 0.407 | 0.168 | **0.003** | 1.024 | 0.160 | **0.000** | 0.939 | 0.248 | **0.000** | 0.536 | 0.481 | **0.040** | 0.171 | 0.097 | **0.006** | 0.284 | 0.561 | 0.374 |
| *Icam1* | ↑ ↑ ↑ | 3.335 | 0.556 | **0.000** | 2.783 | 0.308 | **0.001** | 3.168 | 0.992 | **0.020** | 3.004 | 0.398 | **0.000** | 1.884 | 0.518 | **0.000** | 1.587 | 1.268 | **0.049** |
| *Il6* | ↑ ↔ ↔ | 6.262 | 2.915 | **0.005** | 2.584 | 2.258 | **0.035** | 1.412 | 1.825 | 0.126 | 0.644 | 1.820 | 0.485 | 0.000 | 0.000 | 1.000 | 0.869 | 2.300 | 0.486 |
| *Nfe2l2* | ↑ ↑ ↑ | 0.836 | 0.275 | **0.001** | 0.896 | 0.202 | **0.000** | 0.773 | 0.217 | **0.001** | 1.798 | 0.437 | **0.000** | 1.148 | 0.285 | **0.000** | 0.892 | 0.628 | **0.026** |
| *Ngb* | ↓ ↔ ↓ | -0.123 | 0.457 | 0.784 | -0.840 | 0.408 | **0.004** | -1.210 | 0.497 | **0.005** | -0.617 | 0.708 | 0.361 | -0.102 | 0.545 | 0.734 | -0.759 | 0.258 | **0.016** |
| *Nr1i2* | ↔ ↔ ↑ | 0.181 | 0.363 | 0.667 | -0.076 | 0.512 | 0.895 | 0.327 | 0.300 | 0.485 | 0.036 | 0.382 | 0.899 | 0.853 | 0.461 | **0.006** | 0.039 | 0.619 | 0.895 |
| *Slc15a1* | ↔ ↔ ↔ | -0.278 | 0.369 | 0.360 | 0.224 | 0.784 | 0.727 | -0.037 | 0.212 | 0.856 | 0.199 | 0.280 | 0.537 | 0.213 | 0.141 | 0.476 | 0.163 | 0.125 | 0.555 |
| *Slc15a2* | ↓ ↔ ↑ | -0.279 | 0.077 | **0.004** | -1.486 | 0.187 | **0.000** | -1.954 | 0.592 | **0.000** | 0.430 | 0.373 | 0.061 | 0.543 | 0.287 | **0.004** | 0.469 | 0.050 | **0.032** |
| *Slc16a1* | ↓ ↑ ↑ | -0.037 | 0.050 | 0.853 | -0.539 | 0.169 | **0.000** | -0.351 | 0.204 | **0.008** | 0.692 | 0.196 | **0.035** | 0.619 | 0.122 | **0.003** | 0.594 | 0.516 | 0.066 |
| *Slc22a2* | ↔ ↔ ↔ | 0.245 | 0.359 | 0.555 | -0.208 | 0.389 | 0.555 | -0.054 | 0.069 | 0.895 | 0.772 | 1.003 | 0.467 | 0.362 | 0.750 | 0.608 | 0.189 | 0.728 | 0.818 |
| *Slc22a4* | ↑ ↑ ↑ | -0.034 | 0.552 | 0.931 | 1.044 | 0.131 | **0.017** | 1.338 | 0.483 | **0.001** | 2.320 | 0.493 | **0.000** | 1.863 | 0.231 | **0.000** | 1.787 | 0.244 | **0.003** |
| *Slc22a6* | ↓ ↔ ↔ | 0.227 | 0.362 | 0.485 | -4.589 | 2.116 | **0.002** | -3.646 | 2.216 | **0.008** | 0.102 | 0.626 | 0.895 | 0.164 | 1.545 | 0.856 | 0.991 | 0.844 | 0.413 |
| *Slc22a8* | ↓ ↔ ↔ | -0.337 | 0.111 | 0.254 | -1.680 | 0.644 | **0.001** | -1.442 | 0.440 | **0.000** | -0.420 | 0.371 | 0.341 | 0.392 | 0.750 | 0.369 | -0.322 | 0.255 | 0.220 |
| *Slc28a2* | ↔ ↑ ↔ | 0.091 | 0.180 | 0.764 | -0.536 | 0.349 | 0.084 | 0.664 | 0.801 | 0.122 | 1.502 | 1.011 | **0.013** | -0.247 | 0.429 | 0.374 | 0.169 | 0.290 | 0.537 |
| *Slc29a1* | ↓ ↔ ↔ | -0.702 | 0.490 | 0.137 | -0.793 | 0.358 | **0.002** | -0.411 | 0.263 | **0.032** | 0.137 | 0.386 | 0.550 | 0.514 | 0.179 | 0.128 | 0.517 | 0.253 | 0.112 |
| *Slc2a1* | ↑ ↔ ↑ | 0.457 | 0.139 | 0.076 | 0.614 | 0.186 | **0.001** | 0.664 | 0.394 | **0.007** | 0.546 | 0.428 | 0.055 | 0.331 | 0.091 | 0.060 | 0.438 | 0.326 | **0.049** |
| *Slc47a1* | ↔ ↔ ↔ | -0.608 | 1.464 | 0.728 | -0.856 | 1.437 | 0.597 | 1.379 | 0.939 | 0.413 | 2.092 | 1.511 | 0.188 | 0.039 | 2.154 | 0.963 | 1.382 | 1.150 | 0.369 |
| *Slc7a1* | ↔ ↔ ↔ | -0.132 | 0.213 | 0.287 | -0.192 | 0.202 | 0.360 | -0.173 | 0.104 | 0.374 | -0.189 | 0.324 | 0.537 | -0.163 | 0.265 | 0.540 | -0.257 | 0.096 | 0.233 |
| *Slc7a5* | ↑ ↑ ↑ | 0.327 | 0.185 | **0.021** | 0.613 | 0.244 | **0.001** | 0.694 | 0.152 | **0.000** | 0.426 | 0.104 | **0.020** | 0.582 | 0.092 | **0.004** | 0.375 | 0.178 | **0.009** |
| *Slco1a1* | ↔ ↔ ↔ | 0.058 | 0.494 | 0.939 | 0.469 | 0.869 | 0.291 | 0.796 | 1.123 | 0.161 | -0.593 | 1.384 | 0.567 | -0.367 | 0.893 | 0.748 | -0.368 | 0.240 | 0.669 |
| *Slco1a2* | ↓ ↔ ↔ | -0.988 | 0.193 | **0.009** | -1.540 | 0.380 | **0.000** | -0.775 | 0.472 | **0.016** | -0.189 | 0.431 | 0.550 | 0.105 | 0.350 | 0.592 | 0.108 | 0.140 | 0.529 |
| *Slco1a5* | ↔ ↔ ↔ | -1.362 | 1.864 | 0.413 | -0.076 | 0.421 | 0.853 | 0.319 | 0.415 | 0.430 | -0.155 | 0.115 | 0.112 | 1.715 | 2.717 | 0.220 | -0.399 | 0.330 | 0.380 |
| *Slco1b2* | ↔ ↔ ↔ | 0.000 | 0.000 | 1.000 | 0.000 | 0.000 | 1.000 | 0.121 | 0.341 | 0.485 | 0.021 | 0.060 | 0.485 | 0.000 | 0.000 | 1.000 | 0.000 | 0.000 | 1.000 |
| *Slco2b1* | ↓ ↑ ↑ | -0.610 | 0.398 | **0.021** | -0.884 | 0.207 | **0.000** | -0.594 | 0.189 | **0.002** | 0.688 | 0.200 | **0.001** | 0.968 | 0.387 | **0.001** | 0.741 | 0.514 | **0.023** |
| *Vim* | ↑ ↑ ↑ | 0.485 | 0.154 | **0.016** | 1.881 | 0.308 | **0.000** | 3.909 | 0.407 | **0.000** | 5.091 | 0.957 | **0.000** | 3.286 | 0.667 | **0.000** | 2.609 | 1.055 | **0.005** |

EML*: Early (3-24 hours), Mid (72 hours), Late (7-14 days); ↑: expression increased, ↓: expression decreased, ↔: expression did not change significantly
